# Expansion and re-classification of the extracytoplasmic function (ECF) σ factor family

**DOI:** 10.1101/2019.12.11.873521

**Authors:** Delia Casas-Pastor, Raphael Rene Müller, Anke Becker, Mark Buttner, Carol Gross, Thorsten Mascher, Alexander Goesmann, Georg Fritz

## Abstract

Extracytoplasmic function σ factors (ECFs) represent one of the major bacterial signal transduction mechanisms in terms of abundance, diversity and importance, particularly in mediating stress responses. Here, we performed a comprehensive phylogenetic analysis of this protein family by scrutinizing all proteins in the NCBI database. As result, we identified ∼10 ECFs per bacterial genome on average and classified them into 157 phylogenetic ECF groups that feature a conserved genetic neighborhood and a similar regulation mechanism. Our analysis expands the number of unique ECF sequences ∼50-fold relative to previous classification efforts, enriches many original ECF groups with previously unclassified proteins and identifies 22 entirely new ECF groups. The ECF groups are hierarchically related to each other and are further composed of subgroups with closely related sequences. This two-tiered classification allows for the accurate prediction of common promoter motifs and the inference of putative regulatory mechanisms across subgroups composing an ECF group. This comprehensive, high-resolution description of the phylogenetic distribution of the ECF family, together with the massive expansion of classified ECF sequences, enables the application of *in silico* tools for the prediction of important functional residues, and serves as a powerful hypothesis-generator to guide future research in the field.

## Introduction

Bacterial homeostasis is achieved through signal transduction mechanisms that connect the extracellular environment with the cytoplasm. Extracytoplasmic function σ factors (ECFs) are amongst the major signal transduction components in bacteria in terms of abundance and importance of the stress responses they mediate (1). As members of the σ^70^ family, ECFs guide the RNA polymerase (RNAP) to specific promoter sequences, and thereby enable bacteria to redirect gene expression in response to deteriorating environmental conditions (2, 3). Although ECFs are generally less prevalent than one-component systems (1CS) and two-component systems (2CS), previous studies revealed a large ECF abundance, with an average of six ECFs per bacterial genome (1), a large diversity, with more than 90 phylogenetic groups (1, 4–6), and a diverse range of activation mechanisms (7).

Members of the σ^70^ family are modular proteins composed of up to 4 core domains (σ_1-4_) that are classified into 4 groups. While group 1 represents the essential, full-length version of the σ^70^ family, σ^70^ groups 2-4, also known as alternative σ factors, represent truncations of this general structure that are usually non-essential (8, 9). Members of group 2 lack the σ_1.1_ region and members of group 3 lack the full σ_1_ domain (8). Compared to other members of the σ^70^ family, ECFs (or group 4 σ^70^s) are truly minimalistic regulators, since they only contain the σ_2_ and σ_4_ domains, essential for contact with RNAP and transcription initiation (8). The functions of both domains are well differentiated: while σ_4_ initiates the first step of promoter recognition by binding to the -35 element (10), σ_2_ is responsible for recognition and melting of the -10 element (11). Given that bacteria typically contain several alternative σ factors that compete for the binding to the RNAP core complex, it is key to regulate the activity of ECFs in response to changing conditions. The most common mode of regulation of ECFs is their sequestration by anti-σ factors (3). Anti-σ factors typically contain a cytoplasmic anti-σ domain (ASD) that sequesters the cognate ECF into an inactive conformation. Upon the onset of the inducing signal, anti-σ factors undergo conformational changes or get degraded, thereby releasing their cognate ECFs, which can then guide RNAP to their specific alternative promoter signatures to initiate target gene expression (2). While ECFs are often activated in response to external stimuli involved in cell envelope homeostasis and stress adaptation (8, 12, 13), some ECFs also detect cytoplasmic stimuli and regulate functions such as detoxification of reactive oxidative species, stationary phase survival (14, 15), metal resistance and homeostasis (12), morphological changes throughout the cell-cycle (1, 8) and virulence (16, 17). Besides the regulation via anti-σ factors, ECFs may also be regulated by various other mechanisms, such as conformational changes (18, 19), two-component systems (15, 20), C-terminal extensions (21, 22) and the proposed, but never proven, phosphorylation by serine/threonine kinases, many of which were first predicted by genomics approaches (see (7) for a review).

Research of the ECF family is heavily dependent on bioinformatic tools from the date of their first description (24). Indeed, ECFs are especially suitable for the application of prediction tools since they usually autoregulate their own expression, facilitating the identification of their target promoter, and because they are co-encoded with regulators of their activity and/or with genes regulated by the ECF. Accordingly, the first bioinformatic classification of the ECF σ factor family grouped ECFs from less than 400 genomes into 67 phylogenetic groups, based on sequence similarity, and revealed that conservation at protein level is often accompanied by conservation of the target promoter motif and a conserved genomic neighborhood (1). Altogether, this previous work proposed that it is possible to predict the target promoter of an ECF, its regulatory mechanism and its target genes from sequence information alone. Following studies expanded the number of phylogenetic groups by focusing on nine planctomycetal (4) and 100 actinobacterial (5) genomes, again identifying correlations between primary protein sequence and function.

Although these initial ECF classification studies helped to understand the large diversity of ECFs across the tree of life, they so far addressed ECFs from a limited number of genomes and/or focused on specific phyla (1, 4, 5). Based on the relatively sparse sequence basis, some of the initially defined ECF groups featured natural limitations in that they either defined groups with only very few (less than 10) proteins – so-called “minor” groups (ECF100, ECF102, etc.) (1) – or in that they clustered divergent sequences into groups that share only few unifying characteristics (e.g. ECF01, ECF10, ECF20) (1, 4). However, the explosion of annotated sequences in databases, not only from re-sequenced species but also from new species of underrepresented phyla, nowadays enables a more comprehensive view of the ECF family. Moreover, the availability of more ECF variants now allows the application of advanced bioinformatic tools, such as co-variation-based methods. One of the possible outcomes of these studies are detailed predictions of conserved protein-protein contacts (25) that may guide experimental analyses of the interplay between ECFs and they regulatory partners.

In the light of the above-mentioned limitations of the initial studies, a comprehensive and robust expansion of the ECF classification that reflects the massive increase in bacterial genomic sequence space is overdue. In this work we searched for ECF σ factors in all available genomes and metagenomes of the National Center for Biotechnology Information (NCBI) database, thereby expanding the number of identified ECF proteins 50-fold. We clustered the new ECFs into 2,380 subgroups with a high degree of sequence conservation. Subgroups were further grouped into 157 ECF groups according to genetic context conservation and their putative mode of regulation. As a result, we defined 22 novel ECF groups with no significant similarity to previously described ECF groups. Furthermore, 62 original groups were preserved and expanded with previously unclassified sequences. The conservation of the subgroups facilitated downstream *in silico* analyses such as prediction of conserved target promoter elements, conserved protein domains in the genetic neighborhood and putative anti-σ factors. Even though the large amount of information collected for each of the 157 ECF groups only allows us to focus on the most interesting findings, we provide an extensive compendium of all the information gathered for each group in the supplementary text. The possibilities offered by this hierarchical, comprehensive classification are illustrated by a recent study that predicts the functional role of C-terminal extensions in two ECF groups, only possible due to the expansion of the number of proteins assigned to these groups (26). This wealth of data represents a comprehensive resource to both computational and experimental researchers and helps guiding the characterization of ECF σ factors of unknown function.

## Results

### Rationale of this study

Previous ECF classification efforts (1, 4, 5) were based on 495 genomes and identified a total of 3,554 ECFs, which were classified into 94 ECF groups (summarized in (6)). Upon initiation of this work (February, 2017), the NCBI database contained 180,909 genomes, of which 4,106 were bacterial genomes tagged as “reference” or “representative”, discarding GenBank assemblies when a RefSeq assembly was available for the same genome (see https://www.ncbi.nlm.nih.gov/refseq/about/prokaryotes/). This indicates that the presently available number and diversity of ECF sequences should be much larger than previously observed. To expand the library of ECFs, we here performed a comprehensive ECF search in the NCBI database and re-classified the resulting protein sequences in a hierarchical approach. First, we clustered protein sequences into fine-grained ECF subgroups with a high degree of sequence similarity, and then aggregated subgroups into coarse-grained groups that share a common genetic neighborhood and a putative type of anti-σ factor. Whenever appropriate, the ECF group names were preserved from the original groups with which they share the same characteristics, e.g. groups ECF41 and ECF42. New ECF groups that are product of redistributions of original groups or that have no significant similarity to any of the original groups were designated with running numbers from 201 onwards, e.g. ECF201, ECF202, etc. Similarly, ECF subgroups were named by adding an ‘s’ to their group’s name and a running number indicating the rank in number of sequences contained in the subgroup, e.g. ECF201s1, ECF201s2, etc. (for more details see Methods). The similarity among the ECFs contained in groups allows the identification of common putative target promoter motifs and ECF regulators, as will be described in detail in the following sections. These hypotheses are confirmed whenever experimental reports on members of the group exist.

### The number of identified ECFs is 50-fold larger than in the founding ECF classification

The number of protein sequences in public databases has expanded extensively since the founding ECF classification efforts (1) and we expected a proportionate increase in the number of ECF sequences. To identify novel ECFs, we first extracted the sequences from all previous ECF classification efforts (1, 4, 5), aligned them and created a general Hidden Markov Model (HMM) for the ECF core region, including the linker between σ_2_ and σ_4_, but excluding any potential protein domains fused N- or C-terminally to the ECF (Fig. 1A). To discriminate ECFs from other σ factors, we first scored this generic ECF HMM against two sets of training sequences – true ECFs from the founding classification and a negative control set of σ factors from groups 1, 2 and 3 that additionally contain domains σ_3_ and σ_1_ in some cases. This allowed us to define a threshold score that maximizes true positive ECFs (Fig. 1B; *green*) while minimizing the number of false positive σ factors (Fig. 1B; *red*). We then selected the non-redundant protein sequences from the NCBI database (see Methods for selection of relevant genomes), for which the generic ECF HMM yielded scores higher than this threshold (Fig. 1C). As further quality controls, we filtered for sequences containing the Pfam domains σ_2_ and σ_4_ but lacking the σ_3_ domain, and discarded proteins with non-amino acidic characters, such as X or J. This resulted in a library of 177,910 non-redundant ECF sequences. Some of the candidate ECF σ factors included in this list cluster together with group 3 σ factors, indicating the presence of a cryptic σ_3_ domain, which prompted us to remove them from the list of ECF σ factors. This left us with 177,341 non-redundant ECFs, accounting for a ∼50-fold expansion over the number of ECFs in the founding ECF classification (1, 4, 5) (Fig. 1C). The full list of ECFs extracted during this study can be found in Table S1.

Next, we analyzed the taxonomic origin of this expanded ECF library to extract/determine the typical/average ECF numbers found in individual bacterial phyla. To enable such statistics, we focused on the subset of complete genomes of non-metagenomic origin, classified as NCBI “reference genomes” or “representative genomes”, thereby mitigating bias towards heavily sequenced species. Analysis of 12,539 ECFs extracted from 1,234 of these genomes showed that the taxonomic distribution of species became more diverse than in the original classification efforts (Fig. 2A; *Genomes*). In particular, the fraction of the three most abundant phyla – Proteobacteria, Actinobacteria and Firmicutes – was reduced from 86.9% in the original to 77.6% in the new classification. This reduction was accompanied by an increase in the number of species from underrepresented phyla, such as Bacteroidetes and Cyanobacteria (Fig. 2A; *Genomes*). In addition, 19 new ECF-containing phyla emerged (Table S3, *New phyla*). Yet, these 19 phyla have a limited contribution to the overall ECF database, given their low number of sequenced genomes. This difference in the taxonomic origin of the species included in original and new classifications naturally changes the taxonomic origin of ECFs gathered in each library. For instance, the fraction of ECFs from underrepresented genomes, such as Bacteroidetes and Planctomycetes, is larger in the new ECF library (Fig. 2A; *ECFs*). This is not the case for Cyanobacteria and Acidobacteria, which contribute a smaller percentage of ECFs than in the original library (Fig. 2A; *ECFs*). These differences in taxonomic composition in the ECF library are reflected in the average number of ECFs per genome, which increases from approx. seven ECFs per genome in the original ECF libraries (1, 4, 5) to about ten ECFs per genome in the new library (Fig. 2B). Confirming the findings of previous reports (1, 5), the number of ECFs per genome is directly proportional to genome size (Fig. S1), although the average number of ECFs per genome depends on the phyla of origin (Fig. 2B). Bacteroidetes and Actinobacteria have the greatest abundance of ECFs, with an average of 22.5 and 17.7 ECFs per genome, respectively (Fig. 2B). Phyla with a lower abundance of ECFs include Cyanobacteria and Spirochaetes, with an average of 2.7 and 3.7 ECFs per genome, respectively (Fig. 2B). Firmicutes and Proteobacteria contain an intermediate number of ECFs, 7.1 and 7.5, respectively (Fig. 2B). These differences might indicate different dependence on ECFs as signal-transduction system in different phyla, as previously noticed for Actinobacteria, which are particularly rich in ECFs, but also in 1CS and 2CS (5).

### ECF classification 2.0

The wealth of new proteins identified in the initial library expansion prompted us to reclassify ECF σ factors into groups with common characteristics. To this end, we first subjected the 177,910 protein sequences of the new ECF library to the rapid MMSeqs2 clustering algorithm (27), followed by a quality control that bisects the resulting clusters until the maximum pairwise k-tuple distance between sequences was ≤ 0.60 (Fig. 3A, see Methods for more details). Clusters with ≤ 10 proteins were discarded to ensure high sequence coherence within clusters, while preventing an explosion of small clusters with limited statistical relevance (Fig. 3A). This procedure yields a total of 2,380 ECF clusters (referred to as “subgroups”) with a median of 22 non-redundant proteins per subgroup (Fig. 3D). Subgroups capture 77.3% of the proteins, while 22.7% of the proteins remain unclassified, similar to the statistics in the original classification (1) (Table S1). Permutation tests on subgroups showed that the average k-tuple distance is significantly lower (two-tailed Student’s t-test; p-value < 1e-16) in our subgroups as compared to random clusters of the same size distribution, indicating that subgroups are well defined (Fig. 3F).

Then, we computed a phylogenetic tree based on the consensus sequence of each subgroup. This tree helps to identify the evolutionary relationship between the ECF subgroups (Fig. 3A, bottom; Fig. 4). As outgroups we included sequences with a low-scoring σ_3_ domain, as well as the consensus sequence of all σ_3_-containing proteins in Pfam, the latter of which we selected as root of the tree. Not surprisingly, proteins with a low-scoring σ_3_ domain clustered at the base of the tree (Fig. 4) and formed three groups with significant similarity to the sporulation σ factor SigF from Firmicutes and Actinobacteria, the flagellum biosynthesis σ factor FliA and the stationary phase σ factor SigH from *Bacillus* spp. Although they are not part of the ECF classification, these groups constitute the link between the group 3 and group 4 (ECF) σ factors (9) and account for the quality of our clustering approach. Other sequences with σ_3_ domain remained unclassified (0.18% of the unclassified sequences).

To identify subgroups with common characteristics, we performed an in-depth analysis of the genomic context of ECFs in each subgroup, and aggregated subgroups into a total of 157 ECF groups. For the definition of these ECF groups, the phylogenetic tree was manually split into monophyletic clades, unless clades shared a similar genetic context and putative anti-σ factor type (Fig. 3B and methods for more details). Genetic neighborhood and anti-σ factor type were evaluated in the subset of ECFs that come from “representative” or “reference” genomes, as defined by NCBI (https://www.ncbi.nlm.nih.gov), including only RefSeq genomes when both RefSeq and GenBank assemblies exist. This helps to mitigate the bias towards commonly sequenced bacteria. As a result, 76.0% of the ECFs were captured in groups, displaying a median group size of 243 non-redundant proteins (Fig. 3D). As an example, figure 3B shows a close-up view on 19 ECF subgroups within the ECF tree, together with the proteins in their genetic neighborhood that feature >50% domain architecture conservation (i.e., a combination of their Pfam domains). Here, it is evident that ECFs in subgroups ECF02s1, ECF02s2 and ECF02s3 share a conserved genomic context with the anti-σ factor RseA, and the regulators RseB and RseC for the former two subgroups, suggesting that ECFs in these subgroups feature the same mode of regulation as RpoE from *E. coli* (belonging to ECF02 subgroup 1). Likewise, the subgroups aggregated into group ECF32 display strong conservation with a 2CS and a large number of genes encoding a type III secretion system (T3SS) (Fig. 3B). The assignment of ECFs to ECF groups and subgroups can be found in table S1. These results underline the previous notion that ECFs with close phylogenetic distance often share a conserved genomic context, the gene products of which are typically involved in the regulation of ECF activity and/or direct transcriptional targets of the ECF (1). This not only provides the basis for the definition of an ECF group, but also helps to predict putative functions and regulatory mechanisms to ECF groups with no experimentally described members (Table S2).

To provide a systematic overview on the conserved genomic context in each ECF group, we analyzed the frequencies of genes with a conserved protein domain architecture encoded up- and downstream of the ECF (Fig. 3C). For group ECF02, for instance, this reveals that downstream of the regulators RseA-C there is enrichment of genes encoding translation regulators (e.g. EF-Tu), even though the specific position of individual genes is less conserved (Fig. 3C). However, despite the overall conservation of the genomic context within an ECF group, we often find subgroup-specific traits with respect to the positioning and the specific type of conserved genes (Supplementary Text; Table S2), clearly indicating that the definition of ECF subgroups is highly relevant to the biological function of an ECF.

In addition to the conserved genomic context, ECFs often auto-regulate the expression of their own genes, allowing bioinformatic prediction of their putative (sub)group-specific target promoters from conserved bi-partite DNA elements upstream of the ECF-encoding operon (1, 28). When applying a similar analysis to ECF subgroups and groups (Fig. 3B and C), we found overrepresented promoter motifs in many groups, e.g. ECF02, while others did not show significant motifs, e.g. ECF32, consistent with observations that the latter are not auto-regulated (20). Interestingly, even though predicted target promoter motifs were not used in the definition of the ECF groups, split points that define ECF groups (based on conserved genomic context) usually agree well with similar promoter elements (Fig. 3B). However, as for the conservation of the genomic context, we sometimes find subgroup-specific putative target promoters (e.g. in group ECF30 and others in supplementary text), highlighting the added value of the fine-grained clustering approach taken here.

The definition of ECF groups based on genomic context conservation is further supported by the statistical properties of the ECF subgroup tree, which typically displays high bootstrap support scores at the rooting branches of ECF groups (Fig. 3B and E), indicating that these groups are robust with respect to re-sampling of the original data set. To further check the performance of the new classification approach, we created profile Hidden Markov models (HMMs) from the conserved σ_2_ and σ_4_ domains of all sequences at the ECF group and subgroup level and tested whether these models were capable of faithfully classifying ECF sequences from their own groups (Fig. 3G and Fig. S2). This analysis showed that sequences were assigned to the correct ECF group in 99.3% of cases, while assignment to the correct subgroup was successful in 94% of the cases. The lower performance of subgroup-assignment was not surprising, given that neighboring ECF subgroups share higher sequence similarity than neighboring ECF groups. These results confirm that the definition of ECF groups and subgroups is based on a rational statistical approach and our findings suggest that the used HMMs are specific and sensitive to allow for the classification of novel ECF σ factors in the future.

### ECF classification 2.0 refines original and identifies novel ECF groups

As a proof of concept, we compared the original ECF classification and the classification presented in this work. To this end, we created HMMs from the original ECF groups and used them to classify the new ECFs gathered here. We saw that many of the sequences classified into a particular new group were also classified into a particular original group (Fig. 4, ring #1), indicating that there is a broad degree of correlation between the different classification approaches. Accordingly, for these groups of high coherence we maintained the original group names to label ECF groups presented in this work. Further in-depth analysis of the composition of the new groups revealed that 62 out of the 94 original groups are preserved, 21 are merged into larger groups, five remain mainly ungrouped, three are scattered across several subgroups, and three are present only in small percentages in some groups (Table 1 and Fig. S3).

One case of an extremely scattered original group is ECF01 (Fig. S3). This group was already considered highly diverse in the first ECF classification (1) and, based on the relatively unspecific HMM model of this group, it acquired more sequences in subsequent classification efforts (4, 5). As a result, we did not consider the proteins from ECF01 for the nomenclature of the ECF groups in this work. Another highly scattered original group is ECF20 (Fig. S3). ECF20 is present in four main groups of our classification: ECF281, ECF289, ECF290 and ECF291 (Table S2). ECF281, ECF290 and ECF291 seem to be related to heavy-metal stress, since their genetic neighborhoods contain a conserved heavy-metal resistance protein in position +2 downstream of the ECF-encoding sequence in ECF281 and ECF290, and the full operon of a metal efflux pump in ECF291. This function of ECF291 has been experimentally confirmed for CnrH in *Cupriavidus metallidurans* (ECF291s9) (29). Nevertheless, the anti-σ factors encoded in the genetic context of members of these groups differ. ECF281 features a zinc finger-containing anti-σ factor in position +1, while in the case of ECF289 this protein contains a DUF3520 domain fused to a von Willebrand factor; ECF290 contains a RskA-like anti-σ factor, and, lastly, ECF291 contains a CnrY-like anti-σ factor in position -2 (Supplementary Text, Table S2). Based on this anti-σ factor diversity, it seems likely that the cognate ECFs are regulated in response to different input stimuli, thereby warranting the definition of different ECF groups.

The last scattered group is ECF10. Even though minor parts of the original group ECF10 appear across the new ECF classification, only groups ECF239 and ECF240 receive most of the proteins of the original ECF10. Even though these two groups are both located within the large clade of FecI-like ECFs (Fig. 4), members of ECF239 do not contain genes with a conserved carbohydrate-binding domain in their neighborhood, a characteristic described for members of the original ECF10 (1). These proteins have been suggested to be involved in carbohydrate degradation. However, experimental confirmation for this biological function is still pending.

Even though scattered original groups are interesting, group merging events are more common. Its high occurrence is probably due to the incorporation of new sequences that bridge previously isolated ECF groups. Indeed, this possibility was considered in the founding ECF classification (1). One example of a merged group is ECF243, which constitutes the largest group of the new classification and contains the proteins previously associated to original FecI-like groups ECF05 to ECF09 (Fig. S3). The reasons for merging were 1) members of these original groups form a monophyletic clade in the ECF tree (Fig. 4) and 2) they contain a common genetic neighborhood with a FecI-like anti-σ factor typically in position +1 from their coding sequence and a TonB-dependent receptor in position +2 (Supplementary Text, Table S2). Another example of a merged group is ECF238, which contains sequences from the original groups ECF24 and ECF44 (Fig. S3, Table S2). Members of ECF238 contain a cysteine-rich C-terminal extension of approximately 20 amino acids (Supplementary text), which is likely required for the activation of members of ECF238 when the appropriate metal in the right redox state is present in the cytoplasm, as found for CorE2 from *Myxococcus xanthus* (ECF238s15) (30).

What is likely the most interesting contribution of the new classification are the entirely new groups. We found 22 new groups that could not be assigned to any original group (Table 2). Six of these groups contain less than ten proteins from representative/reference organisms and, therefore, were not further analyzed. From the analyzed groups, 10 share a conserved genetic neighborhood with putative anti-σ factors. A special case of these is ECF241, which is located in the FecI-like clade and represents an evolutionary intermediate between groups ECF240 (derived from original ECF10) and the iron uptake FecI-like group ECF242, although ECF241 shows no significant similarity to any original FecI-like group. Instead of the canonical FecR-like anti-σ factor from FecI-like groups, members of ECF241 contain a conserved two-transmembrane helix protein in position +1 or -1 from their coding sequence that in some cases hits the Pfam model for heavy-metal resistance proteins (Pfam: PF13801). Given the lack of an anti-σ factor in a group within the FecI-like clade, we further analyzed this conserved protein. Its N-terminus, the region that typically contains the ASD in anti-σ factors, is not long enough to feature a typical ASD. However, a multiple sequence alignment of this protein with the ASDs of canonical FecR-like anti-σ factors revealed that a putative, divergent ASD might be located in the C-terminal cytoplasmic part of the conserved protein. To our knowledge, this is the first time an anti-σ domain has been predicted C-terminally from transmembrane helices. The second most common regulators of ECF activity in these new ECF groups are C-terminal extensions (four out of 22), with groups ECF287 and ECF288, from Actinobacteria and Firmicutes, respectively, containing cysteine-rich C-terminal extensions, and group ECF294 with a SnoaL-like extension (Table S2, supplementary text). A potential regulator was not found for members of ECF201 and ECF282. In the case of ECF282, the regulation could be carried out by a novel mechanism that involves transcriptional regulation and ClpXP proteolysis, as explained below.

Taken together, the ECF groups presented in this work preserve many of the original groups, expanding them with more proteins, and splitting or merging them in some cases. Here, we described the new findings concerning the 22 new ECF groups with no significant similarity to any original group. However, a full overview of all the ECF groups and their occurrence in different bacterial phyla is shown in Fig. 5 and the full description of the groups is available in Supplementary Text.

### ECF σ factors feature diverse, often multi-layered, modes of regulation

Given the large diversity of the ECF σ factor family, it is essential to focus on individual groups in order to extract conclusions concerning their biological function, regulation and DNA binding site. Genetic neighborhoods of ECF σ factors typically contain an anti-σ factor with a single transmembrane helix, encoded in position +1 downstream of the ECF coding sequence. However, it is well known that other regulatory elements might be substituting it, ranging from fused C-terminal extensions, to two-component systems and serine/threonine protein kinases (1, 7, 23). Here, we provide an overview of the different modes of regulation present across the groups in the present ECF classification. A comprehensive description of all ECF groups can be found in Supplementary Text and in Table S2.

Most of the ECF groups (114 out of 157) contain a putative anti-σ factor, as defined by 1) the presence of Pfam domains of known anti-σ factors, 2) detectable similarity to anti-σ factors of the founding classification (1) and 3) presence of transmembrane helices (see Methods for details). This anti-σ factor is typically encoded in position +1 from the ECF coding sequence. A list of putative anti-σ factors identified in this study can be found in Table S4. In most of the cases, the putative anti-σ factor does not match any Pfam domain of experimentally addressed anti-σ factors. In order to decipher common types of anti-σ factors present across the ECF tree, we built HMMs from the cytoplasmic area (when it harbors a conserved region representing a putative ASD) of the extracted putative anti-σ factors that did not fit any Pfam domain. With these models, we searched the proteins encoded by 10 genes up- and downstream of the ECF coding sequence in all ECF groups (Fig. 6F). Interestingly, we found that most of the anti-σ factors are ECF group-specific, in agreement with previous experimental observations showing orthogonality between anti-σ factors of different groups (28). However, exceptions are the clade that contains ECF222, ECF51, ECF38 and ECF39, which share the same type of one-transmembrane helix anti-σ factor, groups regulated by FecR-like anti-σ factors, mostly located in the FecI-like clade of the ECF tree (ECF239, ECF240, ECF242 and ECF243), groups regulated by RskA-like anti-σ factors, anti-σ factors with a putative zinc-finger, and anti-σ factors with a DUF4179, almost exclusively present in Firmicutes (Fig. 6F). The number of transmembrane helices on putative anti-σ factors is usually one (82 ECF groups), followed by two (14 groups), six (five groups), four (three groups) and three (one group) (Fig. 6E). Soluble anti-σ factors, as defined by the absence of a predicted TM helix, are present in ten ECF groups (Fig. 6E). However, given that our analysis can only identify soluble anti-σ factors with detectable similarity to existing anti-σ factors, it is likely that other soluble anti-σ factor variants may exist. Additionally, even though we evaluated the similarity of new ECF σ factors to known ASDs, it is not guaranteed that all the new putative anti-σ factors function as such, given the vast diversity, lack of sequence conservation and lack of studies confirming their function.

ECF107 contains two putative anti-σ factors, which could be part of the same protein complex or compete for binding the ECF (6), thereby illustrating the complexity and diversity of anti-σ factor mediated regulation. A second example is ECF102, whose only described member, SigX from *P. aeruginosa*, has been suggested to have a role in mechanosensing (31). SigX is part of a seven-gene operon which includes a mechanosensitive ion channel (CmpX) encoded in position -1, a putative anti-σ factor (CfrX) encoded in position -2 and an outer membrane porin (OprF) encoded in position +1 (31). Even though original reports hypothesized that the regulation of SigX is carried out by the putative anti-σ factor CfrX (6), new reports suggest that its regulatory mechanism is more complex and involves also CmpX and OprF (31). We observed that these proteins are conserved in ECF102s1. Moreover, the mechanosensitive ion channel is conserved in subgroups 2 and 5, which indicates a similar regulation as members of subgroup 1. A similar case, in which proteins in addition to the anti-σ factor are required for ECF regulation, is ECF31. The only characterized member of ECF31, SigY from *B. subtilis* (subgroup 1), requires both the anti-σ factor YxlC, encoded in +1, and YxlD, encoded in +2, for its regulation, presumably forming a protein complex with the ECF (32). YxlCD homologs are conserved across ECF31.

The second most common regulatory mode of ECF σ factors is the presence of C-terminal protein extensions to the ECF (19 groups) (Fig. 6B), which is typically correlated with the lack of putative anti-σ factors (Fig. 6F). This agrees with the idea that the extension is substituting the anti-σ factor in the regulation of the ECF (33). ECFs with the same type of C-terminal extension cluster together in the same group, i.e., members of ECF42, with tetratricopeptide repeats in their extension, or in neighboring groups, i.e. members of ECF41, ECF56, ECF115, ECF294, and ECF295, with SnoaL-like C-terminal extensions. Given that only the core ECF domains were inputs of the ECF classification process, this supports the notion that the extension interacts with the core ECF regions in a unique manner depending on the type of domain that it bears (26). An interesting exception is ECF205, which also has a SnoaL-like extension but is located in proximity to the base of the ECF tree (Fig. 4), indicating that more factors, in addition to its C-terminal extension, determine the sequence conservation of this group. Aside from the Pfam domains identified in C-terminal extensions of the founding ECF classification (1, 4, 5), we identified a domain of unknown function (DUF1835) in ECF264, an extension with five or seven transmembrane helices in ECF263, and a CGxxGxGxCxC motif in ECF288.

Canonical C-terminal extensions are usually longer than 50aa, but we found that some groups contain short C-terminal extensions difficult to identify when only looking at protein length. These groups usually lack any other discernable means of regulation, which points towards the short extension as a modulator of ECF activity. One of these groups is ECF238, which merges the original groups ECF44 and ECF24. Members of ECF238 contain conserved cysteine residues in their short (∼20aa) C-terminal extension and also in the linker in some instances. One of the described members of ECF238, CorE2 from *Myxococcus xanthus* (subgroup 15), is known to be activated by Cd^2+^ and Zn^2+^ via this cysteine-rich C-terminal domain (30, 34). Another characterized member of ECF238 is SigZ from *Bacillus subtilis* (subgroup 9). SigZ is not regulated by any anti-σ factor and the studies about its function are very limited, since its deletion is not linked to any important phenotype (35). However, association of SigZ with group ECF238 suggests that the two cysteine residues present in its C-terminus could have a functional role. Another group with a short C-terminal extension is ECF29, which contains a conserved RCE/D motif in its ∼30 extra amino acids. Unfortunately, no member of ECF29 has been experimentally addressed, but the absence of a putative anti-σ factor similarly suggests a regulatory role of this short extension.

N-terminal extensions of the ECF core regions occur less often, they are generally shorter than canonical C-terminal extensions and they are prone to be overlooked whenever the ECF is translated from non-canonical start codons. The only well-described N-terminal extension appears in ECF121 (Fig. 6C). This extension has been studied in BldN (subgroup 1) from *Streptomyces coelicolor*, where it has to be proteolytically degraded to process the proprotein to its mature ECF, which then is subject to anti-σ factor regulation (36). Nonetheless, subgroups from several groups contain N-terminal extensions (Table S2). For instance, in ECF36s4, represented by SigC from *M. tuberculosis*, the N-terminal extension has been proposed to inhibit the DNA contact in the uninduced state, since members of this subgroup lack of an obvious anti-σ factor (37). Alternatively, the N-terminal extension of two members of ECF12s1, σ^R^ from *S. coelicolor* and SigH from *Mycobacterium smegmatis*, generates an unstable isoform produced from an earlier start codon upon exposure to thiol oxidants (38). This makes σ^R^ susceptible towards σ^R^-activated ClpP1/P2 proteases and thus implements a negative feedback loop that contributes to turning off the stress response (38).

Other putative regulators of σ factor activity that we often found in the conserved genetic neighborhood of ECFs were serine/threonine protein kinases (Fig. 6D). ECF σ factors of five original groups have been hypothesized to be directly phosphorylated by a protein kinase (ECF43 and ECF59-ECF62 (1, 4)). We add to the list of protein kinase-associated groups ECF217, ECF267 and ECF283 (Fig. 6D). Other groups such as ECF40, ECF27 or ECF210 contain protein kinases only in certain subgroups. Proteins from original group ECF60 were not classified by the pipeline since only eight members of ECF60 were extracted. Protein kinase-related ECF groups typically lack co-encoded anti-σ factors (Fig. 6D), consistent with the notion that direct phosphorylation of the σ factor regulates its ability to interact with RNAP and/or the promoter DNA. The only exception is group ECF267, which contains a putative FecR-like anti-σ factor with tetratricopeptide repeats in position +1. Given that ECFs from group ECF267 are very distant from the FecI-like clade (ECF239-ECF243) (Fig. 4), it seems possible that this anti-σ factor does not target members of ECF267, but other FecI-like ECFs. However, none of the organisms that contain members of ECF267 contain any FecI-like ECF σ factor. Whether the anti-σ factor and/or the protein kinase regulate the activity of members of ECF267 is unclear.

A total of four groups contain two-component systems in their genetic context. These regulators can co-occur in conjunction with anti-σ factors, as in the case of ECF15 and ECF246, or not, as in ECF32, ECF234 and subgroups 1, 2 and 3 of ECF39. These possibilities reflect the different regulatory mechanisms exerted by two-component systems. On one hand, members of ECF15, the main general stress response σ factors in alphaproteobacteria, are activated by a partner-switching mechanism where an anti-anti-σ factor sequesters the cognate anti-σ factor, thereby releasing the ECF (15). The anti-anti-σ factor is fused to the response regulator of a two-component system, also encoded in the same genetic context, and is able to bind the anti-σ factor when phosphorylated (15). This is unlikely the case for members of ECF246, since their response regulator is fused to a transcriptional regulator. Future analysis of members ECF246 could determine whether the putative anti-σ factor and/or the two-component system encoded in their genetic context regulates ECF activity. For members of ECF32, it was indeed shown that the two-component system indirectly regulates transcription of the ECF σ factor by inducing the expression of the transcription factors HrpSR (20, 40, 41). Members of ECF32 in turn activate the synthesis of the type III secretion system *hrp*, required for plant infection (20, 41). In the case of ECF39, 2CSs are directly regulating the transcription of the ECF, as described for SigE from *S. coelicolor* and σ^25^ from *Streptomyces avermitilis* (42, 43). Members of this group are involved in antibiotic synthesis and cell-wall stress resistance (42, 43). This direct regulation of the 2CS over the ECF expression could also occur in members of ECF234, given the absence of a putative anti-σ factor and the fusion of the response regulator to a transcriptional regulator. The physiological function of ECF234 seems to be related to an ABC transporter present in its genetic context, but the substrate of the transporter is unknown.

On top of these proteins, we found that some ECF groups contain conserved transcriptional regulators encoded in their genetic contexts, such as TetR-like repressors, which appear in groups with anti-σ factors (ECF125) and, remarkably, in ECF203, which lacks any obvious regulator (Fig. 6D). Given the lack of characterized members of ECF203, it is unclear whether this TetR repressor regulates the expression of members of ECF203 or is part of their response. In favor of the former, members of ECF203 do not seem to be auto-regulated, given the lack of a conserved (predicted) target promoter motif (Table S2). Other transcriptional regulators include LysR- and MerR-like repressors, which appear in several ECF groups associated with anti-σ factors (Fig. 6D).

A total of 16 ECF groups are not linked to any of the above-mentioned regulators (Fig. 7), inspiring us to predict novel, putative regulators of ECF activity. So far, only three of the 16 groups have experimentally addressed members, namely ECF228, ECF282 and ECF114. SigP from *Porphyromonas gingivalis* (ECF228s7) is only present in measurable concentrations when stabilized by direct interaction with the response regulator PorX from the two-component system PorXY (44). Even though response regulators are not conserved in the genetic context of members of ECF228 (Fig. 7), it is possible that members of ECF228 are unstable and require other proteins as chaperons. In the case of the novel group ECF282, σ^AntA^ from *Streptomyces albus* (subgroup 2) is regulated at the level of transcription and might be target of ClpXP proteolysis (45). Indeed, homologs of σ^AntA^ have been considered a new group of ECF σ factors that control the expression of antimycins (45). Even though the C-terminal AA dipeptide, suggested as target of ClpXP proteolysis (45), is only present in members of subgroup 2, members of other ECF282 subgroups could be regulated in a similar manner, since different ClpXP proteases have different binding specificities (46). In ECF114, SigH from *Porphyromonas gingivalis* (subgroup 4) plays a role in aerotolerance. SigH it is induced upon exposure to O_2_ and promotes oxidative stress protection and hemin uptake (47). Even though it is speculated that SigH is transcriptionally activated (47), no transcription factor in charge of this task has been identified.

Given that the genetic neighborhood of the remaining 13 groups does not feature canonical ECF regulators, but other conserved elements (Fig. 7), we speculated about their putative function. However, a general issue of this analysis is that it is hard to discriminate whether these elements are regulators and/or targets of ECF activity, suggesting that both options should be considered in downstream experimental analyses. Interestingly, we found new putative regulators/targets of regulation of the original groups ECF54 and ECF130. ECF54 is encoded in close proximity to a protein with a 4Fe-4S cluster, whereas ECF130 is encoded in proximity to a helix-turn-helix (HTH) containing protein (Fig. 7). Since HTH motifs are usually related to DNA binding, this protein could be a new type of transcriptional factor involved in transcriptional control of members of ECF130. A similar case is found in the new group ECF201, which is usually co-encoded with HTH proteins in position -1 (Fig. 7). Another interesting case is found in members of ECF237, which contain several “killing trait” proteins (Pfam: PF11757) in their vicinity (Fig. 7). These domains were described for RebB, one of the three proteins necessary for the assembly of R-bodies in the Paramecium endosymbiont *Caedibacter taeniospiralis* (48). Given the absence of conservation for the rest of the proteins from the R-body and the presence of several copies of the killer domain in members of ECF237 (which are mainly present in Bacteroidetes unrelated to the alphaproteobacterium *C. taeniospirails*), it is possible that proteins with this domain have an alternative function, not related to R-body assembly, but potentially involved in controlling ECF activity. Lastly, members of ECF286 and ECF292 share genetic neighborhood with several copies of Asp23 proteins (Pfam: PF03780) (Fig. 7). Asp23 is one of the most abundant proteins of *Staphylococcus aureus* and its deletion leads to upregulation of the cell wall stress response (49). Therefore, Asp23 proteins could be acting as a new type of anti-σ factor that regulate the activity of members of ECF286 and ECF292.

## Discussion

This work unifies, refines and greatly expands previous ECF classification efforts. Thanks to its two-tiered clustering approach, it provides a high-resolution view of the ECF family. ECF subgroups, composed of closely related proteins, are further hierarchically clustered into 157 ECF groups, defined on the basis of a common genetic neighborhood, which indicates a similar mode of regulation. As part of the *in silico* characterization of ECF groups, we predicted their putative regulators, their target promoter motifs and their most likely function (Supplementary Text 1 and Table S2). As already observed for the previous classification, these predictions are biologically meaningful in that they correctly reflect results of experimentally studied members, whenever available. The comprehensive description of the ECF groups serves as a source of testable hypotheses that will support the experimental description of new ECFs, which will lead, in turn, to more precise and detailed group descriptions.

Furthermore, the fine-grained resolution of the ECF family and the great enrichment in phylogenetically diverse proteins allows for the application of *in silico* prediction tools for individual groups. These types of analyses have led to the finding of functional differences between the C-terminal extensions from ECF41 and ECF42 (26), where C-termini from ECF41 had an inhibitory role over ECF activity, while ECF42’s were essential for transcriptional activation (26). These results were only possible due to the large number of sequences present in these groups, with >12,000 and >10,000 in ECF41 and ECF42, respectively (26).

The new ECF classification presented in this work has changes with respect to the founding classifications. Even though 62 of the 94 original groups are preserved, 21 are merged, five were ungrouped, and three each were scattered or present in the new classification but composing only small parts of their new group (Table 1). The new ECF groups are monophyletic clades of the ECF phylogenetic tree, that are subdivided into hierarchically-distributed ECF subgroups. This high-resolution, comprehensive classification provides advantages with respect to partial updates. One example comes from ECF54 and ECF58, identified in two different works (4, 5) and in two phyla, Actinobacteria and Planctomycetes, respectively. Within our ECF tree, these two groups are direct neighbors with a bootstrap support value of 17, indicating a significant protein similarity between them. None of them has a putative anti-σ factor or any other clear regulator, and they contain different elements in their genetic context (Fig. 7). These results suggest that ECF58 and ECF54 have the same origin, but they evolved independently in Actinobacteria and Planctomycetes, acquiring the regulation of different genes in their genetic neighborhood. What remains unclear is whether the regulation of members of ECF54 and ECF58 has common features, as expected for ECFs with a common origin.

As part of the description of ECF groups, we analyzed their most likely regulators and the types of putative anti-σ factors encoded in their genetic neighborhood (Fig. 6). Most of the predicted anti-σ factors are highly specific for their own groups (Fig. 6F). Exceptions occur in neighboring ECF groups, e.g. in the FecI-like clade (ECF239 to ECF243) or in the clade formed by groups ECF214, ECF18 and ECF19, indicating co-evolution between ECF and anti-σ factor sequences. The general lack of the same type of anti-σ factors in neighboring groups reflects their large diversity and their specificity, which has been exploited for the construction of orthogonal genetic circuits (28). Anti-σ factors are not the only genes conserved in the genetic context of ECF σ factors. In this study, we identified the ECF groups associated to other known ECF regulators such a C-terminal and N-terminal extensions, two-component systems, serine/threonine kinases (7), and other regulators such as TetR repressors (Fig. 6).

One important insight of this work is that ECF groups controlled by several regulatory layers are more common than originally thought. For instance, members of ECF121 are dually regulated by anti-σ factors and N-terminal extensions, some members of ECF12 are regulated by both anti-σ factors and alternative promoters that generate an unstable longer versions of the ECF σ factor (38) and members of ECF18 and ECF19 are not only regulated by RskA-like anti-σ factors, but also by a pair of conserved cysteine residues known to form a disulphide bridge that senses oxidative stress in SigK from *M. tuberculosis* (ECF19s1) (50). While these regulatory layers have only been deciphered for a few well-studied ECF σ factors, they might point towards a broader means of regulation also implemented in additional ECF groups. For instance, several ECF groups feature conserved cysteine residues potentially able to form disulphide bridges (Table S2), and members of ECF267 contain both a FecR-like anti-σ factor and a conserved protein kinase in their genomic neighborhood. Given their multi-layered regulation, abundance and diversity, we suggest that ECF σ factors have higher signal integration capabilities than previously anticipated.

With an average of approx. 10 ECFs per genome, these regulators are more abundant than previously thought (1). Confirming previous reports (1, 5, 51), we find that the number of ECFs is proportional to genome size (Fig. S1), with species thriving in diverse environments typically featuring larger genomes that provide them with the ability to sense and respond to a variety of external signals. One example is the bacterium *Sorangium cellulosum* So0157-2, which features a genome that is more than 1Mbp larger than its close relative *S. cellulosum* So ce56, allowing the former to adapt to alkaline conditions (51). Accordingly, the number of ECFs in *S. cellulosum* So0157-2 (82 ECFs) is significantly larger than in *S. cellulosum* So ce56 (70 ECFs), emphasizing the increased regulatory capacity incurred by genome expansion. Among the ECF groups acquired exclusively in *S. cellulosum* So0157- 2, we found ECF03 (one extra member), ECF26 (one extra member), ECF41 (two extra members) and ECF56 (one extra member). ECF03 and ECF26 are novel acquisitions present in *S. cellulosum* So0157-2 but not in *S. cellulosum* So ce56. Indeed, members of ECF03 are mainly present in Bacteroidetes (Table S2), and could have been acquired by horizontal gene transfer. However, this protein is not overexpressed under alkaline conditions (51), indicating that this ECF is either not autoregulated, or not responsible for alkaline resistance in *S. cellulosum* So0157-2. In contrast, the additional member of ECF26 contained in *S. cellulosum* So0157-2 is overexpressed at pH 10 (51) and could therefore be part of the alkaline resistance observed for *S. cellulosum* So0157-2. This ECF belongs to ECF26s1, which shares a conserved genetic neighborhood with a catalase (−1 from the ECF coding sequence) and a cytochrome b561 (position -2). Whether ECF26 or any other of these ECFs provides *S. cellulosum* So0157-2 with alkaline resistance needs further investigation.

The search of ECF σ factors presented in this work has some limitations related to the quality filters that we applied during the ECF retrieval. These filters limit the diversity of the extracted sequences, while ensuring that the collected proteins function as real ECF σ factors. In particular, we noticed that two main types of ECF σ factors could not be captured, namely, ECF σ factors from phages and ECFs whose conserved σ_2_ and σ_4_ domains are divergent. σ factors of phage origin have been described in literature (see review (52)); nevertheless, they are usually divergent from traditional σ factors (52), in some cases incorporating alternative domains replacing σ core domains. For instance, in *Bacillus* phage vB_BceM-HSE3, the ECF Gp17 contains a double zinc ribbon domain (Pfam: PF12773) in the position of the σ_2_ domain, while the generic Pfam domain for σ_4_ is not found at all (53). Similarly, the σ factors Gp01 and Gp103 contain only σ_2_ domain or no Pfam domain, respectively (53). Another reason for the lack of phage proteins in the present work is that viral genomes are usually not annotated in NCBI (54) and did not enter the ECF search in most of the cases. Other types of ECF-like σ factors not included in the current version are ECFs whose σ_4_ (e.g. SigI from *Bacillus subtilis*) or σ_2_ domain (such as VP0055 from *Vibrio parahaemolyticus* or ComX from *Streptococcus pneumoniae*) do not hit their Pfam models. A special example of this are σ^I^-like ECFs, which contain a σ_I-C_ domain instead of a canonical σ_4_ domain (55). These ECFs are involved in the synthesis of components of the cellulosome in cellulolytic clostridia (55). Attempts to classify these proteins against the current ECF classification were unsuccessful. The group with the highest probability of containing σ^I^-like ECFs is ECF201 (probability = 1.12e-19), the outermost group of the ECF classification, indicating that σ^I^-like ECFs are distant from canonical ECFs and might have evolved in parallel to them from group 3 σ^70^ factors. Lowering the stringency in our extraction pipeline would allow to study these non-canonical ECFs in further depth.

The average number of identified ECF σ factors per genome varies for different groups and bacterial phyla (Fig. 5). In this analysis it turned out that the number of unclassified ECFs per organism is larger in bacterial phyla underrepresented in biological databases (Fig. 5). In the future, clustering strategies could specifically target these proteins, which are likely too diverse and scarce to be clustered with the currently available dataset. We also found that some ECF groups are particularly enriched in certain phyla. For instance, we observed an average of 5.3 copies of ECF57 per planctomycetal genome, and an average of 3.4 copies of ECF240 per Bacteroidetes genome (Fig. 5). In these cases, a question that remains unsolved is whether the function of members of the same group is redundant in the same organism, or they rather hold specialized functions. In favor of the latter, members of ECF240, which inherits most of its characteristics from the original group ECF10, have been associated to carbohydrate scavenging in Bacteroidetes (1). Even though no member has been experimentally addressed to date, it is possible that the different members of ECF240 present in the same genome are involved in the uptake of different carbohydrates. A similar case occurs in the proteobacterial group ECF243, which merges original FecI-like groups ECF05-09, and is in charge of iron uptake (1, 56). We found an average of 1.13 members of ECF243 per proteobacterial genome. However, under closer inspection only 33% of the proteobacterial genomes contain members of ECF243, indicating that, when present, members of ECF243s are duplicated and appear in 3.4 proteins per organism on average. Interestingly, only 8.9% of the organisms contain ECF243s from the same subgroup, suggesting that different subgroups fulfill different physiological functions. One possibility is that members of different subgroups detect signals from different FecR-like anti-σ factors, which in turn, detect the presence of iron-siderophore complexes from different FecA transporters (see (56) for a review). Future analyses have to answer whether the different members of the same ECF group in the same genome have acquired different specificities and whether this specificity is a general feature of ECF σ factors.

In summary, the updated ECF classification presented in this work serves as a detailed source of testable hypotheses to guide the experimental characterization of this important class of bacterial regulators. The ECF classification comes together with a full description of ECF groups, including the putative group-specific regulators of ECF activity, conserved proteins encoded in the same genetic neighborhood, and predicted target promoter motifs (Supplementary Text and Table S2). Collectively, this information allows prediction of the potential function of the members of the group, which is verified by experimentally described members whenever they are available (Supplementary Text). Moreover, our hierarchical two-level classification provides a broad sequence collection with an appropriate degree of similarity (or variability) required for *in silico* prediction tools that employ sequence variation-based algorithms, such as co-variation-based prediction of protein-protein interactions or structural predictions.

## Methods

### General bioinformatic tools

The data was processed with custom scripts written in Python when nothing else is stated (available upon request from the corresponding author). Multiple Sequence Alignments (MSAs) were generated by Clustal Omega 1.2.3 (57) and were visualized in CLC Main Workbench 7.7.2 (QUIAGEN®). HMMER suite 3.1b2 (58) was utilized for generating and employing Hidden Markov Models (HMMs). HMMs were produced by hmmbuild (58) and proteins were searched against HMMs using hmmscan. The maximum-likelihood phylogenetic tree of ECF subgroups and bootstrap values were retrieved by IQ-TREE 1.5.5 with default options and automatic model selection (59). Phylogenetic trees were visualized using iTOL (60). Transmembrane helix predictions were carried out with TopCons (61) and PRED_TMBB2 for outer membrane proteins (62).

### ECF retrieval

The full retrieval process is depicted in Figure 1. The sequence between the start of σ_2_ and the end of σ_4_ domains (core ECF region) was extracted from the MSA of the 3,755 ECFs from the original library (1, 4, 5) (position 912 and 2059 in the MSA) and was used to build the general HMM for ECF sequence retrieval. Then, ECFs from the original library and proteins with a σ_3_ domain (Pfam: PF04539) retrieved from Pfam (release 31.0 (63)) were scored against this model. The resulting bit-scores were used to construct a Receiver Operator Characteristic (ROC) curve (using the function “roc_curve” from the “scikit-learn” Python package (64)) in order to select the threshold score able to capture the greatest number of ECFs (highest sensitivity), while minimizing the number of σ_3_-containing proteins (highest specificity). This resulted in an optimum bit-score threshold of 60.8, a sensitivity of 0.92, a specificity of 0.98 and an Area Under the Curve (AUC) of 0.98, accounting for a robust performance of the classifier. Then, proteins with a score higher than the optimum threshold were selected as putative ECF σ factors for the next steps. For this search, we considered all protein sequences from genomes with annotation (.gff), protein (.faa) and genome (.fna) file available in NCBI (as of February 2017), including RefSeq and GenBank entries, accounting for a total of 156,241 genomes and 554,108,437 proteins. Subsequent quality controls were applied to ensure that all the putative ECFs contain σ_2_ (Pfam: PF04542) and σ_4_ (Pfam: PF04545 and PF08281) domains and lack the σ_3_ domain using Pfam HMM profiles (hmmsearch with default settings). Only the non-redundant proteins without ambiguous amino acids were considered further. Sequence redundancy was assessed with Cd-hit (65) at 100% sequence identity. In summary, from the 554,108,437 annotated proteins, 714,848 had a positive score against the ECF model. Of those, 177,910 had σ_2_ and σ_4_ domains, lacked σ_3_, lacked non-amino acidic symbols and were non-redundant, constituting the extended ECF library used in the next steps.

Within this dataset we found that 1,217 proteins generated low-scoring hits against the model of the σ_3_ domain (Pfam: PF04539) when hmmscan was applied for σ_3_’s HMM alone. This is likely due to the size of the ECF database, where E-values are less significant than for smaller datasets (see HMMER documentation (58)). Given that the Pfam HMM of σ_3_ wrongly hits the σ_2_ or σ_4_ domain in some cases, only the proteins where the highest scoring σ_2_, σ_3_ and σ_4_ domains are not overlapping are considered as σ_3_-containing proteins. After their identification, the 1,217 σ_3_-containing proteins were used as outliers for the clustering, since they are the closest σ factors to ECFs.

The average number of ECFs per genome was computed considering only the 12,539 ECFs from the 1,234 complete genomes tagged as ‘representative’ or ‘reference’ in NCBI, giving priority to RefSeq genomes over GenBank if both exist for the same organism, unless stated otherwise.

### Clustering and ECF group definition

Non-redundant ECF sequences were stripped to σ_2_ and σ_4_ regions using hmmscan (HMMER suite 3.1b2 (58)) and Pfam models for σ_2_ and σ_4_. We selected the ‘envelope’ region of the hit with the lowest (i.e. best) E-value. For the first step of clustering, we applied MMseqs2 (27) with default parameters. However, the phylogenetic distance between pairs of sequences within clusters, as calculated from the k-tuple distance (66) obtained from Clustal Omega 1.2.3. with options ‘--full, --full-iter and --distmat-out’ (57), was large in some cases (Fig. S4), indicating imperfect clustering at this stage. To solve this issue, MMseqs2 clusters were split using a bisecting k-Means algorithm until the maximum pairwise distance between sequences of the clusters was ≤ 0.6. This threshold was chosen since it was the largest maximum k-tuple distance whose associated clusters contained proteins similar enough to produce homogeneous MSAs. The resulting 2,380 clusters with more than 10 sequences are defined as ECF subgroups. A total of 137,452 ECFs (77.26%) were classified into subgroups. Then, the consensus sequences of subgroups (computed with Biopython (67)) were used for the construction of a phylogenetic tree. As outliers of the tree we included the consensus of a MSA calculated from all proteins included in Pfam (release 31.0 (63)) that contain all of the three domains σ_2_, σ_3_ and σ_4_. We also included the 1,217 closest proteins to ECF σ factors with σ_3_ domain. The subsequent phylogenetic tree was manually split into monophyletic clades using a divisive strategy, where two clades were kept together in the same ECF group unless the genetic context or the putative anti-σ factor (when present) differed. ECF groups featuring only a single subgroup were only assigned if they have significant similarity to an original ECF group, while all other single subgroups were maintained as singleton subgroups (for nomenclature see next paragraph). This strategy resulted in 157 ECF groups that contained 135,259 ECF σ factors, corresponding to 76.03% of the new ECF library. For a graphical summary of the clustering strategy, see Fig. 3.

### Nomenclature

Names of original ECF groups are maintained for groups with the same characteristics. When several original groups are represented in an ECF group or the ECF group has no significant similarity to any original group, the name of this ECF group follows the pattern ECF2XX, standing for ECF classification 2.0, where XX is a running number assigned according to the position in the phylogenetic tree (Fig. 4). For instance, ECF201 is closer to the base of the tree than ECF260. Subgroups are referred with the name of the ECF group they are part of, followed by ‘s’, standing for <u>s</u>ubgroup, followed by a running number that increases for decreasing subgroup size. For instance, subgroups ECF02s1 (ECF02 subgroup 1) and ECF02s2 (ECF02 subgroup 2) are both part of group ECF02, and s1 contains more non-redundant proteins than s2. Subgroups that are not part of any ECF group are named ‘ECFs’ followed by a running number according to their position in the phylogenetic tree.

### Clustering validation

Groups and subgroups were evaluated according to the performance of their HMMs, built from the concatenated σ_2_ and σ_4_ domains of the constituting protein sequences. These HMMs were used to score proteins from all groups or subgroups. Bit-scores below the reporting threshold were considered to be equal to 0. The average score of members of each group against each HMM was normalized by dividing by the score of the group against its own HMM. The average normalized scores are plotted in a heatmap (Fig. S2A). We validated the subgroups by generating 100 randomly permutated sets of proteins with the same size distribution as the subgroups (Fig. 3F). The mean average k-tuple distance in the permutated data was 0.79±0.01, whereas this value was 0.29±0.11 for ECF subgroups. The difference between the distributions of average pairwise k-tuple distances of ECF subgroups and permutated clusters was statistically significant (two-tailed Student’s t-test p-value < 1e-16). Furthermore, we evaluated the support of the branches of the phylogenetic tree by running 100 bootstrap replicates, as implemented in IQ-TREE (59). As a further plausibility test, we verified the agreement between original and new classification (Fig. 4).

### Classification of proteins against ECF clusters

In order to classify proteins against original groups and new groups/subgroups, we derived two bit-score cut-offs for each group/subgroup: (i) The trusted cut-off is defined as the minimum bit-score achieved by a true member of a group/subgroup, whereas (ii) the noise cut-off is the maximum bit-score achieved by all ECFs that are non-members of this group/subgroup. Trusted and noise cut-offs provide the first step in the evaluation of the membership of a protein against a group/subgroup. Then, we obtained the probability that a protein belongs to a cluster using a logistic fit, as described in (68). First, we represented in the x-axis the bit-score and in the y-axis the probability that a bit-score is produced by a real member of the group/subgroup under evaluation, this is 1 for members and 0 for non-members. We fitted the data to a logistic regression describing the transition of the bit scores from non-members to members of a cluster (Fig. S5). The logistic function describing the probability *P*(*q, i*) of protein sequence *q* belonging to group/subgroup *i* reads

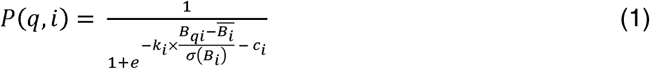

with *B*_*qi*_ being the bit-score obtained for the protein sequence *q* scored against the HMM model of group/subgroup *i* and *B*_*i*_ and *σ*(*B*_*i*_) being the mean and standard deviation of all bit-scores obtained for member sequences of group/subgroup *i*, respectively. The parameters *k*_*i*_ and *c*_*i*_ were fitted using the function ‘curve_fit’ (with a Levenberg-Marquardt algorithm) from the Python package ‘scipy.optimise’. These parameter values are available for groups, subgroups and original groups (Table S5). Some small subgroups could not to be fitted. In that case, the probability is considered 1 if the protein has a bit score larger that the trusted cut-off and it does not match any other subgroup, or 0 otherwise. The selection of a unique probability threshold was carried out with a ROC curve, using the members of the group/subgroup as true positives and members of all other groups/subgroups as true negatives. This probability threshold, together with trusted and noise cut-offs, and fitting parameters of the logistic curve are available for groups, subgroups and original groups (Table S5).

In the classification process itself only protein sequences stripped to their σ_2_ and σ_4_ domains were used. The regions of these domains were defined by the HMMER envelope region of the highest scoring hit of their respective Pfam models. In a first step of the classification only the groups/subgroups against which the bit-score of a protein is higher than the trusted cut-off (or the noise cut-off if none exceeds the trusted cut-off) are considered as putative groups/subgroups. In a second step, the protein is assigned to the cluster with the highest probability, calculated with the logistic curve of each cluster, only if this probability is higher than the ROC-optimized probability threshold.

### Promoter motif prediction

Since ECFs often auto-regulate their own transcription, their putative target promoters can be predicted from the regions upstream of the operon in which the ECF is encoded. To this end we searched for conserved bipartite nucleotide motifs in the 200 base pairs (bp) upstream of the ECF operon, by executing BioProspector (69) with the parameters ‘-W 7 -w 5 -G 18 -g 15 -n 150 -d 1’, as previously described (28). Operons were considered to be the set of coding sequences transcribed in the same direction as the ECF and with an intergenic distance shorter than 50 bp. Only the highest scoring motif of each input sequences was further considered. The region containing the -35 and -10 motifs in addition to ±10 bp up- and downstream is represented in a sequence logo produced by WebLogo 3 (70).

### ECF group analysis

For the analysis of ECF group characteristics we included only proteins from ‘representative’ and ‘reference’ genomes as defined by NCBI (https://www.ncbi.nlm.nih.gov), thereby reducing the bias towards frequently sequenced organisms. Only RefSeq assemblies were considered when both RefSeq and GenBank assemblies are available. To define coherent ECF groups and to elucidate the putative function of members of each group, we analyzed the protein domain composition of the proteins encoded in a distance of ±10 coding sequences from the ECFs coding sequence. First, we queried these proteins against the HMMs of Pfam 31.0 (63). For every protein, we only considered the non-overlapping Pfam domains with the lowest (best) E-value, leading to a set of specific domains (which we define as the ‘domain architecture’) for each protein in the genomic neighborhood of the ECF. For each ECF subgroup we analyzed the conservation of the domain architecture in specific positions up- and downstream of the ECF. A domain architecture is defined as conserved if it appears in more than 75% of the genomic contexts of an ECF subgroup. To avoid biases due to low number of ECFs, we only analyzed the genetic context of subgroups with more than 10 ECFs.

For anti-σ factor identification we used (i) Pfam domains of known anti-σ factors, (ii) detectable sequence similarity to anti-σ factors of the founding classification (1) and (iii) presence of transmembrane helices, as described in the following. Most of the anti-σ factors cannot be predicted due to the lack of Pfam domains that describe them. Therefore, we used the anti-σ factors retrieved by (1) as the database to query candidate anti-σ factors using BLAST (71) with an E-value < 0.01. Moreover, since anti-σ factors are usually transmembrane proteins, we predicted the presence of transmembrane helices using the consensus prediction of TopCons (61). In cases where the presence of the transmembrane helix was not clear, we aligned the sequences of the putative anti-σ factors to determine the presence of conserved hydrophobic regions likely to be transmembrane domains. Since anti-σ factors are usually located in positions ±2 from the ECF coding sequence, those were the main positions we focused our search on.

## References

1. Staroń, A., Sofia, H.J., Dietrich, S., Ulrich, L.E., Liesegang, H. and Mascher, T. (2009) The third pillar of bacterial signal transduction: classification of the extracytoplasmic function (ECF) σ factor protein family. Mol. Microbiol., 74, 557–581.

2. Helmann, J.D. (2002) The extracytoplasmic function (ECF) sigma factors. Adv. Microb. Physiol., 10.1016/S0065-2911(02)46002-X.

3. Paget, M.S.B. and Helmann, J.D. (2003) The σ70 family of sigma factors. Genome Biol., 10.1186/gb-2003-4-1-203.

4. Jogler, C., Waldmann, J., Huang, X., Jogler, M., Glöckner, F.O., Mascher, T. and Kolter, R. (2012) Identification of proteins likely to be involved in morphogenesis, cell division, and signal transduction in Planctomycetes by comparative genomics. J. Bacteriol., 194, 6419–30.

5. Huang, X., Pinto, D., Fritz, G. and Mascher, T. (2015) Environmental Sensing in Actinobacteria: a Comprehensive Survey on the Signaling Capacity of This Phylum. J. Bacteriol., 197, 2517–2535.

6. Pinto, D. and Mascher, T. (2016) The ECF Classification: A Phylogenetic Reflection of the Regulatory Diversity in the Extracytoplasmic Function σ Factor Protein Family. In Stress and Environmental Regulation of Gene Expression and Adaptation in Bacteria.

7. Mascher, T. (2013) Signaling diversity and evolution of extracytoplasmic function (ECF) σ factors. Curr. Opin. Microbiol., 16, 148–155.

8. Paget, M.S. (2015) Bacterial Sigma Factors and Anti-Sigma Factors: Structure, Function and Distribution. Biomolecules, 5, 1245–65.

9. Lonetto, M., Gribskov, M. and Gross, C.A. (1992) The sigma 70 family: sequence conservation and evolutionary relationships. J. Bacteriol., 174, 3843–9.

10. Lane, W.J. and Darst, S.A. (2006) The structural basis for promoter -35 element recognition by the group IV sigma factors. PLoS Biol., 4, e269.

11. Campagne, S., Marsh, M.E., Capitani, G., Vorholt, J.A. and Allain, F.H.T. (2014) Structural basis for - 10 promoter element melting by environmentally induced sigma factors. Nat. Struct. Mol. Biol., 10.1038/nsmb.2777.

12. Grosse, C., Friedrich, S. and Nies, D.H. (2007) Contribution of extracytoplasmic function sigma factors to transition metal homeostasis in Cupriavidus metallidurans strain CH34. J. Mol. Microbiol. Biotechnol., 12, 227–40.

13. Mascher, T., Hachmann, A.B. and Helmann, J.D. (2007) Regulatory overlap and functional redundancy among Bacillus subtilis extracytoplasmic function σ factors. J. Bacteriol., 10.1128/JB.00904-07.

14. Staroń, A. and Mascher, T. (2010) General stress response in a-proteobacteria: PhyR and beyond. Mol. Microbiol., 10.1111/j.1365-2958.2010.07336.x.

15. Francez-Charlot, A., Frunzke, J., Reichen, C., Ebneter, J.Z., Gourion, B. and Vorholt, J.A. (2009) Sigma factor mimicry involved in regulation of general stress response. Proc. Natl. Acad. Sci., 10.1073/pnas.0810291106.

16. Llamas, M.A., Van Der Sar, A., Chu, B.C.H., Sparrius, M., Vogel, H.J. and Bitter, W. (2009) A novel extracytoplasmic function (ECF) sigma factor regulates virulence in Pseudomonas aeruginosa. PLoS Pathog., 10.1371/journal.ppat.1000572.

17. Kazmierczak, M.J., Wiedmann, M. and Boor, K.J. (2005) Alternative Sigma Factors and Their Roles in Bacterial Virulence. Microbiol. Mol. Biol. Rev., 10.1128/mmbr.69.4.527-543.2005.

18. Campbell, E.A., Greenwell, R., Anthony, J.R., Wang, S., Lim, L., Das, K., Sofia, H.J., Donohue, T.J. and Darst, S.A. (2007) A conserved structural module regulates transcriptional responses to diverse stress signals in bacteria. Mol. Cell, 27, 793–805.

19. Li, W., Stevenson, C.E.M., Burton, N., Jakimowicz, P., Paget, M.S.B., Buttner, M.J., Lawson, D.M. and Kleanthous, C. (2002) Identification and structure of the anti-sigma factor-binding domain of the disulphide-stress regulated sigma factor σR from Streptomyces coelicolor. J. Mol. Biol., 10.1016/S0022-2836(02)00948-8.

20. Nizan-Koren, R., Manulis, S., Mor, H., Iraki, N.M. and Barash, I. (2003) The Regulatory Cascade That Activates the Hrp Regulon in Erwinia herbicola pv. gypsophilae. Mol. Plant-Microbe Interact., 10.1094/mpmi.2003.16.3.249.

21. Wecke, T., Halang, P., Staroń, A., Dufour, Y.S., Donohue, T.J. and Mascher, T. (2012) Extracytoplasmic function σ factors of the widely distributed group ECF41 contain a fused regulatory domain. Microbiologyopen, 10.1002/mbo3.22.

22. Liu, Q., Pinto, D. and Mascher, T. (2018) Characterization of the Widely Distributed Novel ECF42 Group of Extracytoplasmic Function σ Factors in Streptomyces venezuelae. J. Bacteriol., 10.1128/jb.00437-18.

23. Chandrashekar Iyer, S., Casas-Pastor, D., Kraus, D., Mann, P., Schirner, K., Glatter, T., Fritz, G. and Ringgaard, S. (2019) Transcriptional regulation by σ factor phosphorylation in bacteria. Nat. Microbiol.

24. Lonetto, M.A., Brown, K.L., Rudd, K.E. and Buttner, M.J. (1994) Analysis of the Streptomyces coelicolor sigE gene reveals the existence of a subfamily of eubacterial RNA polymerase sigma factors involved in the regulation of extracytoplasmic functions. Proc. Natl. Acad. Sci., 91, 7573–7577.

25. Weigt, M., White, R.A., Szurmant, H., Hoch, J.A. and Hwa, T. (2009) Identification of direct residue contacts in protein-protein interaction by message passing. Proc. Natl. Acad. Sci. U. S. A., 10.1073/pnas.0805923106.

26. Wu, H., Liu, Q., Casas□Pastor, D., Dürr, F., Mascher, T. and Fritz, G. (2019) The role of C□terminal extensions in controlling ECF σ factor activity in the widely conserved groups ECF 41 and ECF 42. Mol. Microbiol., 10.1111/mmi.14261.

27. Steinegger, M. and Söding, J. (2017) MMseqs2 enables sensitive protein sequence searching for the analysis of massive data sets. Nat. Biotechnol., 35, 1026–1028.

28. Rhodius, V.A., Segall-Shapiro, T.H., Sharon, B.D., Ghodasara, A., Orlova, E., Tabakh, H., Burkhardt, D.H., Clancy, K., Peterson, T.C., Gross, C.A., et al. (2013) Design of orthogonal genetic switches based on a crosstalk map of σs, anti-σs, and promoters. Mol. Syst. Biol., 10.1038/msb.2013.58.

29. Grass, G., Fricke, B. and Nies, D.H. (2005) Control of expression of a periplasmic nickel efflux pump by periplasmic nickel concentrations. In BioMetals.

30. Marcos-Torres, F.J., Perez, J., Gomez-Santos, N., Moraleda-Munoz, A. and Munoz-Dorado, J. (2016) In depth analysis of the mechanism of action of metal-dependent sigma factors: Characterization of CorE2 from Myxococcus xanthus. Nucleic Acids Res., 10.1093/nar/gkw150.

31. Chevalier, S., Bouffartigues, E., Bazire, A., Tahrioui, A., Duchesne, R., Tortuel, D., Maillot, O., Clamens, T., Orange, N., Feuilloley, M.G.J., et al. (2019) Extracytoplasmic function sigma factors in Pseudomonas aeruginosa. Biochim. Biophys. acta. Gene Regul. Mech., 1862, 706–721.

32. Yoshimura, M., Asai, K., Sadaie, Y. and Yoshikawa, H. (2004) Interaction of Bacillus subtilis extracytoplasmic function (ECF) sigma factors with the N-terminal regions of their potential anti-sigma factors. Microbiology, 150, 591–9.

33. Pinto, D., Liu, Q. and Mascher, T. (2019) ECF σ factors with regulatory extensions: The one□component systems of the σ universe. Mol. Microbiol., 10.1111/mmi.14323.

34. Pérez, J., Muñoz-Dorado, J. and Moraleda-Muñoz, A. (2018) The complex global response to copper in the multicellular bacterium Myxococcus xanthus. Metallomics, 10, 876–886.

35. Luo, Y., Asai, K., Sadaie, Y. and Helmann, J.D. (2010) Transcriptomic and phenotypic characterization of a Bacillus subtilis strain without extracytoplasmic function σ factors. J. Bacteriol., 192, 5736–45.

36. Bibb, M.J. and Buttner, M.J. (2003) The streptomyces coelicolor developmental transcription factor σBldN is synthesized as a proprotein. J. Bacteriol., 10.1128/JB.185.7.2338-2345.2003.

37. Thakur, K.G., Joshi, A.M. and Gopal, B. (2007) Structural and biophysical studies on two promoter recognition domains of the extra-cytoplasmic function σ factor σC from Mycobacterium tuberculosis. J. Biol. Chem., 10.1074/jbc.M606283200.

38. Kim, M.S., Hahn, M.Y., Cho, Y., Cho, S.N. and Roe, J.H. (2009) Positive and negative feedback regulatory loops of thiol-oxidative stress response mediated by an unstable isoform of σR in actinomycetes. Mol. Microbiol., 10.1111/j.1365-2958.2009.06824.x.

39. Bayer-Santos, E., Lima, L.D.P., Ceseti, L. de M., Ratagami, C.Y., de Santana, E.S., da Silva, A.M., Farah, C.S. and Alvarez-Martinez, C.E. (2018) Xanthomonas citri T6SS mediates resistance to Dictyostelium predation and is regulated by an ECF σ factor and cognate Ser/Thr kinase. Environ. Microbiol., 20, 1562–1575.

40. Lan, L., Deng, X., Zhou, J. and Tang, X. (2006) Genome-wide gene expression analysis of Pseudomonas syringae pv. tomato DC3000 reveals overlapping and distinct pathways regulated by hrpL and hrpRS. Mol. Plant. Microbe. Interact., 19, 976–87.

41. Merighi, M., Majerczak, D.R., Stover, E.H. and Coplin, D.L. (2003) The HrpX/HrpY two-component system activates hrpS expression, the first step in the regulatory cascade controlling the Hrp regulon in Pantoea stewartii subsp. stewartii. Mol. Plant. Microbe. Interact., 16, 238–48.

42. Luo, S., Sun, D., Zhu, J., Chen, Z., Wen, Y. and Li, J. (2014) An extracytoplasmic function sigma factor, σ(25), differentially regulates avermectin and oligomycin biosynthesis in Streptomyces avermitilis. Appl. Microbiol. Biotechnol., 98, 7097–112.

43. Tran, N.T., Huang, X., Hong, H., Bush, M.J., Chandra, G., Pinto, D., Bibb, M.J., Hutchings, M.I., Mascher, T. and Buttner, M.J. (2019) Defining the regulon of genes controlled by σ E, a key regulator of the cell envelope stress response in Streptomyces coelicolor. Mol. Microbiol., 10.1111/mmi.14250.

44. Kadowaki, T., Yukitake, H., Naito, M., Sato, K., Kikuchi, Y., Kondo, Y., Shoji, M. and Nakayama, K. (2016) A two-component system regulates gene expression of the type IX secretion component proteins via an ECF sigma factor. Sci. Rep., 10.1038/srep23288.

45. Seipke, R.F., Patrick, E. and Hutchings, M.I. (2014) Regulation of antimycin biosynthesis by the orphan ECF RNA polymerase sigma factor σ AntA. PeerJ, 10.7717/peerj.253.

46. Balogh, D., Dahmen, M., Stahl, M., Poreba, M., Gersch, M., Drag, M. and Sieber, S.A. (2017) Insights into ClpXP proteolysis: heterooligomerization and partial deactivation enhance chaperone affinity and substrate turnover in Listeria monocytogenes. Chem. Sci., 10.1039/c6sc03438a.

47. Yanamandra, S.S., Sarrafee, S.S., Anaya-Bergman, C., Jones, K. and Lewis, J.P. (2012) Role of the Porphyromonas gingivalis extracytoplasmic function sigma factor, SigH. Mol. Oral Microbiol., 10.1111/j.2041-1014.2012.00643.x.

48. Heruth, D.P., Pond, F.R., Dilts, J.A. and Quackenbush, R.L. (1994) Characterization of genetic determinants for R body synthesis and assembly in Caedibacter taeniospiralis 47 and 116. J. Bacteriol., 10.1128/jb.176.12.3559-3567.1994.

49. Müller, M., Reiß, S., Schlüter, R., Mäder, U., Beyer, A., Reiß, W., Marles-Wright, J., Lewis, R.J., Pförtner, H., Völker, U., et al. (2014) Deletion of membrane-associated Asp23 leads to upregulation of cell wall stress genes in Staphylococcus aureus. Mol. Microbiol., 10.1111/mmi.12733.

50. Shukla, J., Gupta, R., Thakur, K.G., Gokhale, R. and Gopal, B. (2014) Structural basis for the redox sensitivity of the Mycobacterium tuberculosis SigK-RskA σ-anti-σ complex. Acta Crystallogr. D. Biol. Crystallogr., 70, 1026–36.

51. Han, K., Li, Z., Peng, R., Zhu, L., Zhou, T., Wang, L., Li, S., Zhang, X., Hu, W., Wu, Z., et al. (2013) Extraordinary expansion of a Sorangium cellulosum genome from an alkaline milieu. Sci. Rep., 3, 2101.

52. Nechaev, S. and Severinov, K. (2003) Bacteriophage-induced modifications of host RNA polymerase. Annu. Rev. Microbiol., 57, 301–22.

53. Peng, Q. and Yuan, Y. (2018) Characterization of a novel phage infecting the pathogenic multidrug-resistant Bacillus cereus and functional analysis of its endolysin. Appl. Microbiol. Biotechnol., 102, 7901–7912.

54. Brister, J.R., Ako-Adjei, D., Bao, Y. and Blinkova, O. (2015) NCBI viral Genomes resource. Nucleic Acids Res., 10.1093/nar/gku1207.

55. Ortiz de Ora, L., Lamed, R., Liu, Y.J., Xu, J., Cui, Q., Feng, Y., Shoham, Y., Bayer, E.A. and Muñoz-Gutiérrez, I. (2018) Regulation of biomass degradation by alternative σ factors in cellulolytic clostridia. Sci. Rep., 10.1038/s41598-018-29245-5.

56. Braun, V., Mahren, S. and Ogierman, M. (2003) Regulation of the FecI-type ECF sigma factor by transmembrane signalling. Curr. Opin. Microbiol., 6, 173–80.

57. Sievers, F., Wilm, A., Dineen, D., Gibson, T.J., Karplus, K., Li, W., Lopez, R., McWilliam, H., Remmert, M., Söding, J., et al. (2011) Fast, scalable generation of high-quality protein multiple sequence alignments using Clustal Omega. Mol. Syst. Biol., 10.1038/msb.2011.75.

58. Finn, R.D., Clements, J. and Eddy, S.R. (2011) HMMER web server: Interactive sequence similarity searching. Nucleic Acids Res., 10.1093/nar/gkr367.

59. Nguyen, L.-T., Schmidt, H.A., von Haeseler, A. and Minh, B.Q. (2015) IQ-TREE: a fast and effective stochastic algorithm for estimating maximum-likelihood phylogenies. Mol. Biol. Evol., 32, 268–74.

60. Letunic, I. and Bork, P. (2016) Interactive tree of life (iTOL) v3: an online tool for the display and annotation of phylogenetic and other trees. Nucleic Acids Res., 10.1093/nar/gkw290.

61. Tsirigos, K.D., Peters, C., Shu, N., Käll, L. and Elofsson, A. (2015) The TOPCONS web server for consensus prediction of membrane protein topology and signal peptides. Nucleic Acids Res., 10.1093/nar/gkv485.

62. Tsirigos, K.D., Elofsson, A. and Bagos, P.G. (2016) PRED-TMBB2: Improved topology prediction and detection of beta-barrel outer membrane proteins. In Bioinformatics.

63. El-Gebali, S., Mistry, J., Bateman, A., Eddy, S.R., Luciani, A., Potter, S.C., Qureshi, M., Richardson, L.J., Salazar, G.A., Smart, A., et al. (2019) The Pfam protein families database in 2019. Nucleic Acids Res., 47, D427–D432.

64. Pedregosa, F., Varoquaux, G., Gramfort, A., Michel, V., Thirion, B., Grisel, O., Blondel, M., Müller, A., Nothman, J., Louppe, G., et al. (2012) Scikit-learn: Machine Learning in Python.

65. Li, W. and Godzik, A. (2006) Cd-hit: a fast program for clustering and comparing large sets of protein or nucleotide sequences. Bioinformatics, 22, 1658–9.

66. Wilbur, W.J. and Lipman, D.J. (1983) Rapid similarity searches of nucleic acid and protein data banks. Proc. Natl. Acad. Sci. U. S. A., 80, 726–30.

67. Cock, P.J.A., Antao, T., Chang, J.T., Chapman, B.A., Cox, C.J., Dalke, A., Friedberg, I., Hamelryck, T., Kauff, F., Wilczynski, B., et al. (2009) Biopython: Freely available Python tools for computational molecular biology and bioinformatics. Bioinformatics, 10.1093/bioinformatics/btp163.

68. Brown, D.P., Krishnamurthy, N. and Sjölander, K. (2007) Automated protein subfamily identification and classification. PLoS Comput. Biol., 3, e160.

69. Liu, X., Brutlag, D.L. and Liu, J.S. (2001) BioProspector: discovering conserved DNA motifs in upstream regulatory regions of co-expressed genes. Pac. Symp. Biocomput.

70. Crooks, G.E., Hon, G., Chandonia, J.M. and Brenner, S.E. (2004) WebLogo: A sequence logo generator. Genome Res., 10.1101/gr.849004.

71. Altschul, S.F., Gish, W., Miller, W., Myers, E.W. and Lipman, D.J. (1990) Basic local alignment search tool. J. Mol. Biol., 10.1016/S0022-2836(05)80360-2.

