## Supplementary Figures 1-5 for "Expansion and re-classification of the extracytoplasmic function (ECF) σ factor family"

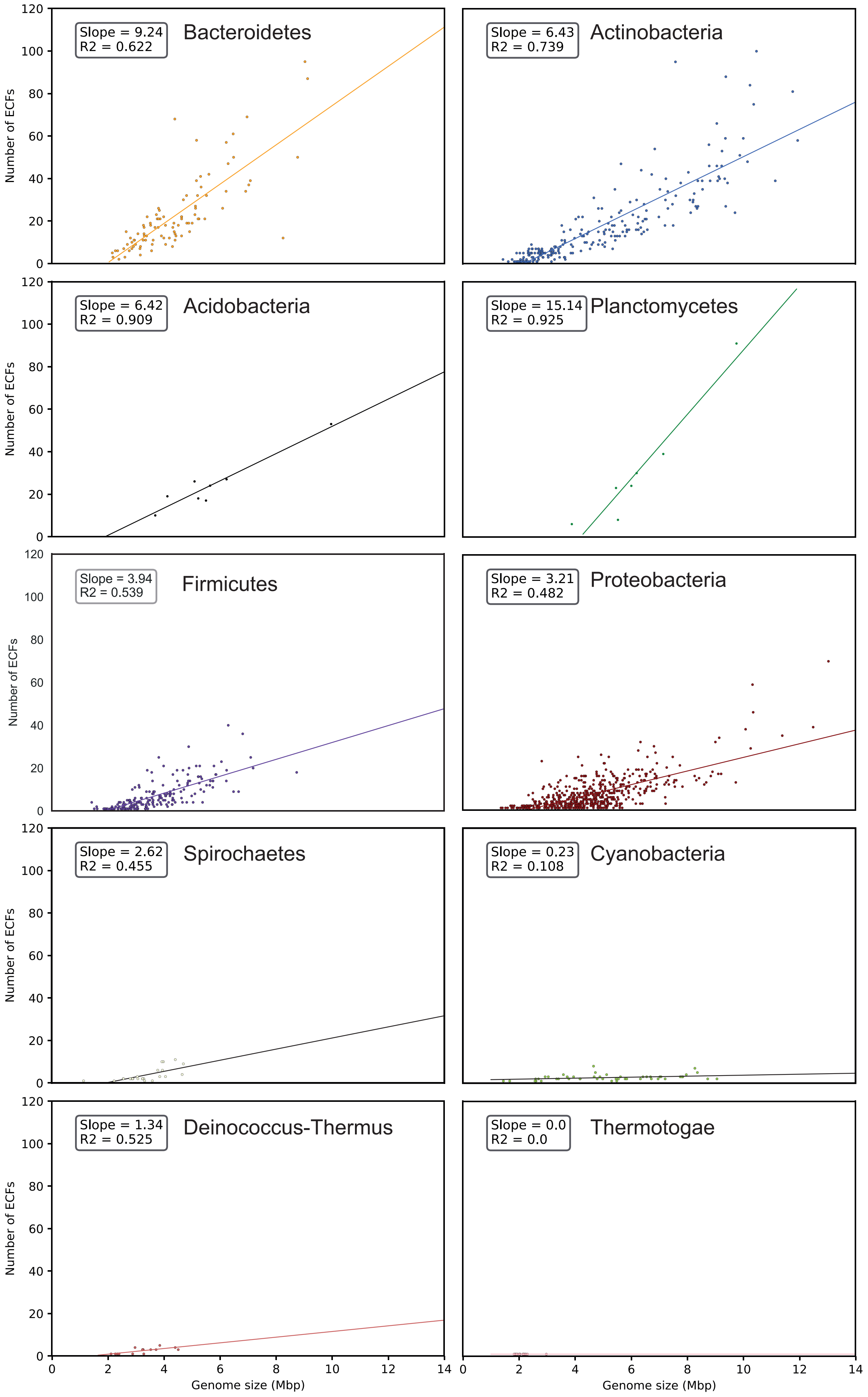
Slope = 9.24
R2 = 0.622
Bacteroidetes
Slope = 6.43
R2 = 0.739
Actinobacteria
Slope = 6.42
R2 = 0.909
Acidobacteria
Slope = 15.14
R2 = 0.925
Planctomycetes
Slope = 3.94
R2 = 0.539
Firmicutes
Slope = 3.21
R2 = 0.482
Proteobacteria
Slope = 2.62
R2 = 0.455
Spirochaetes
Slope = 0.23
R2 = 0.108
Cyanobacteria
Slope = 1.34
R2 = 0.525
Deinococcus-Thermus
Slope = 0.0
R2 = 0.0
Thermotogae
Number of ECFs
Genome size (Mbp)

**Figure S1. Number of ECFs per genome according to genome size for different phyla**. A linear regression with its fitting parameters is shown for each graph as a guide to the eye. Larger genomes, associated to more complex life styles, tend to have a larger number of ECFs.

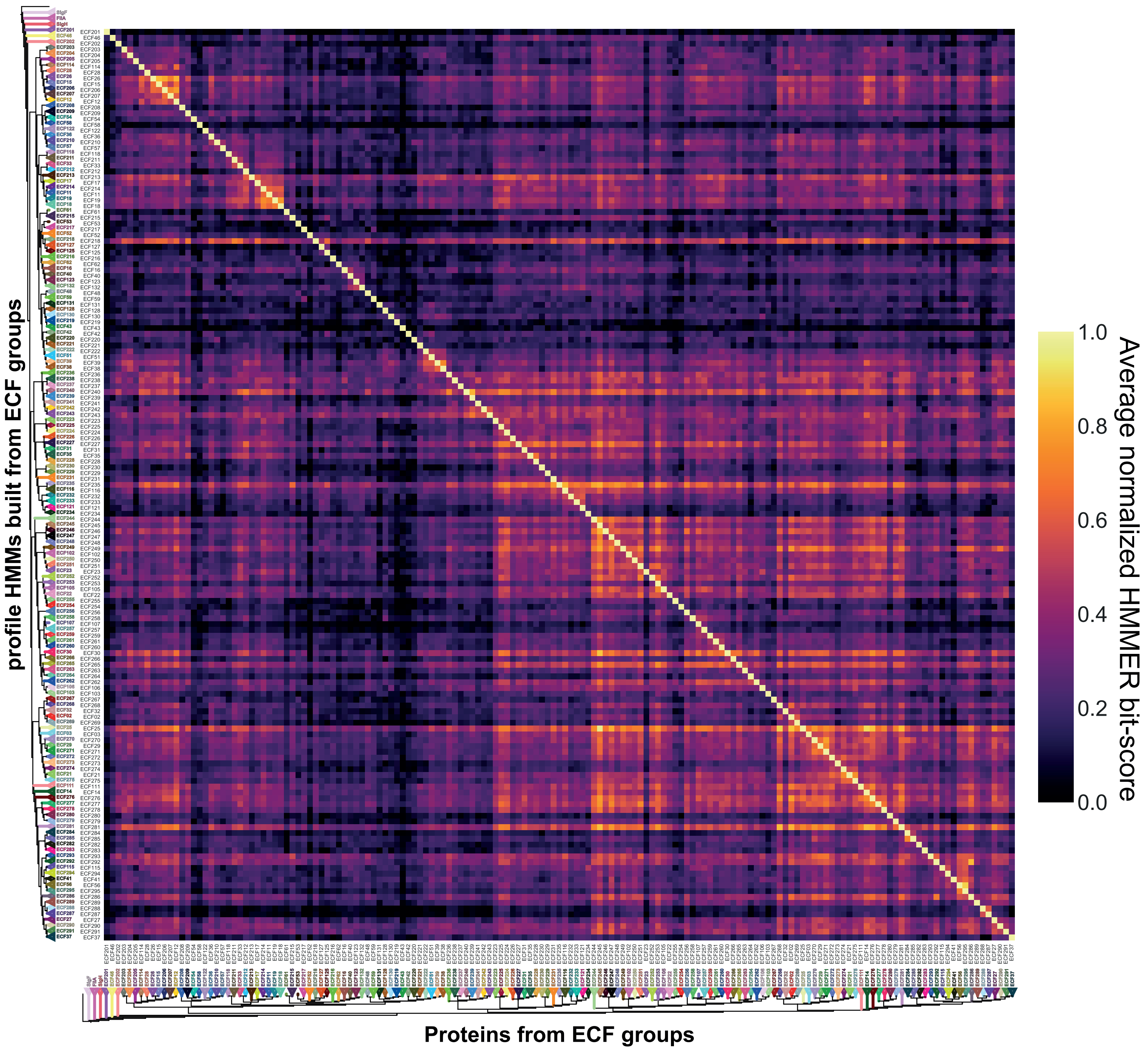


**Figure S2. Heatmap of the average normalized bit-score of each ECF group (x-axis) against each HMM (y-axis).** A schematic ECF tree is included in each axis as a guide. Values off diagonal tend to zero, indicating a good selectivity of the HMMs for the proteins of its own group. Areas with higher bit-scores (brighter spots) are observed more frequently between phylogenetically close groups, as in the case of ECF12-ECF26-ECF15-ECF206-ECF207 and ECF11-ECF19-ECF18.

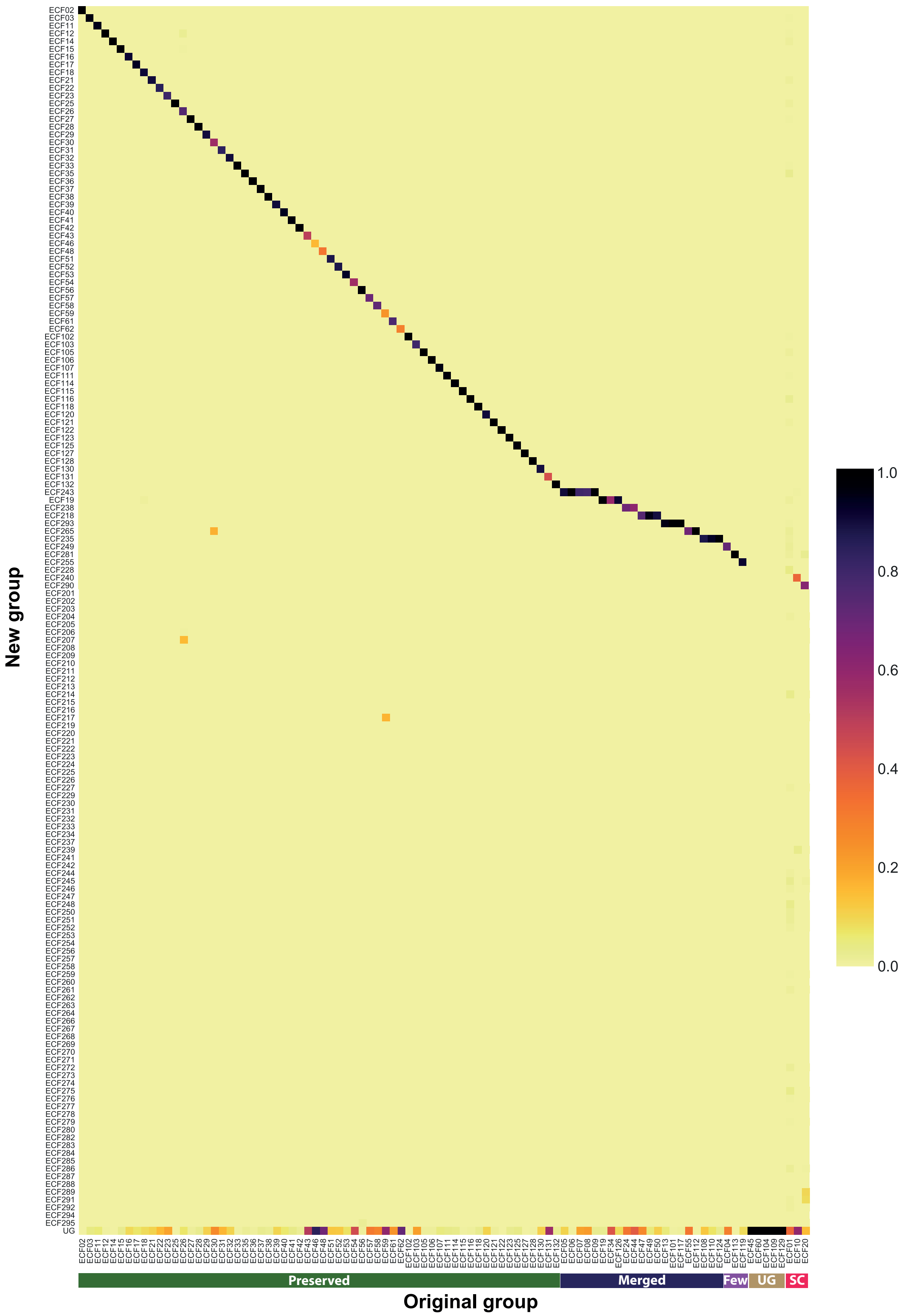



**Figure S3. Agreement between original and new ECF groups**. The heatmap shows the frequency of proteins in each original group (x-axis) associated to each new group (y-axis). Original groups could be: preserved (“Preserved”), if there is a new group that contains most of their elements; merged (“Merged”): when several original groups are merged into a single new group; suppose a low percentage of proteins of the new group (“Few”); be ungrouped (“UG”); or be scatter (“SC”) across several groups of the new classification. New groups are named after their original group its characteristics are preserved.

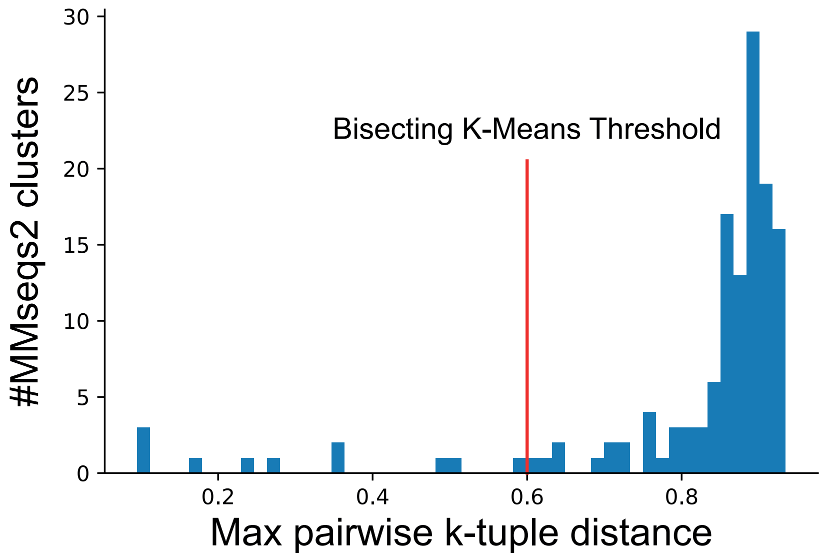


**Figure S4. Selection of maximal pairwise K-tuple distance threshold.** Histogram of the maximum pairwise K-tuple distance for clusters resulting from MMseqs2 execution. The threshold selected as maximum allowed K-tuple distance for any pair of proteins within any cluster was 0.6 (indicated by a red line). This threshold was selected empirically since it was the largest maximum k-tuple distance whose associated clusters contained proteins similar enough to produce homogeneous multiple-sequence alignments. Clusters that have a maximum K-tuple distance above threshold are subjected to bisecting K-means until no children cluster is above threshold.

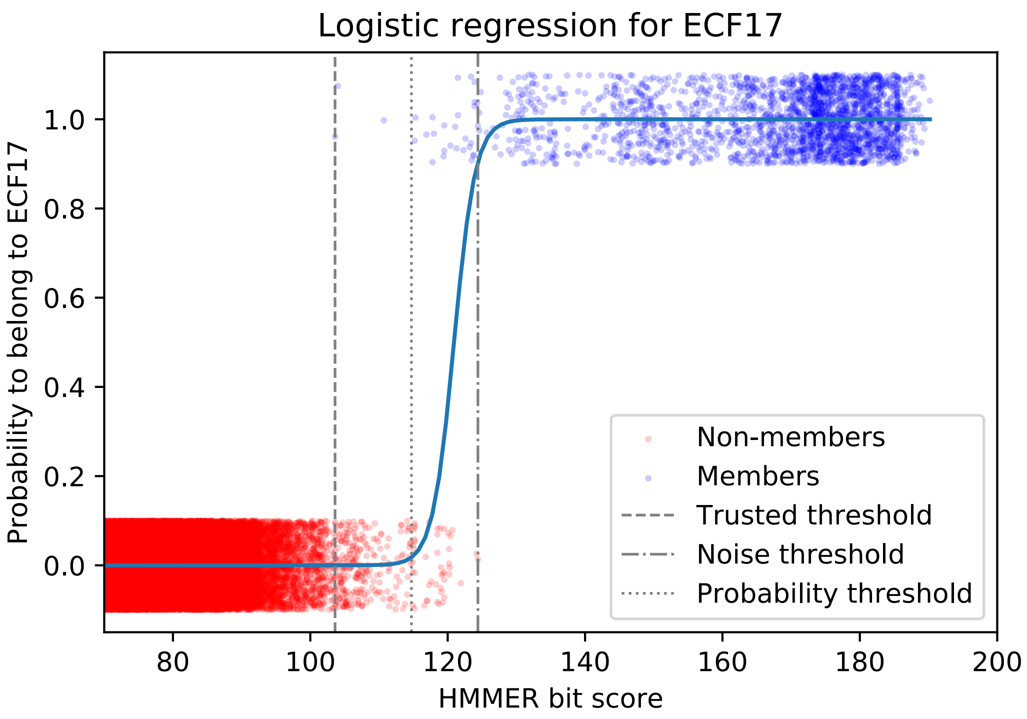


**Figure S5. Example logistic regression.** Logistic regression fitting the distribution of scores of members (y=1, blue) and non-members (y=0, red) of group ECF17. Proteins (represented as points) have been plotted ±0.1 in the y-axis for clarity. X-axis has been capped at bit score=75 for clarity. Three thresholds are used for protein classification, namely, trusted threshold, i.e. bit score of the lowest scoring member, noise threshold, i.e. largest score of any non-member, and probability threshold, i.e. ROC-optimized probability that a protein belongs to a group (see Methods). Thresholds for groups, subgroups and original groups are available in Table S5.
