## Supplementary Text for "Expansion and re-classification of the extracytoplasmic function (ECF) σ factor family"

### Preface

The founding ECF classification showed that similar ECFs (i.e. ECFs from the same group) share similar regulation, input stimuli, response elicited and/or target promoter (Staroń et al., 2009). The number of ECFs characterized *in vivo* is not enough for a comprehensive study of the features of the ECF groups; however, the special features of ECFs, including autoinduction of their own expression and coevolution with the genes involved in their pathway as part of the same genetic neighborhood, made it possible to hypothesize over the functional features of original ECF groups from *in silico* data. In this study we harness the wealth and diversity of new proteins and the data gathered from the original groups (Pinto & Mascher, 2016) in an attempt to describe the function of the new ECF groups. We analyzed the groups in terms of taxonomic distribution, original group composition, sequence features of the ECF (e.g. presence of extensions or transmembrane helices), genetic context conservation, presence of putative regulators such as anti-σ (AS) factors and predicted target promoter motifs. Unless otherwise stated, we only targeted the ECFs from organisms defined as representative or reference by NCBI, including only RefSeq genomes when GenBank and RefSeq assemblies exist for the same genome. In this manner we lessen the bias towards organisms with more genomes sequenced. Domains and domain architectures are conserved in they occur in >75% of the ECFs of the group or subgroup under analysis. We only considered subgroups with more than 10 representatives in the genomic context analysis so as to avoid bias caused by small datasets. In the following we provide a catalog of standardized ECF group descriptions, highlighting the most important common features of the constituting subgroups. More details can be found in Tables S1-S5.

### Description of ECF groups

#### ECF01 (original classification; no longer present)

Members of original ECF01 are broadly scattered across the new classification. The new group that receives most of the sequences of ECF01 is ECF228, with 4% of the proteins from ECF01.

#### ECF02

1. **General description:** Members of ECF02 have homology against proteins of original ECF02 (99.29%) and appear in Proteobacteria (100**%**).
2. **Anti-σ factor:** Members of ECF02 are regulated by RseA-like AS factors with one transmembrane helix encoded in position +1. Even though TopCons failed to predict the transmembrane helix in most of the cases, an MSA revealed the presence of one transmembrane helix in most of the proteins encoded in +1 of members of ECF02. RseA-like AS factors take their name from RseA, the AS factor of RpoE in *E. coli* (ECF02s1) (Missiakas, Mayer, Lemaire, Georgopoulos, & Raina, 1997).
3. **Genomic context conservation:** Other conserved domains in the context of ECF02 are associated with translation and protein secretion. We found a 50S ribosome-binding GTPase fused to a KH domain, an elongation factor Tu involved in the translation elongation, a ribonuclease-III-like protein involved in rDNA transcription and rRNA processing, a signal peptidase S24 (Pfam: Peptidase_S24), a signal peptidase S26 and a fumarate reductase (only ECF02s2).
4. **Studied members:** Among the characterized members of ECF02 we found RpoE, from *E. coli*, which plays a key role during stationary phase, metal resistance (Egler, Grosse, Grass, & Nies, 2005), folding and degradation of periplasmic proteins, lipopolysaccharide biogenesis, heat shock response (Rowley, Spector, Kormanec, & Roberts, 2006) and virulence (D. R. Smith et al., 2017). Unlike other ECFs, RpoE is essential in *E. coli* (Hiratsu, Amemura, Nashimoto, Shinagawa, & Makino, 1995), potentially due to the lack of a complete lipopolysaccharide in respect to the closely related *Salmonella enterica*, where RpoE is not essential (Amar, Pezzoni, Pizarro, & Costa, 2018). RpoE is regulated by its AS factor, RseA, which is inactivated by Regulated Intramembrane Proteolysis (RIP) sequentially carried out by the proteases DegS, RseP and cytoplasmic proteases (Akiyama, Kanehara, & Ito, 2004; Walsh, Alba, Bose, Gross, & Sauer, 2003). DegS is a membrane-bound periplasmic protein able to sense stress signal originated from different unfolded OMP via its PDZ domain (Hasselblatt et al., 2007). RseP contains a pair of permutated PDZ domains in its periplasmic region that control its activity as site-2 protease (Inaba et al., 2008). Indeed,

proteins with PDZ domain appear in the genetic context of ECF02 (1.17 per ECF). The other two proteins in the operon of RpoE - RseB (+2) and RseC (+3), are an extracytoplasmic stabilizer of RseA and an inner membrane positive regulator of RpoE, respectively (Cezairliyan & Sauer, 2007; Missiakas et al., 1997). While RseB-like proteins are conserved in positions +2 of all the subgroups, RseC-like proteins are not conserved in ECF02s3.

RpoE from *S. enterica* (ECF02s1) is activated by UVA radiation, envelope stress generated by phage infection or hypo-osmotic shock in the absence of periplasmic glycans (OPG) (Amar et al., 2018; Spöring et al., 2018; E. C. Woods & McBride, 2017). RpoE from *Burkholderia pseudomallei* is induced under high salt concentrations and is required for heat and oxidative stress since it turns on the expression of *speG*, stabilizing the levels of spermidine (Duangurai, Indrawattana, & Pumirat, 2018). RpoE from *Xylella fastidiosa* participates in heat shock response due to 21-gene sigmulon (da Silva Neto, Koide, Gomes, & Marques, 2007). Similarly, RpoE from *Shewanella oneidensis* is essential for growth under suboptimal conditions caused by temperature, salinity, oxidative stress, minimal medium, metals, absence of oxygen, among others, and it typically targets the synthesis of extracytoplasmic and outer membrane components (Barchinger et al., 2016; Dai et al., 2015). *Vibrio parahaemolyticus* contains an ECF02 (VP2578) involved in cell envelope stress required for intestinal colonization (Haines-Menges, Whitaker, & Boyd, 2014).

Interestingly, RpoE1 from *Xanthomonas campestris* (ECF02s1) is induced under cultivation in minimal medium or plant extract, and it is required by plant pathogenesis since it induces the expression of T3SS (Yang et al., 2018). This function is typical from group ECF32, which is in close evolutionary proximity to ECF02. Therefore, RpoE1 from *X. campestris* could be the mixture of both systems where the ECF kept the sequence of members of ECF02 but the function of ECF32.

Other members of ECF02 are the AlgU-like proteins — AlgU from *Pseudomonas aeruginosa* in involved in alginate production leading to transition the mucoid and biofilm phenotypes. AlgU helps to relieve oxidative stress, and it is involved in virulence, β-lactamase production, and repression of flagella biosynthesis. AlgU is induced by antibiotics that block peptidoglycan synthesis as in the case of RpoE from *E. coli* (Llamas et al., 2014; E. C. Woods & McBride, 2017) (Chevalier et al., 2018; Delgado et al., 2018; Tart, Wolfgang, & Wozniak, 2005). Homologs of AlgU appear in other *Pseudomonas* species such as *P. putida* and *P. syringae*, but also in *Azotobacter vinelandii*.

5. **Promoter motif conservation:** Promoter motifs are well conserved and contain GAACTTT in -35 and GTCT in -10 in agreement with the binding motif of RpoE in *E. coli* (Rhodius & Mutalik, 2010), *Shewanella oneidensis* (Barchinger et al., 2016) and original ECF02 (Staroń et al., 2009). Nevertheless, some members of ECF02, such as RpoE from *Xylella fastidiosa* are not autoregulated (da Silva Neto et al., 2007).
6. **Summary:** ECF02 keeps the characteristics of the original group ECF02 (Staroń et al., 2009). ECF02 is one of the key factors contributing to cell envelope homeostasis in Proteobacteria. These ECFs are also involved in biofilm formation since they induce the production of alginate in bacteria from the genus *Pseudomonas*. Members of ECF02 are negatively regulated by a RseA-like AS factor encoded in position +1, a RseB-like extracytoplasmic negative regulator that protects RseA from degradation and a RseC-like inner membrane positive regulator.

**ECF03**

1. **General description:** Members of ECF03 are homologous to proteins from original ECF03 (89.41%) and are mainly present in Bacteroidetes (95.44%), but also a variety of Gram-negative organisms such as Proteobacteria (1.27%), Verrucomicrobia (2.28%), Kiritimatiellaeota (0.25%) and Chlorobi (0.25%).
2. **Anti-σ factor:** The genomic context is not conserved although it is possible to identify a putative AS factor with one TM helix (62.03%, ~100% in the MSA)in position +1. These putative AS factors of ECF03 extend their conservation also to their extracytoplasmic site.
3. **Genomic context conservation:** Proteins from ECF03s4 have extended genomic context conservation, with an NADP oxidoreductase in -1, a Maf-like protein, an elongation factor Tu fused to a LepA protein, a haloacid dehalogenase-like hydrolase, a UPF0102 protein, a glycoprotease, and an LD-carboxypeptidase. Subgroups ECF03s3 and ECF03s5 contain one or several conserved transketolases. Last, ECF03s10 contains an LTXXQ motif family protein in +2 (involved in relieving extracytoplasmic stress (Danese & Silhavy, 1998)) and a universal stress protein.
4. **Promoter motif conservation:** Predicted target promoter motifs are usually TTAAACC in -35 and a less conserved TC-containing -10, as described in previous literature (Rhodius et al., 2013).
5. **Summary:** Unlike proteins from original ECF03, new ECF03 seems to be regulated by a new type of AS factors with no registered PFAM domain. These AS factors contain a conserved extracytoplasmic domain. The reason for the lack of conserved PFAM domain in ECF03 could be due to the presence of most of the

sequences from ECF03 only in Bacteroidetes, organisms where genetic studies are limited. The presence of LTXXQ-motif proteins and universal stress proteins indicates that ECF03 could be part of the cell envelope stress response as its neighboring ECF groups.

**ECF04 (original classification; no longer present)**

This group is now present in ECF249. 2.63% of the proteins from ECF249 have homology to original ECF04.

**ECF05 (original classification; no longer present)**

This group is now part of ECF243 (FecI-like group). 45.1% of the proteins from ECF243 have homology to original ECF05. ECF243 merges original ECF05-09.

**ECF06 (original classification; no longer present)**

This group is now part of ECF243 (FecI-like group). 6.67% of the proteins from ECF243 have homology to original ECF06. ECF243 merges original ECF05-09.

**ECF07 (original classification; no longer present)**

This group is now part of ECF243 (FecI-like group). 16.62% of the proteins from ECF243 have homology to original ECF07. ECF243 merges original ECF05-09.

**ECF08 (original classification; no longer present)**

This group is now part of ECF243 (FecI-like group). 3.14% of the proteins from ECF243 have homology to original ECF08. ECF243 merges original ECF05-09.

**ECF09 (original classification; no longer present)**

This group is now part of ECF243 (FecI-like group). 5.5% of the proteins from ECF243 have homology to original ECF09. ECF243 merges original ECF05-09.

**ECF10 (original classification; no longer present)**

This group is scattered across several groups in this classification. Members of ECF10 are mainly present in two new groups: ECF239 and ECF240. ECF249 contains 82.76% of the proteins with homology to original ECF10 and preserves most of the characteristics of original ECF10, whereas members of ECF239 do not contain a carbohydrate-binding domain in their genetic context and contain conserved target promoter motifs, indicating the autoregulation of its members.

**ECF11**

1. **General description:** ECF11 is mainly composed of proteins with homology to original ECF11 (83.58%). Members of this group are present in Proteobacteria (Gamma and Alphaproteobacteria).

2. **Anti-σ factor:** Members of ECF11 are regulated by putative AS factors with a ChrR cupin-like domain (CLD) encoded in position +1, as in the case of members of original ECF11 (Staroń et al., 2009). This protein may contain a zinc finger, and it is absent in ECF11s6, where it could be encoded elsewhere in the genome. Putative AS factors of members of ECF11 are soluble proteins (100%), as expected from the AS of original ECF11 (Staroń et al., 2009).
3. **Genomic context conservation:** Members of ECF11 contain an average of about one flavin-containing amino oxidoreductase per genomic context. This protein is encoded in position -4, -3 or -1 and is part of the set of domains that appears in several subgroups of ECF11, among which we also found the substrate-binding domain of LON ATP-dependent protease (position -1), a SnoaL-like domain-containing protein (positions -2 or -1), a short chain dehydrogenase (position -3 or -2), a DUF1365-containing protein (position -5, -4 or -2) and a mycolic acid cyclopropane synthetase.
4. **Studied members:** The most studied member of this group, $\sigma^E$ from *Rhodobacter sphaeroides* (ECF11s1), is inhibited by its anti-σ factor, ChrR (Campbell et al., 2007). ChrR is a soluble zinc-finger AS factor with a C-terminal CLD, which also binds a zinc atom but with less affinity than the zinc-finger in the AS factor domain (Greenwell, Nam, & Donohue, 2011). The CLD senses the presence of oxidative stress from singlet oxygen or tert-butyl hydroperoxide (Campbell et al., 2007; Greenwell et al., 2011). Another member of this group, SigW from *Pseudomonas syringae* (ECF11s4), responds to the same kind of stresses and is regulated by a ChrR type AS factor (Butcher et al., 2017).
5. **Promoter motif conservation:** The predicted target promoter motifs found upstream of the ECF11 coding sequences are conserved and have a consensus of GTGATC for -35 and CGTA for -10. This promoter matches previous predictions (Staroń et al., 2009) and experimental data (Butcher et al., 2017; Newman et al., 1999). The coherent prediction of the promoter motifs within and across subgroups reflects the autoregulatory role of ECF11 (Butcher et al., 2017; Newman et al., 1999) and the conservation of its members.
6. **Summary:** Members of ECF11 are associated with a soluble AS factors with a cupin-like domain (CLD) involved in the sensing of oxidative stress induced by singlet oxygen or an organic peroxide. The association of ECF11 with soluble zinc-binding AS factors and their involvement in oxidative stress response reminds of ECF12. However, ECF12 has a different target promoter sequence, is present in more taxonomic groups, and usually responds to disulfide stress rather than oxidative stress *per se*.

**ECF12**

1. **General description:** Members of ECF12 are homologous to proteins from original ECF12 (86.15%) and ECF26 (3.5%). Members of ECF12 are mainly present in Actinobacteria (88.76%), but also in Bacteroidetes (6.91%), Proteobacteria (1.75%), Chlorobi (1.24%), Ignavibacteriae (0.31%), Planctomycetes (0.21%), Firmicutes (0.10%) and Calditrichaeota (0.10%).
2. **Regulation:** As in the original group ECF12 (Staroń et al., 2009), the genetic neighborhood of most of the members of new ECF12 contain a putative AS factor in position +1. These putative AS factors contain a zinc-finger and are soluble (96.05%), except in the case of subgroup ECF12s19, where the putative AS factors are transmembrane.
3. **Special features:** No putative AS factor is present for some of the smaller subgroups, which could be encoded somewhere else in the genome. One of these subgroups is ECF12s9. It contains an ECF from ECF12s2 in +1 in three out of four proteins from representative genomes. Interestingly, none of the ECFs from ECF12s9 has a putative AS associated, they are transcribed convergently, and their 3' ends are overlapping, which could indicate that these ECFs are mutually exclusive and only one can be expressed at a time. Regulation by transcriptional interference caused by convergent, overlapping transcripts has been shown to resolve in RNAP collision, truncated RNA (Shearwin, Callen, & Egan, 2005) and was used to generate bistable switch that responds to a signaling molecule in *Streptomyces coelicolor* (Chatterjee et al., 2011). If this is a regulatory mechanism for members of ECF12s9 and some members of ECF12s2, it remains unclear what triggers the switch towards the opposite ECF and what are the differences in the response mediated by these two ECFs, given their high sequence similarity. Subgroup ECF12s22, with a single sequence from representative/reference genomes, contains a σ factor in position +1 and lacks of an evident putative AS factor. Nevertheless, the two σ factors are encoded in tandem, non-overlapping configuration, discarding a transcriptional interference mechanism in this subgroup.
4. **Genomic context conservation:** Other domains conserved in some ECF12 groups are a guanylate kinase (+2 of ECF12s6), a cytidylyltransferase-like protein (ECF12s6) and an EPSP synthase (ECF12s1).
5. **Studied members:** One of the members of ECF12, $\sigma^R$ from *Streptomyces coelicolor* (ECF12s1), responds to thiol-oxidative stress by inducing thioredoxin, mycothiol metabolism, proteases, among others (Kim, Hahn, Cho, Cho, & Roe, 2009). $\sigma^R$ is activated when its AS factor, RsrA, forms disulfide bonds within its zinc-binding domain under exposure to oxidation agents and disulfide stress (Paget et al., 2001; Zdanowski

et al., 2006). Upon stress exposure, $\sigma^R$ induces its expression from an earlier promoter that produces a variant of $\sigma^R$ ($\sigma^{R'}$) 55aa longer and more prone to be degraded by the also upregulated ClpP1/P2 proteases (Kim et al., 2009). It is not clear whether this promoter-shift mechanism is present in other members of ECF12.

SigH from *M. tuberculosis* (ECF12s1) is induced by heat, oxidative and nitric oxide stresses and it activates the expression of a broad set of genes involved in DNA repair, sulfur metabolism and translation recovery (Sharp et al., 2016). The AS factor of SigH, RshA, is located in position +2, coordinates a [2Fe-2S] cluster and binds SigH using salt bridges (Kumar et al., 2012). Apart from oxidants, RshA is also inhibited by the eukaryotic-like Ser/Thr kinase PnkB (Park, Kang, & Husson, 2008), which illustrates the integration of different sensing pathways in bacteria. Members of ECF12s1 do not usually contain AS factors in position +2.

6. **Promoter motif conservation:** SigH from *C. glutamicum* (ECF12s2) binds to the same motif a the prediction for its subgroup – GGAAT in -35 and GTT in -10 (Pátek et al., 2018). The target promoter of SigH (GGAAYR-(N17-18)-GTT) (Sharp et al., 2016) does not match the motif observed for subgroup ECF12s1, but ECF12s2. In general, predicted promoter motifs for the subgroups of ECF12 are not conserved.
7. **Summary:** Members of ECF12 are related to oxidative and disulfide stress response. The soluble AS factors associated to this group coordinate metals that serve to sense oxidative stress and release the inhibition over the ECF. The nature of the metal might differ, even within the same subgroup. Moreover, the putative AS factor is lacking in some subgroups, raising the possibility of another regulatory mechanism such as transcriptional interference in subgroup ECF12s9. Nevertheless, it is possible that their AS factor is encoded somewhere else in the genome.

**ECF13 (original classification; no longer present)**

Members of original ECF13 are present in new ECF293, together with members of original ECF101 and ECF117. 43.98% of the proteins of ECF293 have homology to original ECF13.

**ECF14**

1. **General description:** Proteins from ECF14 have homology to original ECF14 (81.93%), which appears in subgrups from Actinobacteria (93.53%). ECF14 is also present in Proteobacteria (3.07%) (ECF14s5, ECF14s7, ECF14s9) and Armatimonadetes (0.65%) (ECF14s10).

2. **Anti-σ factors:** Proteins from ECF14 are regulated by a ZAS located in +1. The conservation observed for the N-termini of this AS factors does not extend to the transmembrane domain. Putative AS factors of ECF14 are soluble (47%) or with one transmembrane-helix membrane (52.14%).
3. **Genomic context conservation:** Proteins from ECF14s1, ECF14s3, and ECF14s4 (Actinobacteria) encode a conserved O-methyltransferase in -1, in agreement with previous findings in original ECF14 (Staroń et al., 2009). The following proteins are conserved in the genomic context of members of ECF14: The protein mtt/Hcf106 is located in +2 and +3, which is involved in a sec-independent translocation mechanism. A NUBPL iron-transfer P-loop NTPase (PFAM: ParA) fused to an iron-sulfur cluster assembly protein usually located in +4, domains present in ATPases involved in plasmid partitioning in prokaryotes and to a protein required for the biogenesis and export of ribosomal subunits in eukaryotes (Pfam), and a trypsin-like peptidase usually in position +2. Subgroup ECF14s6 is located in one of the outermost leaves of the tree of ECF14 and, even though it contains a ZAS encoded in +1, it lacks the rest of the proteins conserved in the genetic neighborhoods of rest of the subgroups.
4. **Studied members:** Described proteins from ECF14 include SigE from *M. tuberculosis* (ECF14s1), *M. marinum* (ECF14s1), *C. glutamicum* (ECF14s2) and *C. pseudotuberculosis* (ECF14s2). These proteins are involved in oxidative stress response and defense against cell envelope stress caused by SDS, vancomycin, and lysozyme. They might also be involved in fatty acid degradation (*M. tuberculosis* (Souza et al., 2014)), starvation, survival in stationary phase and thermal stress (Pettersson et al., 2015; Souza et al., 2014). In SigE from *M. tuberculosis*, the Ser/Thr protein kinase PknB is required to phosphorylate the AS factor RseA, which is degraded as a consequence (Barik, Sureka, Mukherjee, Basu, & Kundu, 2010).
5. **Promoter motif conservation:** Putative target promoter motifs contain CTACTGG in -35, similarly as original ECF14 (Rhodius et al., 2013), and a non-conserved -10 with a G-tract in some cases.
6. **Summary:** ECF14 is involved in response to a wide range of stresses, including oxidative stress, starvation, thermal shock, and cell envelope stress. The characteristics of new ECF14 match original ECF14 (Staroń et al., 2009), where the response mechanisms involves a ZAS encoded in +1 and an O-methyltransferase encoded in -1. Since the functions of ECF14 are so broad, it could be possible that ECF14 modifies basic cellular processes in an attempt to respond to the stress agent.

**ECF15**

1. **General description:** Group ECF15 is mainly composed by members of original ECF15 (88.52%) with some members of original ECF26 (1.76%), and it is present in Proteobacteria (99.86%) and Cyanobacteria (0.14%).
2. **Regulation:** Original ECF15 is known to be regulated by a partner-switching mechanism (Staroń et al., 2009). The first identified example of partner-switching regulation was EcfG1 from *Methylobactrium extorquens* (Francez-Charlot et al., 2009). EcfG1 is an ECF negatively regulated by the anti-σ factor NepR. Apart from NepR, PhyR, a protein containing a receiver domain and a σ-like region, is encoded in the genomic context of EcfG1. PhyR acts as the response regulator of a two-component system and as an anti-anti-σ factor (AAS) under inducing conditions, titrating NepR and triggering the release of EcfG1 into its active form (Campagne et al., 2012; Francez-Charlot et al., 2009).
   Indeed, we found that $\sigma_2$ domain (1.29 per ECF), $\sigma_4$ domain (1.91 per ECF) and the receiver domain of a response regulator (1.16 per ECF, usually in position -1) were conserved in the genetic context of members of ECF15. Moreover, histidine kinases are also in position +2 of ECF15s5 and ECF15s6, although when looking at all the members of ECF15 we only found an average of 0.56 histidine kinases per ECF. This suggests that in some instances the histidine kinase might be encoded somewhere else in the genome, as in the case of EcfG1 from *Methylobactrium extorquens* (Francez-Charlot et al., 2009), RpoE from *Bradyrhizobium japonicum,* RpoE from *Rhodobacter sphaeroides* (Li, Peng, & Klug, 2018) and RpoE2 from *Sinorhizobium meliloti* (Sauviac, Philippe, Phok, & Bruand, 2007). The anti-σ factors could be identified through a BLAST search using as a library the AS factors from the original ECF classification (Staroń et al., 2009). Their position ranges from -2 to +2 in subgroups ECF15s1, ECF15s4, ECF15s5, ECF15s7 and ECF15s9. AS factors from ECF15 do not bear any PFAM domain and are cytoplasmic (90.32%).
3. **Genomic context conservation:** Genetic context conservation in ECF15s4 and ECF15s5, which share genetic context since they are transcribed consecutively, extends to a SecD/SecF GG motif and a protein-export membrane protein domain (PFAM: Sec_GG and SecD_SecF), which corresponds to the subunit of the SecDFYajC complex involved in the proton motive force-dependent steps of proteins translocation (Tsukazaki et al., 2011). Subgroup ECF15s7 contains a conserved superoxide dismutase, in charge of detoxifying superoxide.

4. **Studied members:** Characterized members of ECF15 include SigT from *Caulobacter crescentus* (Alvarez-Martinez, Lourenço, Baldini, Laub, & Gomes, 2007), RpoE from *Rhodobacter sphaeroides* (Li et al., 2018), EcfG1 from *Methylobactrium extorquens* (Francez-Charlot et al., 2009), RpoE from *Bradyrhizobium japonicum* (Gourion et al., 2009), RpoE4 from *Rhizobium etli* (Martinez-Salazar, Salazar, Encarnacion, Ramirez-Romero, & Rivera, 2009), RpoE2 from *Sinorhizobium meliloti* (Sauviac et al., 2007) and EcfT from *Rhodopseudomonas palustris* (Allen et al., 2015). Members of ECF15 are the main general stress response σ factor in most Alphaproteobacteria (Allen et al., 2015; Alvarez-Martinez et al., 2007; Francez-Charlot et al., 2009; Sauviac et al., 2007). However, in some cases, ECF15 members have more specialized functions in defense against membrane stress (Li et al., 2018).
5. **Promoter motif conservation:** The target promoters of members of ECF15 are conserved. The -35 region is usually GGAAC and the -10 CATT, which fits experimental data (Allen et al., 2015; Alvarez-Martinez et al., 2007; Gourion et al., 2009; Sauviac et al., 2007).
6. **Summary:** In summary, similarily to ECFs from original ECF15 (Staroń et al., 2009), members of ECF15 are regulated by parter-switching involving a response regulator that, when activated by its partner histidine kinase, acts as AAS factor, titrating the AS factor and releasing the active form of the ECF.

**ECF16**

1. **General description:** Members of ECF16 are present in Proteobacteria (96.5%), mainly Alphaproteobacteria, and ECF16s14 to Spirochaetes (2.67%). All the subgroups contain proteins with homology to original ECF16 (85.71%).
2. **Anti-σ factor:** Proteins from ECF16 are regulated by a putative AS factor with a conserved DUF1109 encoded in position +1. This protein contains six transmembrane helices (95.8%) and two conserved cysteine residues, as in the experimentally addressed AS factors of ECF16.
3. **Genomic context conservation:** Other conserved proteins in the genetic context of members of ECF16 are a DUF692-containing protein (ECF16s15 and ECF16s16) a DUF2282-containing protein (ECF16s15 and ECF16s16), a DoxX-domain containing protein (ECF16s15 and ECF16s16), an enoyl-CoA hydratase/isomerase (ECF16s2), a putative DNA-binding domain (ECF16s16), a cytochrome C biogenesis protein fused to a thioredoxin-like domain (ECF16s10), a AhpC/TSA family (ECF16s10), an outer membrane efflux protein (ECF16s2), a glutathione S-transferase (ECF16s19) and a protein from the AcrB/AcrD/AcrF family (ECF16s5).

4. **Studied examples:** Some members of ECF16, SigF from *Caulobacter crescentus* and *Bradyrhizobium japonicum*, have been functionally addressed. They are regulated by an AS factor (NrsF from *C. crescentus* and OsrA from *B. japonicum*) with six transmembrane helices and two cysteine residues facing the periplasm that are essential for sequestering their cognate ECF σ factor (Kohler, Lourenço, Avelar, & Gomes, 2012; Masloboeva et al., 2012).
5. **Promoter motif conservation:** Members of this group respond to heavy-metal stress (Kohler et al., 2012) or oxidative stress (Masloboeva et al., 2012). Both SigF from *C. crescentus* (ECF16s20) and *B. japonicum* (ECF16s2) self-induce their expression, and SigF from *C. crescentus* has the target promoter GTAACC-CGTA (Kohler et al., 2012; Masloboeva et al., 2012). These data do not agree with the predicted promoter elements (normally TTC in -35 and TAC or TAAC in -10).
6. **Summary:** Members of ECF16 are associated with DUF1109-containing AS factors with six transmembrane helices and conserved periplasmic cysteine residues. Their promoter motifs, although conserved, do not match experimental data. Members of ECF16 are involved in heavy metal and oxidative stress response.

**ECF17**

1. **General description:** Group ECF17 is composed of 85.82% of proteins with homology to the original group ECF17. All the proteins are present in Actinobacteria.
2. **Anti-σ factors:** Members of ECF17 might be regulated by putative AS factors with a zinc-finger and one transmembrane helix (83.33%). Putative AS factors are encoded in position +1, except in ECF17s1, where it is located mainly in position +2. Subgroups ECF17s18 and ECF17s19 lack of putative AS factors in their genetic context. Indeed, previous studies identified this protein as an AS factor previous studies (Gordon et al., 2008; Hahn, Raman, Anaya, & Husson, 2005; Staroń et al., 2009).
3. **Genomic context conservation:** Aside from the AS factor, the remaining conserved proteins are subgroup-specific. ECF17s1 contains a conserved protein with a bacterial capsule synthesis protein PGA_cap (Pfam: PGA_cap) in position -1, a conserved formamidopyrimidine-DNA glycosylase, a universal stress protein, and a sortase. ECF17s2 and ECF17s11 contain a conserved protein from the Major Facilitator Superfamily (MFS). Consequent with this conserved genomic context, the only described member of ECF17s2, SigU from *Streptomyces coelicolor* is involved in response to envelope stress and morphological differentiation (Gordon et al., 2008). ECF17s4 contains a conserved metallopeptidase from family M24 in position -1, an

adenylate kinase in position -2, a SecY translocase in position -3 and ~2 copies of a leucine carboxyl methyltransferase. Lastly, ECF17s11 encodes a secreted protein in position -1 and members of ECF17s18 encode a conserved N-acetylmuramoyl-L-alanine amidase in their genetic context.

4. **Studied members**: The genetic context of ECF17s4 agrees with the function of its only characterized member, SigL from *M. tuberculosis*, which is involved in the synthesis of envelope lipids, extracytoplasmic protein modification, and pathogenesis, although none of the genes in its genetic neighborhood are regulated by SigL (Hahn, Raman, Anaya, & Husson, 2005).
5. **Promoter motif conservation:** Promoter motifs are conserved, with the consensus TGAACC in -35 and CGT in -10, although other variations are possible in some subgroups. These consensus promoter motifs agree with reports of original ECF17 (Staroń et al., 2009) and with experimental data from SigL (Hahn et al., 2005) and SigU (Gordon et al., 2008).
6. **Summary:** ECF17 is regulated by anti-σ factors. The conservation of the promoter motifs indicates that it auto-induces its expression in most of the cases. Indeed, this is the case for SigL (Hahn et al., 2005) and SigU (Gordon et al., 2008). It is possible that proteases encoded near ECFs from ECF17 are in charge of the degradation of their cognate anti-σ factor. The functions of members of ECF17 are diverse, and it does seem to be the main factor that contributes to their sequence similarity.

**ECF18**

1. **General description:** This group is almost exclusively composed of proteins of original group ECF18 (95.33%). Proteins from ECF18 belong exclusively to Proteobacteria and have two highly conserved cysteine residues in the $\sigma_4$ domain.
2. **Special features:** Like ECF214, members of ECF18 are associated with RskA-like putative AS factors encoded in position +1. These putative AS factors contain one transmembrane helix (96.64%) and a putative zinc-finger in some cases.
3. **Genomic context conservation:** Other than the AS factor, the only conserved proteins encoded in the genetic context of several groups are a DUF4394-containing protein (-1 of ECF18s15 and ECF18s12) and a fasciclin domain-containing protein (-1 of ECF18s2 and +2 of ECF18s3 and ECF18s11). Faciclin domain-containing proteins are usually transmembrane or cell-surface proteins involved in cell adhesion (Moody & Williamson, 2013). Other conserved domains include a SelR domain (+2 of ECF18s11), a DUF4331 (+2 of ECF18s13), a DUF3455 (-1 of ECF18s1), a flagellar FliJ protein (-1 of ECF18s9), an ATP synthase (-2 of

ECF18s9), a response regulator with a transcription regulator (-3 of ECF18s9), a mycolic acid cyclopropane synthetase (ECF18s14), a CheR methyltransferase (ECF18s9), a protein from FHIPEP family (ECF18s9), a haloacid dehalogenase (ECF18s11), an amino acid kinase (ECF18s11) and a 50S ribosome-binding GTPase (ECF18s11).

4. **Studied members:** RpoT from *Pseudomonas putida* (ECF18s1) is involved in resistance to organic solvents since it regulates an efflux pump (Duque et al., 2007). Another characterized member of this group, Rsph17029_3536 (ECF18s7) from *Rhodobacter sphaeroides,* induces the expression of the alanine synthetase *hemT* in response to two inputs. First, changes in reducing conditions sensed by the two cysteine residues in $\sigma_4$ domain release the ECF from the inhibition of it's AS factor, similarly as SigK from *M. tuberculosis* (Shukla et al., 2014); second, $C_4$-dicarboxylic acids inactivate its AS factor (Coulianos, 2018). The third characterized member of ECF18, LitS from *Burkholderia multivorans* (ECF18s6), is responsible for the light-dependent activation of genes for the biosynthesis cyclopropane-fatty-acyl-phospholipid and the photolyase PhrB2 for DNA repair (Sumi, Shiratori-Takano, Ueda, & Takano, 2018). Lastly, RpoP from *C. metallidurans* (ECF18s4) plays a role in nickel and cobalt resistance (Grosse et al., 2019).
5. **Promoter motif conservation:** In general, the target promoter motifs of ECF18 agree with the prediction for original ECF18 (Staroń et al., 2009), this is TGCATCTT in -35 and CGTA in -10. In general, the -35 elements are more conserved. Some promoter motifs correspond to the target of housekeeping σ factors (for example ECF18s24). The putative promoter motif of Rsph17029_3536 does not agree with the prediction for subgroup ECF18s7 since this ECF is not autoregulated (Coulianos, 2018). Nevertheless, genes regulated by LitS in *B. multivorans* contain the same motif as predicted for ECF18 (Sumi et al., 2018). This motif is not conserved in ECF18s6, the subgroup of LitS, indicating that even though LitS might not be autoregulated, the target promoter motif still matches the predictions of the most conserved motif of ECF18.
6. **Summary:** ECF18 has several regulatory layers - on the one hand it is regulated by a single transmembrane AS factor similar to RskA from *M. tuberculosis* and original ECF18 (Staroń et al., 2009); on the other hand the two cysteines in its $\sigma_4$ domain may form a disulfide bridge that unbinds under reducing conditions, releasing the ECF from its AS factor as in the case of SigK and its AS factor RskA (Shukla et al., 2014). Not all the ECFs from ECF18 are auto-induced.

**ECF19**

1. **General description:** ECF19 is composed by a mixture of proteins from original groups ECF19 (56.46%), ECF18 (1%), ECF126 (3.66%) and ECF34 (2.72%). The taxonomic distribution of this group is diverse and subgroup-specific, with proteins coming from Actinobacteria (86.21%), Cyanobacteria (5.65%), Firmicutes (1.95%), Chloroflexi (0.51%), Proteobacteria (1.65%), Verrucomicrobia (0.51%), Deinococcus-Thermus (1.54%), Acidobacteria (1.03%) and Planctomycetes (0.62%) and Gemmatimonadetes (0.10%).
2. **Regulation:** The genetic context of ECF19 contains a conserved putative AS factor in position +1. Typically this AS factor has a RskA-like domain with or without a zinc-finger and one transmembrane helix (79.71%). In subgroup ECF19s19, the putative AS factor is a soluble protein with a ChrR-like CLD, as in members of ECF11. Moreover, the putative AS factor in subgroups ECF19s7 and ECF19s18 is a soluble protein with a putative zinc-finger fused to the N-terminal domain of a mycothiol maleylpyruvate isomerase. Interestingly, these two subgroups contain an anti-anti-σ (AAS) factor with a STAS domain in -1 (94.44% of the contexts in ECF19s7, 40% of ECF19s18). Only 21.82% of the ECFs from ECF19s4 contain putative AS factors, indicating that the cognate AS factors of the rest of the members of ECF19s4 are encoded somewhere else in the genome. Interestingly, subgroup ECF19s4 contains a MerR HTH family regulatory protein (64.15%) in position -1, which induces the transcription of the ECF LitS from *S. coelicolor* (ECF19s4) in response to light, leading to the transcription of the carotenoid biosynthesis pathway (Takano, Obitsu, Beppu, & Ueda, 2005).
3. **Genomic context conservation:** The rest of the proteins in the context of ECF19 are only conserved in a subgroup-specific manner. We could find several domains conserved within both ECF19s7 and ECF19s18, such as a bifunctional DNA primase/polymerase N-terminal domain, a protein with a conserved ACT domain fused to a formyltransferase domain. Other conserved proteins in include a mttA/Hcf106 protein (-1 of ECF19s27), a sortase (-1 of ECF19s5), a DUF4397 (-2 of ECF19s5), a SsgA protein involved in sporulation and cell division (-1 of ECF19s12), a FeoA Iron dependent repressor (+8 of ECF19s18), a SIS domain-containing protein (+9 of ECF19s18), a cytochrome C biogenesis protein (-2 of ECF19s9), a flavin-containing amine oxidoreductase (ECF19s4), an ABC transporter with a DUF4162 (ECF19s13) and a MbtH-like protein (ECF19s13).
4. **Studied members:** A member of ECF19, SigK (ECF19s2) from *Mycobacterium* spp., has been experimentally addressed. SigK regulon includes *sigK*, the antigens MPT70 and MPT83, and genes for the

synthesis of lipids and oxidative stress response (Saïd-Salim, Mostowy, Kristof, & Behr, 2006; Sklar, Makinoshima, Schneider, & Glickman, 2010; Veyrier, Saïd-Salim, & Behr, 2008). Antigens MPT70 and MPT83 are induced when *M. tuberculosis* resides inside macrophages (Schnappinger et al., 2003). The AS factor of SigK, RskA, is dysfunctional in *M. bovis*, which makes the production of MPT70 and MPT83 higher in *M. bovis* resulting in a higher virulence of *M. bovis* respect to *M. tuberculosis* (Medina, Ryan, LaCourse, & North, 2006; Saïd-Salim, Mostowy, Kristof, & Behr, 2006). SigK is the first σ factor described to be dually activated by the dissociation of a disulfide bridge in $\sigma_4$ domain due to reducing conditions (Shukla et al., 2014) and a signal that inactivates its AS factor via RIP mediated by the intramembrane protease Rip1 (Sklar, Makinoshima, Schneider, & Glickman, 2010). Even though this is the first group where the role of the two cysteines of the $\sigma_4$ domain was described, this is not one of the defining features of ECF19 since it only appears in some subgroups (ECF19s1, ECF19s2, ECF19s6, ECF19s11, and ECF19s13).

Yet other members of ECF19 have been experimentally addressed, SigK from *Rhodococcus hoagii* (ECF19s2). This protein is upregulated during macrophage infection (Rahman, Parreira, & Prescott, 2005). Lastly, LitS from *S. coelicolor* (ECF19s4) is involved in the transcription of the carotenoid biosynthesis pathway when induced by LitR, a MerR-type transcriptional regulator encoded in -1, in response to light (Takano et al., 2005). LitS might also be regulated by a putative AS factor, LitB, encoded in position +2 and with a RskA domain (Takano et al., 2005). Indeed, 21.82% of the ECFs from ECF19s4 encode a RskA-like domain in their genetic context.

5. **Promoter motif conservation:** The predicted target promoter motif of members of ECF19 is only conserved at a subgroup level and among neighboring subgroups. A typical pattern is ATC in -35, and CG or TG in -10. The promoter motifs of LitS (GCATG -35 and CGGAAG -10 (Takano et al., 2005)) do not agree with the promoter predicted by our pipeline even though LitS is autoregulated (Takano et al., 2005), but agree with the motifs for SigK in *Mycobacterium tuberculosis* (Veyrier, Saïd-Salim, & Behr, 2008).
6. **Summary:** Members of ECF19 are regulated by putative AS factors, most of which contain one transmembrane helix and an RskA-like domain, as in the case of original ECF19 (Staroń et al., 2009). Some subgroups have soluble AS factors (ECF19s19, ECF19s18 and ECF19s7). Of those, ECF19s7 and ECF19s18, are associated with AAS factors encoded in -1 and contain a soluble ZAS fused to the N-terminal domain of a mycothiol maleylpyruvate isomerase. Since members of these groups have homology

to original ECF126 and show a different mechanism that the rest of the members of ECF19, we consider ECF19s7 and ECF19s18 specialized members of ECF19. ECF19 illustrates how it is possible to unify different original groups from different taxonomic groups but with similar characteristics based on protein similarity.

**ECF20 (original classification; no longer present)**

Members of original ECF20 are scattered across several groups from the new classification. The largest contribution of original ECF20 is to new groups ECF289 (42.27%), ECF290 (95.41%) and ECF291 (33.76%).

**ECF21**

1. **General description:** ECF21 has homology to original ECF21 (62.35%). Proteins in the base of ECF21's clade are present in Bacteroidetes (99.62%). Subgroups ECF21s12 and ECF21s29 are fused to the putative AS factors with an outer membrane beta-barrel, which are encoded in +1 for the rest of the subgroups.
2. **Anti-σ factor:** Members of ECF21 are mainly regulated by AS factors with one inner membrane transmembrane helix (71.31%) located in position +1 (except ECF21s12). These putative AS factors contain an outer-membrane beta-barrel in most of the cases, although, in some subgroups ECF21s9, ECF21s16, and ECF21s28, the AS factor and the outer membrane beta-barrel are decoupled and located in different proteins encoded in +1 and +2, respectively.
3. **Genomic context conservation:** The conservation of the genetic neighborhood is extensive in some subgroups and involves a protein from the radical SAM superfamily (ECF21s35 and +1 of ECF21s12), an oxidoreductase (ECF21s28, +2 of ECF21s12), a DUF4840 (-1 of ECF21s10), a polyprenyl synthetase (ECF21s1 and -1 of ECF21s26), a DNA primase (+3 of ECF21s26), a response regulator with a LuxR repressor (ECF21s1 and +4 of ECF21s26), a NAD synthase (ECF21s1 and +5 of ECF21s26), a RecA protein (-1 of ECF21s3 and ECF21s6), a FKBP-type peptidyl-prolyl cis-trans isomerase (+2 of ECF21s35), an adenylosuccinate lyase (ECF21s28), a thiolase (ECF21s13), two fumarase C (ECF21s16), a coenzyme A transferase (ECF21s6 and ECF21s3), a thioesterase (ECF21s3), a rhodanase (ECF21s3), an acyltransferase (ECF21s3), a GldH lipoprotein (ECF21s3), a haloacid dehalogenase (ECF21s32), a M42 glutamyl aminopeptidase (ECF21s35), an uracil phosphoribosyltransferase (ECF21s35), a phosphoribosyl transferase (ECF21s35) and a sodium/calcium exchanger (ECF21s35).
4. **Promoter motif conservation:** The most common predicted target promoter motif is GCAACC in -35 and CGTCT in -10, in agreement with original ECF21 (Rhodius et al., 2013).

5. **Summary:** ECF21 might be regulated by AS factors encoded in +1. These AS factors contain one transmembrane helix and, in some cases, are fused to an outer membrane beta-barrel. In other instances, the outer membrane beta-barrel is encoded in the +2. No members of ECF21 have been functionally addressed.

**ECF22**

1. **General description:** ECF22 has homology to original ECF22 (88.1%). Proteins from this group are present in Bacteroidetes (78.81%), Proteobacteria (18.14%), Verrucomicrobia (1.19%), Planctomycetes (1.02%) and Acidobacteria (0.68%).
2. **Anti-σ factor:** Putative AS factors of ECF22 are encoded in +1 and usually contain DUF2207 and four (48.06%) TM helices.
3. **Genomic context conservation:** Other proteins conserved in the genetic context of members of ECF22 are an alpha/beta hydrolase (-1 of ECF22s16 and ECF22s4), σ-54 factor (ECF22s8) and two proteins from ABC transporters (-2 and -3 of ECF22s22).
4. **Promoter motif conservation:** Promoter motifs are conserved and contain TGTGATTTT in -35 and GCGAAT(C/A)AT in -10, in agreement with original ECF22 (Rhodius et al., 2013) and expanding these motifs.
5. **Summary:** ECF22 could be regulated by 4-transmembrane helix AS factors located in +1. This expands the knowledge of original ECF22, where putative AS factors were not identified (Staroń et al., 2009). This group seems to be autoregulated since the promoter motifs are conserved.

**ECF23**

1. **General description:** Members of ECF23 are homologous to original ECF23 (55.79%) and are present in Firmicutes (100%).
2. **Anti-σ factor:** Members of this group contain an AS factor encoded in position +1 with a DUF4179 in most of the cases. Proteins encoded in this position contain one transmembrane helix (88.7%). The genetic context conservation does not extend beyond the AS factor.
3. **Promoter motif conservation:** The only conserved predicted target promoter is TGATAG in -35 and CGTATTA in -10, although it is only evident in ECF23s3, ECF23s9, ECF23s6, and ECF23s7.
4. **Summary:** Members of ECF23 are regulated by single transmembrane AS factors encoded in +1 that contain a DUF4179. The genetic context conservation does not extend beyond the AS factor.

**ECF24 (original classification; no longer present)**

Members of original group ECF24 are present in ECF238, together with members of original ECF44. 84.87% of the members of ECF238 have homology to ECF24.

**ECF25**

1. **General description:** Two subgroups of ECF25, ECF25s1 and ECF25s11, have homology to original ECF25 (30.6%) and ECF20 (0.19%). ECF25 is one of the most diverse ECF groups in terms of taxonomic origin of the sequences. Proteins from ECF25 are present mainly in Cyanobaceria (ECF25s1 and ECF25s11, 50%), but also in Firmicutes (11.88%), Deltaproteobacteria (15.84%), Spirochaetes (ECF25s3, 2.97%), Fibrobacteres (ECF25s13, 0.5%), Nitrospirae (ECF25s12, 0.5%), Deinococcus-Thermus (ECF25s5, 6.44%), Chloroflexi (ECF25s17, ECF25s18 and ECF25s19, 6.44%), Gemmatimonadetes (ECF25s4, 1%), Ignavibacteriae (ECF25s9, 1%), Acidobacteria (1%) , Bacteroidetes (1.98%), and Calditrichaeota (0.5%).
2. **Anti-σ factor:** Similarly to original ECF25 (Staroń et al., 2009), proteins from ECF25 contain a putative AS factor in position +1, which are mainly single membrane-pass (74.75%). Nevertheless, variants with two or three transmembrane helices are possible. For instance, putative AS factors from Firmicutes order Thermoanaerobacterales (ECF25s14), and Chloroflexi (ECF25s16, ECF25s18, ECF25s19) contain two transmembrane helices, and proteins ECF25s20 contain three transmembrane helices. These putative AS factors typically contain a zinc-binding domain and/or a DUF4384 or a DUF4349.
3. **Genomic context conservation:** Other proteins conserved in the genetic neighborhood of ECF25 are a transglycosylase SLT from phages, T2SS, T3SS or T4SS (ECF25s2), a DNA polymerase family A with 5'-3' exonuclease activity involved in DNA repair (ECF25s2), a dephospho-CoA kinase (ECF25s2), a DUF1800 (ECF25s5) and an iron-dependent repressor (ECF25s5).
4. **Studied members:** The only member of ECF25 experimentally addressed, SigG from *Nostoc punctiforme* (ECF25s1) is in charge of the defense against cell wall stress arising from external sources (EDTA, heat, lysozyme, cold) or cellular differentiation (Bell, Lee, & Summers, 2017). SigG is regulated by a single membrane pass AS factor SapG.
5. **Promoter motif conservation:** The predicted target promoter motifs of ECF25s1 and ECF25s2 are GGAAC in -35 and GTC in -10. This promoter motif is similar to original ECF25 (Rhodius et al., 2013) and the binding motif of SigG in *N. punctiforme* (Bell et al., 2017).

6. **Summary:** ECF25 is regulated by AS factors located in +1 and seem to be involved in cell wall homeostasis.

**ECF26**

1. **General description:** Proteins from ECF26 have homology to proteins from the original group ECF26 (94.35%). Group ECF26 is present in Proteobacteria (99.87%), mainly Alpha- and Betaproteobacteria, and Bacteroidetes (0.13%) from genus *Bacteroides*.
2. **Anti-σ factor:** Members of ECF26 contain a putative AS factor in position +1, as described for the original group ECF26 (Staroń et al., 2009). Putative AS factors of ECF26 contain one TM helix (85.83%) and a zinc finger.
3. **Genomic context conservation:** Position -1 is usually conserved, and its specific function depends on the position of the subgroup in the phylogenetic tree of ECF26; we could find catalases (ECF26s1), subtilases (ECF26s10, ECF26s4, ECF26s27 and ECF26s14), chitin-binding proteins (ECF26s2), enoyl-(acyl carrier protein) reductases involved in fatty acid biosynthesis (ECF26s12), and a conserved protein with a secreted repeat domain (ECF26s6, ECF26s16, ECF26s17 and ECF26s9). Other domains conserved in the context of members of ECF26 are the SsrA-system RNA-binding protein SmpB (ECF26s9), the domain of the inhibitor of apoptosis-promoting Bax1 involved in calcium leakage across the membrane (ECF26s7 and ECF26s22), the prokaryotic cytochrome b561 from the electron transport chain (ECF26s1), a LysR-like transcription regulator (ECF26s8), the small subunit of an acetolactate synthase (ECF26s6), the beta subunit of the nitrile hydratase (ECF26s6), an aldo/keto reductase (ECF26s6), an UvrD-like helicase (ECF26s6), a LysE type translocator (ECF26s6), a DUF1194-containing protein (ECF26s12), a 17 kDa outer membrane surface antigen (ECF26s12), a pyridoxine 5'-phosphate oxidase (ECF26s12), a histidine phosphatase (ECF26s12), a chorismate synthase (ECF26s12), a DnaJ-like protein (ECF26s12), an ubiquitin from the RnfH family (ECF26s9), an IMP dehydrogenase/GMP reductase (ECF26s9) and a polyketide cyclase/dehydrase (ECF26s9).
4. **Studied members:** SigE from *Starkella novella* (ECF26s28) regulates the expression of *sorAB*, located in +2 and +3, which encode a soluble sulfite:cytochrome c oxidoreductase involved in thiosulfate oxidation independent of a membrane-associated thiosulfate-oxidizing complex during chemolithotrophy (Kappler, Friedrich, Trüper, & Dahl, 2001). The protein encoded downstream of SigE has been proposed to be a membrane-bound AS factor with a short cytoplasmic region (Kappler et al., 2001). PrtI (ECF26s1), present

in *Pseudomonas* spp., regulates the temperature-dependent production of a protease in *P. fluorescens* WH6 (Okrent et al., 2014). *Sinorhizobium meliloti* encodes four members of ECF26, RpoE1 (ECF26s6), RpoE3 (ECF26s29), RpoE4 (ECF26s34) and RpoE6 (ECF26s14). RpoE1 and RpoE4 are involved in detoxification of disulfide compounds, and all are regulated by one transmembrane helix AS factors encoded in +1 (Lang et al., 2018).

5. **Promoter motif conservation:** The promoter motifs are diverse but usually follow the pattern GGAATAAA in -35 and GTT in -10 in agreement with previous reports (Rhodius et al., 2013). This conservation indicates the autoregulatory role of members of ECF26. Indeed, SigE from *S. novella* is self-induced, although its described promoter motif (Kappler et al., 2001) does not entirely agree with the predicted target promoter of ECF26. ECFs from group ECF26 in *S. meliloti* are autoregulated (except RpoE3), and their binding motif agrees with the predictions for ECF26, although the specific predicted target promoter motifs of their subgroups might vary (Lang et al., 2018).

**ECF27**

1. **General description:** Members of ECF27 are homologous to members of original ECF27 (90.2%) and ECF20 (0.06%), and are present in Actinobacteria (100%).
2. **Anti-σ factor:** Position +1 contains a putative AS factor with one transmembrane helix (78.65%) and with a zinc-binding domain in ECF27s1 or non-conserved domains in other groups.
3. **Genomic context conservation**: As in the original group ECF27 (Staroń et al., 2009), the genetic context of members of ECF27 is extensively conserved. Position +2 encodes a pyridine nucleotide-disulfide oxidoreductase; position +3 encodes a thioredoxin; position +4 a N-acetylmuramoyl-L-alanine amidase with a peptidoglycan binding domain or an acetyltransferase of GNAT family (ECF27s1); position -1 contains a MviN-like protein (Pfam: MVIN) (-2 in ECF27s1 and ECF27s2) involved in flipping the lipid II peptidoglycan precursor to the periplasmic surface of the inner membrane in *Salmonella typhimurium, E. coli* and other gram-negative bacteria (Ling, Moore, Surette, & Woods, 2006). Furthermore, ECF27s1 contains a protein kinase in position +1, a sugar-specific transcription regulator TrmB in +5, an AAA domain-containing protein in +6, an rRNA small subunit methyltransferase G in +7, a KH domain-containing protein fused to a R3H domain in -8, possibly functioning as nucleic acid binding protein, a 60Kd inner membrane protein in +9, a polymerase A in charge of DNA repair and replacing Okazaki fragments with DNA in -4 and also conserved in ECF27s2 and ECF27s4, a major facilitator superfamily

protein, and a PadR transcriptional regulator. There is a NUDIX domain protein in -3 of ECF27s5 and in ECF27s4.

4. **Studied members:** SigM, from M*ycobacterium smegmatis* (ECF27s4), is required for receiving DNA for conjugation (Clark et al., 2018). SigM is released from its membrane-bound AS factor upon detection of the donor cell and induces the expression of cell wall hydrolases, nucleases and the ESX-4 secretion apparatus (or type VII secretion system), in charge of connecting the donor and the acceptor cells to accomplish the DNA integration (Clark et al., 2018). The genetic context conservation of ECF27 agrees with this function, suggesting that other members of ECF27 have a similar function. Another member of ECF27, SigM from *S. coelicolor* (ECF27s1), plays a negative role in differentiation (Mao et al., 2009).
5. **Promoter motif conservation:** Promoter motifs are not conserved, indicating the lack of direct regulation of members of ECF27 over their expression. Indeed, this is the cases of SigM (Clark et al., 2018). The promoter motifs of original ECF27 (Staroń et al., 2009) are not observed in new ECF27, which agrees with the lack of activity of these promoters in a heterologous organism (Rhodius et al., 2013).
6. **Summary:** Due to the conservation of the proteins in the genetic neighborhood of members of ECF27, it is possible that ECF27 might be required for recipient cells during conjugation in Actinobacteria, as in the case of SigM in M*ycobacterium smegmatis* (ECF27s4).

**ECF28**

1. **General description:** This group has homology to the original group ECF28 (91.24%), and it is present in Proteobacteria from class Proteobacteria (99.34%) from class Gammaproteobacteria and Cyanobacteria (0.66%).
2. **Anti-σ factor:** ECFs from ECF28 are associated with putative AS factors in position +1 with a DUF3379 and one transmembrane helix (93.21%). This protein is not present in ECF28s3.
3. **Genomic context conservation:** Other conserved proteins encoded in the genetic context of members of ECF28 are a thiolase (ECF28s1 and ECF28s2), an outer membrane protein transport protein (OMPP1/FadL/TodX) (ECF28s1 and ECF28s2) involved in degradation of aromatic hydrocarbons (Pfam: Toluene_X), a 3-hydroxyacyl-CoA dehydrogenase (ECF28s2) and a von Willebrand factor type A (ECF28s1 and ECF28s2).
4. **Promoter motif conservation:** The predicted target promoter motifs are not conserved.

5. **Summary:** Likewise original ECF28 (Staroń et al., 2009), ECF28 might be regulated by a membrane-bound AS factor with a DUF3379. The conservation of the genetic context of members of ECF28 indicates that they are involved in the degradation of aromatic compounds.

**ECF29**

1. **General description:** ECF29 has homology to original groups ECF29 (88.98%). Members of ECF29 are present in Proteobacteria (83.64%) and Bacteroidetes (16.36%).
2. **Special features:** ECF29 contains a conserved RCE/D motif in a short (~30aa) C-terminal extension that could be in charge of the modulation of its activity, either participating in sensing or repression of the ECF σ factor. Since the ECFs from ECF29 are soluble, they could sense cytoplasmic signals that could be related to ROS, disulfide or heavy metal stress.
3. **Anti-σ factor:** Proteins from ECF29 do not contain any evident putative AS factor.
4. **Genomic context conservation:** No conserved elements were found in the genetic context of members of ECF29.
5. **Promoter motif conservation:** Predicted target promoters of ECF29 usually contain GGGAACCT in -25 and CATCCAAT in -10, in agreement with original ECF29 (Rhodius et al., 2013). In subgroups ECF29s18 and ECF29s14 from Bacteroidetes, the putative target promoter motifs are TTAGAT (-35) GTTACA in -10.
6. **Summary:** ECF29 might be regulated by a short C-terminal extension with a conserved RCE/D motif. The predicted target promoter motifs are conserved, indicating the positive autoregulation of ECFs from ECF29. Functional characterization of members of ECF29 would shed light on the function of members of ECF29.

**ECF30**

1. **General description:** Proteins from ECF30 are homologous to original group ECF30 (94.68%) and are present in Firmicutes (99.86%).
2. **Anti-σ factor and genomic context conservation:** ECF30 is associated with single TMH AS factors with DUF4179 or DUF3298 fused or not to DUF4163 encoded in +1.
3. **Genomic context conservation:** The largest subgroup, ECF30s1, has an extensively conserved genetic context with the delta subunit of the DNA polymerase III in +2, the ribosomal protein S20 in +3, a metallo-beta-lactamase fused to a competence protein (DNA transport across the membrane) and a DUF4131, a germination protease and a cytidine and deoxycytidylate deaminase.

4. **Studied members:** Described members of ECF30 belong to ECF30s6 (SigV from *Bacillus subtilis* and SigV from *Enterococcus faecalis*), ECF30s11 (CsfT from *Clostridium difficile*), ECF30s48 (CsfV from *Clostridium difficile*) and ECF30s3 (SigW from *Bacillus thuringiensis*). SigV in critical for lysozyme resistance in both *B. subtilis* and *E. feacalis* due to its regulation of genes for the resistance to lysozyme, such as *oatA*, a gene located in +2 that encodes an O-acetylase that acetylates peptidoglycan (Lewerke, Kies, Müh, & Ellermeier, 2018; E. C. Woods & McBride, 2017). This protein is conserved in ECF30s6, where 80% of the proteins in ECF30s6 contain one acetyltransferase, but it is not conserved in other subgroups of ECF30. SigV from *B. subtilis* is activated when membrane proteins are delocalized (Omardien et al., 2018). The degradation of its AS factor, RsiV, occurs when it binds to lysozyme. This degradation is catalyzed by a signal peptidase as site-1 protease and RasP as site-2 protease (Helmann, 2016; Lewerke et al., 2018). The degradation is hampered in the absence of lysozyme by an amphipathic helix encoded in the position of DUF4179 (Lewerke et al., 2018). CsfT and CsfV play a role in antimicrobial resistance, and they are induced by bacitracin and lysozyme (E. C. Woods & McBride, 2017). The protease PrsW releases CsfT from its AS RsiT (E. C. Woods & McBride, 2017). In the case of CsfV, bacitracin binds directly to the AS factor RsiV (E. C. Woods & McBride, 2017). SigW from *B. thuringiensis* regulates the expression of the β-exotoxin I independently of the cry plasmid and it is encoded with its cognate AS factor (position +1) and EcfY (position +2), a negative regulator of the expression of SigW (Espinasse, Gohar, Lereclus, & Sanchis, 2004) conserved in the genetic context of members of ECF30s3.
5. **Promoter motif conservation:** Putative target promoters have differences in different subgroups. They usually contain TGCAACA or TGAAACTTT in -35 and CGTC or CTCTAAT in -10. These motifs agree with the experimentally obtained auto-inducible target promoters of SigV in *B. subtilis* and SigW from *B. thuringiensis* (Espinasse et al., 2004; Helmann, 2016), and with original group ECF30 (Staroń et al., 2009).
6. **Summary:** ECF30 responds to cell wall damage caused by lysozyme or other antimicrobials. ECFs from ECF30 are regulated by AS factors that might be inactivated via RIP and protected from degradation by amphipathic helices since most of the AS factors contain a DUF4179, as in the case of some described members of this group. The final genes activated by the ECF might vary for different subgroups. ECFs from this group seem to auto-induce their expression.

**ECF31**

1. **General description:** Members of ECF31 have homology to proteins from original ECF31 (91.1%) and are exclusively present in Firmicutes.
2. **Anti-σ factor:** Members of ECF31 might be regulated by putative AS factors encoded in +1. These AS factors contain two transmembrane helices (84.35%) with a DUF3545 and a DUF2207 that covers the total length of the sequence. Members of ECF31s5 (12.5%) contain a cytoplasmic zinc-finger in their putative AS factor. Proteins encoded in +2 are conserved and contain two transmembrane helices. Among the domains that these proteins contain we could find a virulence factor BrkB domain or a MAP17 domain, known to interact with PDZ domains (Kocher, Comella, Tognazzi, & Brown, 1998).
3. **Genomic context conservation:** Other than the AS factor, members of ECF31 contain several genes that encode proteins from ABC transporters (positions +3 and +4 in ECF31s1 and ECF31s2) and the N-terminal domain of a phospholipase D-nuclease (Pfam: PLDc_N) (position +5 in ECF31s1 and ECF31s2) in their genetic context. Other domains conserved in the genetic context of members of ECF31 included the C subunit of Glu-tRNAGln amidotransferase (+7 of ECF31s2), an amidase (+8 of ECF31s2), GatB (+9 of ECF31s2), a protein with an acetyltransferase (GNAT) domain (+10 of ECF31s2), a metallo-beta-lactamase (-2 of ECF31s2), a thioredoxin (-3 of ECF31s2), a SNARE-associated Golgi protein (-4 of ECF31s2).
4. **Studied members:** Characterized members of ECF31 include SigY from *Bacillus subtilis* (ECF31s1). SigY's activity is dependent on YxlC, the putative AS factor encoded in +1, and YxlD encoded in +2 (Cao et al., 2003; Tojo et al., 2003; Yoshimura, Asai, Sadaie, & Yoshikawa, 2004). SigY is induced under nitrogen starvation (Tojo et al., 2003) and it is needed for maintaining the SPβ prophage, which encodes sublacin and its resistance cassette (Mendez, Gutierrez, Reyes, & Márquez-Magaña, 2012). When active, SigY induces only its expression and the one of the protein of unknown function YbgB (Cao et al., 2003). Homologs of YxlD are conserved across ECF31.
5. **Promoter motif conservation:** Promoter motifs are conserved and contain TGAAC in -35 and CGT in -10, in agreement with original ECF31 (Staroń et al., 2009).
6. **Summary:** Members of ECF31 might be regulated by AS factors with two transmembrane helices encoded in +1. The function of members of ECF31 might be related to the presence of conserved transmembrane proteins in +2, as in the case of SigY from *B. subtilis* (Cao et al., 2003; Tojo et al., 2003; Yoshimura et al.,

2004). Other proteins that could participate in ECF31 activity are ABC transporters and phospholipase D-nuclease, encoded in their genetic context.

**ECF32**

1. **General description:** Members of ECF32 are present in Proteobacteria (100%) from orders Enterobacterales and Pseudomonadales and have homology to original ECF32 (77.73%).
2. **Regulation:** Subgroups ECF32s1, ECF32s2 and ECF32s5 do not contain any proteins from representative/reference organisms. The remaining members of ECF32 encode a histidine kinase and a response regulator in their genetic context. The number of histidine kinases and receiver domains per ECF genetic contexts ranges from an average of 0.6 in ECF32s6 to 1 in ECF32s8 and ECF32s7. Histidine kinases are located in position +1 and response regulators in position +2 and fused to a LuxR transcriptional regulator. Interestingly, the histidine kinase does not contain any predicted transmembrane helix. Indeed, members of ECF32 such as HrpL from *Erwinia herbicola* (ECF32s4) and *Pseudomonas syringae* (ECF32s2) are transcriptionally regulated by 2CSs (Nizan-Koren, Manulis, Mor, Iraki, & Barash, 2003; Yu et al., 2014).
3. **Genomic context conservation:** The conservation of the genetic context of ECF32 is extensive and includes genes for the expression of T3SS, as observed for original ECF32 (Staroń et al., 2009). Indeed, members of ECF32 such as HrpL from *E. herbicola* (ECF32s5) and *P. syringae* (ECF32s2) activate this cluster of T3SS genes (*hrp* gene cluster) as a requirement for plant infection (Nizan-Koren et al., 2003; Yu et al., 2014).
4. **Promoter motif conservation:** Information content of predicted target promoter motifs is low, although several subgroups contain TTGC in -35 and GGC in -10. Some proteins from ECF32 do not seem to auto-regulate their expression, but rather to be regulated by members of σ54 (Nizan-Koren et al., 2003). These motifs could be part of the σ54 binding site. The funding ECF classification extracted the target promoter of members of ECF32 from upstream of the *hpr* gene cluster as GGAAC in -35 and CCAC in -10 (Staroń et al., 2009).
5. **Summary:** Similarly to original ECF32 (Staroń et al., 2009) ECF32 is involved in bacteria plant infection and is regulated by 2CSs encoded in their genetic context. Members of ECF32 regulate the expression of the *hrp* gene cluster, which encodes a T3SS involved in the infection.

**ECF33**

1. **General description:** Members of ECF33 are present in Proteobacteria (100%) and have homology against original ECF33 (44.11%).
2. **Anti-σ factor:** Members of ECF33 might be regulated by putative AS factors encoded in +1 with one transmembrane helix (73.64%) and a zinc finger.
3. **Genomic context conservation:** Other than this protein, conserved domains in the genetic context of members of ECF33 include a subtilase (+2 of ECF33s3 and ECF33s2) and a binding-protein-dependent transport system inner membrane component (ECF33s3), which is usually present in the periplasmic transporters of ABC transport systems.
4. **Studied members:** One described member of ECF33, EcfQ from *Bradyrhizobium japonicum* (ECF33s1), is responsible for singlet oxygen resistance (Masloboeva et al., 2012).
5. **Promoter motif conservation:** The target -35 element is extensively conserved (TGAACCTTT), and the -10 element varies amongst subgroups, although an AC is usually conserved. This promoter matches previous work on original ECF33 (Rhodius et al., 2013) and agrees with experimentally described target promoter of EcfQ (Masloboeva et al., 2012).
6. **Summary:** ECF33 might be regulated by zinc-finger AS factors located in position +1 with one TM helix (73.64%). Members of this group could be responsible for singlet oxygen resistance, as described for EcfQ of *B. japonicum* (Masloboeva et al., 2012). The conservation of the target promoter elements indicates that members of ECF33 usually are self-regulated.

**ECF34 (original classification; no longer present)**

Members of original group ECF34 are merged in new group ECF19. 2.72% of the proteins from new ECF19 have homology to original ECF34.

**ECF35**

1. **General description:** Members of ECF35 have homology to members of original ECF35 (6.13%) and are present in Proteobacteria (100%).
2. **Anti-σ factor:** Members of ECF35 might be regulated by a putative AS factor encoded in position +1. This AS factor contains one transmembrane helix (67.68%, ~100% when looking at an MSA) and periplasmic tetratricopeptide repeats in some cases. Members of subgroups ECF35s17, ECF35s12, ECF35s1, ECF35s6, ECF35s14, ECF35s20 and ECF35s4 contain an average of ~1 protein with a DUF3520 in their genetic

context, usually in positions -1 of -2. These proteins were identified as putative AS factors in groups ECF226 and ECF289, where they are fused to von Willebrand factors. Nevertheless, DUF3520-containing proteins do not bear any transmembrane helix when associated with members of ECF35.

3. **Genomic context conservation:** Other conserved domains include a UPF0029 (+2 of ECF35s2), the C-terminal domain of a TonB protein (-1 of ECF35s13), a diguanylate cyclase with an EAL and a GAF domain (-1 of ECF35s2), an oligopeptide/dipeptide ABC transporter (ECF35s11), a protein with a transglycosylase SLT domain (ECF35s7) and an AIR carboxylase (ECF35s7).
4. **Studied members:** A characterized member of ECF35 is SbrI from *Pseudomonas aeruginosa* (ECF35s1). SbrI is involved in cell envelope integrity in the control of swarming motility and biofilm formation. It is inhibited by the AS factor SbrR, encoded in +1 in the same operon (see review (Chevalier et al., 2018)).
5. **Promoter motif conservation:** Predicted target promoter motifs are subgroup-dependent. Subgroups ECF35s2 and ECF35s5 contain ACCC in -35 and CGT in -10, whereas subgroups ECF35s1 and ECF35s6 contain TAACCCG in -35 and CGTCT in -10. Indeed, this last motif agrees with the binding promoter motif of SbrI (Aires, Köhler, Nikaido, & Plésiat, 1999).
6. **Summary:** ECF35 is regulated by one-transmembrane helix AS factors encoded in +1. Members of this group might be involved in the control of swarming motility, biofilm formation and cell envelope integrity as in the case of SbrI from *P. aeruginosa* (ECF35s1).

**ECF36**

1. **General description:** Members of ECF36 share homology with members of original ECF36 (48.17%) and are present in Actinobacteria (100%).
2. **Anti-σ factor:** Members of ECF36 might be regulated by zinc-finger containing putative AS factors encoded in position -1. These putative AS factors contain four transmembrane helices (41.85%), which are more evident when looking at the MSA. Proteins from ECF36s3, the only subgroup with 100% homology to proteins from original ECF36, have an average of three transmembrane helices (four when looking at the MSA) in an N-terminal extension with a DUF2275 and are present exclusively in *Mycobacterium*. This extension has homology to the putative AS factors found in -1 in the remaining subgroups of ECF36. It is possible that the coding sequence of the putative AS factor fused to the N-terminus of members of ECF36s3 in the genus *Mycobacterium*, something common in proteins with physical interactions (Marcotte et al., 1999).

3. **Genomic context conservation:** The genetic context conservation of ECF36 does not extend beyond the ECF coding sequence and the putative zinc-finger AS factor. Subgroup ECF36s4 has a conserved catalase in position -1, a semialdehyde dehydrogenase and an amino acid kinase domain fused to an ACT domain that functions as regulatory ligand-binding domains (Chipman, 2001).
4. **Promoter motif conservation:** The putative target promoter of ECF36 is conserved and has the motifs GTC in -35 and GTTCCCG in -10, which differ from the motifs found for original ECF36 (Toyoda & Inui, 2016). One possibility is that these ECFs are not autoregulated. The data from the two characterized members of this group (Abdul-Majid et al., 2008; Sun et al., 2004; Toyoda & Inui, 2016) support this idea.
5. **Studied members:** Two ECFs from ECF36 have been described. SigC from *Mycobacterium tuberculosis* (ECF36s3) is required for virulence (Abdul-Majid et al., 2008; Sun et al., 2004) and its homologs are only present in pathogenic *Mycobacterium* spp. (Sachdeva, Misra, Tyagi, & Singh, 2009), whereas SigC from *Corynebacterium glutamicum* (ECF36s4) controls the synthesis of cytochrome bd and aa3, and represses the production of cytochrome bd1 under oxygen-limiting conditions (Toyoda & Inui, 2016). SigC from *M. tuberculosis* has been proposed to be regulated by an alternative mechanism that involves the occlusion of the Pribnow-binding region by intramolecular contacts between $\sigma_2$ and $\sigma_4$ regions (Thakur, Praveena, & Gopal, 2010). Indeed, SigC is the most abundantly expressed σ factor of *M. tuberculosis* but not the one found more often in complex with the RNAP core enzyme (Sachdeva et al., 2009).
6. **Summary:** In summary, members of ECF36 are regulated by transmembrane AS factors that are fused to the N-terminus of the ECF in bacteria from the genus *Mycobacterium*.

**ECF37**

1. **General description:** Members of ECF37 are homologous to proteins from original ECF37 (93.68%) and appear in Proteobacteria (99.32%) and Nitrospirae (0.68%).
2. **Anti-σ factor:** Members of this group have a broadly conserved genetic context – as in the case of original ECF37 (Staroń et al., 2009), proteins in +1 are putative AS factors and usually contain a DUF3619, or a RskA-like domain in ECF37s2. A conserved soluble DUF3106 domain-containing protein is present in +2.
3. **Genomic context conservation:** Other conserved proteins are a membrane-bound RDD protein (+3 of ECF37s1), a thiamine pyrophosphate enzyme (-1 of ECF37s1), the small subunit of an acetolactate synthase (-2 of ECF37s1), an acetohydroxy acid isomeroreductase (-3 of ECF37s1), a RNA 2'-O ribose methyltransferase (-1 of ECF37s3), a fructose-1-6-bisphosphatase (-2 of ECF37s3), an OmpA (-4 of

ECF37s3), a TolB protein fused to WD40 repeats (-5 of ECF37s3), the 4TM region of pyridine nucleotide transhydrogenase (-1 of ECF37s2), an AI-2E family transporter (ECF37s3), a biopolymer transport protein ExbD/TolR (ECF37s3), a formyl transferase (ECF37s3), an iron-sulfur cluster assembly protein (ECF37s2 and ECF37s5), a NUDIX domain-containing hydrolase (ECF37s2), an alanine dehydrogenase/PNT (ECF37s2), a DUF1631 (ECF37s2), an 5'-3' exonuclease (ECF37s2 and ECF37s5), the beta subunit of the NAD(P) transhydrogenase (ECF37s2), a nitroreductase (ECF37s2 and ECF37s5) and an asparaginase (ECF37s5).

4. **Studied members:** One member of ECF37, RpoR from *Cupriavidus metallidurans* (ECF37s1), is associated to cobalt and nickel resistance in the absence of RpoQ (ECF16s1) and is involved in the protection of iron-sulfur clusters (Grosse et al., 2019).
5. **Promoter motif conservation:** Predicted target promoter motifs are highly conserved and contain TGTCAACC in -35 and CGTT in -10. This promoter motif does not agree with the original group ECF37 (Rhodius et al., 2013). However, the promoter motifs of new ECF37 have larger information content and are coherent among different subgroups. Given this conservation, it is likely that members of ECF37 are autoregulated.
6. **Summary:** ECF37 contain a putative AS factor in +1 with one transmembrane helix and a DUF3619. Protein in +2 is also conserved and contains a DUF3106. Genomic neighborhoods are broadly conserved and are related to nucleic acid metabolism and transport across the membrane. Predicted target promoter motifs are conserved, suggesting an autoregulatory role of members of ECF37. The only described member of ECF37 is associated with heavy-metal resistance and protection of iron-sulfur clusters.

**ECF38**

1. **General description:** Members of ECF38 share homology with members of original ECF38 (73.09%) or original ECF39 (0.34%) and are present in Actiobacteria (100%).
2. **Anti-σ factor:** As in the case of original ECF38 and ECF39 (Staroń et al., 2009), the proteins in position +1 are putative AS factors, except in ECF38s4, which is associated with YCII-related domain protein in position -1 like proteins from ECF42. These putative AS factors contain one transmembrane helix (67.46%) and no Pfam domain.
3. **Genomic context conservation:** Other domains proteins encoded in the genetic context of members of ECF38 are a HIT protein (-1 of ECF38s5 and ECF38s6), a PhoH-like protein (-2 of ECF38s5), a UPF0054

protein (ECF38s5 and ECF38s6), a transporter fused to a DUF21 (ECF38s5 and ECF38s6), a SURF1 protein (+2 of ECF38s1), a DUF3097 (ECF38s5 and ECF38s6), DnaJ (ECF38s5 and ECF38s6), a protein with a DeoR-like helix-turn-helix domain fused to the C-terminal domain of HrcA (ECF38s5 and ECF38s6), a DUF4870 (ECF38s5), a RNA methyltransferase (ECF38s5 and ECF38s6), a 50S ribosome-binding GTPase (ECF38s5), a YbaB/EbfC protein (ECF38s1), an amino acid kinase with an ACT domain (ECF38s1), a DUF5063 (ECF38s1), a semialdehyde dehydrogenase (ECF38s1), RecR (ECF38s1) and a fructose-bisphosphate aldolase class-II (ECF38s4).

4. **Promoter motif conservation:** Predicted target promoter motifs are subgroup-specific. In subgroups ECF38s1, ECF38s6 and ECF38s9 there is AACC in -35 and TC in -10. These motifs agree with original ECF39 (Rhodius et al., 2013).
5. **Summary:** Members of ECF38 are associated with AS factors in position +1 and share the same characteristics as original ECF38 (Staroń et al., 2009).

**ECF39**

1. **General description:** ECF39 is one of the largest groups, and it is composed of 93 subgroups and 5,506 unique sequences, all from Actinobacteria. The members of this group could be classified against original ECF39 (93.15%) except one clade from the order Micrococcales.
2. **Regulation:** Original ECF39 is associated with 2CSs or membrane-bound AS factors (Staroń et al., 2009). We found that the largest subgroups of new ECF39 are associated with two-component systems – in ECF39s1 the response regulator (1.01 proteins with Pfam Response_reg per ECF) is in -1 and the histidine kinase (0.98 proteins with Pfam HATPase_c per ECF) in -2. In ECF39s3, the response regulator (1.12 copies per ECF) is in +2, and the histidine kinase (one copy per ECF) in +3 and in ECF39s2 only the response regulator (1.12 copies per ECF) is conserved. The response regulator contains a fused C-terminal domain of the transcriptional regulatory protein (Pfam: Trans_reg_C) in most cases. The remaining subgroups are likely regulated by putative AS factors encoded in +1 with one transmembrane helix (72.31%). Putative AS factors might contain sensing domains such as PASTA domains, in charge of sensing cell wall antibiotics, or WD40 beta propeller repeats, also present in other ECF groups such as ECF266 or the sensory domain of protein kinases of ECF62 or the C-terminal extension of some members of ECF57.

3. **Genomic context conservation:** The subgroups with the most conserved proteins in their genetic context are part of the clade regulated by 2CSs. They contain an AMP-binding enzyme (+1 of ECF39s2), a VanZ-like protein (-4 of ECF39s1) involved in resistance against cell surface-acting antimicrobials such as teicoplanin and encoded within the skin element of SigK in *Clostridium difficile* (E. C. Woods, Wetzel, Mukerjee, & McBride, 2018), an endonuclease with a Fe-S cluster involved in base excision DNA repair (-1 of ECF39s3), a DisA bacterial checkpoint controller (ECF39s3), a subtilisin inhibitor-like protein (ECF39s2), a DeoC/LacD family aldolase (ECF39s1), an adenosine/AMP deaminase (ECF39s1), an aldehyde dehydrogenase (ECF39s1) and a protein with a PspC domain (ECF39s1). Other proteins conserved in the genetic context of members of ECF39 include an amidinotransferase (ECF39s9), a ferritin-like protein (ECF39s9), an AAA ATPase domain fused to a MarR transcriptional regulator (ECF39s31), an aminotransferase class I and II (ECF39s35), an EPSP synthase (ECF39s35), an adenylosuccinate lyase (ECF39s14), a beta-eliminating lyase (-1 of ECF39s14) and BetI-type transcriptional repressor fused to a TetR repressor (ECF39s14).
4. **Studied members:** Characterized members of ECF39 include SigE (ECF39s3) and SigQ (ECF39s1) from *Streptomyces coelicolor*, and Sig25 from *Streptomyces avermitilis* (ECF39s1). They are associated with 2CSs encoded in their genetic context that promotes their expression. SigE (ECF39s3) is involved in resistance to antibiotics that target the peptidoglycan such as lysozyme (Wood & Ohman, 2015). Instead, Sig25 and SigQ (ECF39s1) induce the synthesis of antibiotics – Sig25 induces expression of oligomycin and represses avermectin (S. Luo et al., 2014), whereas SigQ induces actinorhodin, undecylprodigiosin and calcium-dependent antibiotic (D. Shu et al., 2008). SigQ is also involved in sporulation (D. Shu et al., 2008).
5. **Promoter motif conservation:** Predicted target promoter motifs usually contain CAACC in -35 and CGTC in -10, in agreement with original ECF39 (Rhodius et al., 2013). This promoter motif agrees with the one of RpoE in S. coelicolor (Tran et al., 2019). Nevertheless, the three groups regulated by 2CSs do not have clear promoter motifs, probably because not the σ factor, but the 2CS, is regulating the expression of these ECFs.
6. **Summary:** ECF39 is involved in antibiotic production and cell wall stress response. Proteins from this group are associated with 2CS in ECF39s1, ECF39s3, and ECF39s2, and to AS factors in other subgroups, as in the case of the original group ECF39 (Staroń et al., 2009). Promoter motifs are not conserved in 2CS-

regulated subgroups, potentially because the regulation is carried out by the response regulator. In the remaining subgroups they are similar to the ones of original ECF39 (Rhodius et al., 2013).

**ECF40**

1. **General description:** Members of this group belong to Actinobacteria (100%) and have homology to proteins from the original group ECF40 (98.37%). Proteins from ECF40 contain a pair of cysteine residues in the $\sigma_2$ domain that could take part in their regulation.
2. **Anti-σ factor:** Position +1 contains a putative RsdA-like AS factor in subgroups ECF40s1 (0.81 per ECF), ECF40s3 (0.3 per ECF), ECF40s6 (0.5 per ECF) and ECF40s8 (0.25 per ECF). Nevertheless, when looking at the MSA only putative AS factors from ECF40s1 and ECF40s2 are homologous and contain one clear transmembrane helix (50% in consensus with TopCons, ~100% in the MSA). Interestingly, 17.31% of the putative AS factors from ECF40s1 are fused to a YfdX-like domain. YdfX bind antibiotics in a multi-drug resistant strain of *Salmonella thyphi* (Saha, Manna, Das, & Ghosh, 2016).
3. **Genomic context conservation:** The genetic context of proteins from ECF40 contains one or two proteins with an IMP dehydrogenase/GMP reductase domain (except ECF40s4 and ECF40s5) involved in purine metabolism, ~3 protein kinases per ECF in ECF40s2, a DUF5319-containing protein (+2 of ECF40s1 and ECF40s3), a transcription factor WhiB (-1 of ECF40s3 and -2 of ECF40s2), a LytR cell envelope-related transcriptional attenuator (-8 of ECF40s4), a DUF4193-containing protein (-7 of ECF40s4), a DUF3093-containing protein (-6 of ECF40s4), a dUTPase (-5 of ECF40s4), a DUF3710-containing protein (-4 of ECF40s4), an OB-fold nucleic acid binding domain (-2 of ECF40s4), a DUF3159-containing protein (-1 of ECF40s4), TrkA (+1 and +2 of ECF40s4), an amino acid permease (+3 of ECF40s4), a protein from the TCP-1/cpn60 chaperonin family (ECF40s2, ECF40s6 and -1 of ECF40s9), the 10 Kd subunit of a chaperonin (ECF40s6 and -2 of ECF40s9), a response regulator fused to a LuxR transcriptional regulator (-1 of ECF40s2), a LysR substrate binding domain fused to a bacterial regulatory helix-turn-helix protein of the LysR family (-3 of ECF40s2), a GMP synthase (ECF40s3), a glycoprotease (ECF40s9, ECF40s3 and ECF40s1), an inositol monophosphatase (ECF40s4), DUF541 (ECF40s4), an alpha-glycerophosphate oxidase (ECF40s2 and ECF40s9) and a short chain dehydrogenase (ECF40s2).
4. **Studied members:** Two characterized members of this group, SigD from *Corynebacterium glutamicum* and *Corynebacterium tuberculosis* (ECF40s3) are associated with AS factors encoded in position +1. SigD from *Corynebacterium glutamicum* is involved in mycolate synthesis and lysozyme resistance (Toyoda & Inui,

2017). Another member of ECF40, SigD from *Mycobacterium tuberculosis* (ECF40s1), is associated with nutrient starvation, protein folding and fatty acid degradation, and has been associated the AS factor RsdA, subjected to RIP by Rip1 protease and encoded in +1 (Jaiswal, Prabha, Manjeera, & Gopal, 2013; Schneider, Sklar, & Glickman, 2014).

5. **Promoter motif conservation:** The promoter motifs of members of ECF40 are not conserved, even though SigD from *M. tuberculosis* is auto-inducible (Calamita et al., 2004; Raman, Hazra, Dascher, & Husson, 2004). Analysis of the sigmulon of SigD in *C. glutamicum* revealed conserved promoter binding motifs with GTAACA/G in -35 and GAT in -10 (Pátek et al., 2018), slightly different from SigD in *M. tuberculosis*. Moreover, promoters of SigD in *C. glutamicum* are also recognized by SigH (ECF12s2) and contain overlapping or closely located housekeeping or SigB-regulated promoters (Pátek et al., 2018).
6. **Summary:** ECF40 is regulated by RsdA-like AS factors usually located in position +1, but absent from the genetic context in other cases. Aside from the AS factor, members of ECF40 could be regulated by the binding or release of a putative disulfide bridge between the conserved pair of cysteine residues located in the $\sigma_2$ domain. This pair of residues coexists with the AS factor, indicating a possible two-layered regulation of ECF40.

**ECF41**

1. **General description:** This is the second largest ECF group, with 12,157 unique proteins. Proteins from ECF41 have homology against original group ECF41 (99.09%) and they appear primary in Actinobacteria (78.48%), but also in Proteobacteria (14.49%), Firmicutes (Bacillales) (4.88%), Acidobacteria (0.67%), Cyanobacteria (0.49%), Chloroflexi (0.35%), Bacteroidetes (0.21%), Verrucomicrobia (0.11%), Planctomycetes (0.07%), Gemmatimonadetes (0.04%) and Armatimonadetes (0.04%). Proteins from ECF41 contain a C-terminal extension of ~120 aa with a SnoaL-like domain with structural homology to epoxide hydrolases (Goutam, Gupta, & Gopal, 2017; Sultana et al., 2004), likewise the neighboring groups ECF56, ECF295, and ECF294. This C-terminal extension has a regulatory role since the removal of the distal part of the C-terminal extension yields a variant with enhanced activity, whereas the removal of the proximal part of the C-terminus extension drives the ECF inactive (Wecke et al., 2012). This C-terminal extension is a structural domain that tethers $\sigma_2$ and $\sigma_4$ domains in the right conformation for its activity and might bind small molecules that modulate the ECF $\sigma$ factor conformation (Goutam, Gupta, & Gopal, 2017). Therefore,

the C-terminal extension of members of ECF41 seems to function as a modulator of the ECF activity rather than an inhibitor (Goutam et al., 2017; Wecke et al., 2012).

2. **Genomic context conservation:** Proteins from ECF41 do not contain any conserved protein in their genetic context other than a flavin-containing amine oxidoreductase (Pfam: Amino_oxidase) in -1 of subgroups ECF41s14, ECF41s24, ECF41s30 and ECF41s39, and a carboxymuconolactone decarboxylase (Pfam: CMD) ECF41s8 and in -1 of proteins from ECF41s6 and ECF41s16. These two proteins were already identified in members of the original group ECF41 (Staroń et al., 2009). Other proteins conserved in the genomic context are a DoxX-like protein in -1 of ECF41s13, a F420H(2)-dependent quinone reductase in ECF41s12, a DUF1059 in ECF41s20 and several proteins in the genetic neighborhood of ECF41s25 (NIF3, CobD/Cbib protein, a low molecular weight phosphotyrosine protein phosphatase, a bacterial transferase hexapeptide and an alanine dehydrogenase/PNT).
3. **Studied members:** Among the members of ECF41 there is SigJ (ECF41s9) and SigI (ECF41s27) from *M. tuberculosis*. Both proteins are involved in oxidative stress response. SigI is activating the transcription of an ATP synthase, heat shock proteins, and the catalase/peroxidase KatG, and it plays a role with the response to isoniazid since the deletion mutant is resistant to this antibiotic probably due to the decreased transcription of KatG, which activates isoniazid (Jong-Hee Lee et al., 2012) (reviewed in (Souza et al., 2014)). SigJ is involved in resistance against hydrogen peroxide in a KatG-independent manner (Hu, Kendall, Stoker, & Coates, 2004).
4. **Promoter motif conservation:** Predicted target promoter motifs are conserved and usually contain TGTCACA in -35 and CGTC in -10, in agreement with the original ECF41 (Wecke et al., 2012). Indeed, Ecf41 from *Rhodobacter sphaeroides* and *Bacillus licheniformis* control its expression and one of the coding sequence in -1 (Wecke et al., 2012) exclusively.
5. **Summary:** ECF41 is involved in oxidative stress response. It has a SnoaL-like domain fused in C-terminal that exerts a regulatory role. It has been suggested that this domain can bind subtracts and change the conformation of the $\sigma_2$ and $\sigma_4$ domains, allowing promoter recognition and RNAP binding as a consequence (Goutam et al., 2017). It is possible that this is also part of the regulatory mechanism behind ECF56, ECF294, and ECF295.

**ECF42**

1. **General description:** ECF42 is homologous to original ECF42 (99.9%) and one of the largest groups, with 10,294 unique protein sequences. ECF42 is present in a broad range of bacteria, mainly in Actinobacteria (70.66%) and Proteobacteria (22.49%). Members of ECF42 contain a C-terminal extension of ~200 aa with a tetratricopeptide repeat. This C-terminus is essential for the activity of members of ECF42 (Q. Liu, Pinto, & Mascher, 2018).
2. **Genomic context conservation:** As previously described, members of ECF42 encode a protein with a YCII-related domain in -1 (Staroń et al., 2009). Other conserved domains encoded in the genetic context of members of ECF42 include a MurR protein (ECF42s15), an acetylmuramoyl-L-alanine amidase (ECF42s15), a MraW methylase (ECF42s15), a penicillin-binding protein (ECF42s15), about two copies of a mycolic acid cyclopropane synthetase (ECF42s10) and a MaoC dehydratase (ECF42s10). Therefore, ECF42 seems to be involved in peptidoglycan synthesis and cell-wall stress resistance.
3. **Studied members:** A described members of ECF42, ECF-10 from *Pseudomonas putida* (ECF42s7), controls a small sigmulon of about twelve coding sequences that mediates resistance to antibiotics such as beta-lactams, sulfonamides and chloramphenicol, and is involved in biofilm formation (Tettmann, Dötsch, Armant, Fjell, & Overhage, 2014). Both these functions are related to the overexpression of the efflux pump encoded by *ttgGHI*. Instead, the targets of the proteins encoded by *sven_0747* (ECF42s2) and *sven_4377* (ECF42s3) from *Streptomyces venezuelae* are exclusively the YCII genes sitting directly upstream of their coding sequences (Q. Liu et al., 2018).
4. **Promoter motif conservation:** Promoter motifs of ECF42 share a TGTCGAT in -35 and CGTC in -10, in agreement with the original classification (Staroń et al., 2009) and the target sequences of *sven_0747* and *sven_4377* (Q. Liu et al., 2018).
5. **Summary:** ECF42 has been expanded to over 10,000 unique sequences with the same C-terminal extension, genetic context conservation, and target promoter as original ECF42 (Staroń et al., 2009). Given the absence of putative AS factors in the genetic context of members of ECF42, the regulation could be carried out by the transmembrane helix, found to be essential for the activity of members of ECF42 (Q. Liu et al., 2018).

**ECF43**

1. **General description:** Members of this group are only present in Proteobacteria and have homology to proteins from the original group ECF43 (100%). Interestingly, this group has an ECF σ factor domain (Pfam: Σ70_ECF) instead of the $\sigma_2$ and $\sigma_4$ domains that typically appear in other groups.
2. **Special features:** Proteins from ECF43 are associated with protein kinases encoded in position +1, which have been proposed to be part of the regulatory mechanism in original ECF43 (Staroń et al., 2009). These kinases contain one transmembrane helix (87.8%) and extracytoplasmic tetratricopeptide repeats. Interestingly, the first part of the $\sigma_{2.2}$ region, which contains a DAED motif involved in core RNAP binding (Wilson & Lamont, 2006), is substituted by S/T residues in members of ECF43. This is the phosphorylation site of the protein kinase encoded in +1 (Iyer et al., 2019).
3. **Genomic context conservation:** Other than the protein kinase, genetic contexts of members of ECF43s1 contain a conserved formamidopyrimidine-DNA glycosylase in position -1 and a transposase fused to a fatty acid desaturase.
4. **Promoter motif conservation:** The predicted target promoter motifs contain a conserved GTC in -35 and a CAG in -10.
5. **Summary:** ECF43 has a different sequence, which might impair the retrieval of more proteins of this type. Members of this group could be phosphorylated by a conserved protein kinase encoded in +1. The putative phosphorylation site could be located in an S/T rich area that substitutes the negatively-charged motif that binds to the core RNAP in $\sigma_{2.2}$.

**ECF44 (original classification; no longer present)**

Members of original group ECF44 are present in ECF238, together with members of original ECF24. 13.09% of the members of ECF238 have homology to ECF44.

**ECF45 (original classification; no longer present)**

Members of original ECF45 are mostly ungrouped in the new classification. The reason is likely that only 7 non-redundant proteins were retrieved from original ECF45 in the current search.

**ECF46**

1. **General description:** ECF46 is present in Gram-negative bacteria from the phyla Planctomycetes (44.44%), Proteobacteria (37.04%) and Verrucomicrobia (18.56%) and has homology to original ECF46 (100%).

2. **Anti-σ factor:** Members of ECF46 contain putative AS factors with one transmembrane helix (92.59%) in position +1. These AS factors typically contain AS domain with homology to FecR AS factor (Pfam: FecR) and an extracytoplasmic region with homology to concanavalin A-like lectin/glucanases (Pfam: Laminin_G_3), a carbohydrate binding domain.
3. **Genomic context conservation:** Other proteins conserved in the genomic context include the N-terminal domain of an alpha-L-rhamnosidase (ECF46s4) and a sulfatase fused to a DUF4976 (ECF46s4 and ECF46s3).
4. **Promoter motif conservation:** The promoter motifs are not conserved, which might indicate the lack of autoregulation.
5. **Summary:** ECF46 is regulated by one-TM helix-containing FecR-like AS factors that might be sensing carbohydrates in the periplasm of gram-negative organisms.

**ECF47 (original classification; no longer present)**

This group is present in new ECF218 together with original groups ECF49 and ECF50. 29.27% of the proteins from ECF218 have homology to original ECF47.

**ECF48**

1. **General description:** Proteins from ECF48 are homologous to proteins from the original ECF48 (14%) and are present in Actinobacteria (100%). Members of ECF48 contain C-terminal extensions with a zinc-finger and one transmembrane helix, except in ECF48s2, where the C-terminal extension is shorter and soluble. The length of the C-terminal extension ranges from ~60 aa in ECF48s2 to ~400 aa in ECF48s4.
2. **Genomic context conservation:** We found neither a conserved domain in the context of ECF48 nor any putative AS factor.
3. **Promoter motif conservation:** The predicted target promoter motifs are not conserved, and we did not find the target promoter predicted for original ECF48 (Huang et al., 2015) in our analysis.
4. **Summary:** Therefore, it is likely that ECF48, like in the case of original ECF48 (Huang et al., 2015), is regulated by the zinc-finger of its C-terminal extension.

**ECF49 (original classification; no longer present)**

This group is present in new ECF218 together with original groups ECF47 and ECF50. 33.49% of the proteins from ECF218 have homology to original ECF49.

**ECF50 (original classification; no longer present)**

This group is present in new ECF218 together with original groups ECF47 and ECF49. 28.48% of the proteins from ECF218 have homology to original ECF50.

**ECF51**

1. **General description:** The proteins from ECF51 could be classified against original ECF51 (97.9%), except ECF51s15, which contains ~50 amino acid N-terminal extension. Proteins from ECF51 are present in Actinobacteria (100%).
2. **Regulation:** Even though domains in position +1 are not conserved, proteins in position +1 have homology to AS factors from the original ECF classification (Staroń et al., 2009) (except in subgroup ECF51s4) and contain typically one transmembrane helix (70.73%, ~100% in the MSA). These putative AS factors were also pinpointed in the original ECF51 (Huang et al., 2015). Interestingly, members of ECF51s9 encoded a protein with a $\sigma_4$ domain (80%) and with the $\sigma_2$ domain (20%) in -1. These proteins might function as AAS factors since they are transmembrane proteins (100%), their $\sigma_2$ domain is missing in some cases, and they contain a PknH-like extracellular domain (20%) with sensing functions. Other proteins with $\sigma_2$ or $\sigma_4$ domains appear in members of ECF51 with an average of 0.33 per ECF. Although these proteins are usually ECFs from different groups, some are not included in the ECF library and could be AAS factors as in the case of ECF51s9.
3. **Genomic context conservation:** Other conserved proteins encoded in the context of ECF51 are only present in ECF51s9 and ECF51s13 and include a CoA-binding protein, a formyltransferase, an AICARFT/IMPCHase bienzyme fused to an MGS-like domain, with potential functions in purine sythesis, and an ATP-grasp-domain protein.
4. **Promoter motif conservation:** The promoter motifs are conserved across subgroups with GCAACCG in -35 and GCGTGTC in -10, expanding the motifs of original ECF51 (Huang et al., 2015).
5. **Summary:** In conclussion, the AS factors of ECF51, also identified for original ECF51 (Huang et al., 2015), could be regulated, at least in ECF51s9, by partner-switching via AAS that also contain an extracytoplasmic, potentially sensing, domain. The promoter motifs are conserved and agree with original ECF51 (Huang et al., 2015).

**ECF52**

1. **General description:** All the proteins from ECF52 could be classified against original ECF52 and are present in Actinobacteria.
2. **Special features:** Members of ECF52 have one to two transmembrane helices (except ECF52s8) in a C-terminal extension of up to ~400 amino acids that contains a zinc-finger and, in some cases, a carbohydrate-binding domain such as NPCBM/NEW2 domain (ECF52s1) or ricin-type beta-trefoil lectin-like domain (ECF52s4). Possible sensing of carbohydrates by members of original ECF52 has been speculated (Huang et al., 2015), although in our results it seems that this function might only be present in some subgroups. The $\sigma_4$ domain is divergent in some members of ECF52.
3. **Genomic context conservation:** The genetic context conservation is only extended in ECF52s1 and ECF52s6. In ECF52s1, a conserved pyridine nucleotide-disulfide oxidoreductase is encoded in -2, potentially involved in oxidative stress response, a TetR repressor in -1, and a histidine kinase ATP-biding domain with a PAS fold and a SpoIIE domain in -3. In ECF52s6, ECFs are associated to a peptidase family S49, to a response regulator fused to a LytTr DNA-binding domain and a formamidopyrimidine-DNA glycosylase from the base excision repair system. The protease might release the ECF from the inhibition of the C-terminal extension and the whole signal transduction system could be responding to DNA damage.
4. **Studied members:** One member of ECF52, SCO4117 from *Streptomyces coelicolor* (ECF52s1), activates secondary metabolism, aerial mycelium differentiation, and sporulation (López-García, Yagüe, Gonzalez-Quiñonez, Rioseras, & Manteca, 2018).
5. **Promoter motif conservation:** The predicted target promoter motifs are not conserved among subgroups and, especially the -10 domains, have low information content. The promoter motif from the original group ECF52 (Huang et al., 2015) does not appear in any of the new subgroups. SCO4117 is self-induced, but its binding motifs differs from the one of its subgroup ECF52s1 and the original ECF52.
6. **Summary:** ECF52 inherits the same characteristics of original ECF52, except for the target promoter motifs, which could not be predicted. Some members of ECF52 might be involved in carbohydrate metabolism since their proteins contain a C-terminal extension periplasmic carbohydrate-binding domain. In general, the C-terminal extension of members of ECF52 could function as AS factors.

**ECF53**

1. **General description:** ECF53 is a group with a single subgroup. Members of ECF53 have homology to original ECF53 (100%) and are present in Actinobacteria (100%).
2. **Special features:** In agreement with original ECF53 (Huang et al., 2015), members of ECF53 contain a C-terminal extension of ~350 aa with a zinc-finger (30%) and an average of about one transmembrane helix. No putative AS factors were identified in the genetic contexts of members of ECF53.
3. **Promoter motif conservation:** Predicted target promoter motifs contain TGTTTATC in -35 and TCTCC in -10, in agreement with predictions for original ECF53 (Huang et al., 2015).
4. **Genomic context conservation:** Proteins conserved in the genomic context of members of ECF53 are an aldehyde dehydrogenase, proteins from an ABC transporter and an AsnC-type helix-turn-helix domain.
5. **Summary:** ECF53 contains a zinc-binding domain and a transmembrane helix as a C-terminal extension, as in original ECF53 (Huang et al., 2015). It is possible that the C-terminal extension functions as an AS factor.

**ECF54**

1. **General description:** Members of ECF54 are homologous to members of the original group ECF54 (100%) and are present in Actinobacteria (100%).
2. **Genomic context conservation:** Members of original ECF54 are encoded near carboxypeptidase-regulatory-like domains, subtilases and a CHAT domain protein with tetratricopeptide repeats (Huang, Pinto, Fritz, & Mascher, 2015). Indeed, a carboxypeptidase regulatory-like domain is present in an average of 0.61 copies per ECF always in position +1, a subtilase is found in an average of 0.68 copies per ECF in positions +2 or -1, CHAT domains are found in an average of 0.71 copies per ECF usually in positions +3 or-2, and tetratricopeptide repeats are found in an average of 1.98 per ECF usually in position +2 or -2. Given the conservation of these proteins and the lack of putative AS factor, they could take part in the regulation or response of members of ECF54, as suggested for original ECF54 (Huang et al., 2015).
3. **Promoter motif conservation:** Predicted target promoter motifs are only conserved in the largest subgroups (ECF54s2, ECF54s1, ECF54s3), with GTATCAG in -35 and CTCC in -10, in agreement with original ECF54 (Huang et al., 2015).

4. **Summary:** ECF54 preserves the characteristics of the original group ECF54 (Huang et al., 2015). Members of this group are usually encoded near subtilases, carboxypeptidase regulatory-like domains and CHAT domains, which are fused to tetratricopeptide repeats in some cases.

**ECF55 (original classification; no longer present)**

This group is present in new ECF265 together with original ECF112. 5.84% of the proteins from ECF265 have homology to original ECF55.

**ECF56**

1. **General description:** Proteins from ECF56 have homolog to original ECF56 (99.89%) and are present primary in Actinobacteria (96.15%), but also in Proteobacteria (2.86%), Chloroflexi (0.66%), Gemmatimonadetes (0.22%) and Acidobacteria (0.11%). Members of ECF56 contain a C-terminal extension of ~120 aa with a SnoaL-like domain (Pfam: snoaL_2). SnoaL is a small polyketide cyclase that catalyzes closure steps of the synthesis of polyketide antibiotics in *Streptomyces* spp. (Sultana et al., 2004). However, it has also been observed as part of ECF σ factors from original groups ECF56 and ECF41 (Huang et al., 2015; Staroń et al., 2009).
2. **Genomic context conservation:** The genomic context of members of ECF56 is not conserved beyond in the ECF in most of the cases. In ECF56s11 there is the N-terminal domain of a mycothiol maleylpyruvate isomerase (+1 of ECF56s11). The genetic context of ECF56s10 is conserved and includes an acyl-CoA dehydrogenase, a thiolase, an AMP-binding enzyme, a luciferase-like monooxygenase, a protein from the glyoxalase/bleomycin resistance protein/dioxygenase superfamily and a MlaE permease involved in maintaining the asymmetry of the outer membrane (Malinverni & Silhavy, 2009).
3. **Studied members:** One member of ECF56, SigG from *M. tuberculosis* (ECF56s1), is transcriptional induced by DNA damage from its RecA_NDp promoter as part of the RecA independent DNA damage response, which harnesses glyoxalases, type I estradiol dioxygenases and bleomycin resistance proteins to inactivate toxic compounds that damage DNA (Gaudion, Dawson, Davis, & Smollett, 2013). Indeed, one of these genes is part of the genetic context of ECF56s10.
4. **Promoter motif conservation:** The -10 promoter element predicted for original ECF56 (CGTC) (Huang et al., 2015) appears in ECF56s1 and ECF56s3, but the -35 element is subgroup-specific. The only described member of this group, SigG, is not autoregulated (Gaudion et al., 2013; Jong-Hee Lee, Geiman, & Bishai, 2008). This might explain why the predicted target promoter motifs are not conserved and might indicate

that most of the members of ECF56 are not autoregulated. However, the target promoter elements of SigG (CGATGA-GTCNNTA) (Gaudion et al., 2013) appear in some subgroups, suggesting that some members of ECF56 are autoregulated.

5. **Summary:** ECF56 might be regulated by a SnoaL C-terminal extension and, in the only described member of this group, it activates genes for the RecA-independent DNA damage response.

**ECF57**

1. **General description:** Proteins from ECF57 have homology to original ECF57 (95.8%), are present in Planctomycetes (100%) and contain an average number of transmembrane helices of ~2 in a C-terminal extension that contains a homeodomain-like domain (8.33% of ECF57s1) and a WD40-like beta propeller repeat (21.05% of ECF57s2) as in the case of original ECF57 (Jogler et al., 2012).
2. **Anti-σ factor:** Proteins from ECF57 have a completely conserved pair of cysteine residues, one in $\sigma_2$ and the other in $\sigma_4$. Cysteine residues in $\sigma_4$ have been found to form a disulfide bridge in the ECF SigK from *Mycobacterium tuberculosis* (ECF19s2) that dissociates under reducing conditions, promoting the release of the ECF from the cytoplasmic region of its AS factor, RskA (Shukla et al., 2014). This mechanism is also present in members of ECF18, such as RSPH17029_3536 from *Rhodobacter sphaeroides* (Coulianos, 2018).
3. **Genomic context conservation:** We found a conserved carboxypeptidase regulatory-like in ECF57s3 and tetratricopeptide repeats in the genetic context of the members of ECF57. This protein is present in the genetic context of FecI-like σ factors and ECF54 and might be part of their mechanism of regulation.
4. **Promoter motif conservation:** Predicted target promoter motifs are not conserved.
5. **Summary:** In conclusion, ECF57 is a Plactomycetal group that contains a C-terminal extension (Jogler et al., 2012) with two transmembrane helixes. The activation mechanism of members of ECF57 could involve their release from the membrane upon the onset of specific environmental cues.

**ECF58**

1. **General description:** Members of ECF58 are homologous to proteins from the original group ECF58 (100%) and are present in Planctomycetes (100%) from family Planctomycetaceae.
2. **Genomic context conservation:** Members of ECF58s1 contain a conserved soluble protein with a DUF3470 and a 4Fe-4S binding domain. Like original ECF58 (Jogler et al., 2012), no putative AS factor was identified in the genetic context of members of ECF58.

3. **Promoter motif conservation:** Predicted target promoter motifs are not conserved, indicating the lack of autoregulation.
4. **Summary:** ECF58 was expanded respect to original ECF58 (Jogler et al., 2012), although the regulatory layer that controls its activity it is still elusive.

**ECF59**

1. **General description:** ECF59 has homology to original ECF59 (100%) and is present in Planctomycetes (100%) order Planctomycetales. Proteins from this group have a linker ~10 amino acids longer.
2. **Genomic context conservation:** Original ECF59 is associated with protein kinases encoded in position +1 (Jogler et al., 2012). Indeed, we found an average of 0.94 protein kinase domains per ECF in ECF59. These kinases contain six transmembrane helixes in ECF59s1 (ECF59s1: 77.78%) and are fused to a phosphotransferase domain in 33.33% and an ABC-2 family transporter protein in 22.22%. Instead, the protein kinases of ECF59s2 usually contain one transmembrane helix (ECF59s2: 60%) and are fused to tetratricopeptide repeats in 20% of the cases. Interestingly, 40% of the protein encoded in +1 of ECF59s2 are tyrosine kinases, leaving opened if the rest of the protein kinases of ECF59 are also tyrosine kinases.
3. **Promoter motif conservation:** Predicted target promoter motifs include a conserved TC in -35.
4. **Summary:** In conclusion, ECF59 might be regulated by protein kinases, encoded in +1 in most of the cases, as in the case of original ECF59 (Jogler et al., 2012).

**ECF60 (original classification; no longer present)**

Members of original ECF60 are mostly ungrouped in the new classification. The reason is likely that only 8 non-redundant proteins were retrieved from original ECF60 in the current search.

**ECF61**

1. **General description:** ECF61 is a group with a single subgroup with only six proteins from representative organisms. Members of ECF61 have homology to original ECF61 (100%) and are present in Planctomycetes (100%).
2. **Genomic context conservation:** Members of ECF61 contain a conserved protein kinase (average of 0.83 copies per ECF), usually encoded in position -2. This protein contains five or more transmembrane helices (80%).
3. **Promoter motif conservation:** Predicted target promoter motif is not conserved.

4. **Summary:** In summary, members of ECF61 might be regulated by the transmembrane protein kinase encoded in their genetic context, as in the case of original ECF61 (Jogler et al., 2012).

**ECF62**

1. **General description:** Members of ECF62 are present in Planctomycetes (100%) and have homology to original ECF62 (92.59%).
2. **Genomic context conservation:** All the subgroups contain a protein kinase 1 (average of 1.27 copies per genetic context), usually encoded in position +1 and with one transmembrane helix (50.69%, ~100% looking at the MSA). In subgroups ECF62s1 and ECF62s2, the protein kinase is fused to WD repeats (average of 3.18 copies per ECF). WD40 repeats are found mainly in eukaryotes, where they fold into beta propellers that serve as platforms for the reversible binding of protein complexes or the recognition of post-translational modifications (Xu & Min, 2011). In subgroups ECF62s1 (only visible in the MSA), ECF62s2 and ECF62s3, the protein kinase contains a zinc-finger in N-terminal.
3. **Promoter motif conservation:** Predicted target promoter motifs are not conserved.
4. **Summary:** New and original ECF62 share the same characteristics in terms of conservation of a protein kinase in their genetic neighborhood (Jogler et al., 2012). In this study, we can resolve the domain composition of these kinases, which have WD40 repeats in the largest subgroup and a zinc-finger in ECF62s1, ECF62s2, and ECF62s3.

**ECF101 (original classification; no longer present)**

Members of original ECF101 are part of ECF293 together with original groups ECF13 and ECF117. 11.35% of the proteins from ECF293 have homology to original ECF101.

**ECF102**

1. **General description:** ECF122 have homology to proteins of original ECF102 (61.39%). The topology of this clade of the ECF σ factor tree follows the taxonomic origin of its proteins – members of ECF102 are present in Bacteroidetes (66.17%) (ECF102s2, ECF102s3, and ECF102s5) and Gammaproteobacteria (33.83%) (ECF102s1, ECF102s6).
2. **Studied members:** The only characterized member of ECF102 is SigX from *P. aeruginosa* (ECF102s1). SigX contributes to the composition of the cell membrane changing the membrane fluidity, antibiotic resistance (polymixin B and imipenem), iron uptake, virulence, motility and biofilm formation (Chevalier et al., 2018). This ECF is part of a 7-gene operon which includes a mechanosensitive ion channel (CmpX)

encoded in -1, a putative AS factor (CfrX) encoded in -2 and an outer membrane porin (OprF) encoded in +1 (Chevalier et al., 2018). Even though original reports hypothesized that the regulation of SigX is carried out by CfrX, new findings argue a more complex regulatory mechanism involving a mechanosensing pathway in which CmpX, CfrX, and OprF might participate (reviewed in (Chevalier et al., 2018)). The functions of SigX are diverse. SigX contributes to the composition of the cell membrane changing the membrane fluidity, antibiotic resistance (polymixin B and imipenem), iron uptake, virulence, motility and biofilm formation (Chevalier et al., 2018).

3. **Genomic context conservation**: When looking at ECF102, we observed that subgroup ECF102s1 contains homologs of CmpX, CfrX, and OprF in its genomic context; the conservation extends to position -3 with a CorA-like $Mg^{2+}$ transporter and to -4 with an aldolase as in the case of original ECF102 (Staroń et al., 2009). The mechanosensitive ion channel is conserved in subgroups ECF102s2 and ECF102s5, which indicates a similar regulation as members of ECF102s1. Other conserved domains are a kinase/pyrophosphorylase (ECF102s1) and a pyruvate phosphate dikinase (ECF102s1).
4. **Promoter motif conservation:** Predicted target promoter motifs are not conserved, indicating lack of autoregulation. Interestingly, SigX is autoregulated, but the SigX binding motifs were not found upstream of its coding sequence, indicating the presence of an intermediate regulator induced by SigX (Chevalier et al., 2018).
5. **Summary:** Activation of ECF102 might be triggered by mechanic stimuli transmitted to the ECF via a mechanosensor and a soluble molecule that could act as AS factor, as revealed by the genetic context of original ECF120 (Staroń et al., 2009) and by characterization of SigX from *P. aeruginosa* (Chevalier et al., 2018). ECF102 is likely autoregulated via an intermediate transcription factor.

**ECF103**

1. **General description:** Proteins from ECF103 have homology to original ECF103 (83.78%) and are present in Firmicutes (100%).
2. **Anti-σ factor:** Proteins from ECF103 encode AS factors with one transmembrane helix (84%) in position +1.
3. **Genomic context conservation:** Other conserved genes in the genetic context of members of ECF103 include proteins from ABC transporters (+2 and +3 of ECF103s3).

4. **Promoter motif conservation:** Putative target promoter motifs are only conserved in the largest subgroups and contain TGTCACAA in -35 and a less conserved TCT in -10.
5. **Summary:** ECF103 is regulated by AS factors with one transmembrane helix and either a zinc-binding domain or a DUF4179 (ECF103s2). In agreement with original ECF103 (Staroń et al., 2009), members of ECF103 might be involved in the expression of ABC transporters.

**ECF104 (original classification; no longer present)**

Members of original ECF104 are mostly ungrouped in the new classification. The reason is likely because several small (less than 10 proteins) clusters, discarded in later steps, were created from the 13 non-redundant proteins from ECF104 extracted during this work.

**ECF105**

1. **General description:** Members of ECF105 are homologous to proteins from original ECF105 (12.33%) and are present in Firmicutes from order Bacillales (100%).
2. **Anti-σ factors:** Likewise the original group ECF105 (Staroń et al., 2009), proteins from ECF105 are regulated by TM-bound ZAS factors encoded in position encoded in +1. These AS factor contain a single transmembrane helix (85.11%).
3. **Genomic context conservation:** The multiple transmembrane proteins encoded in positions +2 and +3 in original ECF105 (Staroń et al., 2009) were also found in new ECF105 (usually 9 (34.04%) and 3 (57.46%) transmembrane helices, respectively). Other proteins conserved in the genetic context of members of ECF105 are some enzymes in ECF105s1 involved in translation and amino acid metabolism: acetokinase, pyrroline-5-carboxylate reductase, tRNA synthetase class II and 3-hydroxyacyl-CoA dehydrogenase.
4. **Promoter motif conservation:** Predicted target promoter motifs are conserved. Two -35 elements contain TG(A/T)AGGG and the -10 element has CGTCTAT. The predicted target promoter motif expands the knowledge over members of original ECF105 (Staroń et al., 2009), which lacks a conserved target promoter motif.
5. **Summary:** ECF105 expands the number of proteins (from 4 in original ECF105 (Staroń et al., 2009) to 312 in new ECF105), predicts a target promoter motif but keeps the same genetic neighborhood as original group ECF105.

**ECF106**

1. **General description:** Proteins from ECF106 have homology to original ECF106 (33.2%) and are present in Firmicutes (100%).
2. **Anti-σ factors:** In agreement with original ECF106 (Staroń et al., 2009), proteins from new ECF106 encoded AS factors with one transmembrane helix (97.14%) in position +1 (except ECF106s6). These AS factors contain a DUF4367 in most of its subgroups.
3. **Genomic context conservation:** The genetic context conservation only extends to the inner membrane component of an ABC transporter (Pfam: BPD_transp_1) in subgroup ECF106s1.
4. **Promoter motif conservation:** Putative target promoter motifs are not conserved, indicating lack of autoregulation of members of ECF106.
5. **Summary:** ECF106 is regulated by AS factors with one transmembrane helix and a DUF4367. New ECF106 expands original ECF106 while maintaining the same characteristics.

**ECF107**

1. **General description**: ECF107 is composed by a single subgroup with homology to original ECF107 (98.96%) and present in Firmicutes (100%) from genus *Bacillus* in organisms such as *B. cereus*, *B. anthracis* and *B. thuringiensis*.
2. **Genetic context conservation:** Only five proteins from representative/reference genomes are part of ECF107, making difficult the extraction of meaningful conclusions. These proteins contain a DUF4199 in +2, a DUF5065 in +3, a peptidase family M28 in +4, a tRNA synthetases class I (R) with a DALR anticodon binding domain and an arginyl tRNA synthetase N-terminal domain in -1, a DUF1934 in -2, a LD-carboxypeptidase in -3, a DUF1093 in -4 and a phosphotransferase system EIIC in -5.
3. **Anti-σ factors:** In agreement with original ECF017 (Staroń et al., 2009), we observed that position +1 and +2 of the genetic context of members of ECF107 contain a 2-transmembrane helix and 3-transmembrane helix proteins (100%), respectively. Proteins in +1 and +2 are not homologous in the cytoplasmic region, but they could be both AS factors as previously suggested (Staroń et al., 2009).
4. **Promoter motif conservation:** Predicted target promoter motifs are not conserved.
5. **Summary:** ECF107 preserves the characteristics of original ECF107, with putative AS factors encoded in +1 and +2 (Staroń et al., 2009).

**ECF108 (original classification; no longer present)**

Members of original ECF108 are part of ECF235, together with ECF110 and ECF124. 10.79% of the proteins from ECF235 have homology to proteins from original ECF108.

**ECF109 (original classification; no longer present)**

Members of original ECF109 are mostly ungrouped in the new classification. A total of 82 non-redundant proteins extracted during this work have homology to original ECF109. Of those, 18.29% classify against subgroup ECFs47. The rest were split into 23 clusters with sizes smaller than 10 proteins, that did not pass the size filter.

**ECF110 (original classification; no longer present)**

Members of original ECF110 are part of ECF235, together with ECF108 and ECF124. 12.01% of the proteins from ECF235 have homology to proteins from original ECF110.

**ECF111**

1. **General description:** Proteins from ECF111 are homologous to original ECF111 (only ECF111s1 and ECF111s2, 15.25% of the total). ECF111 is rich in proteins from underrepresented organisms since it is present in Cyanobacteria (68.75%) (ECF111s1, ECF111s2 and ECF111s4), Thermotogae (14.06%) (ECF111s3), Proteobacteria (10.94%) (ECF111s5), Acidobacteria (4.69%) (ECF111s6) and Fibrobacteres (1.56%) (ECF111s8).
2. **Special features:** Proteins from ECF111 might be regulated by a putative AS factor encoded in position +1 with one TM helix (64.91%). Position +2 contains an LTXXQ motif in ECF111s2 and heavy-metal resistance proteins in other subgroups. LTXXQ motif appears in periplasmic proteins that participate in response to stress, and it is also present in ECF03 (Danese & Silhavy, 1998).
3. **Genomic context conservation:** The conservation of the genetic context does not go beyond these proteins.
4. **Promoter motif conservation:** Predicted target promoter motifs are conserved and contain TTGAACCT in -35 and GTCTAAA in -10, indicating a possible autoregulatory role of ECFs from ECF111.
5. **Summary:** AS factors located in +1 might regulate members of ECF111. It is likely that ECF111 is involved in cell envelope stress and/or heavy-metal resistance since the protein in position +1 is typically a heavy-metal resistance protein, and the AS factors contain one transmembrane helix.

**ECF112 (original classification; no longer present)**

This group is present in new ECF265 together with original ECF55. 0.51% of the proteins from ECF265 have homology to original ECF112.

**ECF113 (original classification; no longer present)**

This group is now present in ECF281. 0.85% of the proteins from ECF281 have homology to original ECF113.

**ECF114**

1. **General description:** Members of ECF114 are homologous to members of original ECF114 (36.83%) and are present in Bacteroidetes (99.11%).
2. **Genomic context conservation:** The only conserved protein in the genomic context of members of ECF114 is a GDP-mannose 4,6 dehydratase (ECF114s6). One of the smallest subgroups of ECF114, ECF114s11, composed by only three proteins from representative organisms from the gut microbiota (*Bacteroides uniformis* ATCC 8492, *Bacteroides thetaiotaomicron* and *Bacteroides stercoris* ATC 43183), encodes proteins needed for carbohydrate catabolism (phosphoglycerate mutase, fructose-bisphosphate aldolase, 6-phosphofructokinase, methylglyoxal synthase) and anabolism (alpha-glucan phosphorylase), urea cycle (arginase), a $Na^+/H^+$ antiporter, a MarC-like protein of unknown function, a heat-shock protein and two copies of ferritin. Since iron can catalyze the synthesis of ROS in aerobic conditions, ferritin sequesters $Fe^{2+}$ into an insoluble core as $Fe^{3+}$, and releases it in a controlled manner (Orino et al., 2001). The two copies of ferritin in this region of the genome of *B. uniformis* and *B. thetaiotaomicron* are speculated to be required as iron storage to protect against ROS damage or/and as competitive advantage against other gastrointestinal bacteria (Rocha & Smith, 2013). Indeed, the expression of the ferritin in iron-replete media increases 10-fold in anaerobic media over aerobic culture (Rocha & Smith, 2013).
3. **Studied members:** SigH, from the oral cavity dweller *Porphyromonas gingivalis,* belongs to ECF114s4 and plays a role in aerotolerance (Yanamandra, Sarrafee, Anaya-Bergman, Jones, & Lewis, 2012). SigH is induced upon exposure to $O_2$, and its deletion mutant is downregulated in genes involved in oxidative stress protection (ferritin is the gene downregulated in a greater fold) and hemin uptake (Yanamandra et al., 2012).
4. **Promoter motif conservation:** Promoter motifs are not conserved, potentially due to the lack of autoregulation of members of ECF114 as it is the case for SigH from *P. gingivalis* (Yanamandra et al., 2012).
5. **Summary:** It is likely that ECF114 proteins play a role in aerotolerance of anaerobic Bacteroidetes and might participate in carbohydrate metabolism. We did not find any putative AS factor or potential modulator of members of ECF114, as in the case of original ECF114 (Staroń et al., 2009).

**ECF115**

1. **General description:** Members of ECF115 are present in Firmicutes (100%) from order Bacillales and have homology to original group ECF115 (40.3%) (Staroń et al., 2009). Members of ECF115 contain a C-terminal extension with several possible domains. This C-terminal extension is typical of ~70aa long, although in subgroup ECF115s3, where the C-terminal extension contains a SnoaL-like domain (Pfam: SnoaL_2), it extends to up to ~130 aa.
2. **Genomic context conservation:** Members of ECF115 contain a Clp protease in their genetic context (except ECF115s3). Clp proteases are in charge of the last step of the degradation of AS factors regulated by RIP (Barchinger & Ades, 2013). Genetic context conservation is not conserved beyond the Clp protease.
3. **Anti-σ factor**: no putative AS factor is present in the genetic context of members of ECF115.
4. **Promoter motif conservation:** Predicted target promoter motifs contain ATC in -35 and CGT in -10.
5. **Summary:** Members of ECF115 could be regulated by their C-terminal extension and the Clp protease encoded in their genetic neighborhood. The protease could cleave the C-terminal extension, leaving the ECF free to bind the RNAP and its target promoter. Another possibility is that the Clp protease is the target of the regulation of ECF115. Interestingly, the only subgroup with a different C-terminal extension, ECF115s3 with a SnoaL-like domain, lacks the Clp protease, indicating that only ECF115-like C-termini need the presence of the protease.

**ECF116**

1. **General description:** Members of ECF116 have homology to original ECF116 (25.24%) and belong to Firmicutes from the order Bacillales.
2. **Anti-σ factors:** Proteins from ECF116 contain AS factors in +1 as original ECF116 (Staroń et al., 2009). In subgroups ECF116s1 and ECF116s2, AS factors are fused to GerM-like sporulation and spore germination domains. By their function as periplasmic-sensing AS factors, proteins encoded in +1 of members of ECF116 contain one transmembrane helix in most of the cases (71.37%). Interestingly, some putative AS factors have a SpoIIP domain (4.38%), specially from subgroups ECF116s7 (100%), ECF116s14 (40%) and ECF116s12 (20%). SpoIIP is a cell wall hydrolase required for the dissolution of the septal cell wall (Chastanet & Losick, 2007). The N-terminus of these SpoIIP proteins aligns well with the rest of the putative AS factors of ECF116, indicating that they are indeed real AS factors.

3. **Genomic context conservation:** Other than the AS factor, subgroups ECF116s1 and ECF116s2 are associated with two-component systems typically encoded in -1 (sensor histidine kinase with PAS and HAMP domains) and -2 (response regulator fused to a transcriptional regulator) (frequency of response regulators: 73.77% and 91.12%, respectively; frequency of histidine kinase: 72.13% and 88.24%, respectively). Other conserved genes encoded in the context of members of ECF116 include a ferredoxin I (ECF116s1), a transcriptional regulator fused to a response regulator (ECF116s2), an ECF transporter (ECF116s2) and an RNA pseudouridylate synthase (ECF116s2).
4. **Studied members:** The only characterized protein of ECF116 is SigX (ECF116s1), from *B. subtilis*. SigX is involved in resistance against cationic antimicrobial peptides, and its function is overlapping with SigM and SigV (Cao & Helmann, 2004; Helmann, 2016; Mascher, Hachmann, & Helmann, 2007). SigX is regulated by the single transmembrane pass AS factor RsiX, encoded in +1 (Brutsche & Braun, 1997; Yoshimura, Asai, Sadaie, & Yoshikawa, 2004) and with an extracytoplasmic sporulation and spore germination domain, as predicted for members of ECF116s1. SigX is also regulated by glucose-induced acetylation of the DEAD-box of the RNA-helicase CshA, which induces its expression (Ogura & Asai, 2016).
5. **Promoter motif conservation:** Predicted target promoter motifs are conserved and share TGAAAC in -35 and a less conserved CGTCTAAT in -10. Indeed, the promoter motifs of SigX agree in -35 but not in -10 (Cao & Helmann, 2004).
6. **Summary:** In conclusion, ECF116 seems to be regulated by a membrane-bound AS factor encoded in +1, as in the case of SigX from *B. subtilis*. These AS factors are commonly fused to sporulation, spore germination and SpoIIP domains, which might indicate that members of ECF116 are involved in these processes. The conserved proteins encoded in the genetic context of ECF116 indicate a function in oxidative stress response. Nevertheless, the only characterized member of this group, SigX from *B. subtilis*, is involved in the resistance against cationic microbial peptides.

**ECF117 (original classification; no longer present)**

Members of ECF117 are part of ECF293, together with ECF13 and ECF101. 9.46% of the proteins from ECF293 have homology to original ECF117.

**ECF118**

1. **General description:** Members of this new group belong to Actinobacteria (100%) and have homology against proteins from original ECF118 (70.48%).
2. **Anti-σ factor:** Members of this group might be regulated by a zinc-finger-containing putative AS factor encoded in position +1. These putative AS factors usually contain XXX TM helices (96.03%).
3. **Genomic context conservation:** Members of ECF118 contain a conserved ABC transporter in position +2.
4. **Promoter motif conservation:** The predicted target promoter motifs are only evident in the case of ECF118s1 and contain TGTGAC in -35, TCAC in the spacer and GAAAACC in -10. In general, several groups share TGTGAC in -35 and AACC in -10.
5. **Summary:** ECF118 gained new sequences, and it was predicted to be regulated by six-transmembrane region putative AS factors encoded in +1. ABC transporters are encoded in position +2.

**ECF119 (original classification; no longer present)**

This group is now present in ECF255. 16.8% of the proteins from ECF255 have homology to original ECF119.

**ECF120**

1. **General description:** A percentage of 20.53% of the proteins from new ECF120 has homology to original ECF120. Proteins from ECF120 belong to Bacteroidetes of the orders Bacteroidales, Chitonophagales, Cytophagales, and Flavobacteriales.
2. **Anti-σ factor:** Proteins from ECF120 contain an AS factor with one TM helix in position +1 (71.22%). This AS factor usually contains either a putative zinc-finger domain or a DUF3379.
3. **Genetic context conservation:** The most conserved domain in the genetic context is DUF4252, which appears in positions +2 and +3, in some cases together with a putative auto-transporter adhesin head GIN domain (PFAM: DUF2807, with structural similarity to pectinases), in position +2 of ECF120s13, and in positions +2, -1 and -2 of ECF120s12. These proteins are soluble and might be part of the signaling mechanisms of ECF120. Members of Flavobacteriales encode a peptidase in position -1 (ECF120s2, ECF120s24, ECF120s4, ECF120s6). Other proteins conserved are an outer membrane beta-barrel protein (+2 of ECF120s9, ECF120s7), a protein contain a 4Fe-4S dicluster domain (-1 of ECF120s15), a protein with a Cys-rich domain (-2 of ECF120s15), an amino acid kinase (-1 of ECF120s3), the pyridoxal binding domain of a pyridoxal-dependent decarboxylase (-2 of ECF120s3), a shikimate kinase (ECF120s1 and ECF120s3), a DUF45 (ECF120s3), an Holliday junction DNA helicase ruvB (ECF120s11), a pyruvate

ferredoxin/flavodoxin oxidoreductase (ECF120s13), the TPP binding domain of a thiamine pyrophosphate enzyme (ECF120s13), a putative porin (ECF120s13), a pyruvate flavodoxin/ferredoxin oxidoreductase (ECF120s13), an Adenylosuccinate lyase (ECF120s6) and a periplasmic binding protein (ECF120s4 and ECF120s6).

4. **Studied members:** One characterized member of ECF120 is PG0985 from *P. gingivalis* (ECF120s16). This ECF and the protein coded in +1 (inner membrane protein) and +2 (outer membrane protein with a DUF4252) are highly abundant under heme excess (Veith, Luong, Tan, Dashper, & Reynolds, 2018). Under these conditions, the PG0985 induces the expression of heme efflux pumps (Veith et al., 2018).
5. **Promoter motif conservation:** The promoter motifs are conserved and involved TGTAACAAA in -35 and TCGTCAT in -10, which suggest that they autoregulate their expression.
6. **Summary:** Given the conservation of the large amount of sensing and transport systems in the genetic context of ECF120 and the function of the only described member of this group, it is likely that ECF120 is in charge of sensing and transporting of heme and/or other metal-containing compounds, potentially from the host organisms if the bacteria is a pathogen or a commensal of mammals. ECF120 inherits the disposal of the genetic context of original ECF120 in some subgroups (Staroń et al., 2009), but extends it to other possibilities. Given the conservation of the promoter motifs, it is likely that members of ECF120 are self-regulated.

**ECF121**

1. **General description:** Members of ECF121 share homology to the original group ECF121 (73.36%) and are present in Actinobacteria (100%). Members of ECF121 are associated with subgroup-dependent N-terminal extensions, whose length ranges from ~30 aa in ECF121s5 to ~130 aa in ECF121s5.
2. **Anti-σ factor:** Members of ECF121 might be regulated by a putative AS factor encoded in position +1 (except ECF121s7). The N-terminal part of these putative AS factors is conserved. These AS factors contain one transmembrane helix (27.17% of the ECFs according to TopCons, ~100% when looking at the MSA.)
3. **Genomic context conservation:** The conservation of the genetic context of members of ECF121 is extensive and includes an acyltransferase (ECF121s3, ECF121s5 and ECF121s2, +2 of ECF121s1 and ECF121s4), a NAD dependent epimerase/dehydratase (+3 of ECF121s1 and ECF121s4), a protein with an helix-turn-helix domain (ECF121s2, ECF121s3, ECF121s5 and +4 of ECF121s1 and ECF121s4), a haloacid dehalogenase-like hydrolase (-1 of ECF121s4 and ECF121s1, +2 of ECF121s2 and ECF121s5), an

antibiotic biosynthesis monooxygenase (-1 of ECF121s5), an AMP-binding enzyme (-1 of ECF121s2, -2 of ECF121s4), a glutaredoxin-like domain (DUF836) (-3 of ECF121s4, -2 of ECF121s3, ECF121s1, ECF121s5 and ECF121s2), a putative DNA-binding protein with a CoA binding domain (ECF121s3, -3 of ECF121s1 and ECF121s5), a glutamyl-tRNAGlu reductase (ECF121s2, ECF121s3, ECF121s4, -4 of ECF121s1 and ECF121s5), a porphobilinogen deaminase (hemC) (-5 of ECF121s1 and ECF121s5), an uroporphyrinogen-III synthase HemD (ECF121s2, ECF121s3, ECF121s4, -6 of ECF121s1 and ECF121s5), a delta-aminolevulinic acid dehydratase (HemB) (ECF121s1, ECF121s2, ECF121s3 and -7 of ECF121s5), the N-terminal domain of a bifunctional DNA primase/polymerase (-8 of ECF121s5), a pyrroline-5-carboxylate reductase (ECF121s4 and ECF121s2), a phosphatase (ECF121s1 and ECF121s5), a histone deacetylase (ECF121s1 and ECF121s5), a NAD dependent epimerase/dehydratase (ECF121s5).

4. **Studied members:** One member of ECF121, BldN from *Streptomyces coelicolor* and *Streptomyces venezuelae* (ECF121s1), has been functionally addressed. In *S. venezuelae*, BldN regulates aerial mycelium formation as it activates the transcription of chaplin and rodlin genes (Maureen J Bibb & Buttner, 2003). Moreover, BldN induces the transcription of its single-pass transmembrane AS factor RsbN, which is encoded in +1 and consists in three helices structurally distinct for other AS domain, the so-called class III ASD (Schumacher et al., 2018). BldN from *S. coelicolor* contains a ~86 amino acid N-terminal extension proteolytically processed into a mature BldN (Maureen J Bibb & Buttner, 2003). The processing of the pro-BldN seems to be typical of members of ECF121 since all of them contain an N-terminal extension.
5. **Promoter motif conservation:** Predicted target promoter motifs of members of ECF121 are not conserved among different subgroups. Indeed, target promoter motifs of *S. venezuelae*'s BldN differs, and it is not autoregulated (Maureen J Bibb, Domonkos, Chandra, & Buttner, 2012).
6. **Summary:** Members of ECF112 are regulated by an AS encoded in +1 and by an N-terminal extension, as in the case of BldN from *S. coelicolor* (Maureen J Bibb & Buttner, 2003). Moreover, members of ECF121 contain an extensively conserved genetic context, which encodes proteins for the synthesis of protoporphyrin-IX.

**ECF122**

1. **General description:** Members of ECF122 are homologous to proteins from the original ECF122 (94.99%) and are present in Actinobacteria (100%).

2. **Genetic context conservation:** There are no putative AS factors present for members of ECF122. Instead, all subgroups of ECF122 contain a conserved 6-O-methylguanine DNA methyltransferase involved in DNA repair encoded in position +1 (an average of 1.26 ribonuclease-like domains and 1.41 DNA binding domains per ECF). Whereas this is the only conserved function in the context of proteins from subgroups ECF122s1, ECF122s2, and ECF122s3, proteins from the outermost subgroup, ECF122s4, are also associated with a glycerophosphoryl diester phosphodiesterase (-1), an histidine kinase-like ATPase domain also known as GHKL domain from Gyrase, Hsp90, Histidine Kinase, MutL since they share a structurally related ATPase domain (Dutta & Inouye, 2000) (-2), one or two copies of the glycosyltransferase family 2 domain involved in transferring sugar from/to a variety of substracts (-4), three repeats of the glycosyltransferase domain from family 87 involved in the synthesis of glycolipids in mycobacteria (Morita et al., 2006) and proteins containing the domain UPF0016 suggested to function as calcium transporters (Demaegd, Colinet, Deschamps, & Morsomme, 2014).
3. **Promoter motif conservation:** The predicted target promoter motifs share the pattern CTCAC in -35 and CGTCTAC in -10 in ECF122s1, ECF122s2, and ECF122s3, whereas the promoter motif of ECF122s4 is not conserved. The information content of these motifs is reduced.
4. **Summary:** All in all, the proteins from ECF122 seem to be involved in DNA damage repair. Moreover, genetic neighborhoods of subgroup ECF122s4 contains conserved proteins related to glycolipid metabolism. A possible reason is that members of ECF122s4 evolved in a different range of bacteria that the rest of the members of ECF122. Supporting this notion, members of ECF122s4 appear only in families Intrasporangiaceae (90%) and Nocardioidaceae (10%), whereas remaining subgroups have a different family composition (at least six families in a subgroup).

**ECF123**

1. **General description:** Group ECF123 contains proteins with homology to original ECF123 (97.48%) and are present in Actinobacteria (100%). Proteins from ECF123s1 and ECF123s3 contain a C-terminal extension of 20-30 amino acids with no homology to any Pfam domain.
2. **Special features:** Position +1 has homology to AS factors from original ECF123. In this position, 73.33% of the encoded proteins contain one transmembrane helix and are most likely the proteins with homology to AS factors.

3. **Genomic context conservation:** Members of ECF123s1 and ECF123s2 encoded one or several transmembrane subunits of the succinate dehydrogenases in their genetic context and a FAD-binding domain, also present in ECF123s6.
4. **Promoter motif conservation:** The predicted target promoter motifs of ECF123 is not conserved.
5. **Summary:** Most of the members of ECF123 have a transmembrane AS factor in +1. The genomic context conservation extends further in ECF123s1, ECF123s2 with functions related to the electron transport chain, pointing out the potential regulatory role of ECF123 on oxidative stress response.

**ECF124 (original classification; no longer present)**

Members of this group are part of ECF235, together with original groups ECF108 and ECF110. 11.8% of the proteins from ECF235 have homology to original ECF124.

**ECF125**

1. **General description:** Proteins from ECF125 have homology against original ECF125 (100%) and are present in Actinobacteria.
2. **Special features:** Members of ECF125 are associated with a soluble glyoxalase/bleomycin resistance protein/dioxygenase encoded in +1. This protein could work as an AS factor due to its homology to AS factors in the original classification and to AS factors of group ECF121.
3. **Genomic context conservation:** Other than position +1, conserved proteins encoded in the genetic context of members of ECF125 include a TetR represor, a protein from the major facilitator superfamily (ECF125s4), an aminopeptidase I zinc metalloprotease M18 (ECF125s2), an enoyl-CoA hydratase/isomerase (ECF125s2), a CobB/CobQ-like glutamine amidotransferase (ECF125s2), a creatinine amidohydrolase (ECF125s1), an aminotransferase class-III (ECF125s1), a GMC oxidoreductase (ECF125s1), a FMN-dependent dehydrogenase (ECF125s1), a radical SAM with an iron-sulfur cluster (ECF125s1), an amidinotransferase (ECF125s1) and a pyridine nucleotide-disulphide oxidoreductase (ECF125s1).
4. **Promoter motif conservation:** The predicted target promoter motifs contain AAAA in -35 and CGTCGGA in -10.
5. **Summary:** In conclusion, ECF125 is associated with soluble glyoxalase/bleomycin resistance dioxygenases encoded in position +1 as in original ECF125 (Huang et al., 2015). These proteins could function as AS factors of ECF125.

**ECF126 (original classification; no longer present)**

Members of this group are part of new ECF19, together with original groups ECF19 and ECF34. 3.66% of the proteins of ECF19 have homology to original ECF126.

**ECF127**

1. **General description:** Proteins of this group have homology to original ECF127 (100%) and are present in Actinobacteria (100%).
2. **Special features:** Members of ECF127 are associated to soluble Rieske [2Fe-2S] domain-containing proteins encoded in +1, which in some cases contain a TAT (tween-arginine translocation) signal sequence (26.32% of ECF127s1 and 10% of ECF127s2). This protein does not have homology against any described AS factors, but shares similarity with AS factors from ECF121 and therefore could function as AS factor.
3. **Genomic context conservation:** Other proteins conserved in the genetic context of members of ECF127 are an acyltransferase (ECF127s1) and a calcineurin-like phosphoesterase (ECF127s1).
4. **Promoter motif conservation:** Predicted target promoter motifs contain a conserved GTAACCC -35 region and a non-conserved -10 element.
5. **Regulation:** Members of ECF127 might be regulated by Riske proteins encoded in +1.

**ECF128**

1. **General description:** Members of this group are only present in Actinobacteria and have homology to proteins from the original group ECF128 (91.59%).
2. **Anti-σ factor**: Members of ECF128 contain a transmembrane protein in +1 of both ECF128s1 and ECF128s2 (ECF128s1: 72.73%, ECF128s2: 75%). These proteins contain a conserved cytoplasmic N-terminal region with no Pfam domain that could serve as AS domain. These putative AS factors bear several extracytoplasmic Pfam domains in ECF128s2, such as a BNR/Asp-box repeat, sortilin and/or YCF48 photosynthesis system II assembly factor, which could serve as sensing domains. Since the BNR/Asp-box is present in neuraminidases, it is possible that putative AS factors of ECF128s2 are sensing degradation products of the cell wall and, therefore, responding to cell wall stress.
3. **Genomic context conservation:** Aside from the transmembrane putative AS factor encoded in +1, likewise members of original ECF128, members of new ECF128 contain a membrane protein in -2 and a sortase in -1 of ECF128s1 (87.5%) (Huang et al., 2015). The transmembrane protein in -2 is present in 75% of the

ECFs of ECF128s1, but this value reduces to 16.67% in ECF128s2. Transmembrane proteins encoded in -2 of ECF128s1 are mainly extracytoplasmic, with a short cytoplasmic region in C-terminal unlikely to contain an AS domain. Other conserved domains of members of ECF128 include an ATP-grasp domain in +2 of ECF128s2.

4. **Studied members:** One protein of this group, SCO3736 from *S. coelicolor* (ECF128s1), is the most upregulated protein in a deletion mutant of *ppm1*, which encodes the donor sugar polyprenol phosphate mannose (PPM), which provides sugars for the extracytoplasmic glycosyltransferases (Howlett et al., 2018). *ppm1* deletion mutant is more sensitive to antibiotics that target the biosynthesis of the cell wall (Howlett et al., 2018). SCO3736 is co-transcribed with the sortase encoded in -1 and the transmembrane protein encoded in -2 (Howlett et al., 2018). It is speculated that the sortase attaches to the transmembrane protein to the cell wall (Howlett et al., 2018).
5. **Promoter motif conservation:** Predicted target promoter motifs are only conserved in ECF128s2, with CCAACC in -35 and GGCGTC in -10.
6. **Summary:** In conclusion, ECF128 could be regulated by CAS encoded in +1, which may respond to extracytoplasmic signals related to the synthesis of the cell wall and/or the process of glycosyl transfer.

**ECF129 (original classification; no longer present)**

Proteins from original ECF129 remain ungrouped, since only 3 non-redundant sequences where retrieved by our pipeline.

**ECF130**

1. **General description:** Members of this group are present in Actinobacteria (100%) mostly in genus *Streptomyces* and have homology to proteins from original ECF130 (73.85%).
2. **Special features:** Original ECF130 contains a putative soluble AS factor in -1 (Huang et al., 2015); nevertheless, we could not find any conserved Pfam domain in this position, proteins in this position do not align well in MSA, and none of the proteins matches any AS factor from the original library (Staroń et al., 2009). Therefore, these proteins were not considered putative AS factors.
3. **Genomic context conservation:** We could find conserved proteins with a helix-turn-helix domain in the genetic context of members of ECF130.
4. **Promoter motif conservation:** Predicted target promoter motifs are not conserved.

5. **Regulation:** The regulation of members of ECF130 remains elusive since the putative AS factors described for members of original ECF130 (Huang et al., 2015) are not present in members of new ECF130.

**ECF131**

1. **General description:** Members of this group are only present in Actinobacteria (100%) and have homology to proteins from the original group ECF131 (60.71%).
2. **Anti-σ factor:** Proteins encoded in -1 contain an N-terminal transmembrane helix (58.33%, at least 75% in the MSA). These proteins contain a cytoplasmic C-terminal region that could function as a new type of ASD.
3. **Genomic context conservation:** We did not find any conserved Pfam domain in the genetic context of members of this group.
4. **Promoter motif conservation:** Putative target promoter motifs contain TGTCA in -35 and CGCAC in -10 in ECF131s2 and ECF131s3, although their information content is low.
5. **Summary:** Members of ECF131 could be regulated by a new type of AS factors with C-terminal AS domain. This protein was not described for original ECF131 (Huang et al., 2015).

**ECF132**

1. **General description:** Proteins from ECF132 are homologous to proteins from original ECF132 (80.22%) and are present in Actionobacteria (100%).
2. **Anti-σ factors:** Members of ECF132 might be regulated by a putative AS factor encoded in position +1. Most of the proteins in this position contain one transmembrane helix (58.14%, ~100% when looking at the MSA) but no Pfam domain. The only member of ECF131s5 contains extracytoplasmic domains related to sulfolipid transport (Pfam: SfLAP) and hemolysis (Pfam: Bacillus_HBL) (source: Pfam).
3. **Genomic context conservation:** Other than the AS factor, genetic neighborhoods of members of ECF131s1 contain a conserved CoA-binding protein in +2 with homology to YccU from *E. coli*, a protein overproduced when the cells grow in oleic acid, and also a conserved adenylosuccinate synthetase involved in purine biosynthesis and a YjbR domain.
4. **Promoter motif conservation:** Promoter motifs are not conserved, which might be due to the few sequences included in each group.
5. **Summary:** In conclusion, ECF132 is associated with single transmembrane helix putative AS factors located in position +1 in ECF132.

**ECF201**

1. **General description:** Group ECF201 is the closest group to the σ3-containing members of the phylogenetic tree. Members of this group are present in Firmicutes (100%) from Family Bacillaceae.
2. **Genomic context conservation:** The genetic context of members of this group is not conserved, and it does not contain any AS factor.
3. **Promoter motif conservation:** Predicted target promoter motifs are not conserved.
4. **Summary:** ECF201 is the new ECF group closest to type 3 σ factors that appear in Firmicutes and mainly in the *Bacillus* genus. The regulation and response of members of this group remain elusive.

**ECF202**

1. **General description:** ECF202 is composed of 35 proteins, all from non-representative/non-reference organisms from *Candidatus* phyla and unclassified bacteria. Sequencing of new organisms will shed light into the nature of this group.

**ECF203**

1. **General description:** Members of ECF203 are present in Actinobacteria (100%).
2. **Genomic context conservation:** Members of this group are associated with an O6-methylguanine methyltransferase (1.44 per ECF) in position +1, a TetR repressor (0.88 per ECF), a helix-turn-helix domain (0.83 per ECF), a protein from the major facilitator superfamily (0.78 per ECF) and the metal binding domain of Ada (Pfam: Ada_Zn_binding) (1.22 per ECF), which repairs O6-methylguanine residues and methyl phosphotriesters in DNA (source: Pfam).
3. **Promoter motif conservation:** The predicted promoter motifs are not conserved, either because of the few members of this group or due to a lack of autoregulation.
4. **Summary:** Members of ECF203 could be involved in DNA modification due to the presence of an O6-methylguanine methyltransferase in their genetic context. The presence of a conserved TetR repressor in the genetic context of members of ECF203 suggests that these proteins are involved in their regulatory mechanism, given that no putative AS factor was identified in the proximity of members of ECF203.

**ECF204**

1. **General description:** Members of this group appear exclusively in Actinobacteria.
2. **Anti-σ factor:** Proteins from ECF204 contain a putative AS factor with one TM region (80.36%). Some of these putative AS factors contain typical AS factor domains such as RskA domain, DUF3040, and

DUF4367. In some cases, the coding sequence of the ECF is fused to the coding sequences of the putative AS factor.

3. **Promoter motif conservation:** Predicted target promoter motifs are conserved only in the largest subgroups, ECF204s1, and ECF204s2, and share a GGAACC in -35 and CGGTGTA in -10. Due to the significant conservation of these motifs, it is likely that members of these subgroups are autoregulated.
4. **Summary:** In summary, members of ECF204 might be regulated by putative AS factors encoded in position +1. The conservation of the genetic context does not extend beyond this point.

**ECF205**

1. **General description:** Proteins from this group do not have any homology to any original group and are present in Actinobacteria (93.75%) and Alphaproteobacteria (6.25%). They contain a C-terminal extension of ~150aa that in some cases contains a SnoaL-like domain (6.7% of ECF205s1 and 33.33% of ECF205s2).
2. **Genomic context conservation:** Other than the ECF coding sequence there is a conserved helix-turn-helix domain in the genetic context of members of ECF205.
3. **Promoter motif conservation:** Promoter motifs are not conserved.
4. **Summary:** ECF205 might be regulated by its C-terminal extension, which is some cases contains a SnoaL-like domain as in the cases of members of ECF41, ECF56, ECF295, and ECF294.

**ECF206**

1. **General description:** Members of ECF206 are homologous to proteins from original ECF26 (63.83%) and are present in Acidobacteria (89.29%) from family Acidobacteriaceae and Proteobacteria (10.71%).
2. **Anti-σ factor:** Members of ECF206 contain putative AS factors with one transmembrane helix (51.85%) in position +1. An MSA revealed that ~100% of the sequences indeed contain a transmembrane region. These AS factors contain a zinc-finger with or without and RskA-like domain.
3. **Genomic context conservation:** Other conserved proteins in the genetic context of members of ECF206 are a calcineurin-like phosphoesterase (-1 of ECF206s2) and a cytochrome c (-2 of ECF206s2).
4. **Promoter motif conservation:** Predicted target promoter motifs are not conserved.
5. **Summary:** Members of ECF206 might be regulated by putative AS factors encoded in position +1 with one transmembrane helix and a putative zinc-finger.

**ECF207**

1. **General description:** Members of ECF207 have homology to original ECF26 (93.49%) and are present in Proteobacteria (97.89%), Verrucomicrobia (1.05%) and Planctomycetes (1.05%).
2. **Genomic context conservation:** Members of ECF207 contain a putative AS factor with one transmembrane helix (46.81%) in position +1, except in members of ECF207s16. An MSA revealed the homology of all these proteins, the presence of the $HX_3CX_2C$ zinc-binding motif and the presence of a transmembrane helix in ~100% of the putative AS factors. Aside from the zinc-finger, these AS factors also contain a RskA-like domain in some cases. Other conserved proteins encoded in the genetic context of members of ECF207 are a calcineurin-like phosphoesterase, usually in position -2 or -1, a cupredoxin (-2 of ECF207s3, -1 of ECF207s4), and several proteins in the genetic context of members of ECF207s4 including a DUF3463 fused to a radical SAM, LytB, an aminotransferase class-III, a GMC oxidoreductase, a peptidase family M20/M25/M40 and a DUF2147.
3. **Promoter motif conservation:** Promoter motifs are conserved and contain GGAATAAA in -35 and a less conserved GTC in -10.
4. **Summary:** Members of ECF207 might be regulated by a putative AS factor with one transmembrane helix and a zinc-finger encoded in position +1 from the ECF coding sequence.

**ECF208**

1. **General description:** ECF208 is present in Spirochaetes (100%) from genera *Leptospira* and *Turneriella*.
2. **Anti-σ factor:** Members of ECF208 might be regulated by putative AS factors encoded in +1. These AS factors contain one transmembrane helix (56%), which is more evident in an MSA. No conserved Pfam domain was found in these putative AS factors.
3. **Genomic context conservation:** Members of ECF208s2 contain an average of 2.25 group 1 glycosyl transferases per ECF. These proteins often contain up to two IDEAL domains. In this subgroup, we could also find a DNA ligase and a DUF218-containing protein in position -2, which appears in the S-adenosylmethionine-dependent enzyme YdcF from *E. coli,* involved in anaerobic respiration (Chao et al., 2008). The C-terminal domain of an acyl-CoA dehydrogenase is present in ECF208s3 and ECF208s2.
4. **Promoter motif conservation:** Predicted target promoter motifs of ECF208 are not conserved among subgroups.

5. **Summary:** Proteins from ECF208 might be regulated by putative AS factor located in position +1. Proteins from this group are likely not autoregulated since the predicted promoter motifs are not conserved.

**ECF209**

1. **General description:** Members of ECF209 is present in Actinobacteria (100%).
2. **Anti-σ factor:** Proteins encoded in position +1 of members of ECF209 contain four transmembrane helices (66.66%). This protein could function as AS factor since they are conserved in all the subgroups from ECF209.
3. **Genomic context conservation:** The genetic context conservation extends only to a protein with a helix-turn-helix domain.
4. **Promoter motif conservation:** Even though this is a small group, we could identify putative promoter binding motifs with the sequence CCGTGAACC in -35, CCGACTGT in -10 and a spacer of variable length.
5. **Summary:** ECF209 could be regulated by a new type of short multi-transmembrane helix AS factor.

**ECF210**

1. **General description:** Members of ECF210 are present in Deltaproteobacteria from the genera *Sorangium* (40%), *Sandaracinus* (20%) and *Chondromyces* (40%). Even though this is not one of the smallest ECF groups, from 139 unique variants found in the search, only five come from representative organisms, which makes a thorough analysis of this group challenge.
2. **Promoter motif conservation:** The predicted target promoter motifs are not conserved.
3. **Summary:** It would be hugely beneficial to increase the set of sequenced genomes from Deltaproteobacteria to be able to analyze ECF210.

**ECF211**

1. **General description:** Members of ECF211 are present in Actinobacteria (100%).
2. **Anti-σ factor:** Members of ECF211 might be regulated by putative AS factors encoded in +1 with six transmembrane helices (75%) and a putative AS domain in N-terminus. Putative AS factors from ECF211s2 contain a zinc-binding motif.
3. **Genomic context conservation:** Members of ECF211 are encoded near proteins from ABC transporters (Pfam: ABC2_membrane_3 with 0.92 copies per ECF, and ABC_tran with 2.15 copies per ECF).

4. **Studied members:** A described member of this group, although not included in the ECF library since its genome annotation is missing, is Sig(PspX) from *Planospora alba*, involved in lantibiotic production (Sherwood & Bibb, 2013).
5. **Promoter motif conservation:** Predicted target promoter motifs are not conserved.
6. **Summary:** In conclusion, members of ECF211 might be regulated by 6-TM helix-containing AS factors with a putative AS domain in N-terminus. ECF211 is associated with ABC transporters.

**ECF212**

1. **General description:** Members of ECF212 are present in Firmicutes (100%) mostly from family Bacillaceae.
2. **Anti-σ factor:** Members of ECF212 might be regulated by putative AS factors with one transmembrane helix (100%) encoded in position +1. These putative AS factors are small proteins (~80 aa) which do not contain any extracytoplasmic domain since the transmembrane helix is located in the C-terminus of the protein.
3. **Genomic context conservation:** Other than this putative AS factor, members of ECF212s1 contain a conserved member of acetyltransferase (GNAT) family.
4. **Promoter motif conservation:** Predicted target promoter motifs have low information content, but they contain a common TGAAC in -35 and TGTATA in -10.
5. **Summary:** In conclusion, members of ECF212 might be regulated by membrane-bound putative AS factors that do not contain any extracytoplasmic domain. It is unclear what type of signal these proteins sense.

**ECF213**

1. **General description:** Their taxonomic composition of ECF213 is diverse – the most divergent branch (ECF213s1) contains proteins from Chloroflexi (8.33%), the clade composed by ECF213s4 and ECF213s7 belongs to Planctomycetes (33.33%) and the clade of ECF213s3, ECF213s5 and ECF213s6 to Firmicutes (class Bacilli) (58.33%).
2. **Anti-σ factor:** Putative AS factors were found in the position +1 in the genetic context of ECFs from subgroups ECF213s1, ECF213s4 and ECF213s7. This putative AS factor contains one transmembrane helix (50%, ~100% when looking at the MSA). These putative AS factors could be encoded somewhere else in the genome for members of other subgroups.
3. **Genomic context conservation:** Genetic context conservation does no extend beyond this protein.

4. **Promoter motif conservation:** The predicted target promoter motifs of the subgroups of ECF213 only agree with each other in subgroups ECF213s3 and ECF213s5, with the motifs CCAACA for -35 and CGTTATCTA for -10. Due to the low number of proteins in ECF213, it is not possible to confidently predict any motif for other subgroups.
5. **Summary:** In summary, members ECF213 (ECF213s1, ECF213s4 and ECF213s7) are regulated by putative AS factors encoded in +1, although ECF213 contains too few members to extract significant conclusions. More sequenced organisms from Chloroflexi or Planctomycetes would shed light into the regulation of ECF213.

**ECF214**

1. **General description:** ECF214 contains proteins with homology to the sequences of original ECF10 (1.03%). ECF214 is present in Firmicutes (24.16%) and Bacteroidetes (75.56%).
2. **Anti-σ factor:** Proteins from this group are associated with RskA-like AS factors in +1. In subgroups from Firmicutes, AS factors are fused to a putative zinc-finger; instead, this domain is not conserved in Bacteroidetes. RskA is the AS factor of SigK from *Mycobacterium tuberculosis* (ECF19s2). RskA is degraded by regulated intramembrane proteolysis (RIP), has one transmembrane helix and contains a non-zinc-binding class I ASD, composed of four alpha helices (Schumacher et al., 2018). Most of the proteins in position +1 of ECF214 contain only one transmembrane helix (94.33%), supporting the idea that proteins in position +1 are RskA-like AS factors that could be regulated RIP, as in the case of RskA.
3. **Genomic context conservation:** The presence of a conserved aspartyl-protease in position -3 of ECF214s10, ECF214s18 and ECF214s6 suggests that this enzyme could be in charge of the first step of the degradation by RIP of the AS factors – 79.31% of the proteins in position -3 in these subgroups contain a signal peptide according to TopCons (Tsirigos, Peters, Shu, Käll, & Elofsson, 2015), a prerequisite for a site-1 protease. The aspartyl protease also contains a PDZ domain (PFAM: PDZ or PDZ_2) in ECF214s18. The PDZ domain of the site-1 protease DegS binds unfolded periplasmic proteins, changing the conformation of DegS and allowing for the proteolysis of the anti-σ factor RseA, thereby activating the RpoE-dependent transcription (Hasselblatt et al., 2007; Sohn, Grant, & Sauer, 2007). This suggests a potential activation of members of ECF214s18 by periplasmic unfolded proteins.

   Other proteins conserved in the genetic context of the members of ECF214 are UDP-N-acetylenolpyruvoylglucosamine reductase (-1 of ECF214s10, ECF214s18 and ECF214s6), an aminotransferase class I and II (-2 of ECF214s10, ECF214s18 and ECF214s6), an aspartyl protease (-3 of ECF214s10 and ECF214s6) fused to a PDZ domain in ECF214s18, a DUF1573-containing protein (-4 of ECF214s10, ECF214s18 and ECF214s6), a tRNA synthetase (-5 of ECF214s10, ECF214s18 and ECF214s6), a extracellular solute-biding protein (ECF214s4), a secreted repeat of unknown function (-1 of ECF214s4), an ECF from ECF245s2 (+3 of ECF214s10), a radical SAM superfamily protein (+4 of ECF214s10), a glyceraldehyde 3-phosphate dehydrogenase (ECF214s18, ECF214s6 and +5 of ECF214s10), an histidine kinase from a 2CS fused to a receiver domain (+6 of ECF214s10), a tetraacyldisaccharide-1-P 4'-kinase (ECF214s18, ECF214s6 and +7 of ECF214s10), NIF3 (+8 of ECF214s10), a C4-type zinc ribbon domain (+9 of ECF214s10), a penicillin-binding protein (ECF214s18), a MreB/MbI protein (ECF214s18), a DUF4331 (ECF214s1), a pyridine nucleotide-disulphide oxidoreductase (-1 of ECF214s11), an enoyl-(Acyl carrier protein) reductase (ECF214s11), a beta-eliminating lyase (ECF214s11), a DUF2721 (ECF214s11), a fasciclin domain containing protein (ECF214s2).

4. **Promoter motif conservation:** The promoter motifs are diverse across subgroups so that a consensus could not be defined. Some groups have TATATACG in -35 and a more variable GGATG in -10 (ECF214s20, ECF214s10, ECF214s18, ECF214s6 and ECF214s2).
5. **Summary:** Members of ECF214 are regulated by a RskA-like AS factors that can contain a zinc-finger. In subgroup ECF214s18, ECF214s10 and ECF214s6 these AS factors seem to be degraded by RIP induced by an extracytoplasmic aspartyl protease encoded in position -3, which contains a PDZ domain (in >75% of the proteins only in ECF214s18), in charge of sensing unfolded peptides in the periplasm in DegS from *E. coli* (Hasselblatt et al., 2007; Sohn et al., 2007).

**ECF215**

1. **General description:** Members of EC215 are exclusively present in bacteria from Candidatus phyla. Therefore, we analyzed ECF215 using all the proteins, including the ones from non-representatives/non-reference organisms. Members of ECF215 are mainly present in Candidatus Gottesmanbacteria (ECF215s1), Candidatus Amesbacteria (ECF215s3), Candidatus Daviesbacteria (ECF215s2), Candidatus Nomurabacteria (ECF215s2) and Candidatus Woesebacteria (ECF215s2).
2. **Anti-σ factor:** Proteins encoded in position +1 could be AS factors in subgroups ECF215s1 and ECF215s3. These putative AS factors contain one transmembrane helix (40.63%).

3. **Genomic context conservation:** Members of ECF215s1 contain an average of 1.62 copies of the N-terminal domain of a GHMP kinase. This protein also appears in ECF215s2, with an average of 0.68 copies per ECF.
4. **Summary:** ECF215 is a group only present in bacteria from Candidatus phyla, which makes its analysis difficult since none of these organisms is designated as representative or reference. Therefore, the analysis of all the variants yields results that are biased. Members of ECF215s1 and ECF215s3 could be regulated by putative AS factors encoded in +1.

**ECF216**

1. **General description:** This small group does not have any homology to any original ECF group. ECF216 is present in Proteobacteria from the order Myxococcales (100%). Some sequences from ECF216s2 contain a short C-terminal extension with one transmembrane helix (average of 0.18 TM helices).
2. **Anti-σ factor:** Proteins in +1 are putative AS. In the case of ECF216s2, the putative AS factor contains a zinc-finger and one transmembrane helix in 87.5% of the cases. Instead, 80% of the putative AS factors of ECF216s2 contains at least one TM helix (40% two TM helices and 40% one TM helix).
3. **Genomic context conservation:** The conserved domains included in ECF216 are a carboxypeptidase regulatory-like in ECF216s1 and two EF-hands (2.2 per genetic context) in ECF216s2.
4. **Promoter motif conservation:** Predicted target promoter motifs are not conserved.
5. **Summary:** Members of ECF216 might be regulated by putative AS factors encoded in +1. ECF216 would benefit from further efforts in the study of Deltaproteobacteria.

**ECF217**

1. **General description:** ECF217 has homology to original group ECF59 (100%) and is present in Planctomycetes (100%) genera *Rhodopirellula* and *Pirellula*.
2. **Genomic context conservation:** We found membrane-bound protein kinases usually encoded in +1, similarly to original ECF59 (Jogler et al., 2012). Other conserved domains are only present in ECF217s1, with a Planctomycetes extracellular protein (Pfam: Planc_extracel), a dockerin type I repeat (Pfam: Dockerin_1), which takes part in the cellulosome together with a cadherin (source: Pfam(Finn, 2004)) and a tetratricopeptide repeat.
3. **Promoter motif conservation:** Predicted target promoter motifs are not conserved.

4. **Summary:** Members of ECF217 are encoded close to proteins with a protein kinase domain, similarly to original ECF59 (Jogler et al., 2012). Due to the lack of functional characterization of any of the members of ECF217, it is unclear whether these kinases are regulating the activity of members of ECF217, as speculated for other ECF groups.

**ECF218**

1. **General description:** Members of ECF218 have homology to proteins from original ECF50 (28.48%), ECF47 (29.27%) and ECF49 (33.49%) and are present in Actinobacteria (100%).
2. **Anti-σ factor:** As members of original ECF47, ECF49 and ECF50 (Huang et al., 2015), members of ECF218 contain a putative AS factor with one transmembrane helix (74.33%) in position +1. This AS factor contains a RskA-like domain, but in most of the cases there is no Pfam domain describing it. In subgroup ECF218s8, the AS factor contains a DUF4349 with two transmembrane helices.
3. **Genomic context conservation:** Genetic context conservation only extends beyond the AS factor coding sequence in ECF218s8, which contains an EPSP synthase and an aconitate hydratase.
4. **Studied members:** There is one described member of this group, Sig(MibX) from *Microbispora corallina*, involved in lantibiotic production (Foulston & Bibb, 2011). This protein is not not included in the ECF library since its genome annotation was missing.
5. **Promoter motif conservation:** Different predicted target promoter motifs are possible depending on the subgroup. For instance, members of ECF218s41, ECF218s28 and ECF218s33 contain TGCAAG in -35 and CGACA in -10; members of ECF218s11 and ECF218s57 contain GAAAT in -35 and ACAT in -10; members of ECF218s16 and ECF218s5 contain GTAACGC in -35 and CCAAGTA in -10.
6. **Summary:** ECF218 merges original groups ECF47, ECF49 and ECF50, all present in Actinobacteria and regulated by putative AS factors with one transmembrane helix encoded in +1.

**ECF219**

1. **General description:** Members of this group are only present in Actinobacteria from genus Streptomyces. Most of the proteins from ECF219s2 and ECF219s3 lack of $\sigma_{2.1}$ region or contain a shorter version of the ECF σ factor.
2. **Anti-σ factor:** Proteins in position +1 of ECF219s2 are putative AS factors with one transmembrane helix (66.67%) and a RskA-like domain in some cases.

3. **Genomic context conservation:** Other conserved proteins in the genetic context of members of ECF219 include an SH3 domain (ECF219s1), present in proteins from signal pathways (Vidal, Gigoux, & Garbay, 2001) and a tetratricopeptide repeat (ECF219s1).
4. **Promoter motif conservation:** Predicted target promoter motifs are not conserved.
5. **Summary:** ECF219 might be regulated by one membrane pass AS factors that are located in position +1 in ECF219s2. Other members of ECF219 might encode this protein somewhere else in their genomes.

**ECF220**

1. **General description:** Members of ECF220 are present in Proteobacteria (100%) from family Polyangiaceae and contain a C-terminal extension of variable length with no Pfam domain except for a tetratricopeptide repeat in one sequence from ECF220s2. C-terminal extensions from EF220 contain ~1 TM helices in ECF220s1.
2. **Genomic context conservation:** Proteins encoded in +1 of ECF220s1 contain an N-terminal transmembrane helix and a C-terminal cytoplasmic domain. Similarly, proteins encoded in -1 of ECF220s1 corresponds to an extracytoplasmic protein docked to the membrane by two transmembrane helices. These proteins could be part of the regulatory mechanisms or the response induced by members of ECF220s1. The genetic contexts of members of ECF220 contain a conserved AAA ATPase and a protein with a tetratricopeptide repeat.
3. **Promoter motif conservation:** Predicted target promoter motifs are not conserved, potentially due to the low number of proteins present in this group.
4. **Summary:** ECF220 is a new group that contains a C-terminal extension, with a transmembrane helix in some cases. This C-terminal extension might regulate the activity of members of ECF220 since no putative AS factor was found in their genetic context.

**ECF221**

1. **General description:** Proteins from ECF221 are present in Actinobacteria (100%), mostly from genus *Streptomyces*.
2. **Anti-σ factor:** Proteins encoded in position +1 are putative AS factors with one transmembrane helix (72.55%). These putative AS factors contain DUF4131 in ECF221s1, DUF2694 in ECF221s2 and RskA-like domain in ECF221s3.

3. **Genomic context conservation:** Other conserved proteins encoded in the genetic context of members of ECF221 are a semialdehyde dehydrogenase (-1 of ECF221s1), a subtilase with a bacteria pre-peptidase domain (+2 of ECF221s1) and a peptidase family M1 with an ERAP1-like domain (ECF221s1 and ECF221s2).
4. **Promoter motif conservation:** Promoter motifs are conserved and contain CGAACGTTTC in -35 and CGTCxTA in -10. Due to the high level of conservation, members of ECF221 may autoregulate their transcription.
5. **Summary:** Members of ECF221 are present mainly in *Streptomyces* spp. They might be regulated by the putative AS factor encoded in position +1. Since this AS factor contains one transmembrane helix and a RskA-like domain in some cases, it might be degraded by RIP harnessing the proteases encoded in the genetic context of members of ECF221.

**ECF222**

1. **General description:** This group is present in Actinobacteria (100%).
2. **Anti-σ factor:** Members of ECF222 might be regulated by putative AS factors encoded in +1. These AS factors contain one transmembrane helix (64.29%, ~100% in MSA) and no conserved Pfam domain.
3. **Promoter motif conservation:** Predicted target promoter motifs are not conserved.
4. **Summary:** ECF222 may be regulated by putative AS factors encoded in +1.

**ECF223**

1. **General description:** Members of ECF223 are present in Firmicutes (71.43%) and Fusobacteria (28.57%) from family Fusobacteriaceae.
2. **Anti-σ factor:** Members of ECF223 contain a putative AS factor in position +1 with one transmembrane helix (69.05%). Some domains found in putative AS factors from ECF223 are the transmembrane region of a lysylphosphatidylglycerol synthase, involved in the defense against cationic agents from the host immune system during bacterial infection, and extracytoplasmic peptidase propeptide and YPEB domain (Pfam: PepSY), hypothesized to have a protease inhibitory function and protect the cell from lysis (Yeats, Rawlings, & Bateman, 2004).
3. **Genomic context conservation:** Other conserved proteins in the genetic context of members of ECF223 include a GcpE (ECF223s2), a VanZ-like protein (ECF223s1) and a tRNA pseudouridine synthase (ECF223s1).

4. **Promoter motif conservation:** Predicted target promoter motifs included AATAAA in -35 and TT in -10.
5. **Summary:** In conclusion, members of ECF223 may be regulated by a putative AS factor with one transmembrane helix encoded in position +1.

**ECF224**

1. **General description:** Members of ECF224 are present only in non-reference/non-representative strains mostly from Candidatus phyla. We analyzed these ECFs in terms of non-representative/non-reference organisms.
2. **Anti-σ factor:** Members of ECF224 might be regulated by putative AS factors encoded in +1 with two transmembrane helices (76.92%). These AS factors do not contain any conserved Pfam domain. Moreover, proteins in +2 contain one transmembrane helix (72.97%) but their homology to AS factors and the conservation of the N-terminus is lower than in the putative AS factors encoded in +1, and therefore we do not consider them putative AS factors.
3. **Genomic context conservation:** Members of ECF224 contain a conserved biotin-lipoyl like domain fused to a HlyD secretion protein, a protein from an ABC transporter and an FtsX-like permease.
4. **Summary:** In conclusion, members of ECF224 are present in Candidatus bacteria and might be regulated by two-transmembrane putative AS factors encoded in +1.

**ECF225**

1. **General description:** Members of ECF225 are present in Proteobacteria (100%) from order Alteromonadales.
2. **Anti-σ factor:** Members of ECF225 contain putative AS factors encoded in +1 with one transmembrane helix (18.18%, ~100% when looking at the MSA).
3. **Genomic context conservation:** Aside from the putative AS factor, the conservation of the genetic context extends to a protein with a LysR repressor (ECF225s2 and ECF225s3), a protein with a PhoU domain (ECF225s1), an ACT domain (ECF225s3), a phosphate transporter (ECF225s2, ECF225s3 and ECF225s4) and the periplasmic transporter of an ABC complex ( ECF225s2 and ECF225s3). The LysR repressor is present in ~80% of the proteins from representative/reference organisms in ECF225.
4. **Promoter motif conservation:** Predicted target promoter motifs contain AAACTTTTT in -35 and CGACT in -10, although these motifs do not appear in all the subgroups.

5. **Summary:** In conclusion, members of ECF225 may be regulated by putative AS factors encoded in +1 and may regulate the uptake of phosphate.

**ECF226**

1. **General description:** Proteins from ECF226 are present in Acidobacteria (55.56%), Planctomycetes (33.33%) and Verrucomicrobia (11.11%).
2. **Anti-σ factor:** Members of ECF226 might be regulated by putative AS factors encoded in position +1 with one transmembrane helix (100%). These AS factors contain a DUF3520 and a von Willebrand factor domain in ECF226s1 and ECF226s3.
3. **Promoter motif conservation:** Predicted target promoter motifs are not conserved.
4. **Genomic context conservation:** Genetic context conservation does not extend beyond this protein.
5. **Summary:** Members of ECF226 might be regulated by putative AS factors encoded in position +1.

**ECF227**

1. **General description:** Members of ECF227 are present in Firmicutes (97.17%) and Bacteroidetes (2.83%).
2. **Anti-σ factor:** Members of ECF227 may be regulated by putative AS factors found in position +1. These AS factors contain six transmembrane helices (79.16%) and no conserved Pfam domain.
3. **Genomic context conservation:** Members of ECF227 encode two proteins from an ABC transporter, usually in positions +2 and +3. Genetic context conservation does not extend beyond this point.
4. **Studied members:** Characterized members of ECF227 include CsfU from *Clostridium difficile* (ECF227s3). CsfU is induced by and confers resistance to bacitracin and lysozyme. Induction is achieved through the proteolytic degradation of its anti-σ factor RsiU (encoded in +1) by the protease PrsW, which activates CsfT (ECF30s11) simultaneously (Ho & Ellermeier, 2011).
5. **Promoter motif conservation:** Predicted target promoter motifs of several subgroups included AAAAC in -35 and TTATAT in -10.
6. **Summary:** Members of ECF227 may be regulated by putative AS factors with six transmembrane helices encoded in +1, as in the case of CsfU from *C. difficile* (ECF227s3) (Ho & Ellermeier, 2011). The only described member of this group, CsfU, is involved in bacitracin and lysozyme resistance (Ho & Ellermeier, 2011). This could be the function of the rest of the members of ECF227. Given the presence of ABC transporter encoded downstream of the ECF, these proteins could be part of this antimicrobial resistance mechanism.

**ECF228**

1. **General description:** Members of ECF228 are present in Bacteroidetes (99.74%).
2. **Anti-σ factor:** Members of ECF228 do not contain any evident AS factor in their genetic context.
3. **Genomic context conservation:** Genetic context conservation of members of ECF228 extends to a ribulose-phosphate 3 epimerase (+1 of ECF228s4), a haloacid dehalogenase-like hydrolase (+1 of ECF228s3), a S-adenosylmethionine synthetase (-2 of ECF228s3), a LysR regulator (-3 of ECF228s3), a catalase (-4 of ECF228s3), a tRNA synthetases class I (ECF228s3), a phage integrase (ECF228s3), an enoyl-CoA hydratase/isomerase (ECF228s3), a cytidine and deoxycytidylate deaminase zinc-binding region (ECF228s3) and a signal peptidase II (ECF228s3).
4. **Studied members:** Characterized members of ECF228 include SigP from *Porphyromonas gingivalis* (ECF228s7). SigP is only present in measurable concentrations when it is stabilized via direct interaction with the response regulator PorX from the two-component system PorXY (Kadowaki et al., 2016). SigP induces a type IX secretion system for the secretion of virulence factors (Dou et al., 2018; Kadowaki et al., 2016) and has overlapping functions with RpoE (PG1660) (ECF23s46) (Dou et al., 2018). Even though response regulators are not conserved in the genetic context of members of ECF228, it is possible that other 2CS play a role in the stabilization of members of ECF228.
5. **Promoter motif conservation:** Predicted target promoter motifs of members of ECF228 are not conserved.
6. **Summary:** The function and regulation of members of ECF228 are not clear. The only described member of ECF228, SigP from *P. gingivalis,* requires the stabilizer role of the response regulator PorX to be active (Kadowaki et al., 2016). AS factors were not found in the genetic context of members of ECF228.

**ECF229**

1. **General description:** Members of ECF229 are present in Spirochaetes (100%) from genus *Leptospira*.
2. **Anti-σ factor:** Members of ECF229 encode a putative AS factor in position +1. This AS factor contains one transmembrane helix (68.42%, ~100% in an MSA) and homology to FecR from *E. coli*.
3. **Genomic context conservation:** Other than the FecR-type AS factor, members of ECF229s1 are associated with an average of 2.56 copies of a proton-conducting membrane transporter (Pfam: Proton_antipo_M) per ECF. Therefore, members of ECF229s1 could be involved in the proton-motive force dependent transport of ions across the inner membrane.
4. **Promoter motif conservation:** Predicted target promoter motifs include ATTC in -35.

5. **Summary:** Members of ECF229 are present in Spirochaetes and contain a FecR-like AS factor in position +1. Members of ECF229 could be involved in the proton-motive force-dependent transport of ions across the inner membrane since members of ECF229s1 contain an average of 2.56 copies of a proton-conducting membrane transporter per ECF.

**ECF230**

1. **General description:** Members of ECF230 are present in Firmicutes (100%).
2. **Anti-σ factor:** Members of ECF230 might be regulated by a putative AS factor with six transmembrane helices encoded in position +1. This protein might be related to the ABC transporter protein encoded downstream since two potential AS factors of ECF230 contain an ABC transporter domain (Pfam: ABC_tran).
3. **Promoter motif conservation:** Predicted target promoter motifs of members of ECF230 are conserved and contain AAACTATTT in -35 and TACGAATATA in -10. This degree of conservation suggests that members of ECF230 are self-induced.
4. **Summary:** ECF230 is associated with ABC transporters and might be regulated by a putative AS factors with six transmembrane helices. These putative AS factors might function as transmembrane components of ABC transporters as well. Future studies will shed light on whether these proteins are indeed AS factors.

**ECF231**

1. **General description:** Proteins from ECF231 are present in Bacteroidetes (96.88%) and Acidobacteria (3.13%) of the order Solibacterales.
2. **Anti-σ factor:** Members of ECF231 encode a putative AS factor with one transmembrane helix (96.88%) in position +1. This AS factor contains a cytoplasmic zinc-finger and a periplasmic HEAT repeat.
3. **Genomic context conservation:** Apart from the AS factor, members of ECF231 share genetic context two adhesins (average of 1.84 copies per ECF), usually encoded in position +3. Moreover, members of ECF231s1 contain a conserved MacB-like periplasmic core domain fused to an FtsX-like permease (average of 3.2 copies per ECF)
4. **Promoter motif conservation:** The putative target promoters are conserved and contain TGTAACCTT in -35 and a less conserved -10 with GTTTCCAC in ECF231s1.
5. **Summary:** ECF231 is associated with a single transmembrane helix zinc-binding AS factor with periplasmic HEAT repeats. Members of ECF231 are associated with adhesins.

**ECF232**

1. **General description:** Members of ECF232 are present in Firmicutes (100%).
2. **Anti-σ factor:** Members of ECF232 might be regulated by a putative AS factor with one transmembrane helix (85.11%) encoded in +1 (except ECF232s3). These AS factors contain a conserved N-terminus and extracellular DUF4367 (ECF232s1 and ECF232s2) or WD40 repeats (ECF232s4 and ECF232s5).
3. **Genomic context conservation:** Other conserved proteins in the genetic context of members of ECF232 include an ABC transporter protein, located in position +3 in ECF232s3 and in a range of other positions in the rest of the subgroups. Moreover, ECF232s3 a peptidase M1 (ECF232s3).
4. **Promoter motif conservation:** Predicted target promoter motifs of members of ECF232 contain TGTT in -35 and CGT in -10. These motifs are extended in ECF232s1 and ECF232s2, with TGTTAAT in -35 and CGTATTAAC in -35.
5. **Summary:** ECF232 is associated with membrane-bound AS factors encoded in +1. These AS factors share AS domain but contain extracellular DUF4367 or WD40 repeats. Members of ECF232 are associated with ABC transporters.

**ECF233**

1. **General description:** Members of ECF233 are exclusively encoded in Candidatus bacteria, mostly in *Candidatus Magasanikbacteria.* Therefore, the analysis will be done in terms of non-representative/non-reference organisms.
2. **Anti-σ factor:** Members of ECF233 might be regulated by putative AS factors of with one transmembrane helix (54.54%) encoded in position +1.
3. **Genomic context conservation:** Other proteins encoded in the genetic context of members of ECF233 included a glycosyltransferase of group 1 (CAZY: GT1) fused to a glycosyltransferase of family 4 (CAZY: GT4) (~1 copy per ECF in ECF233s2 and ~3 copies per ECF in ECF233s1) an O-Antigen ligase, a helix-turn-helix domain and tetratricopeptide repeats (ECF233s1).
4. **Summary:** ECF233 appears only in bacteria from *Candidatus* phyla and seems to be related to carbohydrate metabolism since glycosyltransferases are abundant in their genetic context.

**ECF234**

1. **General description:** Members of ECF234 are present in Firmicutes (100%) from genus *Paenibacillus*. The number of proteins in ECF234 from representative/reference organisms is only 4, which makes it difficult to extract meaningful conclusions on the regulation of members of ECF234.
2. **Genomic context and regulation:** No putative AS factor was found in the genetic neighborhood of members of ECF234. Nevertheless, we observed the presence of a conserved 2CS in the genetic context of members of ECF234 (average of 1 per ECF), including the response regulator fused to a transcriptional regulator and a transmembrane histidine kinase domain. Moreover, there are ~2 proteins from an ABC transporter encoded in the genetic context of members of ECF234. The soluble extracellular component of this ABC transporter is only present in members of ECF234s2. Other conserved proteins include a MacB-like periplasmic core domain fused to an FtsX-like permease.
3. **Promoter motif conservation:** Predicted target motifs include GATA in -35 and ACAA in -10 in ECF234s2.
4. **Summary:** ECF234 might be regulated by the two-component system. An ABC transporter and a MacB-like periplasmic core domain fused to an FtsX-like permease are sharing the same genetic neighborhood with members of ECF234.

**ECF235**

1. **General description:** ECF235 contains proteins with homology to several original groups – ECF124 (11.8%), ECF108 (10.79%), ECF110 (12.01%) and ECF40 (0.11%). All the proteins belong to Firmicutes (99.74%), mainly from genera Ruminococcus, Streptococcus and Bacillus, and 33.33% of ECF235s50 from Actinobacteria (0.26% of the total) from genus Slackia.
2. **Anti-σ factor:** Likewise original ECF124, ECF108 and ECF110 (Staroń et al., 2009), proteins from ECF235 encode AS factors in position +1. These AS factors contain the domains of 'σ factor regulator' (PFAM σ_reg_N and σ_reg_C) or a putative zinc-finger domain. AS factors of ECF235 typically contain one transmembrane helix (74.87%), although putative AS factors from ECF235s18 contain four (6.08%). In subgroup ECF235s15 we could not find any putative AS factor. Interestingly, members of ECF235s15 encode other σ factors in their genetic context (150% frequency of another protein with $\sigma_2$ and $\sigma_4$ domains), usually unclassified (64.29%), member of ECF235s15 (14.29%) or ECF235s42 (7.14%), or other non-ECF σ factors (14.29%).

3. **Genomic context conservation:** Apart from a DUF3139 in +2 of ECF235s18, the conservation in the genetic neighborhood of members of ECF235 does not extend beyond the ECF-AS pair.
4. **Studied members:** The only characterized proteins that belong to this group is YlaC (ECF235s6) from *Bacillus subtilis*. YlaC has overlapping functions with SigV, SigZ, and SigY in exopolysaccharide production as part of the biofilm formation, and in the resistance to cefuroxime, ciprofloxacin, and ofloxacin. YlaC is inhibited by the AS factor YlaD, which coordinates $Zn^{2+}$ and releases it upon oxidation or excess of $Mn^{2+}$, forming a disulfide bond and releasing the active YlaC (Kwak, Ryu, Song, Lee, & Kang, 2018). YlaC stimulates sporulation under oxidative stress and excess of $Mn^{2+}$ since it upregulates SigH and ClpP (Kwak et al., 2018).
5. **Promoter motif conservation:** The predicted target promoter motifs of ECF235 contain a conserved TGTAAC in -35 and a subgroup-specific -10 element. YlaC is encoded in a four-gene operon regulated by YlaC (Kwak et al., 2018). The promoter that YlaC regulates (TTGAAC (-35) GCTGTCTA (-10) (Kwak et al., 2018)) is different from our predictions, raising the question of whether non-optimal promoter motifs regulate YlaC.
6. **Summary:** ECF235 is regulated by membrane-bound AS factors that respond to oxidative stress and/or metal stress. We fused several original minor groups (ECF108, ECF110, and ECF124) into a lager new group. ECF235 does not contain SigM from *Bacillus subtilis* as the original ECF124, which is now unclassified. The AS factor of some members of ECF235 is associated to a pair of cysteines that is sensitive to the redox state and the Mn concentration of the cell as described for YlaC of *B. subtilis* (Kwak et al., 2018).

**ECF237**

1. **General description:** ECF237 is present in Proteobacteria (42.86%), Cyanobacteria (42.86%) Actinobacteria (11.43%), and Chloroflexi (2.86%). Members of ECF237s1 and ECF237s5 contain C-terminal extensions of variable size (~40 to ~100 amino acids), but similar enough to align well in a region of ~22aa. Instead, members of ECF237s4 contain a conserved N-terminal extension of ~220 amino acids.
2. **Anti-σ factor:** No AS factors where identified in the genetic context of members of ECF237.
3. **Genomic context conservation**: ECF237 from Proteobacteria contain from 2.4 to 6.75 “Killing trait” proteins (Pfam: RebB). These proteins, produced by *Paramecium* endosymbionts, are part of the R-bodies,

ribbons able to kill sensitive *Paramecium* strains (Schrallhammer et al., 2012). Other conserved proteins in the genetic context of ECF237 include a glutamine amidotransferase (-3 of ECF237s4).

4. **Promoter motif conservation:** Promoter motifs are not conserved, indicating lack of an autoregulatory role of members of ECF237.
5. **Summary:** Members of ECF237 do not encode any putative AS factor in their genetic context. Members of subgroups from Proteobacteria encoded RebB-like proteins, part of the R-bodies of *Paramecium* endosymbionts. R-bodies are proteinaceous ribbon structures able to kill sensitive *Paramecium* strains (Schrallhammer et al., 2012).

**ECF238**

1. **General description:** Members of ECF238 have homology against original ECF24 (84.87%) and ECF44 (13.09%), the latter only present in an internal clade of ECF238. The taxonomic composition is diverse, with proteins present mainly in Proteobacteria (56.37%) and Bacteroidetes (27.94%), but also in Firmicutes (6.37%), Actinobacteria (4.90%), Acidobacteria (2.94%), Nitrospirae (0.49%), Spirochaetes (0.49%) and Chloroflexi (0.49%).
2. **Special features:** We found that members of ECF238 contain conserved cysteine residues in the linker and a Cys-rich domain (CRD) in C-terminus. The CxC motif of the linker typically appears in the internal clade composed by the members of original ECF44 (ECF238s13, ECF238s18, ECF238s10, ECF238s26, ECF238s15, and ECF238s29).
3. **Studied members:** One of the described members of ECF238, CorE2 from *Myxococcus xanthus* (ECF238s15), is known to be activated by Cd and Zn via its CRD in C-terminus (Marcos-Torres, Pérez, Gómez-Santos, Moraleda-Muñoz, & Muñoz-Dorado, 2016; Torres, Muñoz-Dorado, & Moraleda-Muñoz, 2018). The binding rearranges the conformation of CorE2 and turns it active. CorE2 also contains a CxC motif in the linker between $\sigma_2$ and $\sigma_4$ essential for CorE2 functionality (Marcos-Torres et al., 2016; Torres et al., 2018). Another characterized member is SigZ from *Bacillus subtilis* (ECF238s9). SigZ is not regulated by any AS factor and the studies about its function are very limited since its deletion does not cause a significant phenotype and it is expressed at a low level under a range of growth conditions (Y. Luo, Asai, Sadaie, & Helmann, 2010). SigZ is involved together with sigV, sigY and YlaC in resistance to the antibiotics cephalosporin, ciprofloxacin and ofloxacin (Y. Luo et al., 2010).

4. **Genomic context conservation:** Conserved proteins in the context of ECF238 are limited to subgroup ECF238s2, with a permease and a thioredoxin-domain-containing protein.
5. **Anti-σ factor:** Members of ECF238 are not associated with AS factors.
6. **Promoter motif conservation:** Predicted target promoter motifs are not conserved, indicating that ECF238 is not autoregulated. This seems to be the case of SigZ (Y. Luo et al., 2010).
7. **Summary:** All in all, ECF238 is a new group of proteins that are putatively regulated by the binding of metals to Cys-rich regions present in C-terminus and in the linker, as experimentally supported in the case of CarE2 (ECF238s15) and CarE (ungrouped in our classification) (Marcos-Torres et al., 2016; Torres et al., 2018). This novel mechanism could be part of *B. subtilis* SigZ's response, suggesting a solution for the long-standing enigma of its activation signal. All in all, the presence of metal-binding Cys-rich regions in members of ECF238 indicates that this group is involved in metal homeostasis. This response differs from the traditional metal uptake FecI-like groups in that members ECF238 ease metal-associated toxicity effects rather than inducing metal uptake. Interestingly, members of ECF238 seems to not self-regulate their expression, a typical feature of FecI-like σ factors. This similarity between FecI-like members and ECF238 is reflected in the phylogenetic tree – ECF238 and ECF237 are the closest groups to the clade of FecI-like σ factors. The specific metal that induces the response in ECF238 might be different since the specificity changes from Cu in CorE to Cd in CorE2 due to the lack of only one cysteine in their CRD (Torres, Muñoz-Dorado, & Moraleda-Muñoz, 2018).

**ECF239**

1. **General description:** This is a FecI-like group whose proteins have homology to the original group ECF10 (95.18%), although this is not the largest ECF10-like group. The proteins in ECF239 are mainly present in Bacteroidetes (99.22%) mostly from family Flavobacteriaceae.
2. **Regulation and function:** The genetic context of members of ECF239 follow the rule of FecI-like ECFs, with a FecR-like single TM helix AS factor in +1 and an outer-membrane TonB-dependent receptor in +2 (Staroń et al., 2009). The FecR-like AS factor is fused to a DUF4974 domain (except ECF239s9), as in other FecR-like proteins, and contain one TM helix (96.88%). The TonB-dependent receptor might be fused to a carboxypeptidase regulatory-like domain (Pfam: CarbopepD_reg_2) of unknown function, and it is predicted to be a β-barrel outer membrane protein. In the FecI system of *E. coli*, this outer-membrane protein is FecA, a TonB-dependent transporter of ferric citrate across the outer membrane (reviewed in

(Braun & Mahren, 2005)). Members of original ECF10 are suggested to transport carbohydrates across the membrane since their genetic neighborhood contains an outer membrane protein homolog of the polysaccharide-binding proteins SusC and SusD (Staroń et al., 2009). Nevertheless, despite being part of original ECF10, members of ECF239 lack of this carbohydrate binding protein.

3. **Genomic context conservation:** Other conserved proteins are only present in ECF239s3 and include an SRP54-type protein (-2), a tetrahydrofolate dehydrogenase/cyclohydrolase (-3), an amidinotransferase, a glyoxalase/bleomycin resistance protein/dioxygenase superfamily and an arginyl tRNA synthetase.
4. **Promoter motif conservation:** Predicted target promoter motifs often (6/9) contain a conserved (A|G)ACA in -35 and a TAC in -10. It is possible that members of ECF239 are autoregulated, a new feature for a FecI-like group (Staroń et al., 2009).
5. **Summary:** ECF239 contains sequences with homology to original ECF10. However, unlike original ECF10 (Staroń et al., 2009), AS factors from ECF239 do not contain a carbohydrate-binding domain in their genetic context and contain conserved target promoter motifs, indicating autoregulation.

**ECF240**

1. **General description:** ECF240 is the main group with homology to original ECF10 (82.76%). Proteins of this group are present in Bacteroidetes (99.89%). The internal clade formed by ECF240s106, ECF240s155 and ECF240s180 (family Flavobacteriae) contains a conserved ~55aa C-terminal extension. Several subgroups contain proteins with transmembrane helices (up to an average of 0.89 TMH per protein).
2. **Anti-σ factor and regulation:** Likewise original ECF10 (Staroń et al., 2009), genetic contexts of members of ECF240 encode a FecR-like AS factor in +1, a TonB-dependent outer membrane receptor in +2 (Staroń et al., 2009) and an outer-membrane carbohydrate-binding protein in +3 (Pfam SusD_RagB: 84.20%; Pfam SusD-like_2/3: 98.73%; Pfam Reg_prop: 106.79%). The AS factor is fused to a DUF4974, as in the case of other FecI-like AS factors, and it contains one transmembrane helix (87.7%). In some subgroups, this putative operon is inverted in a way that the carbohydrate-binding protein is encoded -3, the TonB-dependent receptor in -2 and the AS factor in -1 (ECF240s1, ECF240s22).
3. **Genomic context conservation:** Other conserved proteins encoded in the neighborhood of members of ECF240 are a glycosyl hydrolase family 92 (-1 and +2 of ECF240s7) and a lysine exporter LysO (ECF240s4).

4. **Promoter motif conservation:** Predicted target promoter motifs are not conserved, in agreement with the lack of autoregulation associated with FecI-like groups (Staroń et al., 2009).
5. **Summary:** ECF240 maintains the characteristics of the original group ECF10 (Staroń et al., 2009). ECF240 is not autoregulated, and its function might be related to carbohydrate scavenging and degradation in Bacteroidetes, one of the most critical components of gut microbiota.

**ECF241**

**6. General description:** ECF241 does not have homology to any original group. Proteins from ECF241 are present mainly in Bacteroidetes (68.28%), Proteobacteria (24.14%), Acidobacteria (6.21%) and Spirochaetes (0.69%).

6. **Anti-σ factor:** Even though there is no BLAST match for AS factors, proteins encoded either in +1 or -1 (ECF241s1, ECF241s5, and ECF241s7) contain two (57.93%) transmembrane helices. Their MSA reveals a short (~10-20 aa) N-terminal region, two consecutive transmembrane helices and a long (~50-260) charged cytoplasmic C-terminal region. This protein typically matches a heavy-metal resistance protein (Pfam: Metal_resist). Nevertheless, its homology to the AS domain of FecR-like AS factors in C-terminus point out to a dual role as resistance protein and ECF regulator.

6. **Genomic context conservation:** Only subgroup ECF241s4 contains a carboxypeptidase regulatory-like domain fused to a TonB-dependent receptor (~80% of the proteins), indicating a similarity with FecI-like ECFs. Nevertheless, this protein is not conserved in the remaining subgroups and AS factors are divergent from canonical FecR-like proteins. Other conserved domains include a LysR transcriptional regulator in ECF241s1 and a homologous of the activator of the Hsp90 ATPase in ECF241s3.

4. **Studied members:** The only described member of this group, BCAL2462 from *Burkholderia cenocepacia* (ECF241s2), upregulates the transcription of the multidrug efflux pump BCAL1510-1512 (Nunvar, Buroni, Cystic, 2018, n.d.).

5. **Promoter motif conservation:** Promoter motifs are conserved in a subgroup-specific manner. The most common motif is CAATA in -35 and TCT in -10.

6. **Summary:** ECF241 composes the outer branch of the iron-uptake FecI-like ECFs, suggesting that its members are ancestors of the iron-acquisition systems where the autoregulation is still preserved, but the TonB-dependent outer membrane transporter of the siderophore and the FecR-type AS factor are missing. The putative AS factors could be of a new type since their C-terminal region seems to be the one that inhibits the AS factor; they contain

heavy-metal resistance domains and two consecutive transmembrane helices in N-terminus. Given the lack of outer membrane transporter, it is possible that members of ECF241 have a role in metal resistance.

**ECF242**

1. **General description:** Members of ECF242 do not have homology to any original group. The topology of the tree follows the taxonomy of its members. ECF242 contain proteins from Proteobacteria (44.19%) (Gammaproteobacteria from genus Alteromonas (ECF242s7), Alphaproteobacteria (ECF242s3 and ECF242s4) and Epsilonproteobacteria (ECF242s5 and ECF242s6)), and Spirochaetes from genus Leptospira (55.81%) (ECF242s1 and ECF242s2).
2. **Anti-σ factor:** All the subgroups contain a FecR-like AS factor in position +1, in some cases fused to a DUF4974 and DUF4880, as in other FecR-like AS factors. These AS factors contain one transmembrane helix (100%).
3. **Genomic context conservation** Subgroups from Proteobacteria encode at least one TonB-dependent receptor (two in ECF242s7), usually in +2, while this receptor does not appear in subgroups from Spirochaetes (ECF242s1 and ECF242s2). Other proteins conserved in the context of ECF242s1 are a thiolase, a rhodanase (in charge of detoxifying cyanide), a metalloenzyme, an ABC transporter and a DoxX protein (+2) suggesting that ECF242 might be in charge of oxidative stress response at least in ECF242s1, which explains the lack of TonB-dependent receptor.
4. **Promoter motif conservation:** Predicted target promoter motifs are not conserved, indicating lack of autoregulation of ECF242.
5. **Summary:** ECF242 is a new FecI-like group from which members of ECF242s1 might be in charge of detoxification of harmful compounds rather than functioning in metal uptake. As other FecI-like ECFs (Staroń et al., 2009), members of ECF242 do not seem autoregulated since the predicted target promoter motifs are not conserved.

**ECF243**

1. **General description:** Proteins from ECF243 combine the non-ECF10 FecI-like groups of the original classification - ECF05 (45.1%), ECF07 (16.62%), ECF06 (6.67%), ECF09 (5.5%), ECF08 (3.14%), ECF10 (0.39%). Proteins from ECF243s172, ECF243s125 and ECF243s57 are fused to their FecR-like AS factor and, therefore, contain ~1 transmembrane helix. Some ECF243 subgroups contain short N-terminal (ECF243s134) or C-terminal (ECF243s62, ECF243s51, ECF243s1, ECF243s77, ECF243s66, ECF243s23,

ECF243s59, ECF243s47, ECF243s134, ECF243s3, ECF243s38, ECF243s53) extension with a subgroup-dependent sequence. The region $\sigma_{2.3}$ is not conserved in ECF243 and there are different possible types of linker between σ2 and σ4, some of them G-rich.

2. **Anti-σ factor:** Members of ECF243 contain a FecR-like AS factor (average of 1.14 per ECF) with one transmembrane helix (82.33%) in position +1 and a TonB-dependent receptor (average of 1.25 per ECF) in +2, as corresponds to the group of FecI from *E. coli* (ECF243s16). FecI is fused to a DUF4880 (average of 1.28 per ECF) and a DUF4974 (1 per ECF).

3. **Genomic context conservation:** Other proteins conserved in the genomic context of members of ECF243 include a heme-binding protein A (HasA) (+3 of ECF243s15), an protein from an ABC transporter (+4 of ECF243s15), a HlyD membrane-fusion protein of T1SS (+5 of ECF243s15), the beta domain of an autotransporter (+3 of ECF243s75), a beta-lactamase (-1 of ECF243s75), a carbohydrate phosphorylase (-2 of ECF243s75), an alpha amylase (-3 and -6 of ECF243s75), a nucleotidyl transferase (-4 of ECF243s75), a starch synthase (-5 of ECF243s75), a Glu-tRNAGln amidotransferase C subunit (ECF243s75), GatB (ECF243s75), an amidase (ECF243s75), a putative modulator of DNA gyrase (ECF243s24), a LysR regulator (ECF243s5 and ECF243s3), a FecCD transporter (ECF243s16), a periplasmic binding protein (ECF243s16), an amino acid permease (ECF243s2), a DUF454 (ECF243s2) and a heme oxygenase (ECF243s2).

4. **Studied members:** Proteins from ECF243 are involved in the uptake of iron-mediated by different siderophores, both synthesized by the same bacteria or by other organisms (siderophore piracy). This regulation can be made via the upregulation of the ferric-siderophore transporters or the synthesis machinery of the siderophore. FecI from *E. coli* (ECF243s16) participates in the uptake of iron-mediated by citrate via the activation of its transport system encoded in the *fecABCDE* operon. When cells are depleted of $Fe^{2+}$, Fur releases its repression over the transcription of the operon *fecIR*. FecR is the membrane-bound AS factor of FecI. FecIR controls the expression of the operon *fecABCDE,* which encodes the proteins necessary for the uptake of ferric citrate. FecA is an outer membrane-bound TonB-dependent receptor that transports citrate loaded with $Fe^{3+}$. Under the presence of ferric-citrate, the repression of FecR over FecI is released and the complex formed by FecI and the N-terminal part of FecR directs RNAP to the transcription of the transport system *fecABCDE* (reviewed in (Braun & Mahren, 2005)). Other ECFs with a similar regulation belong to *Pseudomonas syringae* (Ecf5 (ECF243s5), Ecf6 (ECF243s73), AcsS (ECF243s9),

PSPTO0444 (ECF243s44), PvdS (ECF243s1)) (Llamas, Imperi, Visca, & Lamont, 2014; Yu et al., 2014), *Pseudomonas aeruginosa* (FiuI (ECF243s1), FemI (ECF243s11), PA2050 (ECF243s20), PA2093 (ECF243s20), FoxI (ECF243s10), FecI (ECF243s5), PA4896 (ECF243s54), PA1300 (ECF243s80), HasI (ECF243s15), VreI (ECF243s56), PvdS (ECF243s1), FpvI (ECF243s40)) (Chevalier et al., 2018; Llamas et al., 2014; Marshall, Stintzi, Gilmour, Meyer, & Poole, 2009) and *Pseudomonas putida* (PfrI (ECF243s1), IutY (ECF243s57, TM helix), PP4611 (ECF243s5), PP0352 (ECF243s44), PP3577 (ECF243s9), PP3086 (ECF243s42), PP0162 (ECF243s69)) (Llamas et al., 2014), *Serratia marcescens* (HasI (ECF243s15)) (Biville et al., 2004), *Burkholderia pseudomallei* (BPSL1787 (ECF243s13)) (Alice, López, Lowe, Ledesma, & Crosa, 2006), *Herbaspirillum seropedicae* (PfrI (ECF243s129)) (Trovero et al., 2018), *Bordetella pertussis* (HurI (ECF243s11)) (Vanderpool & Armstrong, 2003), among others. A special case is the FecI-like σ factor PrhI (ECF243s24), from *Ralstonia solanacea*, since it participates in plant infection, as discussed in (Braun & Mahren, 2005).

5. **Paralogy**: ECF43 is the most abundant and diverse ECF group, accounting for the importance of iron uptake, and yet it only appears in Proteobacteria. This system is not necessary in Gram-positive bacteria, since they lack the outer membrane. Even in Gram-negative bacteria such as *E. coli*, FecA is not the only outer membrane protein able to transport iron – other ferric-siderophore complexes are transported through at least six TonB-dependent outer membrane transporters, of which only one, FecA, is regulating an ECF (reviewed in (Andrews, Robinson, & Rodríguez-Quiñones, 2003)). In general, the primary iron starvation regulators are Fur, or DtxR in high GC Gram-positive bacteria (reviewed in (Andrews et al., 2003)). Indeed, Fur is regulating the expression of *fecIR* in *E. coli* (Braun & Mahren, 2005). In general, FecIR-like systems seem the exception rather than the rule in iron scavenging - even within Proteobacteria: only 33.03% 66.97% of the representative/reference, complete, non-metagenomic genomes contain members of ECF243. Nevertheless, when present, there is usually more than one member of ECF243 per organism (average of 3.44 ECF243 per genome and a standard deviation of 3.91). This notion raises the question of why some Proteobacteria has FecI-like systems if they do not seem essential from iron uptake. One possibility is that FecI-like systems provide bacteria with the ability to overexpress iron-import systems on demand. This is also congruent with the fact that only 8.9% of the Proteobacteria harbor several members of the same subgroup of ECF243, suggesting that different subgroups fulfill different functions. Since each copy could be regulating a different uptake system, the bacterium could upregulate only the systems that are currently

in use. However, looking at the characterized members of ECF243, it does not seem that the same subgroup regulates the uptake of the same ferric-siderophore complex. For instance, members of ECF243s11 use as siderophores heme (HurI from *B. pertussis* (Vanderpool & Armstrong, 2003)) and mycobactin (FemI of *P. aeruginosa* (Chevalier et al., 2018)). Similarly, ECF243s9 uses achromobactin in *P. syringae*'s AcsS (Yu et al., 2014) and mycobactin in *P. putida*'s PP3577 (Llamas et al., 2014). More studies into the specificity of FecIR systems would shed light into the source of specificity of these systems given their paralogy in Proteobacterial genomes.

6. **Promoter motif conservation:** Predicted target promoter motifs are conserved in some subgroups, but they do not have a common pattern. These results agree with the data from original FecI-like groups, which are not autoregulated (Braun & Mahren, 2005; Staroń et al., 2009). Interestingly, some promoter regions contain conserved TAAT regions. These could be the binding site of Fur repressor, as discussed in (Andrews et al., 2003).
7. **Summary:** ECF243 is the largest and more diverse ECF group. It is a canonical FecI-like group that merges original groups ECF05-09. Members of this group are mainly, but not only, responsible for iron uptake, and are regulated by FecR-like AS factors and TonB-dependent receptors that sense and transport the loaded siderophore to the periplasm.

**ECF244**

1. **General description:** Some proteins from ECF244 have homology to members of the original group ECF20 (0.52%). ECF244 is present in Firmicutes (100%) from order Bacillales (genera *Brevibacillus*, *Aneurinibacillus*, *Paenibacillus*, *Lysinibacillus*, *Sporosarcina*, *Ureibacillus*, and *Saccharibacillus*). Proteins from this group contain G-rich C-termini (except ECF244s5).
2. **Anti-σ factor:** Genetic neighborhoods from this group encode a putative AS factor with a zinc-binding domain and a DUF4179 in +1, of which 89.13% contain a single TM helix. The genetic context conservation does not extend beyond this point.
3. **Promoter motif conservation:** Promoters are only conserved in some subgroups. The most common motifs are GAGA in -35 and CGTC in -10 (ECF244s1 and ECF244s3). This group might not be autoregulated.
4. **Summary:** ECF244 is a new group regulated by ZAS with a DUF4179 and one transmembrane helix.

**ECF245**

1. **General description:** Members of ECF245 appear in Firmicutes (100%) from order Bacillales and have homology to members of the original group ECF30 (1.78%) and ECF20 (2.53%).
2. **Genomic context conservation:** Members of ECF245 share a similar genetic context composed by a putative anti-σ factor with an N-terminal zinc-binding domain, one transmembrane helix (97.69%) and an extracytoplasmic bactofilin domain (Pfam: Bactofilin) (+1), a protein with the nucleotide-binding domain from a DisA bacterial checkpoint controller, which acts as a di-adenylate cyclase inhibited by stalled replication forks or recombination intermediates (source: Pfam) (+2), a YbbR-like protein (+3), and a phosphoglucomutase/phosphomannomutase (Pfam: PGM_PMM_IV) (+4). Other conserved proteins are a glutamine amidotransferase fused to an SIS domain, which binds to phosphor-carbohydrates (source: Pfam) (ECF245s2).
3. **Studied members:** One described member of this group is SigW from *Bacillus subtilis* (ECF245s1). SigW has overlapping functions to SigM and SigX (Mascher et al., 2007) and is involved in resistance to several antibiotics that inhibit cell wall synthesis during stationary phase (Cao, Bernat, Wang, Armstrong, & Helmann, 2001; Cao, Wang, Ye, & Helmann, 2002b). It is activated when cell membrane proteins are delocalized, and it regulates the fluidity of the cell membrane by changing the ratio between branched and linear fatty acids (Omardien et al., 2018). Genes controlled by SigW are involved both in detoxification of antibiotics and in the synthesis and transport of bacteriocins (Butcher & Helmann, 2006; Cao et al., 2002a).
4. **Promoter motif conservation:** Predicted target promoter motifs are conserved and contain TGAAAC in -35 and CGTAT in -10, suggesting that proteins from this group are autoregulated. Indeed, SigW is autoregulated, and the promoter that it controls is identical to the predicted in this study (Cao et al., 2001).
5. **Summary:** Members of ECF245 could coordinate cell-wall synthesis (bactofilin domain of putative AS factors encoded in +1) with the end of replication or recombination (DisA encoded in +2). The rest of the proteins encoded in the genetic context of members of ECF245, YbbR of unknown function and a soluble phosphoglucomutase/phosphomannomutase, could be involved in cell-wall biosynthesis. In agreement with this, the function of the only described member of ECF245, SigW from *B. subtilis* (ECF245s1), indicates that members of EC245 are part of the cell envelope stress response and regulate the fluidity of the cell membrane.

**ECF246**

1. **General description:** Members of representative/reference organisms within ECF246 are present in Spirochaetes (93.75%) from genus *Leptospira* (ECF246s2) and Firmicutes (6.25%) from genus *Paenibacillus* (ECF246s5). Nonetheless, most of the subgroups of ECF246 are only present in no-representative/non-reference *Candidatus* organisms.
2. **Genomic context conservation:** Proteins from ECF246 contain a conserved two-component system in their genetic contexts (93.75% of the proteins from representative/reference organisms). Since most of the proteins of ECF246 are from non-representative/non-reference organisms, we decided to include those in the analysis of the genetic context. We found that only subgroups from representative/reference organisms (ECF246s2 and ECF246s5) contain conserved proteins in their genetic context – a membrane transport protein (-1 of ECF246s2), the N-terminal of a phospholipase D-nuclease (+2 of ECF246s5), a response regulator fused to a transcriptional regulator (ECF246s2), an ArgJ-like protein (ECF246s2), an 3'-5' exonuclease fused to a HRDC domain (ECF246s2), a protein with a NUDIX domain (ECF246s2), a glutamine amidotransferase (ECF246s2), a binding-protein-dependent transport system inner membrane component (ECF246s5), the oligopeptide/dipeptide transporter of an ABC transporter (ECF246s5) and a bacterial extracellular solute-binding protein (ECF246s5).
3. **Anti-σ factor:** Potential AS factors are proteins encoded in +1, when looking at their MSA proteins from ECF246s1, ECF246s8, ECF246s3, ECF246s6 and ECF246s7 are very similar and seem to contain three transmembrane helices (16.67% according to TopCons, but evident from the MSA). Members of ECF246s4 also contain three TMHs, but they are different from the ECFs mentioned above. Members of ECF246s5 contain two transmembrane helices, and that also seems the case of ECF246s2, although TopCons is not able to detect them. Whether these proteins are functioning as AS factors is unclear.
4. **Promoter motif conservation:** Promoter motifs could not be extracted since most of the sequences are from non-representative/non-reference sequences.
5. **Summary:** In conclusion, members of ECF246 might be regulated by a new type of AS factor encoded in +1. The function of these ECFs remains unknown. The function of subgroup ECF246s5 might related to ABC transporters encoded in its genetic context. Given that some of the putative AS factors encoded in +1 contain domains related to antibiotic resistance, the role of members of ECF246 could be antibiotic resistance in *Candidatus*, *Leptospira,* and *Paenibacillus*.

**ECF247**

1. **General description:** Members of ECF247 are present in Chloroflexi (87.5%) and unclassified bacteria (12.5%).
2. **Anti-σ factor and genetic context conservation:** Proteins from ECF247 encode putative AS factors with two transmembrane helices (62.5%) in +1 and bactofilins in +2.
3. **Promoter motif conservation:** Predicted target promoter motifs are not conserved.
4. **Summary:** ECF247 is a Chloroflexi group associated with putative multi-transmembrane AS factors in +1 and bactofilins in +2.

**ECF248**

1. **General description:** Members of ECF248 are present in Bacteroidetes (98.39%) and Gemmatimonadetes (0.81%).
2. **Anti-σ factor:** No clear AS factor or other regulator was found in members of ECF248.
3. **Genomic context conservation:** Proteins from ECF248 contain a glycosyltransferase from family 2 in position -1 (except ECF248s3). This protein contains four transmembrane helixes (76.14%) and might be the link between the extracellular input signal and the ECF. The conservation of the genetic context of members of ECF248 extends beyond the glycosyltransferase. In ECF248s4 there is a queuine tRNA-ribosyltransferase (-2), a YjgP/YjgQ permease (-3), an EamA-like transporter (-4), a GHMP kinase in -5, an rRNA small subunit methyltransferase G (usually in +1). Subgroups ECF248s1 and ECF248s3 share a conserved lipoyl synthase of radical SAM family. Other than that, ECF248s1 contains a conserved tetraacyldisaccharide-1-P 4'-kinase, an UDP-N-acetylenolpyruvoylglucosamine reductase and a glyceraldehyde 3-phosphate dehydrogenase.
4. **Promoter motif conservation:** Predicted target promoter motifs are not conserved in ECF248.
5. **Summary:** Members of ECF248 encode a glycosyltransferase in -1 that could be involved in their regulatory mechanism. Glycosyltransferases of this type (CAZy: GT2) include cellulose synthases and chitin synthases, among others. Members of ECF248 could be involved in cell wall synthesis.

**ECF249**

1. **General description:** Proteins of ECF249 have homology to original groups ECF04 (2.63%), ECF10 (2.19%), ECF20 (0.44%) and ECF52 (0.44%) and they are present mainly in Firmicutes (41.81%),

Bacteroidetes (27.68%), Acidobacteria (9.6%), Actinobacteria (4.52%), Chlorobi (4.52%), Planctomycetes (7.91%), Proteobacteria (0.56%) and Verrucomicrobia (3.39%).

2. **Anti-σ factor:** Members of ECF249 are regulated by AS factors in position +1 (except ECF249s6) and contain one (38.21%) or two (43.21%) transmembrane helices (only in ECF249s1). Putative AS factors of ECF249s14, which has homology to original group ECF52, are fused to the ECF.
3. **Genomic context conservation:** Other conserved proteins include several cytochrome proteins, enzymes for the synthesis of tryptophan and proteins with other catalytic activities conserved in ECF249s1, and an EF-hand domain ($Ca^{2+}$ binding) and a SCO1/SenC protein (required for the optimal cytochrome c oxidase activity) in ECF249s3.
4. **Promoter motif conservation:** Predicted target promoter motifs of the contain CGCAACTTT in -35 and GTT in -10.
5. **Summary:** ECF249 combines several original groups. AS factors of ECF249 contain one TM helix in most of the subgroups. Promoter motifs are conserved, which indicates that members of ECF249 are autoregulated.

**ECF250**

1. **General description:** Members of ECF250 are present in Firmicutes (100%) from families Bacillaceae and Paenibacillaceae. Members of ECF250 contain two conserved cysteine residues in their $\sigma_2$ domain.
2. **Anti-σ factor:** Members of ECF250 are associated with single-membrane pass (82.67%) AS factors encoded in +1. These putative AS factors typically contain a DUF4367 or a DUF4179.
3. **Genomic context conservation:** Other proteins conserved in the genetic context of members of ECF250 are a bacterial extracellular solute-binding protein in ECF250s4 and a binding-protein-dependent transport system inner membrane component (Pfam: BPD_transp_1). Both these domains are part of ABC transporters.
4. **Promoter motif conservation:** Predicted target promoter motifs are conserved among subgroups, with TGTCAC or TGTCCCTTT in -35 and GTTATATA in -10.
5. **Summary:** Members of ECF250 are associated with membrane-bound AS factors encoded in +1 and with ABC transporters. The promoter motifs are conserved and similar, especially the -10 element, throughout the group.

**ECF251**

1. **General description:** Members of ECF251 are present in Bacteroidetes (98%) and Ignavibacteriae (1%) from genus *Melioribacter*.
2. **Anti-σ factor:** Proteins in +1 are conserved and contain two transmembrane helices (38% according to TopCons but found in the MSA). These proteins could function as AS factors. A shared domain found in these proteins is DUF2207.
3. **Studied members:** Characterized members of ECF251 include RpoE from *P. gingivalis* (ECF251s13). RpoE activates the expression of T9SS and other virulence factors under oxidative stress conditions together with PG0162 (ECF228s7) (Dou et al., 2018). This ECF could be regulated by an AS factor located in +1 with which it shares operon (Dou et al., 2018).
4. **Promoter motif conservation:** Predicted target promoter motifs contain GAC in -35 and GGT(T|A)ACA in -10. RpoE is autoregulated, but it binds to promoter motifs with low information content (Dou et al., 2018).
5. **Summary:** All in all, members of ECF251 are regulated by a putative AS factor encoded in +1. The only described member of this group, RpoE from *P. gingivalis*, is in charge of T9SS expression involved in virulence (Dou et al., 2018).

**ECF252**

1. **General description:** Members of ECF252 are present in Bacteroidetes (42.11%) (ECF252s3, ECF252s4, ECF252s6), Proteobacteria (38.60%) (ECF252s1, ECF252s2) and Spirochaetes (19.30%) from genus *Treponema* (ECF252s5).
2. **Anti-σ factor:** Members of ECF252 contain a putative AS factor in position +1 with two transmembrane helices (49.12%). In some cases (10% of ECF252s2), this protein has homology to RseA from *E. coli*. These proteins contain zinc-fingers in ECF252s5. Genetic context conservation does not extend beyond this point.
3. **Promoter motif conservation:** Predicted target promoter motifs contain AAA in -35 and TTG in -10.
4. **Summary:** Members of ECF252 appear in a vast range of bacterium phyla, members of this group could be regulated by the transmembrane protein encoded in +1.

**ECF253**

1. **General description:** This group is composed of proteins from Proteobacteria (100%), mostly from *Paenibacillus*.
2. **Anti-σ factor:** Members of this group encode a potential AS factor in position +1 with one transmembrane helix (66.67%). This AS factor contains a zinc-finger and/or a RskA-like domain.
3. **Genomic context conservation:** Other proteins conserved in the genetic context of members of ECF253 include GatB, MgtC, needed for pathogenesis in *Salmonella* spp. (Jang-Woo Lee & Lee, 2015), an amidase, a glycosyl hydrolase from family 18, a copper amine oxidase, nucleotidyltransferase, a ferric uptake regulator (only ECF253s2), the C subunit of a Glu-tRNAGln amidotransferase (only ECF253s2), a protein with an acetyltransferase (GNAT) domain (only ECF253s2) and a zinc-ribbon containing domain (only ECF253s2). In general, the genetic contexts of members of ECF253 are related to tRNA loading and metal homeostasis.
4. **Promoter motif conservation:** Predicted target promoter motifs are conserved and contain TTTGAAGGG in -35 and CGTCTAA in -10.
5. **Summary:** ECF253 is a small group of proteins from Proteobacteria, mainly *Paenibacillus* with extended conservation of the genetic context associated with tRNA loading, metal homeostasis and carbohydrate metabolism. The promoter motifs of members of ECF253 are conserved, indicating an autoregulatory role.

**ECF254**

1. **General description:** Members of ECF254 are present in Firmicutes (100%) from order Clostridiales.
2. **Genomic context conservation:** Members of ECF254 contain a conserved genetic context with a glycosyltransferase family 28 in +1, a DUF4367 in +2, a HIT domain in -1, a histidine kinase from a 2CS in -2, an intramembrane protease from rhomboid family in -3, a TadE-like protein involved in production of pili for surface adherence, a type IV leader peptidase (Pfam: Peptidase_A24) involved in processing the prepilins of T4SS, a protein F from T2SS (Pfam: T2SSF) which forms the platform for the assembly of T2SS and type IV pili machinery, and a TSCPD domain-containing protein.
3. **Anti-σ factor:** The DUF4367 encoded in +2 has homology to an AS factor and contains one transmembrane helix (77.78%). These AS factors could be inactivated by RIP mediated by the protein of rhomboid family encoded in -3.

4. **Pontential function:** A possible function of members of ECF254 is cell envelope homeostasis since glycosyltransferases from family 28 (CAZy: GT28) include enzymes related to glucolipid synthesis (1,2-diacylglycerol 3-β-glucosyltransferase), peptidoglycan synthesis (undecaprenyldiphospho-muramoylpentapeptide β-N-acetylglucosaminyltransferase). Moreover, the presence of genes for the production and type IV pili indicates that members of ECF254 could activate gliding motility, adherence to host cells and/or biofilm formation (Melville & Craig, 2013) under inducing conditions.
5. **Promoter motif conservation:** Predicted target promoter motifs are not conserved.
6. **Summary:** Members of ECF254, present in Clostridiales mostly from genus *Ruminococcus*, are regulated by a DUF4367-containing AS factor encoded in +2, which could be degraded by the rhomboid intramembrane protease encoded in -3. ECF254 could be involved in cell envelope homeostasis since there is a glycosyltransferase of family 28 encoded in +1. Members of ECF254 are encoded close to type IV pili machinery. These types of systems are needed for gliding motility, adherence to host cells and/or biofilm formation in gram-positive bacteria (Melville & Craig, 2013). It has been found that type IV pili are required in adherence of *Ruminococcus albus* to cellulose fibers and their degradation (Rakotoarivonina et al., 2002). The type IV pili could have a similar function in members of ECF254.

**ECF255**

1. **General description:** Members of ECF255 are present in Bacteroidetes (98.13%). Proteins from ECF255 have homology to original ECF119 (16.8%).
2. **Anti-σ factor:** Putative AS factors of members of ECF255 are encoded in +1. This protein harbors one transmembrane helix (78.5%, also present in ECF255s4 and ECF255s10) and contains the C-terminal domain of TonB (PFAM: TonB_C) and a carboxypeptidase regulatory-like domain in the extracytoplasmic region. This region of TonB binds the so-called Ton-box of outer membrane transporters and provides them with energy from the proton-motif force for the active transport of proteins of more than ~600 Da such as iron-siderophore complexes and other organometallic compounds (Shultis, Purdy, Banchs, & Wiener, 2006). Therefore, AS factors from ECF255 could use C-terminal domain of TonB to sense extracellular organometallic compounds harnessing the contact with a Ton-box-containing transporter.
3. **Special features:** Position -1 usually encodes a soluble (41.12%) or secreted (58.89%) protein with von Willebrand factor domain, DUF3520, and a carboxypeptidase regulatory-like domain.
4. **Genomic context conservation:** Conservation in the genetic context does not extend beyond this point.

5. **Promoter motif conservation:** Promoter motifs are conserved and contain TATGGAT in -35 and GCATCT in -10.
6. **Summary:** Members of ECF255 are regulated by AS factors with one transmembrane helix and an extracytoplasmic region that contains TonB C-terminus and a carboxypeptidase regulatory-like domain.

**ECF256**

1. **General description:** Some proteins from ECF256 have homology to original groups ECF31 (1.32%). Proteins from ECF256 are present in Firmicutes (100%), mostly from genus *Bacillus*.
2. **Anti-σ factor:** The regulation of members of ECF256 is not clear. Only one protein from ECF256s2 contains a putative RskA-like AS factor in +1.
3. **Genetic context conservation:** The genetic context of members of ECF256 is not conserved, and the proteins encoded surrounding the ECF are generally soluble.
4. **Promoter motif conservation:** Only ECF256s1 has clear predicted target promoter motifs, with TGAAAC in -35 and CGTTTCAT in -10.
5. **Summary:** ECF256 could be regulated by AS factors, but these are only present in the genetic context of ECF256s2.

**ECF257**

1. **General description:** Members of ECF257 present in Firmicutes (100%) from order Bacillales.
2. **Genomic context conservation:** Genetic contexts of members of ECF257 encode a peptidase family M20/M25/M40 (ECF257s2 and -2 of ECF257s1), a protein with an AAA ATPase (ECF257s2 and ECF257s1), proteins from ABC transporters (ECF257s1 and ECF257s2), a transporter with a DUF21 (ECF257s1), a protein from the ArsC family, a transcriptional regulator under oxidative stress (ECF257s2) and a MarR protein, involved in antibiotic resistance (ECF257s2).
3. **Regulation:** Proteins from ECF257 do not seem to be regulated by AS factors. Instead, all subgroups of ECF257 are associated with ABC transporters and AAA ATPases, which might play a role in the regulation or the response of ECF257.
4. **Promoter motif conservation:** Promoter motifs are conserved and contain CTTACA in -35, TGTAA in the spacer and TGAAG in -10.

5. **Summary:** The regulation of ECF107 is unclear since no AS factors were found in its genomic context. ABC transporters and proteins involved in multidrug resistance, encoded in the neighborhood of members of ECF107 might be part of the response mechanism or the modulation of these ECFs.

**ECF258**

1. **General description:** Members of ECF258 are present in Firmicutes (100%).
2. **Anti-σ factor:** Members of ECF258 contain a putative AS factor with a DUF4179 and one transmembrane helix (77.5%) in position +1 (except ECF258s4).
3. **Genomic context conservation:** Other conserved proteins in the genetic context of members of ECF258 include proteins from ABC transporters with ABC transporter domain (Pfam: ABC_tran) or binding-protein-dependent transport system inner membrane component domain (Pfam: BPD_transp_1).
4. **Promoter motif conservation:** Predicted target promoter motifs are conserved and contain TGAACC in -35 and AATATA in -10.
5. **Summary:** Members of ECF258 are present in Firmicutes. ECF258 might be regulated by DUF4179-containing AS factors located in +1. These ECFs contain ABC transporters in their genetic context, which might be part of the cell response they regulate.

**ECF259**

1. **General description:** Members of ECF259 are present in Firmicutes (100%).
2. **Anti-σ factor:** Members of ECF259 encode a putative AS factor with a DUF4179 and one transmembrane helix (94.6%) in position +1.
3. **Genetic context conservation:** Members of ECF259 share genetic context with proteins from ABC transporters – an ABC transporter (Pfam: ABC_tran) and a binding-protein-dependent transport system inner membrane component (Pfam: BPD_transp_1).
4. **Promoter motif conservation:** Predicted target promoter motifs contain GAACC in -35 and a T-rich -10.
5. **Summary:** Members of ECF259 are present in Firmicutes. DUF4179-containing AS factors located in +1 might regulated them. These ECFs contain ABC transporters in their genetic context, which might be part of the cell response they regulate.

**ECF260**

1. **General description:** Members of ECF260 are present in Firmicutes (100%).

2. **Anti-σ factor:** Members of ECF260 encode putative AS factors with one transmembrane helix (66.67%) in position +1. These AS factors contain DUF4179 in ECF260s1.
3. **Genetic context conservation:** Genetic contexts of members of ECF260 encode an ABC transporter.
4. **Promoter motif conservation:** Promoter motifs are conserved and contain CGAAC in -10 and GTATA in -35.
5. **Summary:** Likewise ECF258 and ECF259, members of ECF260 are present in Firmicutes, and DUF4179-containing AS factors located in +1 might regulate them. These ECFs contain ABC transporters in their genetic context, which might be part of the cell response they regulate.

**ECF261**

1. **General description:** Proteins from ECF261 are present in Firmicutes (100%).
2. **Anti-σ factor:** Members of ECF261 encode putative AS factors with DUF3600 and one transmembrane helix (75%) in +1. Putative AS factors of subgroup ECF261s2 differ from the rest – even though they contain one transmembrane helix that TopCons is not able to predict, they lack ~50 amino acids in N-terminal that the putative AS factors of the remaining groups contain.
3. **Genomic context conservation:** Other than this AS factor, likewise neighboring groups, members of ECF261 encode ABC transporters in their genomic context.
4. **Promoter motif conservation:** Predicted target promoter of ECF261 contain a conserved (T|C)GAAC in -35 and AATATA in -10.
5. **Summary:** ECF261 is a new group that seems to be regulated by a new kind of AS factor with DUF3600 and one transmembrane helix. ABC transporters are encoded in the neighborhood of members of this group.

**ECF262**

1. **General description:** Members of ECF262 belong to Firmicutes order Clostridiales, and contain C-terminal extensions (except ECF262s4) with TM helices in some proteins. The C-terminal extension contains various domains such as POM121, ribosomal protein L35, DUF2749 or TFIIB zinc-binding.
2. **Genomic context conservation:** Only ten proteins from representative organisms are present in this group. ECFs from ECF262s2 and two of the members of ECF262s3 are encoded in consecutive loci and co-transcribed. It is possible that they are involved in a similar pathway or that they regulate each other, potentially via the contact with the partner C-terminal extension.
3. **Anti-σ factor:** No AS factors were found in the context of members of ECF262.

4. **Promoter motif conservation:** Predicted target promoter motifs are not conserved.
5. **Summary:** Members of ECF262 might be regulated by their C-terminal extension. They are sometimes encoded with other members of the same group, which indicates a cross-regulation by the other ECF or a function in the same pathway.

**ECF263**

1. **General description:** Members of ECF263 are present in Proteobacteria (89.8%), Verrucomicrobia (8.16%) and Planctomycetes (2.04%). Proteins from ECF263 contain a C-terminal extension with five transmembrane helices in ECF263s1, ECF263s2 and ECF263s4 (~230 aa) and with seven transmembrane helices in ECF263s3 (~330 aa). This extension has homology to fatty acid desaturases in some examples from ECF263s4 and ECF263s2.
2. **Anti-σ factor:** Proteins from ECF263 from subgroups ECF263s1 and ECF263s2 are associated with AS factors with one transmembrane helix (81.25%).
3. **Genetic context conservation:** Members of ECF263 encode a putative prokaryotic signal transducing domain (Pfam: DUF2007) in -1. Conserved proteins in ECF263 include a thioesterase (ECF263s2 and -2 of ECF263s1), a N-acetylmuramoyl-L-alanine amidase (ECF263s2) and a RNA methylase (ECF263s2), a endonuclease/exonuclease/phosphatase (ECF263s1) and an arginine-tRNA-protein transferase (ECF263s1).
4. **Promoter motif conservation:** Predicted target promoter motifs are conserved and contain TGCAC(A/C) in -35 and CGTC in -10.
5. **Summary:** ECF263 contains proteins with a long C-terminal extension, which might contain five or seven transmembrane helices. Since putative AS factor with DUF2007 are encoded in position -1 in subgroups ECF263s1 and ECF263s2, and it is possible that these AS factors are encoded somewhere else in the genome of the ECF263s3 and ECF263s4-containing organisms, the C-terminal extension of ECF263 might not have a regulatory role.

**ECF264**

1. **General description:** Proteins from ECF264 are present in Firmicutes (100%), mostly from family Paenibacillaceae. Members of ECF264 contain a ~350 C-terminal extension with a DUF1835.
2. **Anti-σ factor:** Proteins from ECF264 are not associated with AS factors.
3. **Genomic context conservation:** The only conserved gene in the genetic context of members of ECF264 encodes part of an ABC transporter.

4. **Promoter motif conservation**: Promoter motifs are conserved and contain GTAACCTT in -35 and GGCACATT in -10.
5. **Summary:** ECF264 contains a C-terminal extension with a conserved DUF1835 domain. Since no AS factor can be idenfied in the genetic context of members of this group, it is likely that proteins from ECF264 are regulated by their C-terminal extension.

**ECF265**

1. **General description:** Proteins from ECF265 have homology to proteins from original ECF30 (46.6%), ECF55 (5.84%) and ECF112 (0.51%), and are present in Firmicutes (95.80%) and Actinobacteria (3.50%).
2. **Regulation:** Members of ECF265 are associated with AS factors with one transmembrane helix (51.87%) in +1. In the case of ECF265s4 and ECF265s7, which appear primary in the *Bacillus* genus, there is no apparent AS factor in their genetic context. In these groups, ECFs share the genetic neighborhood with a transcriptional repressor from PadR family (+1) and a cell cycle protein (+2) involved in peptidoglycan synthesis such as FtsW, RodA or SpoVE (Pfam: PadR and FTSW_RODA_SPOVE) in +1 and +2, respectively. PadR is involved in negative regulation of the phenolic acid metabolism and RodA is a peptidoglycan glycosyltransferase involved in peptidoglycan biosynthesis. It is possible that in these two subgroups the AS factor is substituted by transcriptional control.
3. **Genomic context conservation:** Apart from the proteins mentioned above, the only conserved element in the genomic neighborhood of members of ECF265 are the inner-membrane component of an ABC transporter (Pfam: BPD_transp_1) (ECF265s24).
4. **Promoter motif conservation:** Promoter motifs are subgroup-dependent conserved and usually contain AAC in -35 and CGTC in -10; in agreement with the prediction for original ECF30 (Staroń et al., 2009).
5. **Summary:** In summary, ECF265 seems to be regulated by putative AS factors encoded in +1 with diverse unknown domains, in contrast to original ECF30 (Staroń & Mascher, 2010). Two subgroups, ECF265s4 and ECF265s7, seem to be regulated by a PadR transcriptional repressor encoded in +1 and involved in phenolic acid metabolism. Since RodA is encoded in position +2 in ECF265s4 and ECF265s7, proteins from these subgroups might be involved in peptidoglycan biosynthesis.

**ECF266**

1. **General description:** Members of ECF266 are present in Firmicutes (100%) from order Clostridiales.

2. **Anti-σ factor:** Members of ECF266 contain a protein with a beta-propeller domain (Pfam: Beta_propel) and one inner transmembrane helix (84.38%) in +1. Beta-propeller domains are related to WD40 repeats (source: Pfam), which appear in the sensory domain of AS factors from ECF39 and STKs from ECF62. These proteins could function as AS factors of a new type.
3. **Genomic context conservation:** Conservation of the genetic context of members of ECF266 does not extend beyond this point.
4. **Promoter motif conservation:** Predicted target promoter motifs are conserved and contain TGCAACA in -15 and ACTCT.
5. **Summary:** Members of ECF266 encode a one-transmembrane helix protein with an extracytoplasmic beta-propeller domain in +1. This protein could serve as an AS factor. Proteins from ECF266 seem to be autoregulated.

**ECF267**

1. **General description:** Proteins from ECF267 are present in Proteobacteria (100%) from genus *Sorangium* and *Chondromyces*.
2. **Regulation:** Proteins of ECF267 might be regulated by FecR-like AS factors fused to tetratricopeptide repeats encoded in position +1. Members of ECF267 contain an average of 2.67 and 1.33 protein kinases in their genetic contexts in ECF267s1 and ECF267s2, respectively. The position where this protein is encoded is not conserved.
3. **Promoter motif conservation:** Target promoter motifs are not conserved.
4. **Summary:** ECF267 is regulated by FecR-like AS factors without a FecA-like outer membrane transporter or another TonB-dependent receptor. Members of this groups might be regulated through phosphorylation given the presence of 1-3 protein kinases in their genetic context.

**ECF268**

1. **General description:** Proteins from ECF268 are present in Acidobacteria (100%).
2. **Anti-σ factor:** Proteins from ECF268 might be regulated by a putative AS with one transmembrane helix and a putative zinc-finger encoded in position +1.
3. **Promoter motif conservation:** The information content of the predicted target promoter motifs is low, but both subgroups from ECF268 contain the same motifs: GGGAAC in -35 and GGTGT in -10.
4. **Summary:** ECF268 seems to be regulated by ZAS factors.

**ECF269**

1. **General description:** Proteins from ECF269 are present in Proteobacteria (100%) mostly in genus *Legionella*.
2. **Anti-σ factor:** Members of this group are not encoded with proteins of unknown function and two transmembrane helices (69.23%) in position -1. While looking at their MSA, all proteins encoded in position -1 are homologs and could be AS factors. Proteins in position -2 encode a flavinator of succinate dehydrogenase and position -3 contains a protein with an aminomethyltransferase folate-binding domain.
3. **Promoter motif conservation:** Predicted target promoter motifs are not conserved.
4. **Summary:** In conclusion, members of ECF269 could be regulated by a new type of AS factors with two transmembrane helices and no homology to AS factors of the original library.
5. **Summary:** Members of ECF269 could be regulated by a new type of AS factors with two transmembrane helices and no homology to AS factors of the original library.

**ECF270**

1. **General description:** Proteins from this group are present in Deltaproteobacteria (37.5%), Planctomycetes (12.5%) from genus *Zavarzinella*, Ignavibacteriae (25%) and Nitrospirae (25%).
2. **Anti-σ factor:** Proteins from ECF270 encoded a soluble (100%) ZAS in position +1, except in ECF270s2, where this protein is fused to the ECF coding sequence in 60% of the cases.
3. **Genomic context conservation:** The conservation does not extend beyond this protein.
4. **Promoter motif conservation:** Promoter motifs are not conserved.
5. **Summary:** ECF270 is a new group regulated by a soluble ZAS encoded in +1.

**ECF271**

1. **General description:** Members of ECF271 are homologous to proteins from original ECF03 (2.5%) and are present in Chloroflexi (81.82%) (ECF271s1) and Acidobacteria (9.09%) from the genus *Bryobacter* (ECF271s2).
2. **Anti-σ factor:** Members of this group encode a soluble (100%) zinc-finger putative AS factor in +1.
3. **Promoter motif conservation:** Promoter motifs are only conserved in ECF271s1 and contain GCTGCTG in -35 and GXTCA in -10.
4. **Summary:** Unlike original ECF03 (Staroń et al., 2009), members of ECF271 are regulated by a soluble ZAS encoded in +1.

**ECF272**

1. **General description:** Members of ECF272 are present in Gammaproteobacteria (97%) and Gemmantimonadetes from genus *Gemmatimonas* (3%).
2. **Anti-σ factor:** Members of ECF272 might be regulated by single-transmembrane helix putative AS factors encoded in position +1. Most of these putative AS factors have homology to RskA from *M. tuberculosis*.
3. **Genomic context conservation:** The genetic context conservation extends to position +2, which encodes a putative adhesine (Pfam: DUF4097). Other proteins conserved in the genetic context of members of ECF272 are a DnaB-like helicase (ECF272s3 and +3 of ECF272s4), a ribosomal protein L9 (ECF272s3 and ECF272s4), a ribosomal protein S18 (ECF272s3 and ECF272s4), an alanine racemase (+4 of ECF272s4), a single-strand binding protein (ECF272s4), a RNA pseudouridylate synthase (ECF272s5). This genetic context conservation indicates that members of ECF272 might be involved in translation, at least for members of subgroups ECF272s3 and ECF272s4.
4. **Promoter motif conservation:** Predicted target promoter motifs are conserved and contain TTTAAACCTTT in -35 and a less conserved CG(A|T)CTAA in -10.
5. **Summary:** ECF272 is a new group regulated by RskA-like, one TMH, putative AS factors encoded in position +1. A putative adhesin is encoded in position +2, indicating that the suface contacts of the bacteria are essential for the transduction mechanisms of members of ECF272. Given the genetic context conservation of subgroups ECF272s3 and ECF272s4, these could be regulating translation.

**ECF273**

1. **General description:** Members of ECF273 are present in Proteobacteria (100%) from order Xanthomonadales.
2. **Anti-σ factor:** Members of ECF273 could be regulated by a putative RskA-like AS factor with one transmembrane helix (80%) encoded in position +1.
3. **Genomic context conservation:** Other proteins conserved in members of ECF273 included a protein with a PDZ domain in +2. This protein could be coupling the sensing of unfolded OMP to the degradation of the AS factor as in the case or RseP and DegS proteases in *E. coli* (Hasselblatt et al., 2007; Inaba et al., 2008). Other conserved proteins include the partner-binding domain of proteins from the glycine cleavage system (a glycine cleavage T-protein and a glycine cleavage H-protein), proteins related to stomatins (SPFH domain / Band 7 protein, NfeD-like protein) and GMC oxidoreductase and a beta-lactamase.

4. **Promoter motif conservation:** Promoter motifs are only predicted for ECF273s1, with AGTATAGG in -35 and CC in -10.
5. **Summary:** Members of ECF273 contain an RskA-like single-membrane pass AS factor in +1 and a protein with a PDZ domain in +2. Given the conservation of two of the proteins in the genetic context, members of ECF273 could be responding to cell wall stress caused by different substances and whose response could involve the expression of the glycine cleavage system, beta-lactamases, stomatins, and GMC oxidoreductases.

**ECF274**

1. **General description:** Members of ECF274 are present in Bacteroidetes (100%).
2. **Anti-σ factor:** Members of ECF274 contain a one-transmembrane helix (81.25%) AS factor in position +1.
3. **Genomic context conservation:** The conservation of the genetic neighborhood could not be assessed due to the low number of proteins included in ECF274. Proteins encoded in position +2 of ECF274s1 seem to be adhesins, similarly as in the Proteobacterial neighboring group ECF273.
4. **Promoter motif conservation:** Promoter motifs are conserved and contain CAACCTTT in -35 and CTCTTT in -10.
5. **Summary:** Members of ECF274 might be regulated by putative AS factors with one transmembrane helix encoded in position +1. Elucidating the genetic neighborhood conservation requires more sequences from ECF274.

**ECF275**

1. **General description:** Members of ECF275 are present in Bacteroidetes (99.61%).
2. **Anti-σ factor:** Members of ECF275 encode a single-transmembrane helix (51.95%) putative AS factor in position +1. This protein contains a RskA-like domain, HEAT repeats and/or a DUF3379 in some case. Neverthless, an MSA of the putative AS revealed their conservation and the presence of one transmembrane helix.
3. **Genomic context conservation:** The genetic context conservation extends only to the GIN domain of a putative auto-transporter adhesion (+3 of ECF275s1), a Lon protease (ECF275s1), a cytidylate kinase (ECF275s1).
4. **Promoter motif conservation:** Promoter motifs are conserved and contain TGxAAC in -35 and a less conserved GTC in -10.

5. **Summary:** Members of ECF275 are present in Bacteroidetes and might be regulated by putative AS factors encoded in position +1. Proteins from ECF275 are annotated with two possible starting methionines. In the longest version, the sequences from ECF275 contain a conserved N-terminal cysteine.

**ECF276**

1. **General description:** Some proteins from this group (5.04%) have homology to original ECF30. Proteins from ECF276 are represented in Firmicutes (60.98%) (ECF276s1, ECF276s6 and ECF276s7), Deltaproteobacteria (29.27%) (ECF276s3, ECF276s4 and ECF276s5) and Planctomycetes (9.76%) (ECF276s8). Members of ECF276s3 contain a C-terminal extension.
2. **Anti-σ factor:** Members of ECF276 might be regulated by a zinc-finger-containing AS factor with one (29.27%) or two (56.1%) TM helices encoded in position +1. The genomic context conservation does not extend beyond this point.
3. **Promoter motif conservation:** Predicted target promoter motifs have a low information content and usually display AAC in -35 and TCT in -10.
4. **Summary:** ZAS factors with one or two TM helices located in position +1 migth regulate ECF276.

**ECF277**

1. **General description:** Members of ECF277 are present in Acidobacteria (71.43%), Bacteroidetes (14.29%) from genus Prevotella and Proteobacteria (14.29%).
2. **Anti-σ factor:** Members of ECF277 might be regulated by putative AS factors with a zinc-finger encoded in position +1. These AS factors contain one transmembrane helix (85.86%) and HEAT repeats in the extracytoplasmic region of putative AS factors of ECF277s1. Proteins with HEAT repeats are found in position +2 of ECF277s3, and +2 and +3 of ECF277s5, suggesting that these repeats are essential for the signaling mechanisms that activate members of ECF277.
3. **Genomic context conservation:** Other conserved proteins encoded in the genetic context of members of ECF277 included a protein with PDZ domain in +2 of ECF277s1 (0.94 copies per ECF) and is also present in the genetic context of members of ECF277s5 (0.5 copies per ECF). PDZ domain is present in DegS, the site-1 protease that senses misfolded proteins and degrades RseA, the AS factor of RpoE in *E. coli* (Wilken, Kitzing, Kurzbauer, Ehrmann, & Clausen, 2004).
4. **Promoter motif conservation:** Predicted target promoter motifs are not conserved.

5. **Summary:** Members of ECF277 might be regulated by single-membrane pass AS factors that contain HEAT domains in ECF277s1. In other subgroups, the HEAT domain is present in proteins encoded in +2 and/or +3, suggesting that this is, indeed, part of the sensing domain. Members of ECF277 could sense unfolded OMP since some subgroup (ECF277s1 and ECF277s5) contain a protein with a PDZ domain in their genetic neighborhood.

**ECF278**

1. **General description:** Members of ECF278 are present in Proteobacteria (83.33%) from genus *Desulfovibrio* and *Syntrophobacter* (ECF278s1), and Acidobacteria (16.67%) from genus *Chloracidobacterium* (ECF278s2).
2. **Anti-σ factor:** Members of ECF278 might be regulated by putative AS factors with one transmembrane helix (100%) encoded in position +1. These putative AS factors from ECF278s1 contain the glycogen recognition site of AMP-activated protein kinase in the periplasm. This proteins could be regulating the ECF278-dependent signal transduction according to the presence of glycogen.
3. **Promoter motif conservation:** Predicted target promoter motifs are only conserved for ECF278s1 and contain GTGAC in -35 and GTATA in -10.
4. **Summary:** Members of ECF278 are regulated by an AS factor with one transmembrane helix encoded in position +1 that could be sensing the presence of glycogen, at least in ECF278s1.

**ECF279**

1. **General description:** Members of ECF279 are present in Bacteroidetes (100%) from genus *Bacteroides* and *Prevotella*.
2. **Anti-σ factor:** Members of ECF279 might be regulated by AS factors encoded in position +1 with a DUF5056. These AS factors contain two transmembrane helices (52.5% according to TopCons, but present in most of the putative AS factors when looking at the MSA).
3. **Genomic context conservation:** Genetic neighborhood conservation is extensive and includes a dihydrofolate reductase, a biopolymer transport protein ExbD/TolR, a 16S rRNA methyltransferase RsmB/F, a peptidase M64 (ECF279s2), a thymidylate synthase (ECF279s2), an acetyltransferase (GNAT) family (ECF279s2) and a Lrp/AsnC ligand binding domain fused to an AsnC-type helix-turn-helix domain (ECF279s2).

4. **Promoter motif conservation:** Predicted target promoter motifs included TTGCAA in -35 and CGTCTAA in -10.
5. **Summary:** Members of ECF279 might be regulated by a DUF5056 AS factor with two transmembrane helices encoded in position +1. The genetic neighborhood of members of ECF279 is conserved and encodes proteins for the import of biomolecules and rRNA modification.

**ECF280**

1. **General description:** Members of ECF280 are present in Proteobacteria (100%) from family Caulobacteraceae.
2. **Anti-σ factor:** Members of ECF280 might be regulated by putative AS factors with two transmembrane helices (73.33%) encoded in position +1.
3. **Genomic context conservation:** Genetic context conservation is broad and includes a ketopantoate hydroxymethyltransferase (+2), a tetratricopeptide repeat-like domain (+3), a PQQ-like domain (+4), a 50S ribosome-binding GTPase fused to a KH-domain-like of EngA bacterial GTPases (+5), a glutamine amidotransferase, a phosphoribosyltransferase, and an ArsC-like protein. Interestingly, members of ECF280s1 and ECF280s3 encoded a second ECF σ factor in position -3 or -4 of their genetic neighborhood. These ECFs, even though part of the ECF library, are ungrouped.
4. **Promoter motif conservation:** Predicted target promoter motifs are not conserved.
5. **Summary:** Members of ECF280 might be regulated by putative AS factors with two transmembrane helices encoded in position +1. Genetic context is conserved and contains another ungrouped ECFs in position -3 or -4 of ECF280s1 and ECF280s3. Given the overall conservation of ECF280 and the relatedness of the organisms where it appears (all in Caulobacteraceae), it is possible that members of ECF280 fulfill the same function and it is activated by the same input signal.

**ECF281**

1. **General description:** Proteins from ECF281 have homology to original groups ECF20 (10.45%) and ECF113 (0.85%), and are present in Firmicutes (41.44%), Actinobacteria (23.42%), Deinococcus-Thermus (18.92%) from genus *Deinococcus* and Proteobacteria (16.22%).
2. **Anti-σ factor:** Most of the subgroups contain putative AS factors with one TM helix (79.69%) and a zinc-finger encoded in +1. In subgroups ECF281s3 and ECF281s4, the putative AS factor could be fused to a DUF4349. Instead, subgroups ECF281s1, ECF281s5, ECF281s7 and ECF281s10 do not contain AS factors

in their genomic context. It is likely that the AS factors of these subgroups are encoded somewhere else in the genome.

3. **Genomic context conservation:** Other proteins conserved are a heavy-metal resistance protein (+2 of ECF281s2), a tRNA-dihydrouridine synthase (-1 of ECF281s6), and a protein from family FUN14 (-1 of ECF281s5), which appears in Eubacteria, Archaea and Eukaryotes and is involved in mitophagy induced by hypoxia in mammalian cells (L. Liu et al., 2012).
4. **Promoter motif conservation:** Predicted target promoter motifs are conserved and contain GGAACTT in -35 and a less conserved GTCTAA in -10. This motif is similar, but not identical, to the target promoter motif in the original group ECF20 (Rhodius et al., 2013).
5. **Summary:** Members of ECF281 are regulated by zinc-finger AS factors with one transmembrane helix encoded in +1. Members of ECF281 could be involved in response to heavy metals and their detoxification. This function inherits from original ECF20 (Staroń et al., 2009), even though the only characterized member of original ECF20 is now classified against ECF291.

**ECF282**

1. **General description:** Members of ECF282 are present in Actinobacteria (100%).
2. **Anti-σ factor**: No evident AS factor was found in the genomic context of members of ECF282.
3. **Genomic context conservation:** The genetic context of members of ECF282 is not conserved.
4. **Studied members:** One characterized member of ECF282, $\sigma^{AntA}$ (ECF282s2) from *Streptomyces albus,* is not regulated by AS factors. Instead, $\sigma^{AntA}$ is regulated at the transcriptional level as part of the live cycle of *S. albus*, and it is a target of ClpXP proteolysis, which needs a C-terminal AA motif (Seipke, Patrick, & Hutchings, 2014). Nevertheless, the C-terminal AA is only present in ECF282s2 – therefore, members of ECF282s1 could be regulated differently.
5. **Promoter motif conservation:** Predicted target promoter motifs are not conserved. Indeed, $\sigma^{AntA}$ is not autoregulated.
6. **Summary:** Previous reports found that homologs of $\sigma^{AntA}$ are only found in species containing the *ant* gene clusters for the production of the antibiotic antimycin (Seipke et al., 2014). Indeed, in this work, we also define ECF282 as a new group of ECF σ factors. $\sigma^{AntA}$ (ECF282s2) from *Streptomyces albus*, is regulated transcriptionally and by ClpXP proteolysis that targets an AA motif in C-terminal.

**ECF283**

1. **General description:** Members of ECF283 are present in Proteobacteria (100%).
2. **Genomic context conservation:** Members of ECF283 encode a transmembrane (~100% in the MSA) protein kinase in position -1 or +2 in ECF283s5. Protein sequences of members of ECF283 contain a Ser/Thr rich region in the last part of the linker between $\sigma_2$ and $\sigma_4$ domain. It is possible that this is the phosphorylation site of the ECFs from ECF283 and that this phosphorylation triggers a change on the activity of members of ECF283.
3. **Promoter motif conservation:** Predicted target promoter motifs contain CCTT in -35 and AG in -10.
4. **Summary:** Members of ECF283 seem to be regulated by protein kinases encoded in their genetic neighborhood, which could be phosphorylating a Ser/Thr rich area at the beginning of the linker between $\sigma_2$ and $\sigma_4$ domain.

**ECF284**

1. **General description:** Members of ECF284 are present in Actinobacteria (100%).
2. **Anti-σ factor:** A putative AS factor with two transmembrane helices (57.14%) and some homology to AhpC/TSA family (Pfam: AhpC-TSA) was found in position +1 of the proteins from ECF284. An MSA of the proteins in this position revealed their homology.
3. **Genomic context conservation:** Genetic context conservation does not extend beyond the putative AS factor.
4. **Promoter motif conservation:** Predicted target promoter motifs are conserved and contain GAATCCTTT in -35 and CTCTT in -10.
5. **Summary:** Members of ECF284 might be regulated by putative AS factors encoded in position +1. These AS factors typically contain two transmembrane helices.

**ECF285**

1. **General description:** Members of ECF285 are present in Proteobacteria (100%).
2. **Anti-σ factor:** Members of ECF285 encode 2-transmembrane helix proteins (67.39%) with a DUF2892 in -1. These proteins could be a new type of AS factor with no homology to AS factors from the original library (Staroń et al., 2009).
3. **Genomic context conservation:** Genetic context conservation extends to a DUF692 (+2 of ECF285s5), a putative DNA-binding domain (Pfam: DUF2093) (+3 of ECF285s5), a RNA 2'-O ribose methyltransferase

(ECF285s5), a PBP (Pfam: PBP_like_2) that might bind to phosphatidylethanolamine and have a role in membrane signal transduction (Banfield, Barker, Perry, & Brady, 1998) (ECF285s5), an exonuclease (Pfam: RNase_T) (ECF285s5), the periplasmic component of an ABC transporter (ECF285s5) and a PhoU, mainly as part of the putative AS factor in position -1, but also possibly involved in phosphate uptake (ECF285s5).

4. **Promoter motif conservation:** Predicted target promoter motifs of members of ECF285 contain AA in -35 and CCTCT in -10.
5. **Summary:** members of ECF285 might be regulated by a new type of single-transmembrane helix-containing AS factors encoded in position -1.

**ECF286**

1. **General description:** Some members of this group are homologous to proteins from the original group ECF20 (2.2%). Members from ECF286 are present in Actinobacteria (100%).
2. **Anti-σ factor:** Proteins from ECF286 are encoded together with several soluble proteins from Asp23 family (Pfam: Asp23), which is one of the most abundant proteins of *S. aureus* and is related with cell envelope homeostasis (Müller et al., 2014). Deletion of Asp23 in *S. aureus* leads to upregulation of the cell wall stress genes (Müller et al., 2014); therefore, this protein could function as AS factor.
3. **Genomic context conservation:** Other than Asp23, the genetic context of ECF286s4 contains a conserved RibD C-terminal domain in -1 and an AFG1-like ATPase in -2.
4. **Promoter motif conservation:** Predicted target promoter motifs are conserved and contain a conserved GAC in -35 and TC in -10.
5. **Summary:** In contrast to original ECF20 (Staroń et al., 2009), members of ECF286 do not seem to be regulated by canonical AS factors. Instead, they contain several Asp23-like proteins, indicating that members of ECF286 are involved in cell envelope homeostasis as in the case of Asp23 in S. *aureus* (Müller et al., 2014). Provided that these Asp23 proteins function as AS factors of members of ECF286, it remains unclear why each ECF from ECF283 encodes an average of 3.33 Asp23 proteins in its genetic neighborhood.

**ECF287**

1. **General description:** Members of ECF287 are present in Actinobacteria (100%) and contain a conserved ~80 aa C-terminal extension of an unknown domain.

2. **Anti-σ factor:** No putative AS factor was found in the genetic context of members of ECF287.
3. **Genomic context conservation:** Members of ECF287 are encoded with transcriptional regulators from MerR family (0.44 and 0.22 per ECF of ECF287s1 and ECF287s2, respectively) and TetR family (0.22 and 0.57 per ECF of ECF287s1 and ECF287s2, respectively). Other conserved proteins in the genetic context of members of ECF287 include a soluble alpha/beta hydrolase encoded in position -1.
4. **Promoter motif conservation:** Predicted target promoter motifs are only conserved in -35, with a common TCG in ECF287s1 and ECF287s2.
5. **Summary:** Members of ECF287 could be regulated by their C-terminal extension since they do not contain putative AS factors in their genetic context. The regulation could be carried out by disulfide bridges since the ECFs from ECF287 contain several conserved cysteine residues, also within their C-termini. A similar mechanism controls the activity of CorE2 from *M. xanthus* (Marcos-Torres et al., 2016; Torres et al., 2018) (ECF238s15) and SigK from *M. tuberculosis* (Shukla et al., 2014) (ECF19s2).

**ECF288**

1. **General description:** Members of ECF288 appear in Firmicutes (100%) and contain a C-terminal extension of ~100aa. Interestingly, both $\sigma_4$ domain and the C-terminal extension of members of ECF288 contain multiple cysteine residues, which indicates that disulfide-bridge formation could be in charge of the modulation of their activity, as in the case of SigK from *M. tuberculosis* (Shukla et al., 2014) (ECF19s2) and CorE2 from *M. xanthus* (Marcos-Torres et al., 2016; Torres et al., 2018) (ECF238s15).
2. **Anti-σ factor:** No putative AS factors were found in the genetic context of members of ECF288.
3. **Genomic context conservation:** There is a soluble conserved DUF2461-containing protein in +1, which might be part of the regulatory mechanism of ECF288.
4. **Promoter motif conservation:** Predicted target promoter motifs are highly conserved, with TGTCACA in -35 and TGTCTAAT in -10.
5. **Summary:** As in the case of ECF287, proteins ECF288 contains C-rich C-terminal extensions that could be in charge of their activation or inactivation as in the case of CorE2 from *M. xanthus* (Marcos-Torres et al., 2016; Torres et al., 2018) (ECF238s15) and SigK from *M. tuberculosis* (Shukla et al., 2014) (ECF19s2).

**ECF289**

1. **General description:** Members of ECF289 are homologous to proteins from the original group ECF20 (42.27%) and are present in Proteobacteria (100%).

2. **Anti-σ factor:** Members of ECF289 encode a putative AS factor in position +1. This putative AS factor contains a von Willebrand domain fused to a DUF3520 in C-terminal and harbors one transmembrane helix (87.34%). In ECF289s6 the N-terminal domain ASD is truncated even though the rest of the protein is similar other putative AS factors from ECF289.
3. **Genomic context conservation:** Other conserved domains found in the genetic context of members of ECF289 are a trimethylamine methyltransferase (ECF289s1) and an aldehyde dehydrogenase (ECF289s4).
4. **Promoter motif conservation:** Predicted target promoter motifs are conserved and contain CCCAAGGA in -35 and CGTCTxA in -10. In subgroups ECF289s2 and ECF289s4 the up-element (AAAATTT) is more conserved than the -10 element. The overall conservation of the predicted binding motifs indicates that members of ECF289 self-regulate their transcription.
5. **Summary:** ECF289 is regulated by putative AS factors encoded in +1 with a DUF3520 fused to a von Willebrand domain. These AS factors are bound to the membrane by a single transmembrane helix. Original ECF20 was found close to von Willebrand factors, but they were not identified as AS factors (Staroń et al., 2009). New experiments will determine if these proteins are functioning as AS factors.

**ECF290**

1. **General description:** Members of this group have homology members of original ECF20 (95.41%) and appear in Proteobacteria (97.64%) and Planctomycetes (2.36% of the total) from family Planctomycetaceae.
2. **Anti-σ factor:** Proteins in +1 are putative AS factors (except in ECF290s3) and harbor one transmembrane helix (42.54%, ~100% of the sequences from subgroups except ECF290s3 in an MSA). These proteins do not appear in ECF290s3, which contains the coding sequence of flagellin instead.
3. **Genomic context conservation:** Conserved proteins encoded in the context of ECF290 include a membrane-bound heavy-metal resistance protein (+2 of ECF290s1 and ECF290s2), a diguanylate cyclase fused to a response regulator receiver domain (+3 of ECF290s2), an EF-hand (-1 of ECF290s2), a DUF983 (-2 of ECF290s2), a NUDIX domain (-3 of ECF290s2) from nucleoside diphosphate hydrolases. This genetic context, and especially the metal-resistance protein encoded in +2, is conserved in most groups from ECF290. Nevertheless, members of subgroup ECF290s3 are encoded near proteins involved in flagellum biosynthesis such as flagellin in +1, flagellar hook-associated protein 2 in +2 and FliS in +3.

4. **Promoter motif conservation:** Predicted target promoter motifs have low information content. The largest subgroups, ECF290s1 and ECF290s2, share an ACGG in -35 and CGT is conserved in several -10 elements. These motifs agree with the original group ECF20 (Staroń et al., 2009).
5. **Summary:** Members of ECF290 are associated with putative AS factors usually encoded in position +1. These putative AS factors contain one transmembrane helix and no Pfam domain. A conserved heavy metal resistance protein is encoded in +2, which points to metal detoxification as the function of members of ECF290. Subgroup ECF290s3 does not seem to encode any AS factor in its genomic context and it contains instead proteins for the synthesis of the flagellum.

**ECF291**

1. **General description:** Members of ECF291 have homology against proteins from original ECF20 (33.76%) and appear in Proteobacteria (98.45%) and Cyanobacteria (1.55%).
2. **Studied members:** One member of ECF291, CnrH from *Cupriavidus metallidurans* (ECF291s9), has been experimentally addressed. CnrH is encoded as part of a metal resistance determinant that includes *cnrYHX* and *cnrCBAT* operons (Grass, Fricke, & Nies, 2005). CnrH is sequestered by the AS factor CnrY, encoded in -2 from the ECF (Maillard et al., 2014). CnrY is a type II AS factor since it only contains two transmembrane helices that wrap around the ECF protein (Maillard et al., 2014). CnrX, the transmembrane heavy metal resistance protein encoded in -1, serves as the periplasmic sensor of $Ni^{2+}$ or $Co^{2+}$ (Trepreau et al., 2011). Upon sensing of these metals, CnrY in inhibited by an unknown mechanism and CnrH (Maillard et al., 2014). CnrH then directs the RNAP towards the *cnrYXH* promoter and the *cnrCBA* efflux pump, which secretes heavy metals (Grass et al., 2005).
3. **Genomic context conservation:** Indeed, this metal resistance determinant is conserved in our analysis of the genetic neighborhood of members of ECF291. Proteins from ECF291 encode a multicopper oxidase in +2, a copper resistance protein B precursor (CopB) in +2, a heavy metal resistance protein in -1. Members of ECF291s1 and ECF291s4 also encode CopC in –3 and CopD in -4. The Pfam domain of CnrY is only conserved in ECF291s9 and ECF291s12. Nevertheless, it is known that AS factors of this type have a variable sequence (Maillard et al., 2014) and the proteins in -2 of the rest of the groups contain one transmembrane helix (81.4%), have homology to AS factors from the original classification (Staroń et al., 2009) and align well with the CnrY-like AS factors from ECF291s9 and ECF291s12. Other conserved

proteins include a LuxR transcriptional regulator fused to a response regulator receiver domain (ECF291s1), a TonB dependent receptor (ECF291s4) and an E1-E2 ATPase.

4. **Promoter motif conservation:** Predicted target promoter motifs are only conserved in some subgroups such as ECF291s1, where there is TCTCC in -35 and TACGCA in -10. These motifs do not agree with the original group ECF20 (Rhodius et al., 2013). CnrH regulates the transcription of its operon, *cnrYXH*, in *C. metallidurans* (Grass et al., 2005).
5. **Summary:** Members of ECF291 are in charge of heavy metal resistance since they upregulate the expression of an efflux complex synthesized in coding sequences from +1 to +4. Members of this group are regulated by single-transmembrane helix AS factors encoded in -2 with homology to CnrY from *C. metallidurans* and a transmembrane protein with a heavy metal resistance domain encoded in -1 with homology to CnrX in *C. metallidurans*, which serves as a periplasmic sensor of the $Ni^{2+}$ or $Co^{2+}$.

**ECF292**

1. **General description:** Proteins from ECF292 appear in Actinobacteria, mainly in Micrococcales and Corynebacteriales.
2. **Anti-σ factor:** Genomic contexts of members of ECF292 contain an average of 2.08 soluble proteins (94.32%) from Asp23 family (Pfam: Asp23), which is one of the most abundant proteins *S. aureus* and is related with cell envelope homeostasis (Müller et al., 2014). The BLAST search recognizes proteins from the Asp23 family as AS factors since the ungrouped putative AS factor 1534 (YP_118547 from *Nocardia farcinica*) contains this domain in C-terminus (Staroń et al., 2009). Two different Asp23 proteins appear in our analysis, the ones with Asp23 domain and longer versions with an N-terminal extension whose length is subgroup-specific (~100 aa in ECF292s1 to ~20 in ECF292s2) and are encoded in position +1. Whether Asp23 family proteins are AS factors is unknown, but it could be possible since their deletion leads to upregulation of the cell envelope stress genes (Müller et al., 2014). Asp23 from *S. aureus* is attached to the membrane via DUF322 family protein (AmaP) (Müller et al., 2014). Interestingly, Asp23 proteins are also putative regulators of ECF286.
3. **Genomic context conservation:** In ECF292s4 there is also a conserved DUF2273-containing protein in -3.
4. **Promoter motif conservation:** Promoter motifs are not conserved.
5. **Summary:** ECF292 might be regulated by a new kind of soluble AS factor that belongs to the Asp23 family. This group could be involved in cell envelope stress response.

**ECF293**

1. **General description:** Proteins from ECF293 have homology to original groups ECF101 (11.35%), ECF117 (9.46%) and ECF13 (43.98%), and are present in Proteobacteria (53.03%), Bacteroidetes (29.65%), Actinobacteria (16.88%), Verrucomicrobia (0.22%) and Spirochaetes (0.22%). Proteins from ECF293 contain an extended linker between the $\sigma_2$ and $\sigma_4$ domains. Most subgroups also contain three cysteines in the $\sigma_4$ domain. It is unclear whether these modifications respect to the canonical ECF sequence affect the activity of members of ECF293.
2. **Anti-σ factor:** Similarly to proteins from original ECF13, ECF101 and ECF117 (Staroń et al., 2009), proteins from ECF293 encode a conserved soluble (97.92%) zinc-finger putative AS factor in position +1.
3. **Genomic context conservation:** Other conserved proteins are an ACP synthase III (-1 of ECF293s2), an acyl-CoA dehydrogenase (-1 of ECF293s8), a glycine cleavage system P-protein (+3 of ECF293s3), a DUF417 (+2 of ECF293s6), an EamA-like transporter (ECF293s2), a LysR transcriptional regulator (ECF293s2), OsmC (ECF293s8), a DUF3097 (ECF293s8), an universal stress protein (ECF293s8), an ABC transporter with a TOBE domain (ECF293s8) and a peptide methionine sulfoxide reductase (ECF293s6).
4. **Studied members:** Four members of ECF293 have been experimentally addressed - Ecf from *Neisseria gonorrhoeae* (ECF293s9), RpoE from *Nisseria meningitidis* (ECF293s9), PA3285 from *Pseudomonas aeruginosa* (ECF293s10) and CHU_3097 from *Cytophaga hutchinsonii* (ECF293s18). Ecf is overexpressed under treatment with hydrogen peroxide and is thought to participate in oxidative stress response since Ecf controls the expression of a methionine sulfoxide reductase and the genes from -4 to -1 in the locus of Ecf (Gunesekere et al., 2006). RpoE also controls the expression of its operon, which contains a methionine sulfoxide reductase (Huis in 't Veld et al., 2011). Even though this methionine sulfoxide reductase is not encoded in the proximity of Ecf in *N. gonorrhoeae* genome, a similar protein is encoded in the genetic context of members of ECF293s6. PA3285 from *P. aeruginosa* is predicted to be regulated by the FecI-like ECF PvdS (ECF05) (Chevalier et al., 2018). A short, soluble, zinc-binding anti-σ factor (MseR) has been identified for RpoE (Huis in 't Veld et al., 2011). CHU_3097 from *C. hutchinsonii* is released from the membrane upon induction with cellulose and upregulates proteins in charge of its degradation (X. Wang et al., 2019). An AS factor located in +1 and found in the membrane fraction has been proposed to be in charge of the shift (X. Wang et al., 2019). This AS factor does not contain any transmembrane helix when looking at the TopCons prediction.

5. **Promoter motif conservation:** Predicted target promoter motifs are not conserved among different subgroups. Indeed, Ecf from *N. gonorrhoeae* does not appear to be autoregulated (Gunesekere et al., 2006).
6. **Summary:** ECF293 merges three original groups (ECF13, ECF101, and ECF117) that encode a putative zinc-binding soluble AS factor in +1. Future analysis could determine if this protein is, indeed, an AS factor. ECF293 is likely involved in oxidative stress response.

**ECF294**

1. **General description:** Members of ECF294 are present in Proteobacteria (96.15%) and Acidobacteria (3.95%). Members of this group contain a ~120aa C-terminal extension with a SnoaL-like domain (Pfam: SnoaL_2). SnoaL is a small polyketide cyclase that catalyzes closure steps of the synthesis of polyketide antibiotics in *Streptomyces* spp. (Sultana et al., 2004). However, it has also been observed as part of ECF σ factors from original groups ECF56 and ECF41 (Huang et al., 2015; Staroń et al., 2009).
2. **Genomic context conservation:** The genetic context of members of ECF294 is not conserved beyond the ECF coding sequence.
3. **Promoter motif conservation:** Predicted target promoter motifs are not conserved among different subgroups.
4. **Summary:** Members of ECF294 compose a new group of SnoaL-like C-terminal extension σ factors with no homology to any original group. As in the case of ECF56 and ECF41 from the original ECF classification (Huang et al., 2015; Staroń et al., 2009), this C-terminal extension could be regulating ECF activity given the absence of any putative AS factor in the genetic context of members of ECF294.

**ECF295**

1. **General description:** Members of ECF295 are homologous to sequences of original ECF56 (100%) and are present in Firmicutes (78.57%) and Actinobacteria (21.43%). Members of ECF295 contain a C-terminal extension with homology to SnoaL-like domain (Pfam: snoaL_2) in ECF295s1 and ECF295s2, or to nuclear transport factor 2 (NTF2) domain (Pfam: NFT2) in ECF295s3. An MSA of the ECFs from ECF295 revealed that all the C-terminal extension are homologous to each other.
2. **Genomic context conservation:** Members of ECF295 contain an activator of Hsp90 ATPase homolog 1 in position +1 (0.71 copies per ECF). Hsp90 is a chaperone required for proper folding of proteins. Other than this protein and a tetratricopeptide repeat, the genetic context is not conserved.
3. **Promoter motif conservation:** Predicted target promoter motifs are not conserved.

4. **Summary:** Members of ECF295 might be regulated by their SnoaL-like C-terminal extension, and they could be controlling the activity of Hsp90. Members of ECF295 could be involved in stress response.

### References


Abdul-Majid, K.-B., Ly, L. H., Converse, P. J., Geiman, D. E., McMurray, D. N., & Bishai, W. R. (2008). Altered cellular infiltration and cytokine levels during early Mycobacterium tuberculosis sigC mutant infection are associated with late-stage disease attenuation and milder immunopathology in mice. *BMC Microbiology*, *8*(1), 151. http://doi.org/10.1186/1471-2180-8-151

Aires, J. R., Köhler, T., Nikaido, H., & Plésiat, P. (1999). Involvement of an Active Efflux System in the Natural Resistance of Pseudomonas aeruginosa to Aminoglycosides. *Antimicrobial Agents and Chemotherapy*, *43*(11), 2624–2628. http://doi.org/10.1128/AAC.43.11.2624

Akiyama, Y., Kanehara, K., & Ito, K. (2004). RseP (YaeL), an Escherichia coli RIP protease, cleaves transmembrane sequences. *The EMBO Journal*, *23*(22), 4434–4442. http://doi.org/10.1038/sj.emboj.7600449

Alice, A. F., López, C. S., Lowe, C. A., Ledesma, M. A., & Crosa, J. H. (2006). Genetic and Transcriptional Analysis of the Siderophore Malleobactin Biosynthesis and Transport Genes in the Human Pathogen Burkholderia pseudomallei K96243. *Journal of Bacteriology*, *188*(4), 1551–1566. http://doi.org/10.1128/JB.188.4.1551-1566.2006

Allen, M. S., Hurst, G. B., Lu, T.-Y. S., Perry, L. M., Pan, C., Lankford, P. K., & Pelletier, D. A. (2015). Rhodopseudomonas palustris CGA010 Proteome Implicates Extracytoplasmic Function Sigma Factor in Stress Response. *Journal of Proteome Research*, *14*(5), 2158–2168. http://doi.org/10.1021/pr5012558

Alvarez-Martinez, C. E., Lourenço, R. F., Baldini, R. L., Laub, M. T., & Gomes, S. L. (2007). The ECF sigma factor σT is involved in osmotic and oxidative stress responses in Caulobacter crescentus. *Molecular Microbiology*, *66*(5), 1240–1255. http://doi.org/10.1111/j.1365-2958.2007.06005.x

Amar, A., Pezzoni, M., Pizarro, R. A., & Costa, C. S. (2018). New envelope stress factors involved in σE activation and conditional lethality of rpoE mutations in Salmonella enterica. *Microbiology (Reading, England)*, *117*, 483. http://doi.org/10.1099/mic.0.000701

Andrews, S. C., Robinson, A. K., & Rodríguez-Quiñones, F. (2003). Bacterial iron homeostasis. *FEMS Microbiology Reviews*, *27*(2-3), 215–237. http://doi.org/10.1016/S0168-6445(03)00055-X

Banfield, M. J., Barker, J. J., Perry, A. C., & Brady, R. L. (1998). Function from structure? The crystal structure of human phosphatidylethanolamine-binding protein suggests a role in membrane signal transduction. *Structure (London, England : 1993)*, *6*(10), 1245–1254.

Barchinger, S. E., & Ades, S. E. (2013). Regulated Proteolysis: Control of the Escherichia coli σE-Dependent Cell Envelope Stress Response. In *Regulated Proteolysis in Microorganisms* (Vol. 66, pp. 129–160). Dordrecht: Springer Netherlands. http://doi.org/10.1007/978-94-007-5940-4_6

Barchinger, S. E., Pirbadian, S., Sambles, C., Baker, C. S., Leung, K. M., Burroughs, N. J., et al. (2016). Regulation of Gene Expression During Electron Acceptor Limitation and Bacterial Nanowire Formation in Shewanella oneidensis MR-1. *Applied and Environmental Microbiology*, *82*(17), AEM.01615–16–5443. http://doi.org/10.1128/AEM.01615-16

Barik, S., Sureka, K., Mukherjee, P., Basu, J., & Kundu, M. (2010). RseA, the SigE specific anti-sigma factor of Mycobacterium tuberculosis, is inactivated by phosphorylation-dependent ClpC1P2 proteolysis. *Molecular Microbiology*, *75*(3), 592–606. http://doi.org/10.1111/j.1365-2958.2009.07008.x

Bell, N., Lee, J. J., & Summers, M. L. (2017). Characterization and in vivoregulon determination of an ECF sigma factor and its cognate anti-sigma factor in Nostoc punctiforme. *Molecular Microbiology*, *104*(1), 179–194. http://doi.org/10.1111/mmi.13620

Bibb, Maureen J, & Buttner, M. J. (2003). The Streptomyces coelicolor developmental transcription factor sigmaBldN is synthesized as a proprotein. *Journal of Bacteriology*, *185*(7), 2338–2345. http://doi.org/10.1128/JB.185.7.2338-2345.2003

Bibb, Maureen J, Domonkos, Á., Chandra, G., & Buttner, M. J. (2012). Expression of the chaplin and rodlin hydrophobic sheath proteins in Streptomyces venezuelae is controlled by σBldN and a cognate anti-sigma factor, RsbN. *Molecular Microbiology*, *84*(6), 1033–1049. http://doi.org/10.1111/j.1365-2958.2012.08070.x

Biville, F., Cwerman, H., Létoffé, S., Rossi, M. S., Drouet, V., Ghigo, J. M., & Wandersman, C. (2004). Haemophore-mediated signalling in Serratia marcescens: a new mode of regulation for an extra cytoplasmic function (ECF) sigma factor involved in haem acquisition. *Molecular Microbiology*, *53*(4), 1267–1277. http://doi.org/10.1111/j.1365-2958.2004.04207.x

Braun, V., & Mahren, S. (2005). Transmembrane transcriptional control (surface signalling) of the Escherichia coliFec type. *FEMS Microbiology Reviews*, *29*(4), 673–684. http://doi.org/10.1016/j.femsre.2004.10.001

Brutsche, S., & Braun, V. (1997). SigX of <Emphasis Type="Italic">Bacillus subtilis</Emphasis> replaces the ECF sigma factor FecI of <Emphasis Type=“Italic”>Escherichia coli</Emphasis> and is inhibited by RsiX. *Molecular and General Genetics MGG*, *256*(4), 416–425. http://doi.org/10.1007/s004380050585

Butcher, B. G., & Helmann, J. D. (2006). Identification of Bacillus subtilis sigmaW-dependent genes that provide intrinsic resistance to antimicrobial compounds produced by Bacilli. *Molecular Microbiology*, *60*(3), 765–782. http://doi.org/10.1111/j.1365-2958.2006.05131.x

Butcher, B. G., Bao, Z., Wilson, J., Stodghill, P., Swingle, B., Filiatrault, M., et al. (2017). The ECF sigma factor, PSPTO_1043, in Pseudomonas syringae pv. tomato DC3000 is induced by oxidative stress and regulates genes involved in oxidative stress response. *PloS One*, *12*(7), e0180340. http://doi.org/10.1371/journal.pone.0180340

Calamita, H., Ko, C., Tyagi, S., Yoshimatsu, T., Morrison, N. E., & Bishai, W. R. (2004). The Mycobacterium tuberculosis SigD sigma factor controls the expression of ribosome-associated gene products in stationary phase and is required for full virulence. *Cellular Microbiology*, *7*(2), 233–244. http://doi.org/10.1111/j.1462-5822.2004.00454.x

Campbell, E. A., Greenwell, R., Anthony, J. R., Wang, S., Lim, L., Das, K., et al. (2007). A conserved structural module regulates transcriptional responses to diverse stress signals in bacteria. *Molecular Cell*, *27*(5), 793–805. http://doi.org/10.1016/j.molcel.2007.07.009

Cao, M., & Helmann, J. D. (2004). The Bacillus subtilis extracytoplasmic-function sigmaX factor regulates modification of the cell envelope and resistance to cationic antimicrobial peptides. *Journal of Bacteriology*, *186*(4), 1136–1146. http://doi.org/10.1128/JB.186.4.1136-1146.2004

Cao, M., Bernat, B. A., Wang, Z., Armstrong, R. N., & Helmann, J. D. (2001). FosB, a cysteine-dependent fosfomycin resistance protein under the control of sigma(W), an extracytoplasmic-function sigma factor in Bacillus subtilis. *Journal of Bacteriology*, *183*(7), 2380–2383. http://doi.org/10.1128/JB.183.7.2380-2383.2001

Cao, M., Kobel, P. A., Morshedi, M. M., Wu, M. F. W., Paddon, C., & Helmann, J. D. (2002a). Defining the Bacillus subtilis σW regulon: a comparative analysis of promoter consensus search, run-off transcription/macroarray analysis (ROMA), and transcriptional profiling approaches. *Journal of Molecular Biology*, *316*(3), 443–457. http://doi.org/10.1006/jmbi.2001.5372

Cao, M., Salzberg, L., Tsai, C. S., Mascher, T., Bonilla, C., Wang, T., et al. (2003). Regulation of the Bacillus subtilis extracytoplasmic function protein sigma(Y) and its target promoters. *Journal of Bacteriology*, *185*(16), 4883–4890. http://doi.org/10.1128/JB.185.16.4883-4890.2003

Cao, M., Wang, T., Ye, R., & Helmann, J. D. (2002b). Antibiotics that inhibit cell wall biosynthesis induce expression of the Bacillus subtilisσW and σM regulons. *Molecular Microbiology*, *45*(5), 1267–1276. http://doi.org/10.1046/j.1365-2958.2002.03050.x

Chao, K. L., Lim, K., Lehmann, C., Doseeva, V., Howard, A. J., Schwarz, F. P., & Herzberg, O. (2008). The Escherichia coli YdcF binds S-adenosyl-L-methionine and adopts an alpha/beta-fold characteristic of nucleotide-utilizing enzymes. *Proteins*, *72*(1), 506–509. http://doi.org/10.1002/prot.22046

Chastanet, A., & Losick, R. (2007). Engulfment during sporulation in Bacillus subtilis is governed by a multi-protein complex containing tandemly acting autolysins. *Molecular Microbiology*, *64*(1), 139–152. http://doi.org/10.1111/j.1365-2958.2007.05652.x

Chatterjee, A., Drews, L., Mehra, S., Takano, E., Kaznessis, Y. N., & Hu, W.-S. (2011). Convergent transcription in the butyrolactone regulon in Streptomyces coelicolor confers a bistable genetic switch for antibiotic biosynthesis. *PloS One*, *6*(7), e21974. http://doi.org/10.1371/journal.pone.0021974

Chevalier, S., Bouffartigues, E., Bazire, A., Tahrioui, A., Duchesne, R., Tortuel, D., et al. (2018). Extracytoplasmic function sigma factors in Pseudomonas aeruginosa. *Biochimica Et Biophysica Acta (BBA) - Gene Regulatory Mechanisms*. http://doi.org/10.1016/j.bbagrm.2018.04.008

Chipman, D. (2001). The ACT domain family. *Current Opinion in Structural Biology*, *11*(6), 694–700. http://doi.org/10.1016/S0959-440X(01)00272-X

Clark, R. R., Judd, J., Lasek-Nesselquist, E., Montgomery, S. A., Hoffmann, J. G., Derbyshire, K. M., & Gray, T. A. (2018). Direct cell-cell contact activates SigM to express the ESX-4 secretion system in Mycobacterium smegmatis. *Proceedings of the National Academy of Sciences of the United States of America*, *2*, 201804227. http://doi.org/10.1073/pnas.1804227115

Coulianos, N. N. G. (2018). Regulation of hemT expression in Rhodobacter sphaeroides wild type strain 2.4.9.

da Silva Neto, J. F., Koide, T., Gomes, S. L., & Marques, M. V. (2007). The single extracytoplasmic-function sigma factor of Xylella fastidiosa is involved in the heat shock response and presents an unusual regulatory mechanism. *Journal of Bacteriology*, *189*(2), 551–560. http://doi.org/10.1128/JB.00986-06

Dai, J., Wei, H., Tian, C., Damron, F. H., Zhou, J., & Qiu, D. (2015). An extracytoplasmic function sigma factor-dependent periplasmic glutathione peroxidase is involved in oxidative stress response of Shewanella oneidensis. *BMC Microbiology*, *15*(1), 34. http://doi.org/10.1186/s12866-015-0357-0

Danese, P. N., & Silhavy, T. J. (1998). CpxP, a stress-combative member of the Cpx regulon. *Journal of Bacteriology*, *180*(4), 831–839.

Delgado, C., Florez, L., Lollett, I., Lopez, C., Kangeyan, S., Kumari, H., et al. (2018). Pseudomonas aeruginosa Regulated Intramembrane Proteolysis (RIP): Protease MucP can Overcome Mutations in the AlgO Periplasmic Protease to Restore Alginate Production in Nonmucoid Revertants. *Journal of Bacteriology*, JB.00215–18. http://doi.org/10.1128/JB.00215-18

Demaegd, D., Colinet, A.-S., Deschamps, A., & Morsomme, P. (2014). Molecular Evolution of a Novel Family of Putative Calcium Transporters. *PloS One*, *9*(6), e100851. http://doi.org/10.1371/journal.pone.0100851

Dou, Y., Rutanhira, H., Chen, X., Mishra, A., Wang, C., & Fletcher, H. M. (2018). Role of extracytoplasmic function sigma factor PG1660 (RpoE) in the oxidative stress resistance regulatory network of Porphyromonas gingivalis. *Molecular Oral Microbiology*, *33*(1), 89–104. http://doi.org/10.1111/omi.12204

Duangurai, T., Indrawattana, N., & Pumirat, P. (2018). Burkholderia pseudomallei Adaptation for Survival in Stressful Conditions. *BioMed Research International*, *2018*(1), 1–11. http://doi.org/10.1155/2018/3039106

Duque, E., Rodríguez-Herva, J.-J., la Torre, de, J., Domínguez-Cuevas, P., Muñoz-Rojas, J., & Ramos, J.-L. (2007). The RpoT regulon of Pseudomonas putida DOT-T1E and its role in stress endurance against solvents. *Journal of Bacteriology*, *189*(1), 207–219. http://doi.org/10.1128/JB.00950-06

Dutta, R., & Inouye, M. (2000). GHKL, an emergent ATPase/kinase superfamily. *Trends in Biochemical Sciences*, *25*(1), 24–28.

Egler, M., Grosse, C., Grass, G., & Nies, D. H. (2005). Role of the extracytoplasmic function protein family sigma factor RpoE in metal resistance of Escherichia coli. *Journal of Bacteriology*, *187*(7), 2297–2307. http://doi.org/10.1128/JB.187.7.2297-2307.2005

Espinasse, S., Gohar, M., Lereclus, D., & Sanchis, V. (2004). An extracytoplasmic-function sigma factor is involved in a pathway controlling beta-exotoxin I production in Bacillus thuringiensis subsp. thuringiensis strain 407-1. *Journal of Bacteriology*, *186*(10), 3108–3116. http://doi.org/10.1128/JB.186.10.3108-3116.2004

Finn, R. D. (2004). Pfam: the protein families database. In *Encyclopedia of Genetics, Genomics, Proteomics and Bioinformatics* (Vol. 215, p. 403). Chichester, UK: John Wiley & Sons, Ltd. http://doi.org/10.1002/047001153X.g306303

Foulston, L., & Bibb, M. (2011). Feed-forward regulation of microbisporicin biosynthesis in Microbispora corallina. *Journal of Bacteriology*, *193*(12), 3064–3071. http://doi.org/10.1128/JB.00250-11

Francez-Charlot, A., Frunzke, J., Reichen, C., Ebneter, J. Z., Gourion, B., & Vorholt, J. A. (2009). Sigma factor mimicry involved in regulation of general stress response. *Proceedings of the National Academy of Sciences of the United States of America*, *106*(9), 3467–3472. http://doi.org/10.1073/pnas.0810291106

Gaudion, A., Dawson, L., Davis, E., & Smollett, K. (2013). Characterisation of the Mycobacterium tuberculosis alternative sigma factor SigG: its operon and regulon. *Tuberculosis (Edinburgh, Scotland)*, *93*(5), 482–491. http://doi.org/10.1016/j.tube.2013.05.005

Gordon, N. D., Ottaviano, G. L., Connell, S. E., Tobkin, G. V., Son, C. H., Shterental, S., & Gehring, A. M. (2008). Secreted-protein response to sigmaU activity in Streptomyces coelicolor. *Journal of Bacteriology*, *190*(3), 894–904. http://doi.org/10.1128/JB.01759-07

Gourion, B., Sulser, S., Frunzke, J., Francez-Charlot, A., Stiefel, P., Pessi, G., et al. (2009). The PhyR-σEcfG signalling cascade is involved in stress response and symbiotic efficiency in Bradyrhizobium japonicum. *Molecular Microbiology*, *73*(2), 291–305. http://doi.org/10.1111/j.1365-2958.2009.06769.x

Goutam, K., Gupta, A. K., & Gopal, B. (2017). The fused SnoaL_2 domain in the Mycobacterium tuberculosis sigma factor σJ modulates promoter recognition. *Nucleic Acids Research*, *45*(16), 9760–9772. http://doi.org/10.1093/nar/gkx609

Grass, G., Fricke, B., & Nies, D. H. (2005). Control of expression of a periplasmic nickel efflux pump by periplasmic nickel concentrations. *Biometals : an International Journal on the Role of Metal Ions in Biology, Biochemistry, and Medicine*, *18*(4), 437–448. http://doi.org/10.1007/s10534-005-3718-6

Greenwell, R., Nam, T.-W., & Donohue, T. J. (2011). Features of Rhodobacter sphaeroides ChrR Required for Stimuli to Promote the Dissociation of σE/ChrR Complexes. *Journal of Molecular Biology*, *407*(4), 477–491. http://doi.org/10.1016/j.jmb.2011.01.055

Grosse, C., Poehlein, A., Blank, K., Schwarzenberger, C., Schleuder, G., Herzberg, M., & Nies, D. H. (2019). The third pillar of metal homeostasis in Cupriavidus metalliduransCH34: Preferences are controlled by extracytoplasmic function sigma factors. *Metallomics : Integrated Biometal Science*. http://doi.org/10.1039/C8MT00299A

Gunesekere, I. C., Kahler, C. M., Ryan, C. S., Snyder, L. A. S., Saunders, N. J., Rood, J. I., & Davies, J. K. (2006). Ecf, an alternative sigma factor from Neisseria gonorrhoeae, controls expression of msrAB, which encodes methionine sulfoxide reductase. *Journal of Bacteriology*, *188*(10), 3463–3469. http://doi.org/10.1128/JB.188.10.3463-3469.2006

Hahn, M. Y., Raman, S., Anaya, M., & Husson, R. N. (2005). The Mycobacterium tuberculosis extracytoplasmic-function sigma factor SigL regulates polyketide synthases and secreted or membrane proteins and is required for virulence. *Journal of Bacteriology*, *187*(20), 7062–7071. http://doi.org/10.1128/JB.187.20.7062-7071.2005

Haines-Menges, B., Whitaker, W. B., & Boyd, E. F. (2014). Alternative sigma factor RpoE is important for Vibrio parahaemolyticus cell envelope stress response and intestinal colonization. *Infection and Immunity*, *82*(9), 3667–3677. http://doi.org/10.1128/IAI.01854-14

Hasselblatt, H., Kurzbauer, R., Wilken, C., Krojer, T., Sawa, J., Kurt, J., et al. (2007). Regulation of the sigmaE stress response by DegS: how the PDZ domain keeps the protease inactive in the resting state and allows integration of different OMP-derived stress signals upon folding stress. *Genes & Development*, *21*(20), 2659–2670. http://doi.org/10.1101/gad.445307

Helmann, J. D. (2016). ScienceDirect Bacillus subtilis extracytoplasmic function (ECF) sigma factors and defense of the cell envelope. *Current Opinion in Microbiology*, *30*, 122–132. http://doi.org/10.1016/j.mib.2016.02.002

Hiratsu, K., Amemura, M., Nashimoto, H., Shinagawa, H., & Makino, K. (1995). The rpoE gene of Escherichia coli, which encodes sigma E, is essential for bacterial growth at high temperature. *Journal of Bacteriology*, *177*(10), 2918–2922. http://doi.org/10.1128/jb.177.10.2918-2922.1995

Ho, T. D., & Ellermeier, C. D. (2011). PrsW is required for colonization, resistance to antimicrobial peptides, and expression of extracytoplasmic function σ factors in Clostridium difficile. *Infection and Immunity*, *79*(8), 3229–3238. http://doi.org/10.1128/IAI.00019-11

Howlett, R., Read, N., Varghese, A., Kershaw, C., Hancock, Y., & Smith, M. C. M. (2018). Streptomyces coelicolor strains lacking polyprenol phosphate mannose synthase and protein O-mannosyl transferase are hyper-susceptible to multiple antibiotics. *Microbiology*, *164*(3), 369–382. http://doi.org/10.1099/mic.0.000605

Hu, Y., Kendall, S., Stoker, N. G., & Coates, A. R. M. (2004). The Mycobacterium tuberculosis sigJ gene controls sensitivity of the bacterium to hydrogen peroxide. *FEMS Microbiology Letters*, *237*(2), 415–423. http://doi.org/10.1111/j.1574-6968.2004.tb09725.x

Huang, X., Pinto, D., Fritz, G., & Mascher, T. (2015). Environmental sensing in Actinobacteria: a comprehensive survey on the signaling capacity of this phylum. *Journal of Bacteriology*, *197*(15), 2517–2535. http://doi.org/10.1128/JB.00176-15

Huis in 't Veld, R. A. G., Willemsen, A. M., van Kampen, A. H. C., Bradley, E. J., Baas, F., Pannekoek, Y., & van der Ende, A. (2011). Deep sequencing whole transcriptome exploration of the σE regulon in Neisseria meningitidis. *PloS One*, *6*(12), e29002. http://doi.org/10.1371/journal.pone.0029002

Inaba, K., Suzuki, M., Maegawa, K.-I., Akiyama, S., Ito, K., & Akiyama, Y. (2008). A pair of circularly permutated PDZ domains control RseP, the S2P family intramembrane protease of Escherichia coli. *Journal of Biological Chemistry*, *283*(50), 35042–35052. http://doi.org/10.1074/jbc.M806603200

Jaiswal, R. K., Prabha, T. S., Manjeera, G., & Gopal, B. (2013). Mycobacterium tuberculosis RsdA provides a conformational rationale for selective regulation of σ-factor activity by proteolysis. *Nucleic Acids Research*, *41*(5), 3414–3423. http://doi.org/10.1093/nar/gks1468

Jogler, C., Waldmann, J., Huang, X., Jogler, M., Glockner, F. O., Mascher, T., & Kolter, R. (2012). Identification of Proteins Likely To Be Involved in Morphogenesis, Cell Division, and Signal Transduction in Planctomycetes by Comparative Genomics. *Journal of Bacteriology*, *194*(23), 6419–6430. http://doi.org/10.1128/JB.01325-12

Kadowaki, T., Yukitake, H., Naito, M., Sato, K., Kikuchi, Y., Kondo, Y., et al. (2016). A two-component system regulates gene expression of the type IX secretion component proteins via an ECF sigma factor. *Scientific Reports*, *6*(1), 23288. http://doi.org/10.1038/srep23288

Kappler, U., Friedrich, C. G., Trüper, H. G., & Dahl, C. (2001). Evidence for two pathways of thiosulfate oxidation in<Emphasis Type="Italic"> Starkeya novella</Emphasis> (formerly<Emphasis Type="Italic"> Thiobacillus novellus</Emphasis>). *Archives of Microbiology*, *175*(2), 102–111. http://doi.org/10.1007/s002030000241

Kim, M.-S., Hahn, M. Y., Cho, Y., Cho, S.-N., & Roe, J. H. (2009). Positive and negative feedback regulatory loops of thiol-oxidative stress response mediated by an unstable isoform of sigmaR in actinomycetes. *Molecular Microbiology*, *73*(5), 815–825. http://doi.org/10.1111/j.1365-2958.2009.06824.x

Kocher, O., Comella, N., Tognazzi, K., & Brown, L. F. (1998). Identification and partial characterization of PDZK1: a novel protein containing PDZ interaction domains. *Laboratory Investigation; a Journal of Technical Methods and Pathology*, *78*(1), 117–125.

Kohler, C., Lourenço, R. F., Avelar, G. M., & Gomes, S. L. (2012). Extracytoplasmic function (ECF) sigma factor σF is involved in Caulobacter crescentus response to heavy metal stress. *BMC Microbiology*, *12*(1), 210. http://doi.org/10.1186/1471-2180-12-210

Kumar, S., Badireddy, S., Pal, K., Sharma, S., Arora, C., Garg, S. K., et al. (2012). Interaction of Mycobacterium tuberculosis RshA and SigH is mediated by salt bridges. *PloS One*, *7*(8), e43676. http://doi.org/10.1371/journal.pone.0043676

Kwak, M.-K., Ryu, H.-B., Song, S.-H., Lee, J.-W., & Kang, S.-O. (2018). Anti-sigma factor YlaD regulates transcriptional activity of sigma factor YlaC and sporulation via manganese-dependent redox-sensing molecular switch in Bacillus subtilis. *The Biochemical Journal*, BCJ20170911. http://doi.org/10.1042/BCJ20170911

Lane, W. J., & Darst, S. A. (2010). Molecular evolution of multisubunit RNA polymerases: structural analysis. *Journal of Molecular Biology*, *395*(4), 686–704. http://doi.org/10.1016/j.jmb.2009.10.063

Lang, C., Barnett, M. J., Fisher, R. F., Smith, L. S., Diodati, M. E., & Long, S. R. (2018). Most Sinorhizobium meliloti Extracytoplasmic Function Sigma Factors Control Accessory Functions. *mSphere*, *3*(5), 619. http://doi.org/10.1128/mSphereDirect.00454-18

Lee, Jang-Woo, & Lee, E.-J. (2015). Regulation and function of the <Emphasis Type="Italic">Salmonella</Emphasis> MgtC virulence protein. *The Journal of Microbiology*, *53*(10), 667–672. http://doi.org/10.1007/s12275-015-5283-1

Lee, Jong-Hee, Ammerman, N. C., Nolan, S., Geiman, D. E., Lun, S., Guo, H., & Bishai, W. R. (2012). Isoniazid resistance without a loss of fitness in Mycobacterium tuberculosis. *Nature Communications*, *3*(1), e1000160. http://doi.org/10.1038/ncomms1724

Lee, Jong-Hee, Geiman, D. E., & Bishai, W. R. (2008). Role of stress response sigma factor SigG in Mycobacterium tuberculosis. *Journal of Bacteriology*, *190*(3), 1128–1133. http://doi.org/10.1128/JB.00511-07

Lewerke, L. T., Kies, P. J., Müh, U., & Ellermeier, C. D. (2018). Bacterial sensing: A putative amphipathic helix in RsiV is the switch for activating σV in response to lysozyme. *PLOS Genetics*, *14*(7), e1007527. http://doi.org/10.1371/journal.pgen.1007527

Li, Q., Peng, T., & Klug, G. (2018). The PhyR homolog RSP_1274 of Rhodobacter sphaeroides is involved in defense of membrane stress and has a moderate effect on RpoE (RSP_1092) activity. *BMC Microbiology*, *18*(1), 18. http://doi.org/10.1186/s12866-018-1161-4

Ling, J. M. L., Moore, R. A., Surette, M. G., & Woods, D. E. (2006). The mviN homolog in Burkholderia pseudomallei is essential for viability and virulence. *Canadian Journal of Microbiology*, *52*(9), 831–842. http://doi.org/10.1139/w06-042

Liu, L., Feng, D., Chen, G., Chen, M., Zheng, Q., Song, P., et al. (2012). Mitochondrial outer-membrane protein FUNDC1 mediates hypoxia-induced mitophagy in mammalian cells. *Nature Cell Biology*, *14*(2), 177–185. http://doi.org/10.1038/ncb2422

Liu, Q., Pinto, D., & Mascher, T. (2018). Characterization of the widely distributed novel ECF42 group of extracytoplasmic function σ factors in Streptomyces venezuelae. *Journal of Bacteriology*, JB.00437–18. http://doi.org/10.1128/JB.00437-18

Llamas, M. A., Imperi, F., Visca, P., & Lamont, I. L. (2014). Cell-surface signaling in Pseudomonas: stress responses, iron transport, and pathogenicity. *FEMS Microbiology Reviews*, *38*(4), 569–597. http://doi.org/10.1111/1574-6976.12078

López-García, M. T., Yagüe, P., Gonzalez-Quiñonez, N., Rioseras, B., & Manteca, A. (2018). The SCO4117 ECF sigma factor pleiotropically controls secondary metabolism and morphogenesis in Streptomyces coelicolor. *Frontiers in Microbiology*, *9*. http://doi.org/10.3389/fmicb.2018.00312

Luo, S., Sun, D., Zhu, J., Chen, Z., Wen, Y., & Li, J. (2014). An extracytoplasmic function sigma factor, σ(25), differentially regulates avermectin and oligomycin biosynthesis in Streptomyces avermitilis. *Applied Microbiology and Biotechnology*, *98*(16), 7097–7112. http://doi.org/10.1007/s00253-014-5759-7

Luo, Y., Asai, K., Sadaie, Y., & Helmann, J. D. (2010). Transcriptomic and phenotypic characterization of a Bacillus subtilis strain without extracytoplasmic function σ factors. *Journal of Bacteriology*, *192*(21), 5736–5745. http://doi.org/10.1128/JB.00826-10

Maillard, A. P., Girard, E., Ziani, W., Petit-Härtlein, I., Kahn, R., & Covès, J. (2014). The Crystal Structure of the Anti-σ Factor CnrY in Complex with the σ Factor CnrH Shows a New Structural Class of Anti-σ Factors Targeting Extracytoplasmic Function σ Factors. *Journal of Molecular Biology*, *426*(12), 2313–2327. http://doi.org/10.1016/j.jmb.2014.04.003

Malinverni, J. C., & Silhavy, T. J. (2009). An ABC transport system that maintains lipid asymmetry in the gram-negative outer membrane. *Proceedings of the National Academy of Sciences of the United States of America*, *106*(19), 8009–8014. http://doi.org/10.1073/pnas.0903229106

Mao, X.-M., Zhou, Z., Cheng, L.-Y., Hou, X.-P., Guan, W.-J., & Li, Y.-Q. (2009). Involvement of SigT and RstA in the differentiation of Streptomyces coelicolor. *FEBS Letters*, *583*(19), 3145–3150. http://doi.org/10.1016/j.febslet.2009.09.025

Marcotte, E. M., Pellegrini, M., Ng, H.-L., Rice, D. W., Yeates, T. O., & Eisenberg, D. (1999). Detecting Protein Function and Protein-Protein Interactions from Genome Sequences. *Science (New York, N.Y.)*, *285*(5428), 751–753. http://doi.org/10.1126/science.285.5428.751

Marshall, B., Stintzi, A., Gilmour, C., Meyer, J. M., & Poole, K. (2009). Citrate-mediated iron uptake in Pseudomonas aeruginosa: involvement of the citrate-inducible FecA receptor and the FeoB ferrous iron transporter. *Microbiology*, *155*(1), 305–315. http://doi.org/10.1099/mic.0.023531-0

Martinez-Salazar, J. M., Salazar, E., Encarnacion, S., Ramirez-Romero, M. A., & Rivera, J. (2009). Role of the Extracytoplasmic Function Sigma Factor RpoE4 in Oxidative and Osmotic Stress Responses in Rhizobium etli. *Journal of Bacteriology*, *191*(13), 4122–4132. http://doi.org/10.1128/JB.01626-08

Mascher, T., Hachmann, A.-B., & Helmann, J. D. (2007). Regulatory overlap and functional redundancy among Bacillus subtilis extracytoplasmic function sigma factors. *Journal of Bacteriology*, *189*(19), 6919–6927. http://doi.org/10.1128/JB.00904-07

Masloboeva, N., Reutimann, L., Stiefel, P., Follador, R., Leimer, N., Hennecke, H., et al. (2012). Reactive oxygen species-inducible ECF σ factors of Bradyrhizobium japonicum. *PloS One*, *7*(8), e43421. http://doi.org/10.1371/journal.pone.0043421

Medina, E., Ryan, L., LaCourse, R., & North, R. J. (2006). Superior virulence of Mycobacterium bovis over Mycobacterium tuberculosis (Mtb) for Mtb-resistant and Mtb-susceptible mice is manifest as an ability to cause extrapulmonary disease. *Tuberculosis*, *86*(1), 20–27. http://doi.org/10.1016/j.tube.2005.04.003

Melville, S., & Craig, L. (2013). Type IV Pili in Gram-Positive Bacteria. *Microbiology and Molecular Biology Reviews*, *77*(3), 323–341. http://doi.org/10.1128/MMBR.00063-12

Mendez, R., Gutierrez, A., Reyes, J., & Márquez-Magaña, L. (2012). The extracytoplasmic function sigma factor SigY is important for efficient maintenance of the Spβ prophage that encodes sublancin in Bacillus subtilis. *DNA and Cell Biology*, *31*(6), 946–955. http://doi.org/10.1089/dna.2011.1513

Missiakas, D., Mayer, M. P., Lemaire, M., Georgopoulos, C., & Raina, S. (1997). Modulation of the Escherichia coli sigmaE (RpoE) heat-shock transcription-factor activity by the RseA, RseB and RseC proteins. *Molecular Microbiology*, *24*(2), 355–371.

Moody, R. G., & Williamson, M. P. (2013). Structure and function of a bacterial Fasciclin I Domain Protein elucidates function of related cell adhesion proteins such as TGFBIp and periostin. *FEBS Open Bio*, *3*(1), 71–77. http://doi.org/10.1016/j.fob.2013.01.001

Morita, Y. S., Sena, C. B. C., Waller, R. F., Kurokawa, K., Sernee, M. F., Nakatani, F., et al. (2006). PimE is a polyprenol-phosphate-mannose-dependent mannosyltransferase that transfers the fifth mannose of phosphatidylinositol mannoside in mycobacteria. *Journal of Biological Chemistry*, *281*(35), 25143–25155. http://doi.org/10.1074/jbc.M604214200

Müller, M., Reiß, S., Schlüter, R., Mäder, U., Beyer, A., Reiß, W., et al. (2014). Deletion of membrane-associated Asp23 leads to upregulation of cell wall stress genes in Staphylococcus aureus. *Molecular Microbiology*, *93*(6), 1259–1268. http://doi.org/10.1111/mmi.12733

Nizan-Koren, R., Manulis, S., Mor, H., Iraki, N. M., & Barash, I. (2003). The regulatory cascade that activates the Hrp regulon in Erwinia herbicola pv. gypsophilae. *Molecular Plant-Microbe Interactions : MPMI*, *16*(3), 249–260. http://doi.org/10.1094/MPMI.2003.16.3.249

Nunvar, J., Buroni, S., Cystic, V. M. J. O., 2018. (n.d.). … The effect of 2-thiopyridine derivative 11026103, a novel antibacterial compound, on Burkholderia cenocepacia: transcriptomic response and resistance mechanisms. *Cysticfibrosisjournal.com*.

Ogura, M., & Asai, K. (2016). Glucose Induces ECF Sigma Factor Genes, sigX and sigM, Independent of Cognate Anti-sigma Factors through Acetylation of CshA in Bacillus subtilis. *Frontiers in Microbiology*, *7*(e18179), 4478. http://doi.org/10.3389/fmicb.2016.01918

Okrent, R. A., Halgren, A. B., Azevedo, M. D., Chang, J. H., Mills, D. I., Maselko, M., et al. (2014). Negative regulation of germination-arrest factor production in Pseudomonas fluorescens WH6 by a putative extracytoplasmic function sigma factor. *Microbiology*, *160*(Pt_11), 2432–2442. http://doi.org/10.1099/mic.0.080317-0

Omardien, S., Drijfhout, J. W., van Veen, H., Schachtschabel, S., Riool, M., Hamoen, L. W., et al. (2018). Synthetic antimicrobial peptides delocalize membrane bound proteins thereby inducing a cell envelope stress response. *Biochimica Et Biophysica Acta (BBA) - Biomembranes*. http://doi.org/10.1016/j.bbamem.2018.06.005

Orino, K., Lehman, L., Tsuji, Y., Ayaki, H., Torti, S. V., & Torti, F. M. (2001). Ferritin and the response to oxidative stress. *The Biochemical Journal*, *357*(Pt 1), 241–247.

Park, S. T., Kang, C.-M., & Husson, R. N. (2008). Regulation of the SigH stress response regulon by an essential protein kinase in Mycobacterium tuberculosis. *Proceedings of the National Academy of Sciences of the United States of America*, *105*(35), 13105–13110. http://doi.org/10.1073/pnas.0801143105

Pátek, M., Dostálová, H., Busche, T., Holátko, J., Rucká, L., Štěpánek, V., et al. (2018). Overlap of promoter recognition specificity of stress response sigma factors SigD and SigH in Corynebacterium glutamicum ATCC 13032. *Frontiers in Microbiology*, *9*. http://doi.org/10.3389/fmicb.2018.03287

Pettersson, B. M. F., Das, S., Behra, P. R. K., Jordan, H. R., Ramesh, M., Mallick, A., et al. (2015). Comparative Sigma Factor-mRNA Levels in Mycobacterium marinum under Stress Conditions and during Host Infection. *PloS One*, *10*(10), e0139823. http://doi.org/10.1371/journal.pone.0139823

Pinto, D., & Mascher, T. (2016). The ECF Classification: A Phylogenetic Reflection of the Regulatory Diversity in the Extracytoplasmic Function σ Factor Protein Family. In *Stress and Environmental Regulation of Gene Expression and Adaptation in Bacteria* (Vol. 7, pp. 64–96). Hoboken, NJ, USA: John Wiley & Sons, Inc. http://doi.org/10.1002/9781119004813.ch7

Rahman, M. T., Parreira, V., & Prescott, J. F. (2005). In vitro and intra-macrophage gene expression by Rhodococcus equi strain 103. *Veterinary Microbiology*, *110*(1-2), 131–140. http://doi.org/10.1016/j.vetmic.2005.08.005

Rakotoarivonina, H., Hebraud, M., Mosoni, P., Forano, E., Jubelin, G., & Gaillard-Martinie, B. (2002). Adhesion to cellulose of the Gram-positive bacterium Ruminococcus albus involves type IV pili. *Microbiology*, *148*(6), 1871–1880. http://doi.org/10.1099/00221287-148-6-1871

Raman, S., Hazra, R., Dascher, C. C., & Husson, R. N. (2004). Transcription regulation by the Mycobacterium tuberculosis alternative sigma factor SigD and its role in virulence. *Journal of Bacteriology*, *186*(19), 6605–6616. http://doi.org/10.1128/JB.186.19.6605-6616.2004

Rhodius, V. A., & Mutalik, V. K. (2010). Predicting strength and function for promoters of the Escherichia coli alternative sigma factor, sigmaE. *Proceedings of the National Academy of Sciences of the United States of America*, *107*(7), 2854–2859. http://doi.org/10.1073/pnas.0915066107

Rhodius, V. A., Segall-Shapiro, T. H., Sharon, B. D., Ghodasara, A., Orlova, E., Tabakh, H., et al. (2013). Design of orthogonal genetic switches based on a crosstalk map of σs, anti-σs, and promoters. *Molecular Systems Biology*, *9*(1), 702–702. http://doi.org/10.1038/msb.2013.58

Rocha, E. R., & Smith, C. J. (2013). Ferritin-like family proteins in the anaerobe Bacteroides fragilis: when an oxygen storm is coming, take your iron to the shelter. *Biometals : an International Journal on the Role of Metal Ions in Biology, Biochemistry, and Medicine*, *26*(4), 577–591. http://doi.org/10.1007/s10534-013-9650-2

Rowley, G., Spector, M., Kormanec, J., & Roberts, M. (2006). Pushing the envelope: extracytoplasmic stress responses in bacterial pathogens. *Nature Reviews Microbiology*, *4*(5), 383–394. http://doi.org/10.1038/nrmicro1394

Sachdeva, P., Misra, R., Tyagi, A. K., & Singh, Y. (2009). The sigma factors of Mycobacterium tuberculosis: regulation of the regulators. *FEBS Journal*, *277*(3), 605–626. http://doi.org/10.1111/j.1742-4658.2009.07479.x

Saha, P., Manna, C., Das, S., & Ghosh, M. (2016). Antibiotic binding of STY3178, a yfdX protein from <i>Salmonella</i> Typhi. *Scientific Reports*, *6*(1), 21305. http://doi.org/10.1038/srep21305

Saïd-Salim, B., Mostowy, S., Kristof, A. S., & Behr, M. A. (2006). Mutations in Mycobacterium tuberculosis Rv0444c, the gene encoding anti-SigK, explain high level expression of MPB70 and MPB83 in Mycobacterium bovis. *Molecular Microbiology*, *62*(5), 1251–1263. http://doi.org/10.1111/j.1365-2958.2006.05455.x

Sauviac, L., Philippe, H., Phok, K., & Bruand, C. (2007). An extracytoplasmic function sigma factor acts as a general stress response regulator in Sinorhizobium meliloti. *Journal of Bacteriology*, *189*(11), 4204–4216. http://doi.org/10.1128/JB.00175-07

Schnappinger, D., Ehrt, S., Voskuil, M. I., Liu, Y., Mangan, J. A., Monahan, I. M., et al. (2003). Transcriptional Adaptation of Mycobacterium tuberculosis within Macrophages: Insights into the Phagosomal Environment. *Journal of Experimental Medicine*, *198*(5), 693–704. http://doi.org/10.1084/jem.20030846

Schneider, J. S., Sklar, J. G., & Glickman, M. S. (2014). The Rip1 Protease of Mycobacterium tuberculosis Controls the SigD Regulon. *Journal of Bacteriology*, *196*(14), 2638–2645. http://doi.org/10.1128/JB.01537-14

Schrallhammer, M., Galati, S., Altenbuchner, J., Schweikert, M., Görtz, H.-D., & Petroni, G. (2012). Tracing the role of R-bodies in the killer trait: absence of toxicity of R-body producing recombinant E. coli on paramecia. *European Journal of Protistology*, *48*(4), 290–296. http://doi.org/10.1016/j.ejop.2012.01.008

Schumacher, M. A., Bush, M. J., Bibb, M. J., Ramos-León, F., Chandra, G., Zeng, W., & Buttner, M. J. (2018). The crystal structure of the RsbN-σBldN complex from Streptomyces venezuelae defines a new structural class of anti-σ factor. *Nucleic Acids Research*, *68*, 357. http://doi.org/10.1093/nar/gky493

Seipke, R. F., Patrick, E., & Hutchings, M. I. (2014). Regulation of antimycin biosynthesis by the orphan ECF RNA polymerase sigma factor σAntA. http://doi.org/10.7287/peerj.preprints.203v1

Sharp, J. D., Singh, A. K., Park, S. T., Lyubetskaya, A., Peterson, M. W., Gomes, A. L. C., et al. (2016). Comprehensive Definition of the SigH Regulon of Mycobacterium tuberculosis Reveals Transcriptional Control of Diverse Stress Responses. *PloS One*, *11*(3), e0152145. http://doi.org/10.1371/journal.pone.0152145

Shearwin, K. E., Callen, B. P., & Egan, J. B. (2005). Transcriptional interference--a crash course. *Trends in Genetics : TIG*, *21*(6), 339–345. http://doi.org/10.1016/j.tig.2005.04.009

Sherwood, E. J., & Bibb, M. J. (2013). The antibiotic planosporicin coordinates its own production in the actinomycete Planomonospora alba. *Proceedings of the National Academy of Sciences of the United States of America*, *110*(27), E2500–9. http://doi.org/10.1073/pnas.1305392110

Shu, D., Chen, L., Wang, W., Yu, Z., Ren, C., Zhang, W., et al. (2008). <Emphasis Type="Italic">afsQ1</Emphasis>-<Emphasis Type="Italic">Q2</Emphasis>-<Emphasis Type="Italic">sigQ</Emphasis> is a pleiotropic but conditionally required signal transduction system for both secondary metabolism and morphological development in <Emphasis Type="Italic">Streptomyces coelicolor</Emphasis>. *Applied Microbiology and Biotechnology*, *81*(6), 1149–1160. http://doi.org/10.1007/s00253-008-1738-1

Shukla, J., Gupta, R., Thakur, K. G., Gokhale, R., Gopal, B., IUCr. (2014). Structural basis for the redox sensitivity of the Mycobacterium tuberculosis SigK–RskA σ–anti-σ complex. *Acta Crystallographica Section D: Biological Crystallography*, *70*(4), 1026–1036. http://doi.org/10.1107/S1399004714000121

Shultis, D. D., Purdy, M. D., Banchs, C. N., & Wiener, M. C. (2006). Outer Membrane Active Transport: Structure of the BtuB:TonB Complex, *312*(5778), 1396–1399. http://doi.org/10.1126/science.1127694

Sklar, J. G., Makinoshima, H., Schneider, J. S., & Glickman, M. S. (2010). M. tuberculosis intramembrane protease Rip1 controls transcription through three anti-sigma factor substrates. *Molecular Microbiology*, *77*(3), 605–617. http://doi.org/10.1111/j.1365-2958.2010.07232.x

Smith, D. R., Price, J. E., Burby, P. E., Blanco, L. P., Chamberlain, J., & Chapman, M. R. (2017). The Production of Curli Amyloid Fibers Is Deeply Integrated into the Biology of Escherichia coli. *Biomolecules*, *7*(4), 75. http://doi.org/10.3390/biom7040075

Sohn, J., Grant, R. A., & Sauer, R. T. (2007). Allosteric activation of DegS, a stress sensor PDZ protease. *Cell*, *131*(3), 572–583. http://doi.org/10.1016/j.cell.2007.08.044

Souza, B. M., Castro, T. L. de P., Carvalho, R. D. de O., Seyffert, N., Silva, A., Miyoshi, A., & Azevedo, V. (2014). σ ECFfactors of gram-positive bacteria. *Virulence*, *5*(5), 587–600. http://doi.org/10.4161/viru.29514

Spöring, I., Felgner, S., Preuße, M., Eckweiler, D., Rohde, M., Häussler, S., et al. (2018). Regulation of Flagellum Biosynthesis in Response to Cell Envelope Stress in Salmonella enterica Serovar Typhimurium. *mBio*, *9*(3), e00736–17. http://doi.org/10.1128/mBio.00736-17

Staroń, A., & Mascher, T. (2010). General stress response in α-proteobacteria: PhyR and beyond. *Molecular Microbiology*, *78*(2), 271–277.

Staroń, A., Sofia, H. J., Dietrich, S., Ulrich, L. E., Liesegang, H., & Mascher, T. (2009). The third pillar of bacterial signal transduction: classification of the extracytoplasmic function (ECF) σ factor protein family. *Molecular Microbiology*, *74*(3), 557–581. http://doi.org/10.1111/j.1365-2958.2009.06870.x

Sultana, A., Kallio, P., Jansson, A., Wang, J.-S., Niemi, J., Mäntsälä, P., & Schneider, G. (2004). Structure of the polyketide cyclase SnoaL reveals a novel mechanism for enzymatic aldol condensation. *The EMBO Journal*, *23*(9), 1911–1921. http://doi.org/10.1038/sj.emboj.7600201

Sumi, S., Shiratori-Takano, H., Ueda, K., & Takano, H. (2018). Role and Function of Class III LitR, a Photosensor Homolog from Burkholderia multivorans. *Journal of Bacteriology*, JB.00285–18. http://doi.org/10.1128/JB.00285-18

Sun, R., Converse, P. J., Ko, C., Tyagi, S., Morrison, N. E., & Bishai, W. R. (2004). Mycobacterium tuberculosis ECF sigma factor sigC is required for lethality in mice and for the conditional expression of a defined gene set. *Molecular Microbiology*, *52*(1), 25–38. http://doi.org/10.1111/j.1365-2958.2003.03958.x

Takano, H., Obitsu, S., Beppu, T., & Ueda, K. (2005). Light-induced carotenogenesis in Streptomyces coelicolor A3(2): identification of an extracytoplasmic function sigma factor that directs photodependent transcription of the carotenoid biosynthesis gene cluster. *Journal of Bacteriology*, *187*(5), 1825–1832. http://doi.org/10.1128/JB.187.5.1825-1832.2005

Tart, A. H., Wolfgang, M. C., & Wozniak, D. J. (2005). The alternative sigma factor AlgT represses Pseudomonas aeruginosa flagellum biosynthesis by inhibiting expression of fleQ. *Journal of Bacteriology*, *187*(23), 7955–7962. http://doi.org/10.1128/JB.187.23.7955-7962.2005

Tettmann, B., Dötsch, A., Armant, O., Fjell, C. D., & Overhage, J. (2014). Knockout of Extracytoplasmic Function Sigma Factor ECF-10 Affects Stress Resistance and Biofilm Formation in Pseudomonas putida KT2440. *Applied and Environmental Microbiology*, *80*(16), 4911–4919. http://doi.org/10.1128/AEM.01291-14

Thakur, K. G., Praveena, T., & Gopal, B. (2010). Structural and biochemical bases for the redox sensitivity of Mycobacterium tuberculosis RslA. *Journal of Molecular Biology*, *397*(5), 1199–1208. http://doi.org/10.1016/j.jmb.2010.02.026

Tojo, S., Matsunaga, M., Matsumoto, T., Kang, C.-M., Yamaguchi, H., Asai, K., et al. (2003). Organization and expression of the Bacillus subtilis sigY operon. *Journal of Biochemistry*, *134*(6), 935–946.

Torres, J. P., Muñoz-Dorado, J., & Moraleda-Muñoz, A. (2018). The complex global response to copper in the multicellular bacterium Myxococcus xanthus. *Metallomics : Integrated Biometal Science*. http://doi.org/10.1039/C8MT00121A

Toyoda, K., & Inui, M. (2016). The extracytoplasmic function σ factor σC regulates expression of a branched quinol oxidation pathway in Corynebacterium glutamicum. *Molecular Microbiology*, *100*(3), 486–509. http://doi.org/10.1111/mmi.13330

Toyoda, K., & Inui, M. (2017). Extracytoplasmic function sigma factor σD confers resistance to environmental stress by enhancing mycolate synthesis and modifying peptidoglycan structures in Corynebacterium glutamicum. *Molecular Microbiology*, *107*(3), 312–329. http://doi.org/10.1111/mmi.13883

Tran, N. T., Huang, X., Hong, H.-J., Bush, M. J., Chandra, G., Pinto, D., et al. (2019). Defining the regulon of genes controlled by σ E, a key regulator of the cell envelope stress response in Streptomyces coelicolor. *Molecular Microbiology*, mmi.14250. http://doi.org/10.1111/mmi.14250

Trepreau, J., Girard, E., Maillard, A. P., de Rosny, E., Petit-Haertlein, I., Kahn, R., & Covès, J. (2011). Structural basis for metal sensing by CnrX. *Journal of Molecular Biology*, *408*(4), 766–779. http://doi.org/10.1016/j.jmb.2011.03.014

Trovero, M. F., Scavone, P., Platero, R., Souza, E. M., Fabiano, E., & Rosconi, F. (2018). Herbaspirillum seropedicae differentially expressed genes in response to iron availability. *Frontiers in Microbiology*, *9*. http://doi.org/10.3389/fmicb.2018.01430

Tsirigos, K. D., Peters, C., Shu, N., Käll, L., & Elofsson, A. (2015). The TOPCONS web server for consensus prediction of membrane protein topology and signal peptides. *Nucleic Acids Research*, *43*(W1), W401–W407. http://doi.org/10.1093/nar/gkv485

Tsukazaki, T., Mori, H., Echizen, Y., Ishitani, R., Fukai, S., Tanaka, T., et al. (2011). Structure and function of a membrane component SecDF that enhances protein export. *Nature*, *474*(7350), 235–238. http://doi.org/10.1038/nature09980

Vanderpool, C. K., & Armstrong, S. K. (2003). Heme-Responsive Transcriptional Activation of Bordetella bhu Genes. *Journal of Bacteriology*, *185*(3), 909–917. http://doi.org/10.1128/JB.185.3.909-917.2003

Veith, P. D., Luong, C., Tan, K. H., Dashper, S. G., & Reynolds, E. C. (2018). The Outer Membrane Vesicle Proteome of Porphyromonas gingivalis is Differentially Modulated Relative to the Outer Membrane in Response to Heme Availability. *Journal of Proteome Research*, acs.jproteome.8b00153. http://doi.org/10.1021/acs.jproteome.8b00153

Veyrier, F., Saïd-Salim, B., & Behr, M. A. (2008). Evolution of the mycobacterial SigK regulon. *Journal of Bacteriology*, *190*(6), 1891–1899. http://doi.org/10.1128/JB.01452-07

Vidal, M., Gigoux, V., & Garbay, C. (2001). SH2 and SH3 domains as targets for anti-proliferative agents. *Critical Reviews in Oncology/Hematology*, *40*(2), 175–186.

Walsh, N. P., Alba, B. M., Bose, B., Gross, C. A., & Sauer, R. T. (2003). OMP peptide signals initiate the envelope-stress response by activating DegS protease via relief of inhibition mediated by its PDZ domain. *Cell*, *113*(1), 61–71.

Wang, X., Zhang, W., Zhou, H., Chen, G., Liu, W., & Kivisaar, M. (2019). An Extracytoplasmic Function Sigma Factor Controls Cellulose Utilization by Regulating the Expression of an Outer Membrane Protein in Cytophaga hutchinsonii. *Applied and Environmental Microbiology*, *85*(5), e02606–18. http://doi.org/10.1128/AEM.02606-18

Wecke, T., Halang, P., Staroń, A., Dufour, Y. S., Donohue, T. J., & Mascher, T. (2012). Extracytoplasmic function σ factors of the widely distributed group ECF41 contain a fused regulatory domain. *MicrobiologyOpen*, *1*(2), 194–213. http://doi.org/10.1002/mbo3.22

Wilken, C., Kitzing, K., Kurzbauer, R., Ehrmann, M., & Clausen, T. (2004). Crystal structure of the DegS stress sensor: How a PDZ domain recognizes misfolded protein and activates a protease. *Cell*, *117*(4), 483–494.

Wilson, M. J., & Lamont, I. L. (2006). Mutational analysis of an extracytoplasmic-function sigma factor to investigate its interactions with RNA polymerase and DNA. *Journal of Bacteriology*, *188*(5), 1935–1942. http://doi.org/10.1128/JB.188.5.1935-1942.2006

Wood, L. F., & Ohman, D. E. (2015). Cell wall stress activates expression of a novel stress response facilitator (SrfA) under σ22 (AlgT/U) control in Pseudomonas aeruginosa. *Microbiology*, *161*(1), 30–40. http://doi.org/10.1099/mic.0.081182-0

Woods, E. C., & McBride, S. M. (2017). Regulation of antimicrobial resistance by extracytoplasmic function (ECF) sigma factors. *Microbes and Infection*, *19*(4-5), 238–248. http://doi.org/10.1016/j.micinf.2017.01.007

Woods, E. C., Wetzel, D., Mukerjee, M., & McBride, S. M. (2018). Examination of the Clostridioides (Clostridium) difficile VanZ ortholog, CD1240. *Anaerobe*.

Xu, C., & Min, J. (2011). Structure and function of WD40 domain proteins. *Protein & Cell*, *2*(3), 202–214. http://doi.org/10.1007/s13238-011-1018-1

Yanamandra, S. S., Sarrafee, S. S., Anaya-Bergman, C., Jones, K., & Lewis, J. P. (2012). Role of the Porphyromonas gingivalis extracytoplasmic function sigma factor, SigH. *Molecular Oral Microbiology*, *27*(3), 202–219. http://doi.org/10.1111/j.2041-1014.2012.00643.x

Yang, L.-Y., Yang, L.-C., Gan, Y.-L., Wang, L., Zhao, W.-Z., He, Y.-Q., et al. (2018). Systematic Functional Analysis of Sigma (σ) Factors in the Phytopathogen Xanthomonas campestris Reveals Novel Roles in the Regulation of Virulence and Viability. *Frontiers in Microbiology*, *9*, 205. http://doi.org/10.3389/fmicb.2018.01749

Yeats, C., Rawlings, N. D., & Bateman, A. (2004). The PepSY domain: a regulator of peptidase activity in the microbial environment? *Trends in Biochemical Sciences*, *29*(4), 169–172. http://doi.org/10.1016/j.tibs.2004.02.004

Yoshimura, M., Asai, K., Sadaie, Y., & Yoshikawa, H. (2004). Interaction of Bacillus subtilis extracytoplasmic function (ECF) sigma factors with the N-terminal regions of their potential anti-sigma factors. *Microbiology*, *150*(3), 591–599. http://doi.org/10.1099/mic.0.26712-0

Yu, X., Lund, S. P., Greenwald, J. W., Records, A. H., Scott, R. A., Nettleton, D., et al. (2014). Transcriptional Analysis of the Global Regulatory Networks Active in Pseudomonas syringae during Leaf Colonization. *mBio*, *5*(5), e01683–14–e01683–14. http://doi.org/10.1128/mBio.01683-14
